# The *pos-1* 3ʹ untranslated region governs germline specification and proliferation to ensure reproductive robustness

**DOI:** 10.1101/2025.11.26.690476

**Authors:** Haik V. Varderesian, Juliet N. Utaegbulam, Hannah E. Brown, Beverly Ramirez, Melina Velcani, Sean P. Ryder

**Affiliations:** Department of Biochemistry and Molecular Biotechnology, University of Massachusetts Chan Medical School, Worcester, MA, USA

**Keywords:** RNA-binding protein, tandem zinc finger, nematode, oocyte, reproduction, fertilization

## Abstract

During fertilization, haploid gametes combine to form a zygote. The male (sperm) and female (oocyte) gametes contribute a similar amount of DNA, but the oocyte contributes nearly all the cytoplasm. Oocytes are loaded with maternal mRNAs thought to be essential for embryonic patterning after fertilization. A conserved suite of RNA-binding proteins (RBPs) regulates the spatiotemporal translation and stability of maternal mRNAs. POS-1 is a CCCH-type tandem zinc finger RBP expressed in fertilized *Caenorhabditis elegans* zygotes from maternally supplied mRNA. POS-1 accumulates in the posterior of the embryo where it promotes posterior cell fate. Here, we show that the *pos-1* 3ʹ untranslated region (UTR) is essential for POS-1 patterning and contributes to maximal reproductive fecundity. We engineered a *pos-1* mutant where most of the endogenous *pos-1* 3ʹUTR was removed using CRISPR genome editing. Our results show that the 3ʹUTR represses POS-1 expression in the maternal germline but increases POS-1 protein levels in embryos after fertilization. In a wild-type background, POS-1 repression via the 3ʹUTR has little impact on fertility. In a sensitized background, the deletion mutant has a complex pleiotropic phenotype where most adult homozygous progeny lack either one or both gonad arms. Most phenotypes become more penetrant at elevated temperature. Together, our results support an emerging model where the 3ʹUTRs of maternal transcripts, rather than being essential, contribute to reproductive robustness during stress.

## INTRODUCTION

According to a World Health Organization news report, infertility impacts one out of six people worldwide [1]. Infertility problems arise from defects in gametogenesis, mutations in the sperm or oocyte that impact embryo viability, and a variety of other causes [2, 3]. It is estimated that 15% of pregnancies end with miscarriage, defined as spontaneous loss of pregnancy prior to 24 weeks of gestation [4]. Environmental factors such as smoking, excessive alcohol consumption, and toxin exposure contribute to the miscarriage rate [5, 6]. Similarly, stressors such as a high fever in the first trimester can lead to miscarriage and an increased risk of birth defects such as cleft palate and congenital heart defects [7]. A thorough characterization of the molecular pathways that govern fertilization, early development, and embryonic robustness is necessary to mitigate the impact of these reproductive issues.

The nematode *Caenorhabditis elegans* is an effective model for studying reproduction. As with most animals, maternal mRNAs produced in the germline are used by the embryo after fertilization [8]. Proteins produced from these transcripts guide early developmental decisions such as axis polarization, segregation of germline and soma, and cell fate specification prior to the onset of zygotic transcription [9–13]. Consistent with this model, forward genetic screens identified numerous RNA-binding proteins (RBPs) essential for germline and early embryonic development [14–18]. Many of the factors involved are conserved from humans to worms [19]. In addition, the 3ʹ untranslated regions (UTRs) of maternal transcripts were shown in a series of reporter studies to be essential for patterned expression in the germline and embryos [20, 21].

Mutations of the human CCCH-type tandem zinc finger (TZF) gene ZFP36L2 have been linked to infertility due to zygotic arrest at day three post-fertilization [22]. Mutations in its murine homolog Zfp36l2 cause embryonic lethality between days two and three post fertilization with embryonic arrest at the two-cell stage [23]. There are multiple TZF family RBPs encoded in the *C. elegans* genome, many of which contribute to oocyte maturation and early embryonic patterning [16–18]. Here, we focused on *pos-1*, required for posterior cell fate specification in the embryo [16, 24].

Null mutation of the *pos-1* gene causes maternal effect embryonic lethality with terminal embryos that fail to specify intestine and germ cells while developing a disorganized region of pharyngeal tissue [16]. POS-1 is thought to direct repression of maternal transcripts in the posterior of early embryos through binding to their 3ʹUTR [24]. POS-1 binds with high affinity to a linear, partially degenerate sequence motif known as the POS-1 repression element (PRE: UAU2-3RDN1-3G), though presence of a PRE is not sufficient to confer POS-1-mediated repression in animals [25]. Notable targets of POS-1 repression include *glp-1* mRNA, a Notch receptor gene required for cell-to-cell signaling that drives anterior patterning decisions [24], and *neg-1*, a nuclear-localized protein expressed in anterior lineage cells that blocks mesoderm cell fates [26].

POS-1 protein is produced from maternal mRNA shortly after fertilization (**Fig. 1A-B**) [16]. It accumulates in the posterior of the one-cell zygote [16]. At the first cell division, POS-1 is asymmetrically inherited in the P1 blastomere. In four cell embryos, POS-1 is observed in the P2 (germline) and EMS (intestine, muscle) blastomeres. POS-1 is observable for a few more rounds of cell division, where it accumulates in the p-granules of germline lineage cells. The anti-correlated pattern of POS-1 with NEG-1 and GLP-1 suggests that it functions to repress expression of these genes in the posterior [16, 26]. Transgenic reporter studies in live embryos confirmed that POS-1 represses both genes through sequence specific interaction with their 3ʹUTRs [26, 27].

**Fig. 1.**
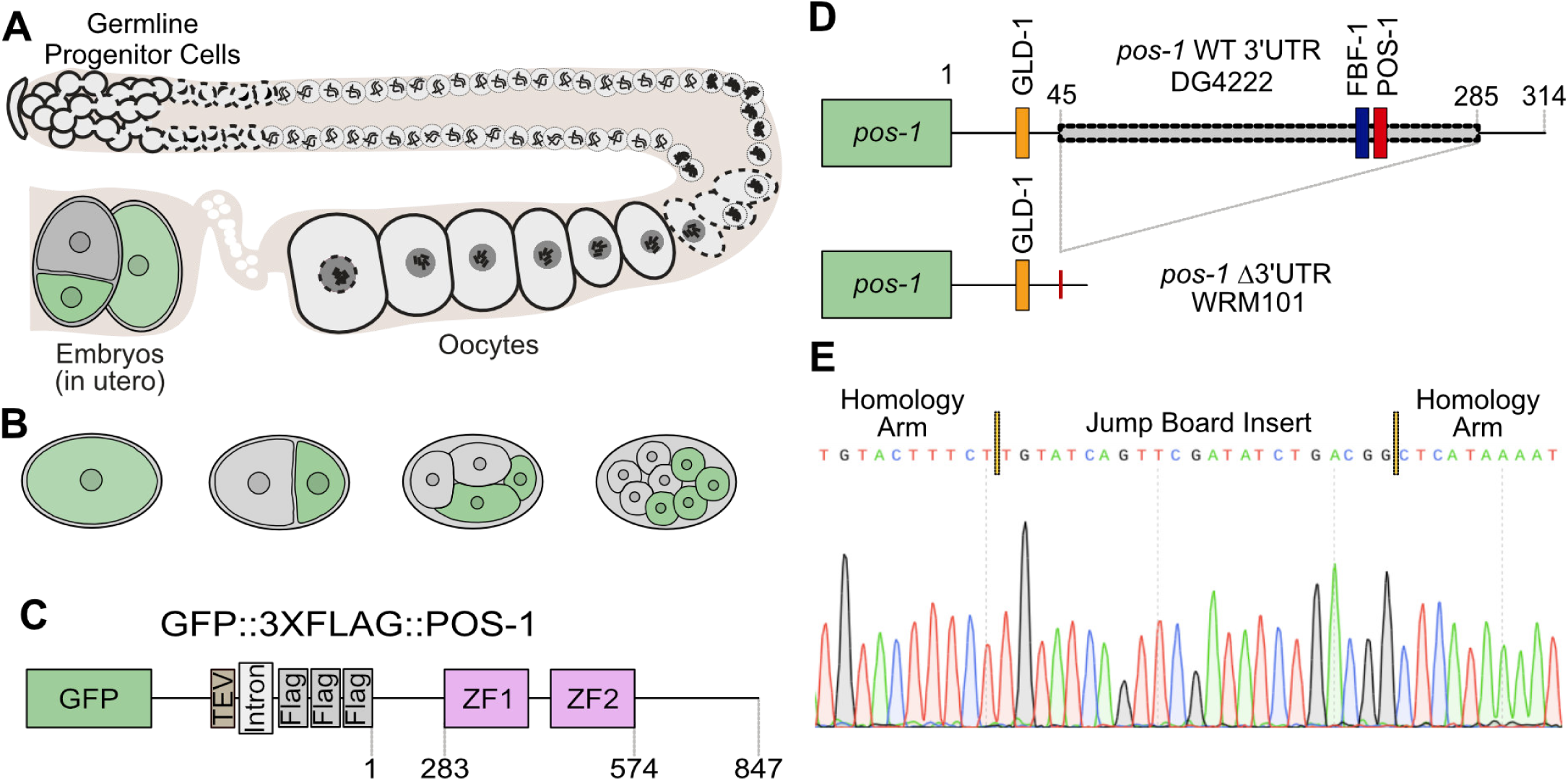
POS-1 is a germline RNA binding protein required for posterior cell fate specification in the early embryo [16]. **A.** Diagram of a *C. elegans* hermaphrodite gonad. **B.** The pattern of POS-1 expression pattern in early embryos is shown in green. In older embryos POS-1 is eventually restricted to the P-lineage [16]. **C.** In the genetic background used in this study (DG4222), the endogenous *pos-1* locus, which contains two CCCH-type zinc finger domains (labeled in pink), has an N-terminal *gfp::tev::3xflag* tag [33]. **D.** The ΔUTR allele replaces 241 of the 314 nucleotides of the *pos-1* 3ʹUTR with a 23 nucleotide jump board sequence [34]. GLD-1, FBF-1, and POS-1 binding motifs are shown. **E.** Chromatogram of the mutant confirming the deletion and jump board insertion. A portion of each flanking homology arm are shown on either side of the incorporated sequence.

Spatial and temporal patterning of POS-1 is thought to involve both post-translational and post-transcriptional regulatory mechanisms. Contributing factors that have been identified include RNA-binding proteins and Polo-like kinases [28, 29]. To test the hypothesis that *pos-1* mRNA is regulated at the post-transcriptional level, we used CRISPR-Cas9 genome editing to generate a large deletion allele in the endogenous *pos-1* 3ʹUTR (**Fig. 1C-D**). Worms that harbor this allele are viable as homozygotes. However, in a sensitized background, they display a complex pleiotropic phenotype that is distinct from the *pos-1* null mutant phenotype. Most progeny are sterile with absent or undersized gonads that do not undergo oogenesis. Others show defective in oocyte development or maturation. Most of the phenotypes become more penetrant at higher temperatures. Fertile animals show high POS-1 expression throughout the germline but reduced POS-1 expression in embryos compared to wild-type. In this regard, the *pos-1* 3ʹUTR mutant behaves similar to a 3ʹUTR deletion mutation recently characterized in the maternal *mex-3* gene [30, 31]—another maternally-supplied RBP that determines cell fate [14]—though the reproductive phenotypes are different. This parallel adds support to an emerging model where the 3ʹUTRs of maternal mRNAs contribute to silencing in the maternal germline and buffering expression in embryos, especially during periods of stress.

## RESULTS

### Mutating the pos-1 3′UTR using CRISPR-Cas9 genome engineering

We set out to test whether the endogenous *pos-1* 3ʹUTR plays a role in reproductive fecundity. Following the method of Ghanta et al., we used CRISPR-Cas9 genome engineering to remove the majority of the *pos-1* 3ʹUTR [32] in both the wild-type N2 (Bristol) strain and in strain DG4222 (**Table 1**), which includes an N-terminal *gfp::tev::3xflag* fusion tag knocked into the endogenous *pos-1* locus [33] (**Fig. 1C**). We used a repair template that includes a jump board sequence containing a non-native guide RNA (gRNA) site and protospacer adjacent motif (PAM) site to simplify further editing [34]. The resulting alleles are identical in both genetic backgrounds, removing 239 of the 314 nucleotides found in the *pos-1* 3’UTR and replacing them with the 23 nucleotide jump board (**Fig. 1D**). The deletion removes several motifs predicted to be recognized by RNA-binding proteins including OMA-1/2, FBF-1, as well as POS-1 itself [35–37]. Both alleles (denoted as GFP-ΔUTR and ΔUTR hereafter) were confirmed by PCR, gel electrophoresis, and sequencing (**Fig. 1E**). Mutant worms harboring the 3ʹUTR deletion can be propagated as homozygotes under standard growth conditions in both genetic backgrounds. (NGM Agar, *Escherichia coli* OP50 food, 20°C) [38].

**Table 1.**

| Strain ID | Genotype | Technology |
| --- | --- | --- |
| Published Strains |  |  |
| N2 | *wild isolate* | *n/a* |
| DG4222 | *pos-1(tn1730[gfp::tev::3xflag::pos-1]) V* | CRISPR |
| Strains Produced for this Study |  |  |
| WRM85 | *pgl-1(spr20[mCherry::pgl-1]) IV* | CRISPR |
| WRM101 | *pos-1(spr28[tn1730\*(gfp::tev::3xFlag::pos-1::ΔUTR)]) V* | CRISPR |
| WRM102 | *pos-1(spr29 [ΔUTR]) V* | CRISPR |
| WRM104 | *pgl-1(spr20[mCherry::pgl-1]) IV; pos-1(spr28[tn1730\*(gfp::tev::3xFlag::pos-1::ΔUTR)]) V* | CRISPR |
| WRM105 | *pgl-1(spr20[mCherry::pgl-1]) IV; pos-1(tn1730[gfp::tev::3xflag::pos-1]) V* | CRISPR |

### Deleting the pos-1 3′UTR impacts reproductive fecundity

We set out to quantify the reproductive health of untagged ΔUTR worms compared to the parent strain (N2: WT UTR hereafter). We also assessed the fertility of each worm’s F1 offspring. To do so, we measured the brood size (total embryos laid), hatch rate (hatched embryos / total brood), and sterility rate (number of sterile progeny / total viable hermaphrodite progeny) of ΔUTR worms compared to WT UTR controls. All three parameters were determined at two culture conditions—20°C, the optimal temperature for *C. elegans* growth [38], and 25°, a mild stress condition that can enhance reproductive phenotypes [39].

The average brood produced by untagged ΔUTR hermaphrodites is similar to N2 worms at 20°C (Average Brood_ΔUTR_ = 245 ± 14, Average Brood_N2_ = 250 ± 9, p_adj_ = 1.0) and 25°C (Average Brood_ΔUTR_ = 127 ± 7, Average Brood_N2_ = 157 ± 7, p_adj_ = 0.25, **Fig. 2A**). Similarly, the average hatch rate between WT N2 and the untagged *pos-1* Δ3′UTR strain is indistinguishable at both temperatures. One hundred percent of the brood of N2 hermaphrodites hatched under standard and mild stress temperature (N2; Average Hatch_WT-20°C_ = 100% ± 0% and Average Hatch_WT-25°C_ = 100% ± 0%, p_adj_ = 1.0). The ΔUTR strain had a minor reduction in hatchings from total brood at both temperatures, but neither was statistically significant (untagged ΔUTR; Average Hatch_ΔUTR-20°C_ = 99.4% ± 0.5%, and Average Hatch_ΔUTR-25°C_ = 95% ± 3%, p_adj_ = 0.39, **Fig. 2B**). These data indicate that mutating the *pos-1* 3′UTR has an insignificant impact on the brood size and hatch rate of *C. elegans* embryos in an otherwise wild-type background.

**Fig. 2.**
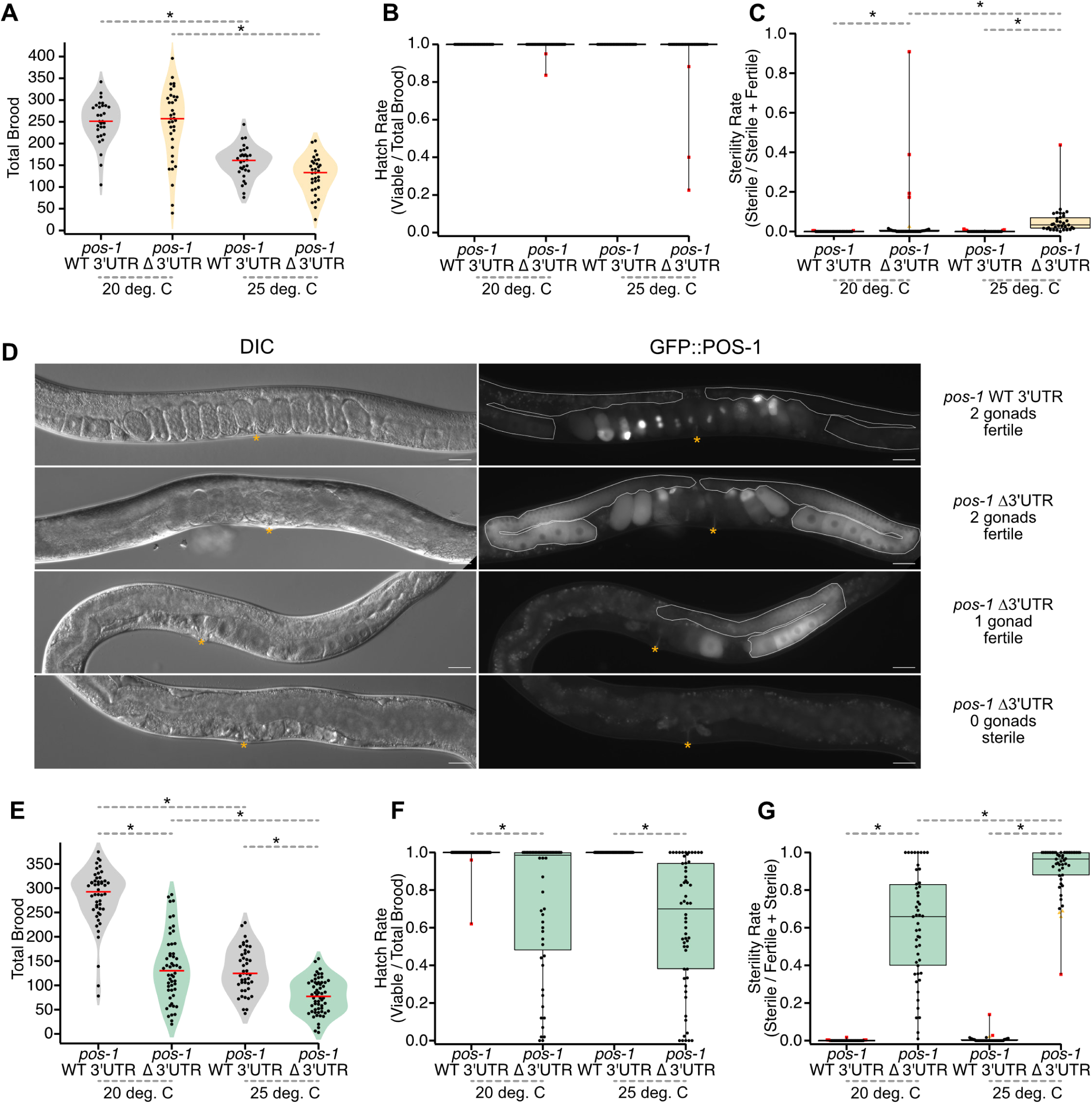
The *pos-1* 3ʹUTR contributes to reproductive robustness and germline formation**. A.** Violin plots of total brood for WT UTR (gray) and ΔUTR (orange) strains. The red lines indicate the median. Each point represents the total number of embryos produced by a single hermaphrodite. Asterisks indicate statistical significance in a one-way ANOVA with Bonferroni correction for multiple hypothesis testing. **B.** Box-and-whisker plot representing the hatch rate (viable / total brood) of embryos from panel A. Each dot represents the hatch rate of embryos from a single hermaphrodite. Red points indicate far outliers in a Tukey analysis (far outliers are at least 3X larger or smaller than the inner quartile range). **C.** Box-and-whisker plot depicting the sterility rate (sterile / sterile + fertile) of viable worms from panel B once they reach adulthood. Each dot represents the ratio of sterile F1 hermaphrodites to total viable hermaphrodites. Orange points indicate outliers in a Tukey analysis (1.5X larger or smaller than the inner quartile range. Far outliers and statistical significance are represented as in panel A. **D.** 20X DIC and GFP images of GFP-WT UTR and GFP-ΔUTR mutants at 20°C. The yellow asterisk indicates the vulva, and the dotted white line represents each gonad arm, if present. Scale bars: 30 μm. **E.** Violin plot of total brood comparing between GFP-WT UTR (gray) and the GFP-ΔUTR (green) at 20°C and 25°C. Median bars, statistical significance, and individual points are represented as in panel A. **F.** Box-and-whisker plot depicting the hatch rate of embryos from panel E. Each dot represents the hatch rate of embryos produced from a single worm as in panel B. **G.** Box-and-whisker plot depicting the sterility rate of viable progeny from panel F at adulthood. The representation is as described for panel C.

By contrast, some of the viable progeny of ΔUTR worms are sterile at both temperatures. At 20°C, approximately half of the worms produced at least some sterile F1 progeny (n=17/35). Four animals had relatively high sterility rates (red dots, **Fig 2C**), including one animal that produced 91% sterile progeny. WT UTR animals also produced some sterile progeny at 20°C, but the sterility rate was much lower (n=4/30, max sterility rate = 0.42%, p_adj_ = 0.015). At 25°C, nearly all ΔUTR animals produced sterile progeny (n=32/33), with a maximum sterility rate of 44%. WT UTR animals had a much lower sterility rate with a maximum sterility rate of 1.2% (n=5/30, p_adj_ < 6.04e-10). The data show that loss of the *pos-1* 3ʹUTR causes a partially penetrant F1 sterility phenotype that is exacerbated by elevated temperature.

In the *gfp::tev::3xflag* tagged background, the ΔUTR phenotypes appear to be much stronger. While observing homozygous GFP-ΔUTR mutants on plates, we noticed that a large fraction of adult animals appears to lack gonads and do not contain embryos (**Fig. 2D**). These worms are easily distinguished from fertile animals under a light microscope by the appearance of a “dark gut” morphology and the absence of embryos [40]. We also noted an accumulation of dead embryos on the plate.

To quantify the fecundity of fertile GFP-ΔUTR worms, we repeated the brood size, hatch rate, and sterility rate measurements, comparing GFP-ΔUTR mutants to GFP-WT UTR (DG4222, **Table 1**) controls. Fertile GFP-ΔUTR worms had a lower total brood than GFP-WT UTR controls at both temperatures (**Fig. 2E**). At 20°C, GFP-WT UTR worms produced an average of 281 ± 9 embryos. By contrast, GFP-ΔUTR worms produced an average of 135 ± 10 total embryos (Fold Effect_ΔUTR/WT UTR_=0.48, p_adj_<1e-8). At 25°C, the total brood is further reduced in both strains. GFP-WT UTR worms produced an average of 130 ± 7 total embryos, while GFP-ΔUTR worms produced 78 ± 5 (Fold Effect_ΔUTR/WT UTR_=0.6, p_adj_=5.6e-7).

The fraction of hatching embryos was also impacted in the GFP-ΔUTR mutant at both temperatures. At 20°C, the fraction of GFP-WT UTR embryos that hatch is 0.99 ± 0.008, while the fraction of GFP-ΔUTR embryos that hatch is 0.73 ± 0.05 (**Fig. 2F**, p_adj_=2e-5). Similarly, at 25°C, all the GFP-WT UTR embryos hatched, while only 0.62 ± 0.04 GFP-ΔUTR embryos hatched (p_adj_=3e-10). As such, adult GFP-WT UTR animals produce an average of 278 viable progeny at 20°C and 130 viable progeny at 25°C, while GFP-ΔUTR adults produce an average of 99 viable progeny at 20°C and just 48 at 25°C. This reduction in fecundity encompasses both reduced embryo production and reduced embryo viability, suggesting problems in the germline as well as the embryo.

In addition to reduced brood size and hatch rate, F1 sterility is a common phenotype in GFP-ΔUTR homozygotes (**Fig. 2G**). At 20°C, GFP-WT UTR animals produced at least some sterile progeny 15% of the time (n=7/48), but the average sterility rate was very low (0.1%). In contrast, all GFP-ΔUTR animals produced at least some sterile progeny (n=49/49, p_adj_ <1e-10), and the average sterility rate was much higher (68%, p_adj_ < 1e-10), with all sterile animals lacking both gonad arms. At 25°C, the fraction of GFP-WT UTR worms producing sterile animals increased (GFP-WT UTR = 34%, n=15/44), but the sterility rate remained low (0.7%). In GFP-ΔUTR animals, once again all worms produced at least some sterile progeny (n=52/52, padj < 1e-10), but the average sterility rate increased to 91% (p_adj_ < 1e-10). The results suggest a defect in germline development or germline maintenance in mutant F1 animals compared to genotype-matched controls.

### Additional defects in pos-1 3ʹUTR mutants

We observed two additional low penetrance phenotypes in fertile GFP-ΔUTR adults compared to GFP-WT UTR controls. The first is the presence of immature oocytes with multiple nuclei (polynucleated oocytes, **Fig. 3A, B**), suggesting errors in cytokinesis, apoptosis, or gamete fusion during oogenesis [41]. GFP-WT UTR worms do not form polynucleated oocytes (20°C, n=0/68; 25°C, n=0/84), but the GFP-ΔUTR mutants show this phenotype at both growth temperatures (20°C, n=11/95 [11.6%], p_adj_ΔUTR20vsWTUTR20_=0.022; 25°C, n=7/80 [8.75%], p_adj_ΔUTR25vsWTUTR25_=0.034). We also observed the presence of unusually oblong oocytes, wider than they are tall (**Fig. 3C, D**). Oblong oocytes were present in both GFP-ΔUTR and GFP-WT UTR control animals, but they were most prevalent in the GFP-ΔUTR mutants at elevated temperature. At 20°C, oblong oocytes are found in 2.9% (20°C, 2/68) of GFP-WT UTR control animals and 7.4% of GFP-ΔUTR animals (20°C, n=7/95), a nonsignificant difference (p_adj_ΔUTR20vsWTUTR20_=1). At 25°C, the phenotype becomes more prevalent, with 51.3% of GFP-ΔUTR animals (25°C, 41/80) and 16.7% (25°C, 14/84) of GFP-WT UTR animals showing oblong oocytes. This increase is statistically significant (p_adj_ΔUTR25vsWTUTR25_=1.7e-5). The results are consistent with partially penetrant oogenesis and/or oocyte maturation phenotypes in the fertile subset of GFP-ΔUTR mutant worms compared to genotype matched control. Some phenotypes depend on the temperature of growth, but the impact of temperature appears unique for each phenotype observed.

**Fig. 3.**
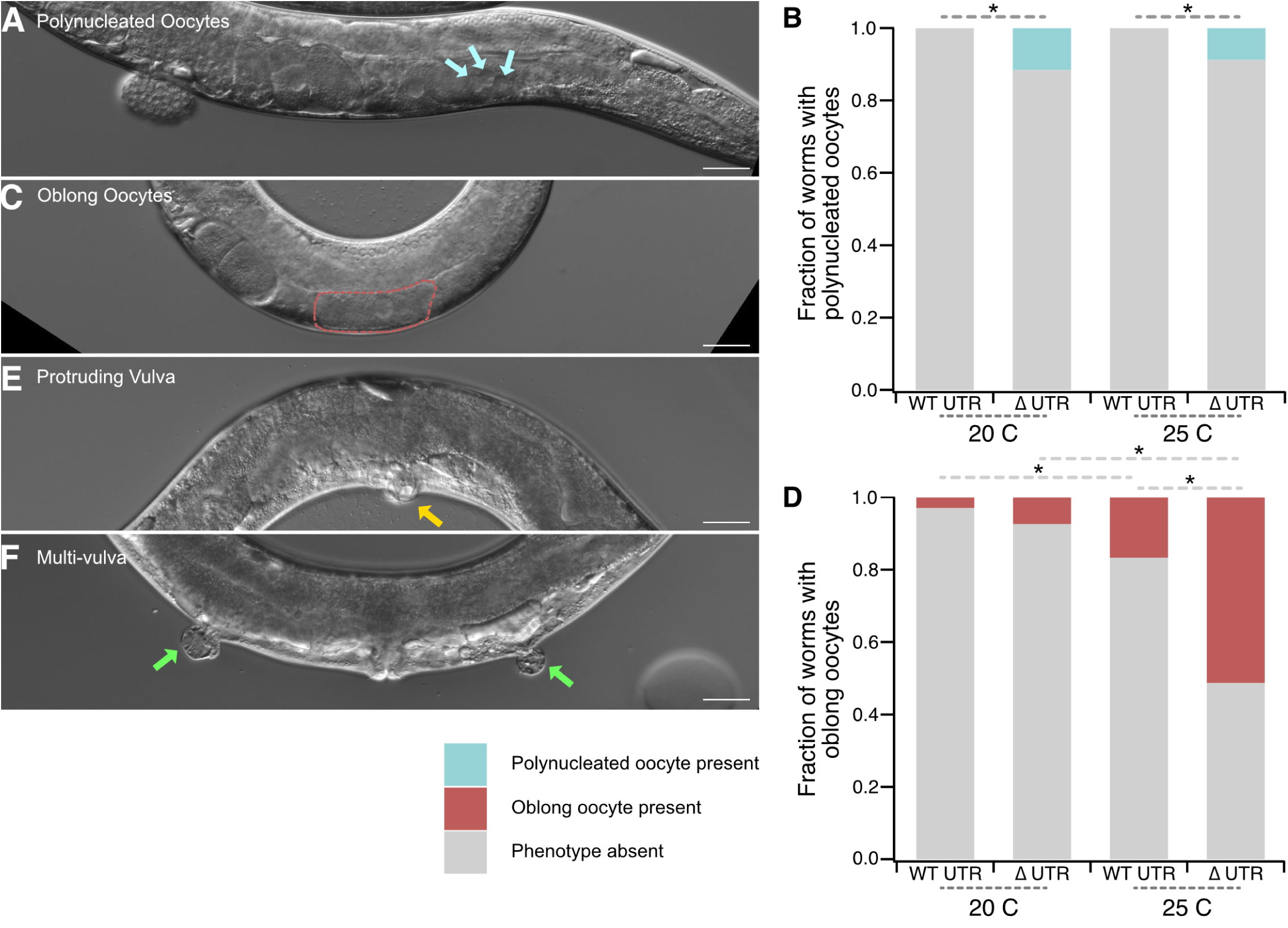
Germline phenotypes in the *pos-1* 3ʹΔUTR mutant. **A.** DIC image of a GFP-ΔUTR mutant with a polynucleated oocyte (cyan arrows–each points to a different nucleus within the same oocyte). **B.** Stacked bar graph showing the fraction of worms with polynucleated oocytes (cyan) or normal oocytes (gray) in germline images. Asterisks indicate statistical significance in a Pearson’s chi square test with Bonferroni correction for multiple hypothesis testing. **C.** DIC image of a GFP-ΔUTR oblong oocyte phenotype. The length of most proximal oblong oocyte is marked in red. **D.** Stacked bar graph showing fraction of worms with oblong oocytes (red) or normal oocytes (gray) in germline images. Statistical significance is denoted as in B. **E.** DIC image of a representative GFP-ΔUTR mutant with protruding vulva phenotype (pvul, yellow arrow). **F.** DIC image of a GFP-ΔUTR mutant with a multiple vulva (muv) phenotype. Each abnormal vulva location is marked with a green arrow. The scale bars represent 30 μm in all images.

In addition to these, we observed a variety of incompletely penetrant somatic phenotypes including protruding vulva and multivulva animals (**Fig. 3E, F**). These phenotypes only occurred in GFP-ΔUTR mutants. We did not quantify their penetrance or temperature dependence. We also note that vulval morphogenesis defects may be the root cause of our difficulty in outcrossing the ΔUTR allele.

### Whole genome sequencing of ΔUTR strains

We were surprised that the phenotypes appeared much stronger in the GFP-tagged background than in the untagged mutants. We also noted that some phenotypes manifest at low penetrance in the GFP-WT UTR control strain, including low penetrance sterility (**Fig. 2G**) and oblong oocyte (**Fig. 3C**) phenotypes. Homozygous GFP-WT UTR animals were outcrossed three times prior to CRISPR mutagenesis. However, the resultant GFP-ΔUTR mutant animals mate inefficiently, and as such could not be outcrossed. Attempted matings resulted in animals dying due to bursting through the vulva, potentially due to the vulval morphogenesis defects described above. As such, it remained possible that background mutations arising during CRISPR mutagenesis could be responsible for the strong phenotypes we observed.

To identify potential confounds in the GFP-tagged genetic background, we performed whole genome sequencing on the ΔUTR strain (WRM102), the GFP-ΔUTR strain (WRM101), and the GFP-WT UTR strain (DG4222). The results confirmed the presence of the ΔUTR allele in the *pos-1* locus of both the ΔUTR and GFP-ΔUTR strains (**Supplemental Data Set 1**, **Supplemental Fig. 1A**). The data also identify 105 exonic single nucleotide polymorphisms (SNPs) in the GFP-ΔUTR animals with an allele frequency of one compared to the reference genome (**Supplemental Fig. 1B**). All 105 are also found in the GFP-WT UTR background, although one SNP—a nonsynonymous substitution in the unstudied *Y22D7AR.2* gene—is found at lower allele frequency. Fifty of the 105 SNPs are also found in the untagged ΔUTR mutant. The data also revealed 32 exonic indels in the GFP-ΔUTR strain (**Supplemental Fig. 1C**). All but one are also found in the GFP-WT UTR background. The sole exception is found in the untagged ΔUTR background, which shares 26 of the 31 exonic indels with the GFP-ΔUTR strain, and does not display strong phenotypes. Because none of the SNPs or indels are unique to the GFP-ΔUTR strain, it is unlikely that the phenotypes observed are caused solely by a background mutation. However, the differences in phenotype between both GFP-tagged strains and the untagged ΔUTR strain suggest that the GFP-tagged background is sensitized, leading to higher penetrance phenotypes. This may be the result of the 55 exonic SNPs and 6 exonic indels found in the GFP-tagged background but not in the untagged ΔUTR strain. Alternatively, it is possible that the *gfp::tev::3xflag* tag itself contributes to the stronger ΔUTR phenotype. Either way, the data show that the GFP-WT UTR strain is a suitable control for the GFP-ΔUTR strain.

### Mutation of the endogenous pos-1 3ʹUTR causes strong germline expression

To define the pattern of GFP::POS-1 expression, GFP-WT UTR and GFP-ΔUTR young adult worms were imaged under a fluorescence microscope at both temperatures. Though both sterile and fertile progeny of GFP-ΔUTR worms were imaged (**Fig. 2D**), the comparisons described below focus on the subset of ΔUTR animals that appear fertile and contain two intact gonads. In GFP-WT UTR worms, the GFP::POS-1 expression pattern was restricted to young embryos (**Fig. 4A**) as expected [16]. By contrast, GFP::POS-1 protein expression is strongly dysregulated in fertile GFP-ΔUTR mutants that contain intact gonads. High GFP::POS-1 expression is observed in the germline pachytene region (Bin 11, Fold Increase _ΔUTR/WT_ = 5.7, p_adj_ < 0.00001, **Fig. 4B**), in the loop region (Bin 13, Fold Increase _ΔUTR/WT_ = 4.1, p_adj_ < 0.00001), and in oocytes (Bin 18, Fold Increase _ΔUTR/WT_ = 5.2, p_adj_ < 0.00001). A smaller increase is observed in mitotic germline progenitor cells at the distal end of the germline (Bin 0, Fold Increase _ΔUTR/WT_ = 1.4, p_adj_ = 0.00001). The results suggest that the *pos-1* 3ʹUTR is essential for repressing POS-1 protein production in the maternal germline.

**Fig. 4.**
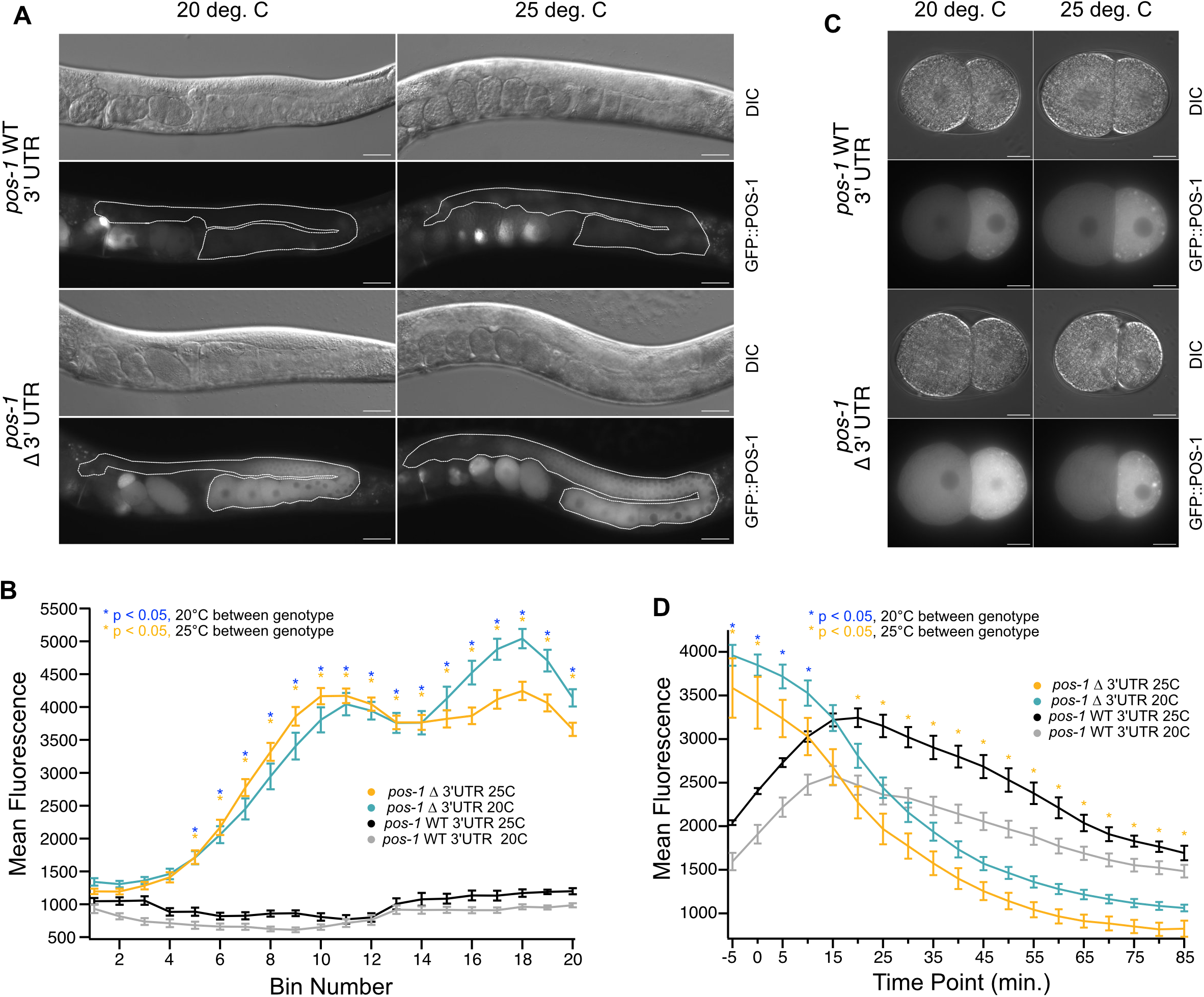
POS-1 protein expression is affected in the germline and embryos of 3ʹΔUTR mutants. **A.** DIC and GFP images of GFP-WT UTR and GFP-ΔUTR germlines (white outline) revealing POS-1 protein overexpression differences between genotype and condition. The scale bars represent 30 μm. **B.** Plot of germline fluorescence intensity as a function of distance from the distal end, with average GFP::POS-1 fluorescence averaged across twenty equal width bins representing the full length of the germline. The points represent the average fluorescence intensity from multiple images within a given genotype and condition; the error bars represent the standard error of the mean. Statistical significance is shown with asterisks as labeled in the key. P values were calculated with a one-way ANOVA with Bonferroni correction for multiple hypothesis testing. **C.** DIC and GFP image of GFP-WT UTR and GFP-ΔUTR embryo at the two-cell stage. The posterior pole is oriented to the right. Scale bars: 10 μm. **D.** Average GFP::POS-1 fluorescence intensity across the entire embryo is plotted as a function of time from embryogenesis movies collected in five-minute intervals over 90 minutes of early embryonic development. Statistical significance is represented as in panel B.

Next, we sought to assess the impact of temperature stress on this pattern of expression. We repeated our imaging experiments with worms cultured at 25°C. As expected, increasing the temperature caused no change in the pattern of expression in GFP-WT UTR worms—germline expression is not observed at either temperature. GFP-ΔUTR mutants continued to display strong fluorescence throughout the germline (**Fig. 4B**). There are no statistically significant differences within a genotype as temperature increases.

We noted that a subset of fertile GFP-ΔUTR young adults had only one gonad arm. For GFP-WT UTR worms cultured at 20°C or 25°C, none of the imaged worms had fewer than two gonad arms (20°C, n=0/68; 25°C, n=0/84). For ΔUTR animals cultured at 20°C, 11.4% had just one gonad arm (20°C, n=10/95, p_adj_=0.00051, **Fig. 2D**). Surprisingly, this phenotype appears to be absent at 25°C. None of the fertile GFP-ΔUTR worms had just one gonad arm (25°C, n=0/80, p_adj_ΔUTR25vs20_=0.00024). This change corresponds with an increase in sterility (**Fig. 2G**), suggesting that at elevated temperature, more animals have zero gonads. The data indicate that *pos-1* 3ʹUTR mutants produce a mixed population of progeny where some animals lack one or both gonad arms, and that elevated temperature alters the distribution.

### GFP::POS-1 abundance rapidly decreases in Δ3′UTR mutant embryos post-fertilization

To determine whether GFP::POS-1 protein abundance remains high after fertilization, we recorded images of embryogenesis under a fluorescence microscope as a function of time at five-minute intervals (**Fig. 4C**, **Supplemental Movie 1, Supplemental Movie 2**). For each frame, the total fluorescence intensity of GFP::POS-1 within the embryo was measured. The first frame in which cellular division is observed was arbitrarily set to time zero to facilitate comparisons between embryos. Total fluorescence values were averaged at each time point over multiple movies (n = 7-10) within each genotype and growth condition (20°C or 25°C).

At early time points (first 15 minutes), GFP-ΔUTR mutant embryos have higher average POS-1 fluorescence values than similarly aged GFP-WT UTR embryos (p_adj_ < 3.5e-6, **Fig. 4D**). At t = 0 and 20°C, ΔUTR mutant embryos have a mean pixel intensity twice that of WT UTR embryos of the same age (t=0 minutes, 20°C, Fold Increase _ΔUTR/WT_ = 2.0, p_adj_ <1e-10). The fold increase comparing the GFP-ΔUTR to the GFP-WT UTR animals at 25°C is not as high, but remains statistically significant (t=0 minutes, 25°C, Fold Increase _ΔUTR/WT_ = 1.5, p_adj_ = 0.01, **Fig. 4D**). The data indicate that the POS-1 protein found within mutant embryos is higher in abundance compared to WT expression at both temperature conditions immediately following fertilization.

The pattern of GFP::POS-1 accumulation does not appear to be affected by the 3ʹUTR mutation. In two-cell embryos, most of the GFP::POS-1 protein accumulates in the P1 cell (posterior) in both genotypes at both temperatures (**Fig. 4C**). However, the total abundance of GFP::POS-1 appears to be impacted by both genotype and temperature. Though GFP::POS-1 is initially much higher in mutant embryos than control embryos, it declines to a level lower than GFP-WT UTR embryos over the course of early embryogenesis (**Fig. 4D**). The decay of GFP::POS-1 in GFP-ΔUTR embryos follows a sigmoidal curve. The time with half maximal concentration of GFP::POS-1 (incorporating the rates of both new GFP::POS-1 translation and decay) is t = 20.3 ± 3 minutes at 20°C and t = 15 ± 8 minutes at 25°C. By contrast, GFP::POS-1 abundance increases in GFP-WT UTR animals following a modified Gaussian curve with a peak at t = 12 ± 2.4 minutes at 20°C and t = 16.3 ± 1.2 minutes at 25°C. After 20 minutes, GFP::POS-1 levels are lower in GFP-ΔUTR_25°C_ embryos than in GFP-WT UTR_25°C_ embryos, and remain lower for the duration of the recording (t=85 min, 25°C, Fold Decrease _ΔUTR/WT_ = 0.49, p_adj_ = 0.009). The trend is similar at 20°C, but the apparent differences are not statistically significant (p_adj_ > 0.05). Together with our previous observations, the data suggest a model where the *pos-1* 3ʹUTR represses POS-1 expression in the germline but enhances POS-1 expression after fertilization. Temperature also appears to impact GFP::POS-1 differently in GFP-ΔUTR embryos compared to GFP-WT UTR embryos.

### Embryonic germ cell specification defects in the pos-1 3ʹUTR mutant

The frequent absence of gonad arms in the GFP-ΔUTR mutant could be caused by germ cell misspecification during embryogenesis, failure of germline proliferation during larval growth, or both. PGL-1 is a core component of germ granules, found exclusively in germline lineage cells, and makes a convenient marker for germ cell differentiation in embryos and germline morphology in adults. In early embryos, PGL-1 is found in the P-lineage cell (P1-P4) [42]. Around three hours post fertilization, the P4 cell divides to produce both the Z2 and Z3 cell, the primordial germ cells (PGCs) that ultimately form the germline in both gonads. The Z2 and Z3 cells remain in a quiescent state until the L1 larval stage, when they begin to divide again.

To test our hypotheses, we crossed the GFP-ΔUTR strain and the GFP-WT UTR control strain with worms expressing mCherry::PGL-1 from the endogenous *pgl-1* locus (WRM85). The resultant strains (WRM104 and WRM105, **Table 1**) have similar brood size, hatch rate, and sterility rate compared to genotype-matched strains lacking mCherry::PGL-1, suggesting that the mCherry tag on PGL-1 does not modify the ΔUTR phenotypes (**Fig. 5A-C**).

**Fig. 5.**
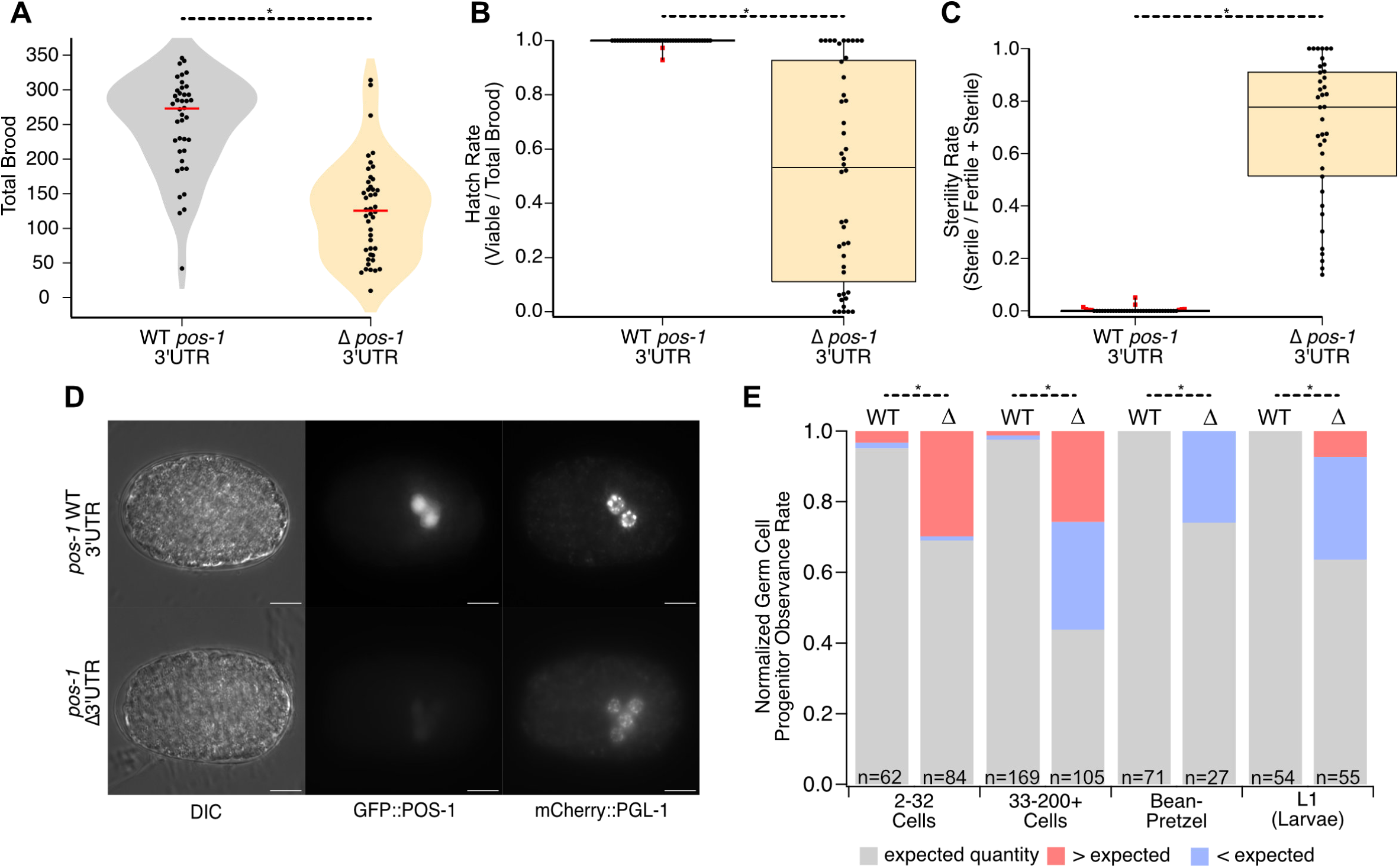
Germline progenitor specification is aberrant in *pos-1* 3ʹΔUTR mutants. **A.** Brood size of *mCherry::pgl-1* marked GFP-WT UTR and GFP-ΔUTR strains. The representations are as in Figure 2A. **B.** Hatch rate of the embryos from panel A. The representation is as in Figure 2B. **C.** Sterility rate of the viable progeny from panel B. The graph is represented as in Figure 2C. **D.** DIC, GFP, and mCherry images of embryos from GFP-WT UTR or GFP-ΔUTR animals tagged with mCherry::PGL-1 showing an example of increased germline progenitor cells in embryos in the 3ʹUTR deletion mutant. Scale Bars: 20 μm. **E.** Fraction of animals with less than (blue), equal to (gray), or more than (red) the expected number of germline progenitor cells in embryo images binned by developmental stage. Statistically significant differences from a Pearson’s Chi Square test with Bonferroni correction are marked with asterisks.

GFP-WT UTR and GFP-ΔUTR worms harboring the mCherry::PGL-1 marker were grown under standard conditions, and fertile young adults were selected for dissection and embryo imaging with a fluorescence microscope (**Figure 5D**). The images were grouped by genotype and by age—estimated from the cell count or morphological appearance—and binned into three groups: (1) early embryos from the 2–32 cell stage, where we expect exactly one mCherry::PGL-1 positive cell, (2) 33–200+ intermediate age embryos where both Z2 and Z3 should score positive for mCherry::PGL-1, and (3) older gastrulating embryos in the “bean” through “pretzel” stages that should also contain two mCherry::PGL-1 cells [42]. The quantity of mCherry positive cells was determined per embryo in each group for both genotypes. We also determined the number of germ cells in synchronized L1 larvae that were recovered by bleaching and allowing embryos to hatch overnight in the absence of food. L1 larvae were randomly selected for imaging shortly after plating.

Most of the embryos from GFP-WT UTR adults in the 2-32 cell stage had the expected number of mCherry positive cells (91%, one cell expected, n=58/64, **Fig. 5E**). In contrast, just 69% (n=58/84, p_adj_ = 0.00028) of GFP-ΔUTR embryos in this stage had the expected number of cells. Nearly 30% of GFP-ΔUTR mutants had more than one cell, with the highest single-embryo quantity at seven (**Fig. 5D**). This phenotype contrasts with *pos-1* null embryos, which fail to specify germ cells [16]. In the 33-200+ cell embryos, we observed something different. Again, nearly all the GFP-WT UTR embryos had the expected number of cells (98%, two cells expected, n=165/169). However, only 44% of GFP-ΔUTR embryos at this age had two cells (n=46/105, p_adj_ = <1e-10). The remaining embryos were split between those with fewer than expected germ cells (30%, 0–1 mCherry positive cells, n=33/105) and those with more than expected germ cells (26%, >2 mCherry positive cells, n=26/105).

As embryos aged, the distribution changed again. By the bean–pretzel stage, all the GFP-WT UTR embryos had the two expected PGCs (n=71), while 74% of GFP-ΔUTR embryos at this stage had the expected amount (two cells expected, n=20/27, p_adj_ = 8.5e-6). We note that it was relatively difficult to recover embryos of this age, presumably due to arrest or death prior to gastrulation in some mutant embryos, consistent with the hatch rate data (**Fig. 2G**). Consistent with that idea, we observed that all the remaining embryos had fewer than the expected number of germ cells (0–1), and none had more than expected (**Fig. 5E**). This suggests that embryos that specified extra germ cells failed embryogenesis before entering gastrulation, while those that specified 0-2 germ cells continued to develop. Interestingly, 62% of the L1 larvae that hatched from the GFP-ΔUTR mutant embryos contained the expected two germ cell progenitors (n=35/55) while all the GFP-WT UTR larvae had the expected number (n=54/54, p_adj_ = 6.0e-6). Most of the remaining larvae had fewer than two germ cells (29%, n=16/55), but we note that a few had more than expected (7%, n=4/55). These animals could represent embryos that successfully hatched after mis-specifying extra germ cells—contrary to our hypothesis—or they could be animals that exited quiescence to initiate germline proliferation early.

### Germline proliferation defects in the pos-1 3ʹUTR mutant

In *C. elegans*, there are four larval stages prior to adulthood. During each of these four stages, the two germ cell progenitors (Z2 & Z3) proliferate, eventually forming the characteristic U-shaped gonad arms [43]. To monitor germ cell proliferation as a function of developmental stage, we synchronized mCherry::PGL-1 expressing GFP-ΔUTR and GFP-WT UTR animals and collected images across development (**Fig. 6A**). The length of the germline was determined from mCherry::PGL-1 positive images using the segmented line tool in Fiji (ImageJ).

**Fig. 6.**
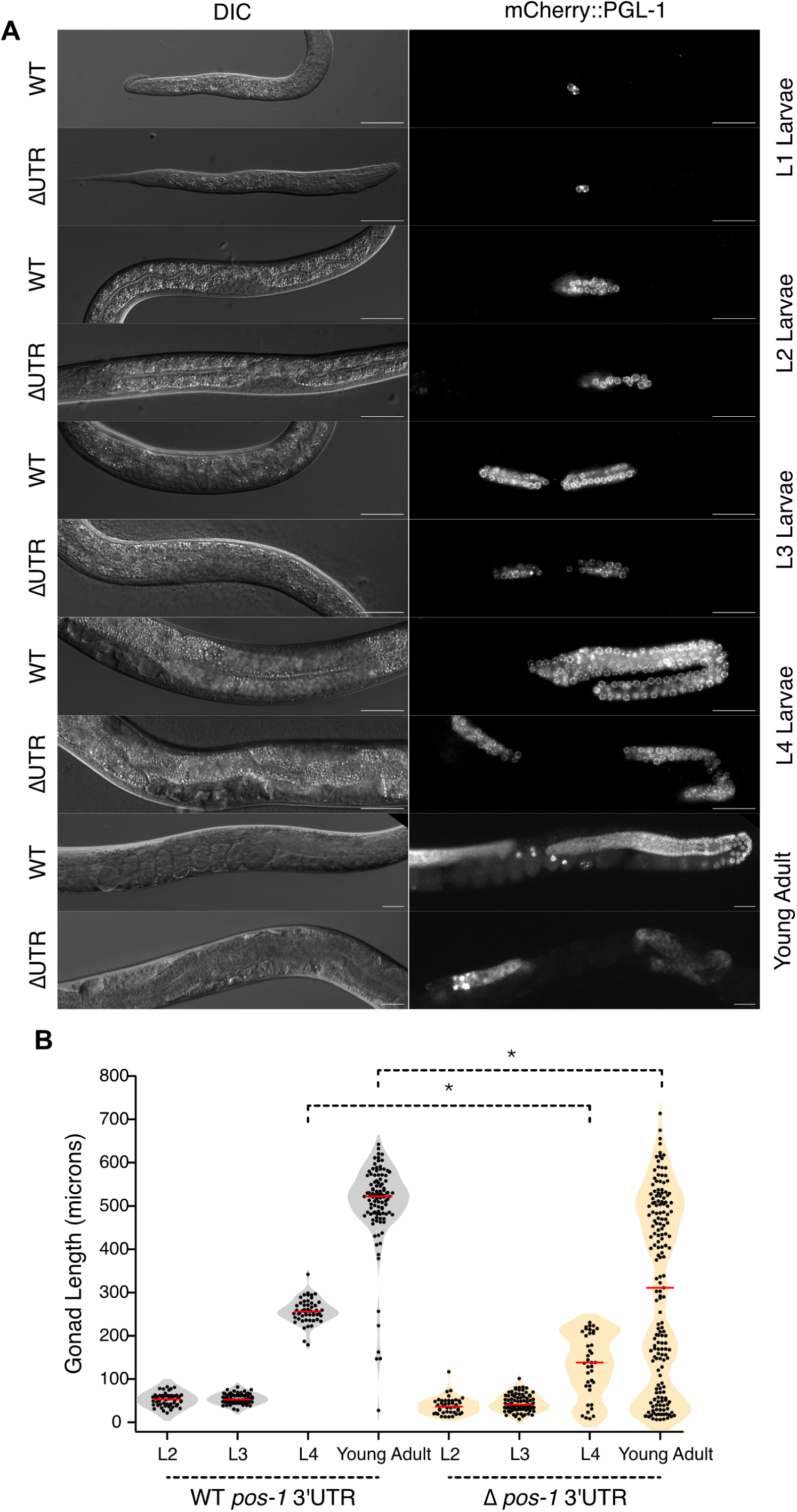
Germline proliferation is impacted in *pos-1* 3ʹUTR mutants. **A.** DIC and mCherry images of mCherry::PGL-1 marked GFP-WT UTR or GFP-ΔUTR mutants as a function of stage from L1 larva to adults. Scale bars: 20 μm. **B.** Violin plots of germline length as a function of developmental stage. Each point is the length of a single gonad arm as defined by the zone of mCherry::PGL-1 expression. Gray represents GFP-WT UTR animals, orange denotes GFP-ΔUTR animals. The red bar indicates the median. The asterisks indicate statistical significance in a one-way ANOVA with Bonferroni correction for multiple hypothesis testing.

At the L2 and L3 larval stages, the average gonad length of GFP-ΔUTR larvae is not significantly different than GFP-WT UTR larvae (Average Length_WT-L2_ = 52 µm ± 2, Average Length_ΔUTR-L2_ = 38 µm ± 3, p_adj_ = 1.0; Average Length_WT-L3_ = 54 µm ± 1.5, Average Length_ΔUTR-L3_ = 45 µm ± 2, p_adj_ = 1.0, **Fig. 6A, B**). However, in L4 larvae, GFP-ΔUTR mutants the germline arms are half the length of GFP-WT UTR controls (Average Length_WT-L4_ = 257 µm ± 4, Average Length_ΔUTR-L4_ = 130 µm ± 12, p_adj_ = 0.00008, **Fig. 6A, B**). In young adults, the GFP-ΔUTR mutant germline arms are on average 40% shorter than their WT UTR counterparts (Average Length_WT-YA_ = 503 µm ± 10, Average Length_ΔUTR-YA_ = 303 µm ± 16, p_adj_ < 1e-10). Some mutant gonads appeared to have unusual morphology. These data suggest that dysregulation of GFP::POS-1 in the germline impacts germline proliferation in the L4–young adult stages but not at earlier stages. The reduced size of the germline could also possibly explain the lower average brood size observed in **Figure 2F**.

### Dysregulation of pos-1 impacts somatic and germline genes in multiple categories

To evaluate the transcriptome-wide impacts of disrupting the *pos-1* 3ʹUTR, we performed RNA-seq on synchronized young adult WT UTR and ΔUTR animals cultured at 20°C. We used DESeq2 to identify differentially expressed transcripts across three biological replicates of each genotype [44]. This analysis identified 1437 significantly upregulated transcripts and 636 significantly downregulated transcripts with a log2 fold change cut off of 0.585, a p_adj_ threshold of 0.05, and a base mean expression level of 100 (**Fig. 7A**, **Supplemental Data Set 2**). The large number of differentially expressed genes likely reflects both direct effects and indirect impacts from the complex pleiotropic phenotype—including animals missing one or more gonads, oogenesis defects, and vulval differentiation issues.

**Fig. 7.**
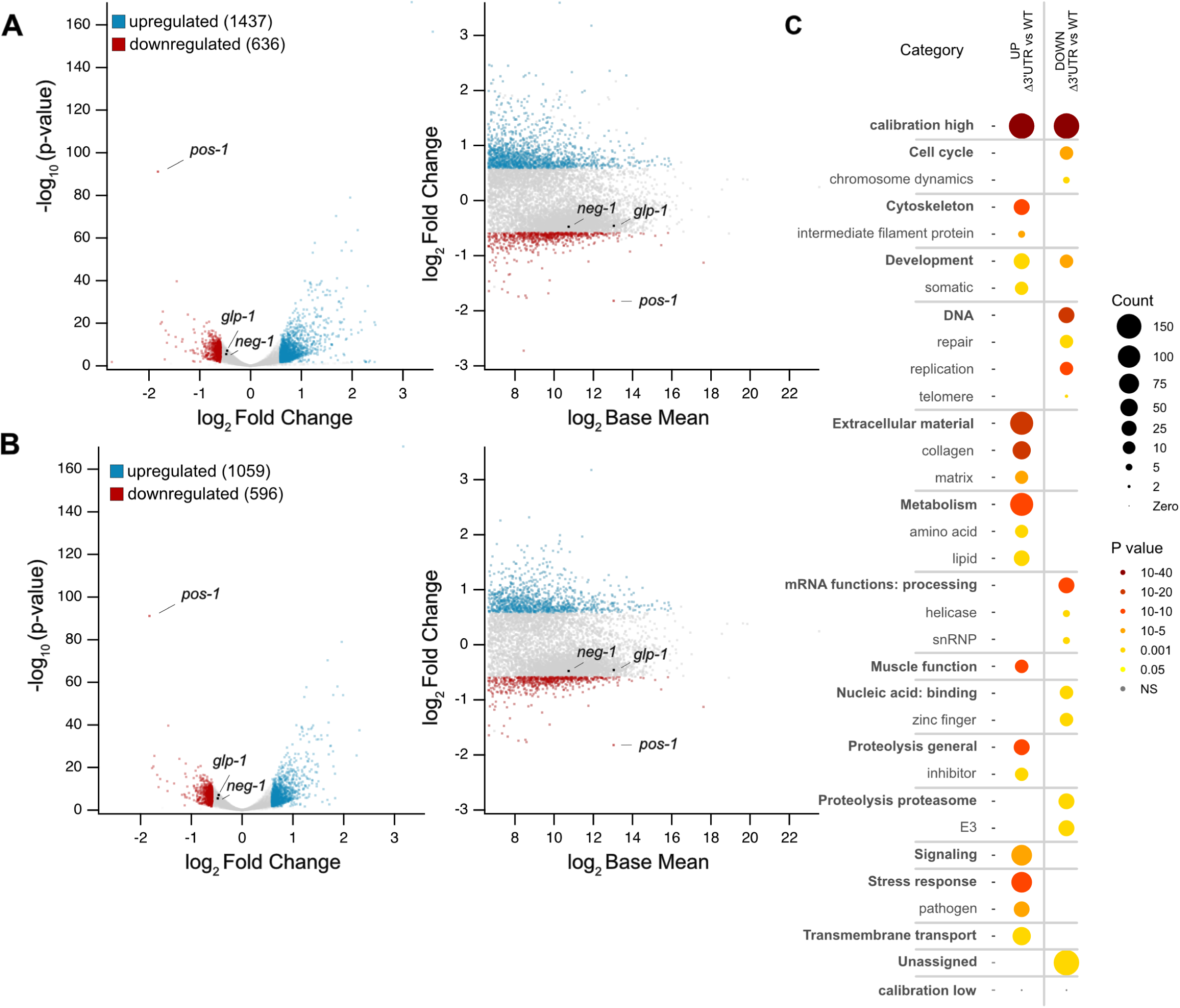
RNA sequencing reveals transcriptome-wide changes in *pos-1* 3ʹUTR mutants. **A.** Volcano and MA plots of differentially expressed genes from GFP-tagged *pos-1* WT and 3ʹUTR mutants grown at 20°C. Downregulated genes that meet filtering criteria are shown in red, upregulated genes are shown in blue. Genes that are unchanged or do not meet the filtering criteria are shown in gray. The *pos-1*, *glp-1*, and *neg-1* genes are labeled in both plots. **B.** Volcano and MA plots of the differentially regulated germline genes. The representation is the same as in panel A. **C.** Differentially expressed gene sets from WormCat analyses of both up and downregulated genes. The WormCat categories are organized by function. The colors denote statistical significance in WormCat 2.0, and the size of the circle represents the number of genes in the category impacted.

Next, we compared the list of differentially expressed genes to an annotated library of all germline transcripts [45]. The comparison identified both upregulated and downregulated germline associated transcripts in GFP-ΔUTR mutants compared to GFP-WT UTR controls using the same cut offs (up=1059, down=596, **Fig. 7B**). Most differentially expressed transcripts are germline associated genes (74% of upregulated genes, 94% of downregulated genes). The data suggest that loss of gonad arms in a large fraction of GFP-ΔUTR animals accounts for most of the downregulated transcripts. However, this phenotype cannot explain the preponderance of upregulated germline genes. We hypothesize that these changes are caused by the expanded germline expression pattern of POS-1 in the GFP-ΔUTR mutant. It is not clear why these transcripts are upregulated.

Next, we used WormCat 2.0 to perform a gene ontology analysis to identify gene sets that are dysregulated in the GFP-ΔUTR mutant [46]. Upregulated germline genes include those involved in extracellular material production (collagen, extracellular matrix), metabolism (lipids, amino acids), and stress response to pathogens, among others. Downregulated germline genes include those involved in DNA replication and repair, mRNA functions, zinc-finger genes, and cell cycle regulation. The data show that dysregulation of POS-1 has widespread impacts on germline genes involved in multiple pathways. How POS-1 exerts these impacts remains unknown.

POS-1 has been shown to regulate both *glp-1* and *neg-1* expression through direct binding to elements found in each gene’s 3ʹUTR [24, 26, 27]. The current model suggests that POS-1 regulates both transcripts at the level of translation, reducing the amount of protein produced per available mRNA without impacting mRNA abundance [26, 27]. Consistent with that model, the amount of *glp-1* and *neg-1* mRNA remains unchanged in the GFP-ΔUTR mutant compared to the GFP-WT UTR control (**Fig. 7A, B**). Intriguingly, the amount of *pos-1* mRNA is much lower in the mutant compared to the control (log2 fold change = -1.82, p_adj_ = 4.3e-88). Yet we observe much more GFP::POS-1 protein produced throughout the germline (**Fig. 4A**). The data suggests that the *pos-1* 3ʹUTR contains elements that stabilize *pos-1* mRNA as well as elements that repress its translation in the germline. It is not yet clear what the elements are, or which trans-acting factors are involved in regulation. Several RBPs are predicted to bind to the *pos-1* 3ʹUTR, including FBF-1/2, GLD-1, OMA-1/2, and POS-1 itself.

## DISCUSSION

In this study we show that the *pos-1* 3ʹUTR plays multiple roles in controlling POS-1 expression in the germline and in early embryos. The most striking consequence of the 3ʹUTR deletion mutant is strong overexpression of POS-1 protein throughout the germline, a tissue where *pos-1* mRNA is normally transcribed but remains silenced at the translational level [16]. The consequence of dysregulated POS-1 translation appears to depend both on the genetic background and the growth condition. Under optimal growth in an otherwise healthy, wild-type background, worms tolerate dysregulation of *pos-1* without much impact on reproductive fecundity. However, when worms are stressed by temperature or sensitized by their genetic background, loss of the 3ʹUTR affects 1) the number of progeny produced, 2) the number of embryos that hatch, and 3) the fraction of viable progeny that are sterile. Specific defects include misspecification of germ lineage cells in the embryo, including some that have too few and others that have too many, as well as germline proliferation defects that manifest in L4 and young adult animals. We also detected multiple issues with oogenesis which could impact all three outcomes. These phenotypes are distinct from a *pos-1* null allele, which produces inviable embryos that fail prior to gastrulation due to cell fate specification defects including absence of germline progenitors, lack of endoderm, and expanded populations of pharynx and other anterior lineage cells [16]. The exact mechanism of how the *pos-1* 3ʹUTR ensures reproductive robustness during environmental or genotoxic stress remains unknown.

### The role of the pos-1 3ʹUTR in the germline

In the GFP-ΔUTR mutant, POS-1 is expressed at a very high level throughout the germline, with maximal expression observed in the pachytene region and in immature oocytes. In a sensitized background, ectopic expression of POS-1 in the germline decreases the maximum brood, impacts oogenesis, and reduces the proliferation of germ cells in the L4 and adult stages. The molecular bases for these phenotypes are not known, but we can surmise that increased POS-1 expression in the germline might lead to competition with other RNA binding proteins normally found in the germline. Indeed, POS-1 has been shown to compete with GLD-1 for binding to *glp-1* mRNA [27]. GLD-1 encodes a STAR/GSG domain RNA-binding protein normally expressed in the pachytene region of the germline that promotes entry into meiosis and the switch from spermatogenesis to oogenesis [47, 48]. Raising the concentration of POS-1 in the germline might impact GLD-1 repression of its mRNA targets, especially for transcripts where POS-1 and GLD-1 binding motifs overlap [27]. The POS-1 motif also partially overlaps with motifs recognized by PUF domain RBPs including FBF-1/2, and MEX-3 [37, 49], both of which are expressed in the distal germline and promote mitosis [37, 50].

One of the most prevalent germline phenotypes observed in the GFP-ΔUTR mutant is that proliferation of germ cells appears to be limited in L4 and adult animals. POS-1 is known to bind to the 3ʹUTR of *glp-1* mRNA (glp: germline proliferation defective) and repress its translation in the posterior of early embryos [24, 25, 27]. GLP-1 is a Notch transmembrane domain receptor that is normally required for both proliferation of mitotically dividing germline progenitor cells in the anterior region of the germline and anterior development in the embryo [12, 51]. Overexpression of a known *glp-1* repressor in the distal germline is expected to cause reduced germ cell proliferation, as we observe in our data. More work will be needed to determine whether POS-1 regulation of *glp-1* mRNA, or dysregulation of many other transcripts, or both, underlie the germline phenotypes we observe.

### The role of the pos-1 3’UTR in embryos

In addition to the germline phenotypes, we also note phenotypes that manifest in the progeny produced by homozygous *pos-1* 3ʹUTR deletion worms. We observe a reduction in hatch rate and an increase in F1 sterility in ΔUTR mutant animals. These phenotypes become stronger in the sensitized GFP-tagged background and at mildly elevated growth temperature. Interestingly, our imaging data show that POS-1 protein levels are high at the point of fertilization in the GFP-ΔUTR background, but they drop precipitously to reach a concentration below that of GFP-WT UTR embryos within the first 85 minutes. The data suggest that mutant embryos inherit GFP::POS-1 from oocytes that express the protein ectopically. They also suggest that loss of the 3ʹUTR prevents further expression of new POS-1 protein after fertilization, leading to the reduced terminal abundance. If so, the 3’UTR might normally encode a repressive function in the germline but an activating function in embryos. Notably, the pattern of POS-1 accumulation is unaffected in the 3ʹUTR mutant, suggesting that posterior localization of POS-1 protein is not governed by the 3ʹUTR of *pos-1* mRNA. This finding is consistent with published reports that show posterior localization of POS-1 occurs at the protein level and requires ZIF-1, an E3-like ubiquitin ligase adaptor protein that promotes POS-1 turnover in the anterior of the embryo [52].

The data show that germ cell specification is frequently dysregulated in GFP-ΔUTR embryos. Intriguingly, most early embryos appear to have the normal number of progenitor cells, but as many as 30% have too many germline progenitor cells. This distribution shifts as the embryos age. After the 32-cell stage but before gastrulation, most have the normal number of germ cells, but some embryos have fewer, while others have too many. By the gastrulation stage, the distribution shifts again such that only embryos with the normal number or too few are found. We hypothesize that embryos that specify too many germline progenitor cells arrest prior gastrulation. This could explain the shift in the germline progenitor cell population as embryos age and account for the reduction in the number of viable embryos observed in the mutant. We also hypothesize that the post-gastrulation embryos that have zero or one germline progenitor cell account for our observation of some F1 adult animals that lack one or both gonad arms. Having said that, other explanations for the shift germ cell distribution and number of gonad arms are possible.

Why are the phenotypes incompletely penetrant? It is possible that variances in the amount of GFP::POS-1 protein and timing of its loss impact early cell fate decisions. Consistent with that idea, we observe that mildly elevated temperature causes reduced POS-1 abundance in immature oocytes. It is likely that less POS-1 ends up in fertilized embryos in these mutant animals grown at higher temperature, which manifests as higher penetrance of inviable embryos and a higher rate of sterility in F1 progeny. However, the changes in POS-1 abundance at elevated temperature and enhanced phenotypes are correlative. More work will be needed to determine whether these observations are connected mechanistically.

### Somatic phenotypes

How does deletion of the *pos-1* 3ʹUTR manifest somatic phenotypes in a subset of the progeny? The imaging data show that POS-1 is expressed in the germline and in germline progenitor cells in the early embryo. Yet we observe two different types of somatic vulval morphogenesis defects, including protruding vulva and multivulva phenotypes in a subset of progeny. We also observe a dark gut phenotype. The RNA-seq data helps rationalize these observations. Loss of the *mrt-2* gene causes a dark gut phenotype like the one observed in some GFP-ΔUTR animals [40]. Transcripts encoding *mrt-2* are downregulated in GFP-ΔUTR compared to GFP-WT UTR controls. This gene encodes a DNA repair enzyme that regulates telomere length and causes a transgenerational mortal germline phenotype [53]. As such, reduction of this gene could contribute to both the dark gut and the loss of gonad phenotypes that we observed.

The genetic pathways that define vulval morphogenesis have been well characterized [54]. The protruding vulva (pvul) and multivulva (muv) phenotypes could be caused by myriad changes to transcript abundance for genes involved in regulating this pathway. We note that *lin-3*, an EGF-like ligand necessary for vulva cell induction [55], is down regulated in the RNA-seq data set. We also observe an increase in *lin-15a*, *lin-15b*, *cwn-1*, *cwn-2*, and *bar-1* transcripts, all genes involved in specifying vulval differentiation in vulva precursor cells. LIN-15A and LIN-15B are transcriptional regulators that negatively regulate vulva cell fate [56]. Loss-of-function alleles in both genes cause muv phenotypes. CWN-1 and CWN-2 are both wnt signaling pathway ligands that contribute to vulval precursor cell polarity [57]. BAR-1 (beta-catenin) is the major effector of wnt signaling pathways [58]. When activated, this protein accumulates in the nucleus of vulval precursor cells to promote vulva fate. The complex mixture of pvul and muv animals in our mutants probably results from the dysregulation of multiple genes involved in vulval morphogenesis. It is not yet known how these genes are impacted by POS-1 dysregulation.

### What is the role of maternal mRNA?

Spirin first hypothesized in 1966 that oocytes contain “masked” mRNAs, produced and stored in the maternal germline, to be used by embryos after fertilization prior to the onset of zygotic gene activation [59]. The idea was that these mRNAs are necessary to ensure robust cell physiology at a time when DNA replication occurs rapidly and repeatedly. We now know that multiple embryonic cell fate determinants are transmitted from the maternal germline to embryos in mRNA form. The regiospecific activation of these transcripts after fertilization is thought to guide early developmental decisions such as cell fate specification, anterior-posterior axis formation, and differentiation of the germline from the soma [8]. The results shown here, similar to a second cell fate determining transcript *mex-3* [30, 31], suggest that regulation of maternal mRNAs may not be as essential for embryonic survival as previously thought. As with *pos-1*, loss of the *mex-3* 3ʹUTR has a modest impact on reproductive success [30]. Only under stress conditions is its importance revealed [31]. The 3ʹUTRs of both transcripts appear to enhance reproductive robustness in conditions that are less than ideal, but neither is essential to reproduction. Additional studies on more maternal transcripts are needed to assess whether the lessons learned from *mex-3* and *pos-1*—two important maternal genes required for very early pattern formation—hold for most maternal mRNAs. Additional deletion alleles will be needed to fully evaluate the role of maternal 3ʹUTRs in early embryogenesis and thus comprehensively test Spirin’s hypothesis.

## MATERIALS AND METHODS

### Strains, Nematode Culture, and CRISPR mutagenesis

All *C. elegans* strains used in this study were propagated on Nematode Growth Media (NGM) plates seeded with *Escherichia coli* OP50 under standard growth conditions unless otherwise noted [38]. The strains referenced within this study are listed in **Table 1**. WRM101 and WRM102 were generated using CRISPR-Cas9 mutagenesis following the procedure of Ghanta and Mello [32]. Custom Alt-R modified crRNA guides and single stranded oligonucleotide donor (ssODN) homology directed repair (HDR) templates were designed and ordered using the Integrated DNA Technologies (IDT, Coralville, IA) Alt-R CRISPR HDR design tool on the IDT website (https://www.idtdna.com/pages/tools/alt-r-crispr-hdr-design-tool). All primer, crRNA, and ssODN sequences used for strain generation, PCR confirmation, or sequencing are listed in **Supplementary Table 1**. Strain DG4222 was obtained from the *Caenorhabditis* Genetics Center (CGC, Minneapolis, MN) [33]. This strain was outcrossed three times through the wild-type N2 strain, and the *gfp::tev::3xflag* integration at the *pos-1* locus was confirmed by Sanger sequencing, prior to use in experiments.

To generate strain WRM101, 3x outcrossed DG4222 hermaphrodites were injected with a mixture containing 10 µg/µL S*treptococcus pyogenes* Alt-R modified Cas9 (IDT, Cat. #1081058), 0.4 µg/µL tracrRNA (IDT, Cat. #1073189), 2nmol crRNA (CD.Cas9.BZLP9054.AB and CD.Cas9.BZLP9054.AM) resuspended to 0.4 µg/µL with IDTE buffer (IDT, Cat. #11-01-02-02), 1 µg/µL ssODN (pos-1_ssODN_UTR-Delete), 500 ng/µL pRF4 (*rol-6*) plasmid as an injection marker, and nuclease free water (IDT, Cat. #11-04-02-01). Because two crRNAs were included, 1.4 µL of each was used. WRM102 was generated using the same injection mix with N2 hermaphrodites. Mutants were identified through PCR using primers that span the entire *pos-1* 3′UTR (pos-1_PCR004F and pos-1_PCR005R). Heterozygotes of the mutant alleles were propagated, and progeny were isolated and reconfirmed via PCR to identify homozygotes. Both strains can be maintained as homozygotes at 20°C on NGM plates.

WRM85 was generated using the same CRISPR-Cas9 protocol described above with the following exceptions. One crRNA (CD.HC9.YJVG2684.AB) was used instead of two, and an HDR template DNA containing the mCherry coding sequence was amplified from a plasmid (pCCM953, a gift from Craig Mello, UMass Chan Medical School, Worcester, MA, USA) using primers Linker1_pgl1_HA_F and Linker1_pgl1_HA_R (**Supplemental Table 1**). The PCR product was purified using the Zymo Research DNA Clean & Concentrator™ kit (Catalog Number D4034, Irvine, CA) prior to use in injections. The genotype of WRM85 was confirmed by PCR and Sanger sequencing before experiments were performed.

WRM105 was generated by crossing DG4222 males with WRM85 hermaphrodites. Upon signs of successful mating, young hermaphrodite F1 cross progeny were isolated and allowed to propagate. F2 progeny were allowed to mature, and fluorescence imaging was used to identify plates where mutants contained both GFP::POS-1 and mCherry::PGL-1. The isolation and fluorescence confirmation process was continued until homozygosity was achieved for both alleles. WRM104 was generated using the same approach, except WRM101 males were crossed with WRM85 hermaphrodites. Both WRM104 and WRM105 can be propagated as homozygotes at 20°C on NGM plates.

### Brood Size, Hatch Rate, and Sterility Assays

Brood size, hatch rate, and sterility assays were performed for strains DG4222, WRM101, WRM102, WRM104, WRM105, and N2. L1 larval synchronization was achieved by bleaching young adult worms with 20% alkaline hypochlorite solution (3 mL Clorox bleach, 3.75 mL filtered 1M NaOH, 8.25 mL ddH_2_O) to recover embryos, which were extensively washed with M9 buffer and allowed to hatch at room temperature overnight. L1 hatchlings were plated onto NGM agar plates seeded with OP50 *E. coli* and allowed to mature to L4 larvae at 20°C. For each genotype, L4 larvae were isolated onto NGM plates with OP50 and placed under standard growth temperature (20°C) or mild temperature stress (25°C). Each worm was transferred daily to a new NGM plate with OP50, and the number of embryos laid onto the previous day’s plate was counted. Viable F1 progeny that hatched were counted 24 hours after each embryo count. Once larval hatchlings matured to adulthood, approximately 72 hours after larval count, F1 hermaphrodites were scored for the presence (fertile) or absence (sterile) of a uterus containing oocytes and/or embryos. For each strain, this entire assay procedure was repeated in triplicate.

The total number of embryos laid per worm (brood size) was calculated by summing the embryo counts over the course of each worm’s reproductive window. The total viable progeny was calculated by summing the larval counts over the same period. The number of viable progeny was divided by the total brood to determine the hatch rate for each animal. The sterility rate was calculated by dividing the total number of F1 hermaphrodite progeny categorized sterile by the total number of F1 progeny assessed. Comparisons of the brood size, hatch rate, and sterility rate were made using a one-tailed ANOVA with Bonferroni correction for multiple hypothesis testing using StatPlus software (V8.0.4.0, AnalystSoft, Alexandria, VA). For yes or no assessments concerning the presence of sterile progeny, comparisons between genotypes and growth conditions were analyzed using a Pearson’s Chi-Square test in StatPlus, and post-hoc pairwise p-values were adjusted with Bonferroni correction for multiple hypothesis testing.

### Whole Genome Sequencing

Worm DNA samples were harvested from two freshly starved medium plates per genotype (WRM101, WRM102, DG4222). Samples were lysed using 200 µL of 1X Worm Lysis Buffer (10 mM Tris-Cl pH 8.0, 50 mM KCl, 2 mM MgCl_2_). DNA was purified from crude lysate using a Qiagen DNeasy blood and tissue kit (Cat. #69504, Germantown, MD) following the manufacturers protocol. DNA samples were analyzed using a QuBit fluorometer (ThermoFisher, Waltham, MA) to determine concentration and quality before being sent to Novogene (Davis, CA) for library construction, sequencing, and data analysis through their whole genome sequencing service. The entire report from Novogene, including quality control, data filtering, and SNP / Indel characterization pipelines, is provided as **supplemental data set 1**. Briefly, FastP v.0.20.0 was used to assess sequencing quality, BWA v.0.7.17 was used to map the reads to the reference genome (WBcel235 GCF_000002985.6), and SamTools v.1.3.1 was used to map indels and single nucleotide polymorphisms.

### Gonad Fluorescence Imaging and Quantitation

For gonad imaging, synchronized DG4222 and WRM101 larvae were propagated at standard growth temperature (20°C) or under mild temperature stress (25°C) and selected as young adult hermaphrodites for imaging. Worms were mounted onto 2% agarose pads (Fisher BioReagents Cat. #BP160-500) on plain glass slides (Corning Cat. #2947-75x25). A volume of 1 µL of 1 mM levamisole was pipetted onto the pad to paralyze the worms, and an additional 1–2 µL of M9 buffer was added before the cover slip (Fisherbrand Cat. No. 12-542-B). DIC and GFP images of gonads were taken using a Zeiss AxioObserver 7 microscope (Oberkochen, Germany) with a 20X objective. Images collected with identical microscope settings were analyzed using the segmented line tool in Fiji V2.14.0 (ImageJ) software with a 20 µm width to define the average GFP::POS-1 fluorescence intensity profile from the distal tip to the most proximal oocyte, measuring through the center of the gonad while avoiding nuclei in oocytes. A control line of approximately equal length and width was drawn outside of the worm and used as a background control. The background corrected intensity values were binned into 20 equal partitions per gonad and averaged across multiple images of worms with the same genotype and growth conditions. Statistical significance between genotypes and growth conditions was assessed using a one-tailed ANOVA with Bonferroni correction for multiple hypothesis testing using StatPlus software, as above.

### Embryogenesis Movies and Quantitation

DG4222 and WRM101 worms were propagated at standard growth temperature (20°C) or under mild temperature stress (25°C) for at least one generation before being selected for dissection. Worms were mounted onto 2% agarose pads on plain glass slides in 1-2 µL of levamisole and dissected at the vulva using a 26-gauge needle (BD PrecisionGlide 305110) to release the embryos. A volume of 1-2 µL of M9 buffer is added before application of the coverslip, and then an additional 2–3 µL of M9 was added under the cover slip to prevent the slide from drying out. One cell embryos were identified under DIC optics, and then DIC and GFP images were collected at five-minute intervals for a total of 90 minutes using a Zeiss AxioObserver 7 microscope with a 63X oil immersion objective (Zeiss Immersion Oil Cat. #444960-0000-000). Focus was maintained with a Zeiss Definite Focus module. All images were collected using identical microscope settings.

The total embryonic fluorescence intensity was measured at each time point using the oval selection tool in Fiji (ImageJ). An oval of equal area was drawn outside of the embryo and was used to determine the background fluorescence. Due to the inability to synchronize fertilization, the first frame where cytokinesis is observed during the first cellular division was arbitrarily set to time point zero. The time axis of all embryos was synchronized to this morphological feature to enable comparison of multiple embryos. The average fluorescence intensity across multiple embryos and multiple movies was plotted as a function of time. A one-tailed ANOVA with Bonferroni correction for multiple hypothesis testing was conducted using StatPlus software to assess statistical significance in intensity between time points.

To define the time point with half maximal fluorescence intensity in strain WRM101, the data were fit to a sigmoid equation using IgorPro software (v9.02, Wavemetrics, Lake Oswego, OR)

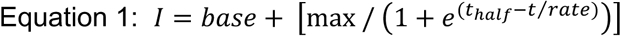

where I is the average fluorescence intensity, t is the time, t_half_ is the time with half maximal fluorescence intensity, rate defines the shape of the sigmoid, and base and max describe the minimal and maximal bounds of the fluorescence intensity. To define the time of peak fluorescence in strain DG4222, the average fluorescence intensity data were fit to a modified Gaussian function using IgorPro software

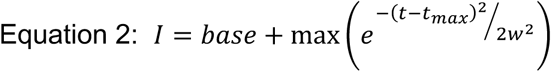

where I is average fluorescence intensity, t is the time, t_max_ is the time with maximal fluorescence intensity, and w is the width of the Gaussian, where w = width of the left tail for t<t_max_, and w = width of the right tail for t>t_max_. In both cases, the fits were weighted by the standard error of the mean for each time point, and the reported error is the error of the fitted parameter.

### Embryonic Germ-Cell Progenitor Imaging and Quantitation

WRM104 and WRM105 strains were grown at 20°C for at least one generation before individual worms were selected for dissection as described above. DIC, mCherry, and GFP images of embryos were acquired using a Zeiss AxioObserver 7 as described above. Images of the embryos for each genotype were binned by the approximate cell count and/or developmental stage into the following categories: 2-32 cells, 33-200+ cells, or bean through pretzel stages. The number of germ cell progenitors was determined by counting the number of cells expressing mCherry::PGL-1. The data sets for each genotype and each bin were compared using a Pearson’s Chi-Square test with Bonferroni correction for multiple hypothesis testing of post-hoc pairwise comparisons using StatPlus as described above.

### Larvae Gonad Length Imaging and Quantitation

WRM104 and WRM105 embryos were synchronized by bleaching as described above. Synchronized L1 worms were plated onto NGM agar plates seeded with *E. coli* OP50. DIC, GFP, and mCherry images were taken for L1 through L4 larval stages using a 40X oil immersion objective on a Zeiss AxioObserver 7 microscope. L1 larvae were imaged between 0–8 hours after plating. Because L1 larvae have yet to begin germ cell proliferation, the quantity of germ cell progenitors was counted rather than the length of the gonad. The subsequent L2 and L3 larval stages were imaged 20–24 hours after plating. Differentiation between each stage was based on the presence or absence of separation of the gonad arms, indicating completion of the L2 to L3 molt. Analysis of gonad length was conducted using the segmented line tool in Fiji (ImageJ) software as described above. To measure the length, a line was drawn from the distal to the proximal end of the gonad using PGL-1 expression as a guide. From the L3 larval stage and onwards, each gonad arm length was measured separately unless a gonad arm was absent (no value recorded). L4 larvae were imaged 40-48 hours after plating and identified based on the observance of the characteristic square vulval lumen. Young adult hermaphrodites were selected at random and imaged 65-70 hours after plating. Young adult gonads were too large to properly fit in frame using the 40X objective, thus the 20X objective was utilized for this developmental stage. If gonads were fragmented, the length of each fragment was summed to avoid erroneous overmeasurement of gaps. We found that taking Z-stack images helped define the distal and proximal ends of each gonad for all stages except L1. Statistical significance between comparison groups was determined using a one-way ANOVA test with Bonferroni correction for multiple hypothesis testing in StatPlus software.

### RNA Sequencing

Five medium plates (60mm x 15mm petri dish seeded with 200 µL of OP50 bacteria) of synchronized young adult stage worms were harvested using 1 mL ddH_2_O per plate and combined into a 15 mL conical tube. Tubes were spun in a room temperature centrifuge for 1 minute at 3000 rpm (2100 x g), and pellets were washed with 3 mL ddH_2_O four more times. Pellets from each replicate were aggregated into one 1.7 mL tube each then washed once more to remove any remaining contaminating OP50 (960 x g). Each tube was flash frozen in liquid nitrogen to lyse the worms. Total RNA was extracted using eight pellet volumes of TRIzol, followed by1.6 volumes of chloroform. The aqueous phase was extracted, and an equal volume of isopropanol was added to precipitate the RNA. Extraction was followed by rRNA depletion using a *C. elegans* optimized protocol [60]. Sample libraries were prepared using the NEBNext Ultra II library prep kit (E7775S) and barcoded using the NEBNext Multiplex Oligos for Illumina (Dual Index Primer Set 1 (E7335S) & Set 2 (E7500S)). Sample concentration was calculated using a Qubit Fluorometer and Fragment analyzer prior to data collection on an Illumina NEXTSeq 1000 (Illumina, San Diego, CA). In total, data from three biological replicates per genotype were collected.

The resulting raw sequencing reads were submitted to the OneStopRNASeq pipeline for quality control and differential gene expression analysis [61]. FastQC v.0.11.5 was used for raw sequencing quality control, and MultiQC v.1.6 was used post-alignment quality control using the default configurations in the OneStopRNASeq pipeline. STAR v2.7.5a was used to align the reads to the reference genome using WBcel235.90 annotations using the following parameters ‘-Q 20 –minOverlap 1 --fracOverlap 0 -p -B -C’ for paired-end strict-mode analysis [62]. Differential expression (DE) analysis was performed with DESeq2 v1.28.1 [44]. Significantly differentially expressed genes were filtered with the criteria FDR < 0.05, absolute log2 fold change (|LFC|) > 0.585, and Base Mean expression level of >100.

### Data availability

The whole genome sequencing data report described in **Supplemental Figure 1** is included as **Supplementary Data Set 1**. The OneStopRNASeq described in **Figure 7** pipeline report is included as **Supplementary Data Set 2.** All other numerical data and statistical analyses described in this manuscript are available in **Supplementary Data Set 3**. Raw sequencing data were uploaded to the NIH Sequence Read Archive and will be made available under Bioproject PRJNA1368703.

## Supporting information

Supplemental Data Set 2

Supplemental Data Set 1

Supplemental Data Set 3

## ACKNOWLEDGEMENTS

The authors thank Dr. Craig Mello for the gift of plasmid pCCM953. We thank Dr. Ye Duan for technical support with the rRNA depletion protocol. We also thank Christable Darko, Sharon Noronha, and Oscar Lam for helpful comments and constructive criticisms. This work was supported by NIH Grant R01HD111505 to S.P.R.

## AUTHOR CONTRIBUTIONS

HV made strains WRM101, WRM102, WRM104, and WRM105. HB made strain WRM85. HV collected and analyzed all the data presented in the manuscript with the following exceptions: JU outcrossed strain DG4222 and confirmed the gene by sequencing, BR and MV assisted with the data collection for Figure 4, SR analyzed the data described in Supplemental Figure 1, and HV and SR analyzed all other data sets together. All experiments were conceived by HV and SR. HV and SR wrote the manuscript under the advisement of all authors.

## SUPPLEMENTAL INFORMATION

1. Supplemental Figure 1
2. Supplemental Movie 1
3. Supplemental Movie 2
4. Supplemental Table 1
5. Supplemental Data Set 1
6. Supplemental Data Set 2
7. Supplemental Data Set 3

### SUPPLEMENTAL FIGURE LEGEND

**Supp. Fig. 1.**
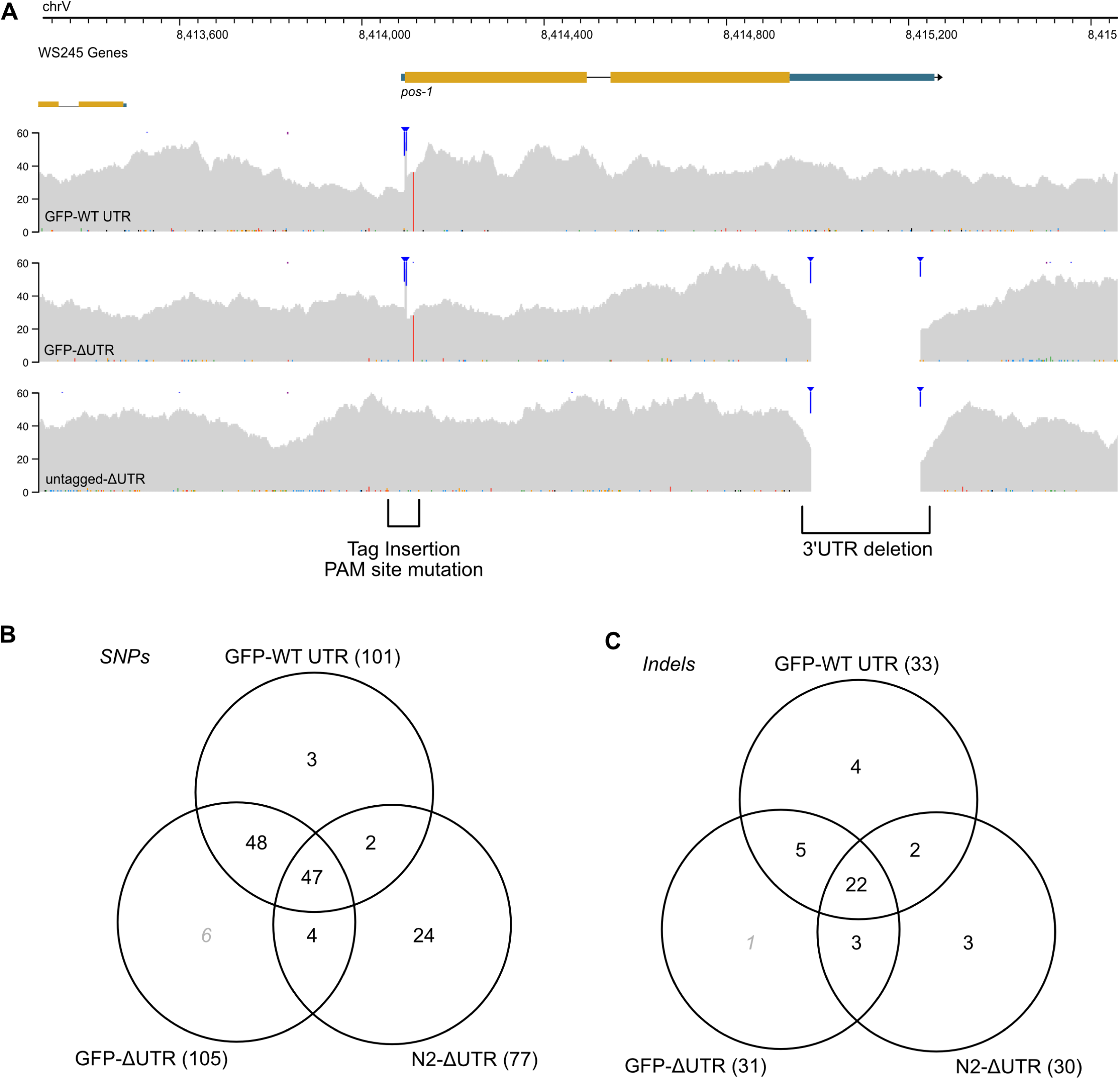
Confirmation of the strain background by whole genome resequencing. **A.** A genome viewer image of the *pos-1* locus with pileup views for GFP-ΔUTR, GFP-WT UTR, and untagged N2-ΔUTR whole genome sequencing data. The brackets show the location of the 3ʹUTR deletion and the point of insertion of the *gfp::tev::3xflag* tag. The vertical red stripe is a silent mutation in the *pos-1* gene that removes a PAM site. **B.** Venn diagram of the overlap between exonic SNP calls from the whole genome sequencing data. The number for the six unique GFP-ΔUTR SNPs is listed in gray because visual inspection of the sequencing tracks in a genome data browser shows that all six alleles are also present in the GFP-WT UTR strain at an allele frequency >0.5, with the exception of *Y22D7AR.2* which has an allele frequency of <0.5, as described in the text. These represent false negatives in the bioinformatic pipeline that likely correspond to the allele frequency cutoff. **C.** Venn diagram of exonic indel alleles in the whole genome sequencing data. The lone candidate allele in the GFP-ΔUTR strain is listed in gray because it was also found in the untagged N2-ΔUTR strain by visual inspection of the genome sequencing tracks.

### SUPPLEMENTAL MOVIE LEGENDS

**Supp. Movie 1**

Time lapse images of GFP-WT UTR and GFP-ΔUTR embryos are presented. Each frame represents five minutes of elapsed time. The scale bar represents 5 microns. The growth temperature was 20°C. DIC and GFP images for both strains are presented side by side.

**Supp. Movie 2**

The time lapse images shown here are labeled as in Supplemental Movie 1. The only difference is that the animals were grown at 25°C.

**Supplemental Table 1.**
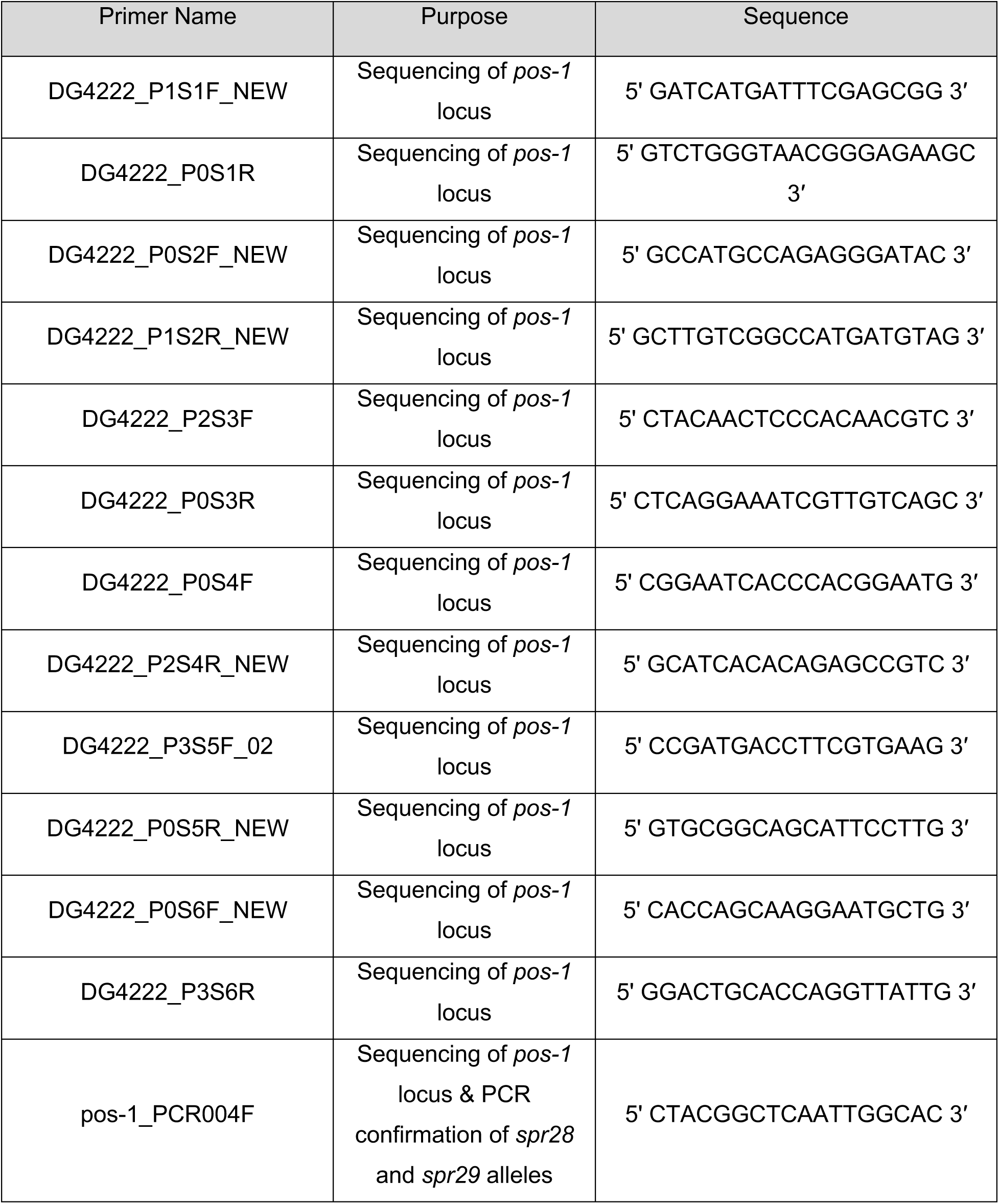
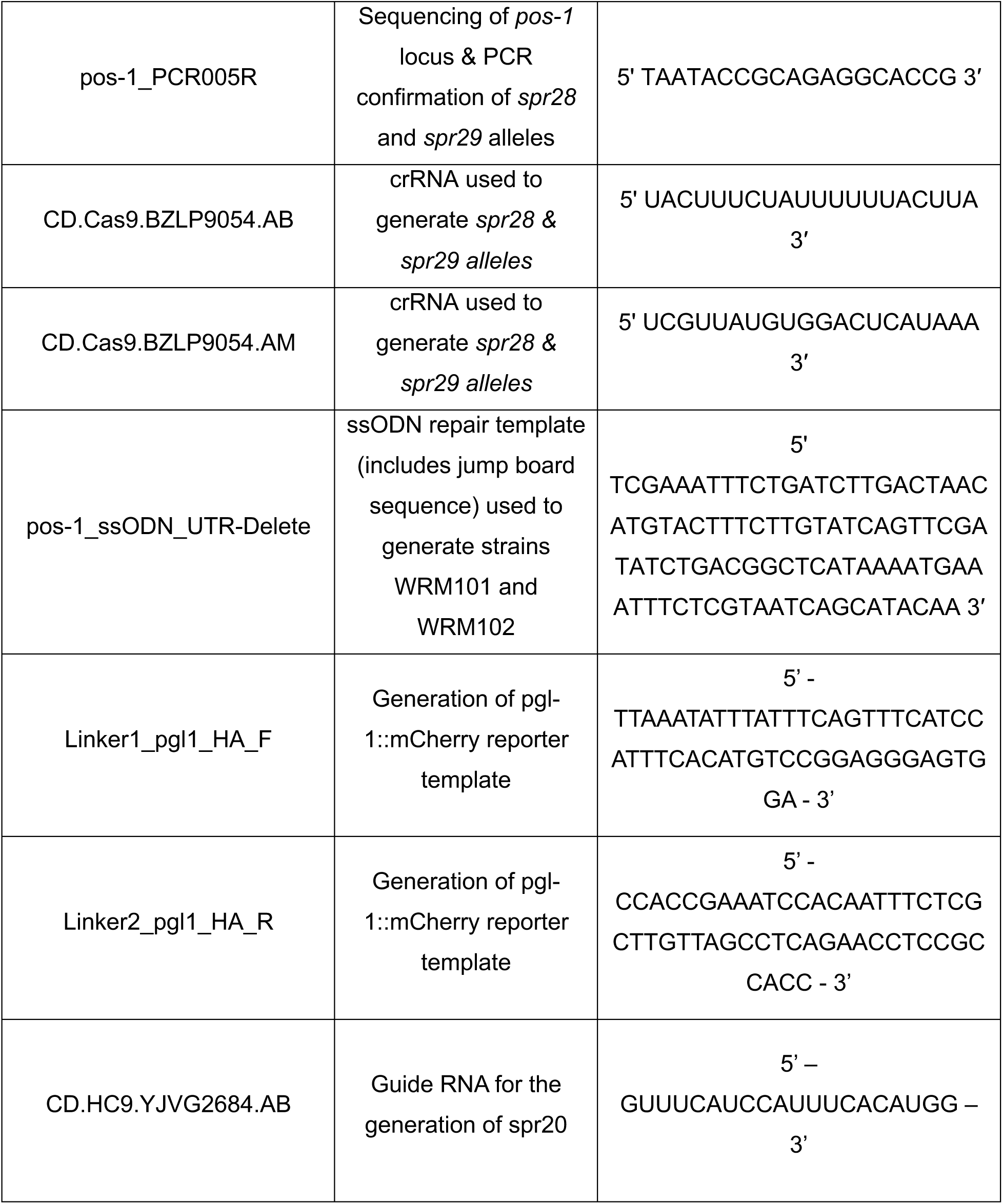
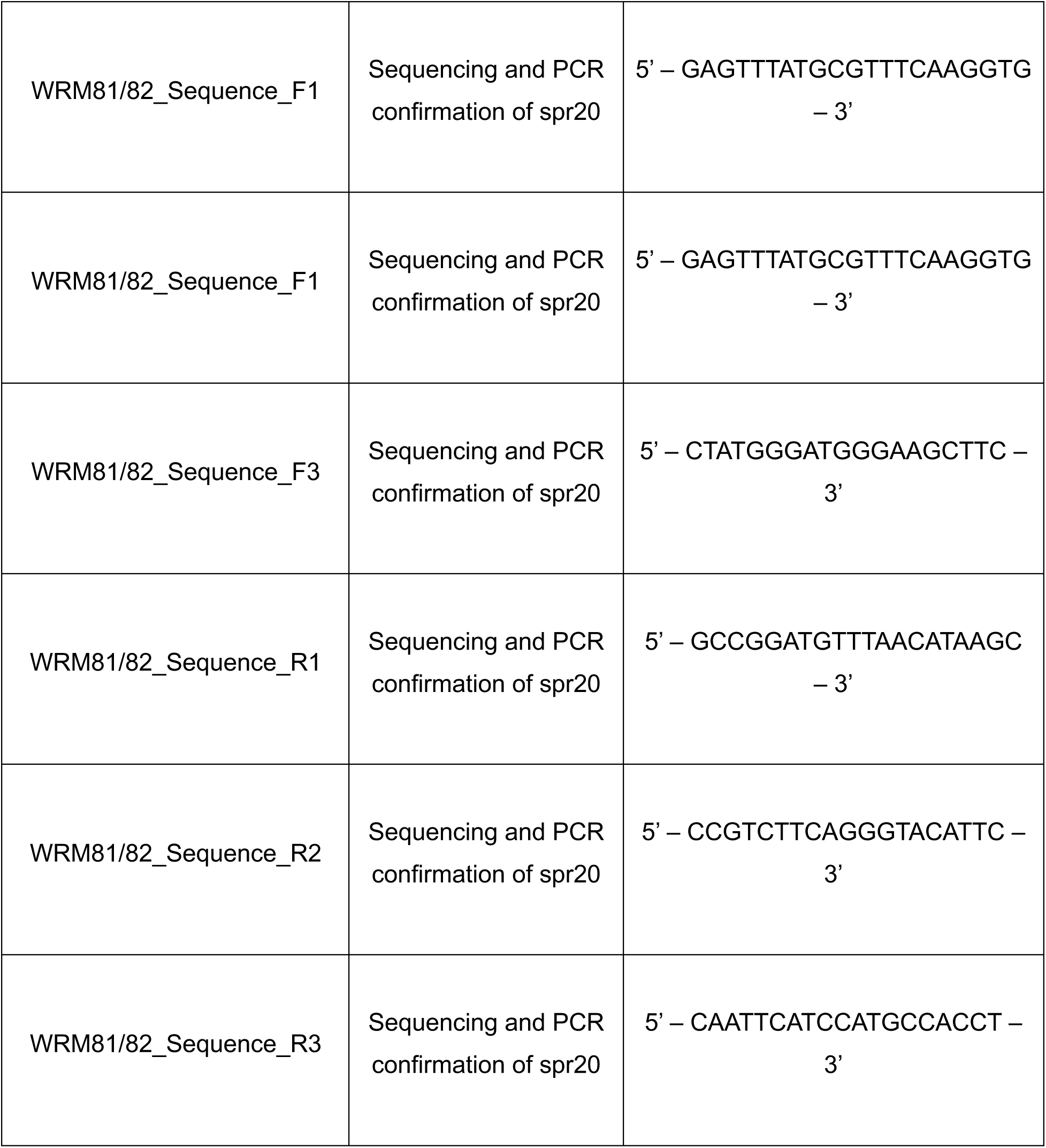

