## Supplemental Data Set 2 for "The *pos-1* 3ʹ untranslated region governs germline specification and proliferation to ensure reproductive robustness": ReadMe.txt

Supplemental Data Set 2
Varderesian et al., 2025

File:
DESeq2_Analysis.xlsx (Microsoft Office)

This file contains the OneStopRNASeq results, and our analysis with filters of >100 Base Mean expression, |log2FC|>0.585, and Padj<0.05. These analyses were used to make Figure 7

Folder:
OneStopRNASeq

This folder contains the complete output of the OneStopRNASeq pipline, including fastqc reports, bam_qc reports, and all data / visualizations from the DESeq2 package.

Folder:
WormCat

This folder contains the complete output of WormCat 2.0 web server, including visualizations, excel spreadsheets, and html files. The analyses were divided into unregulated and down regulated categories.
