## Supplemental Data Set 1 for "The *pos-1* 3ʹ untranslated region governs germline specification and proliferation to ensure reproductive robustness": jquery-ui.min.js

```
/*! jQuery UI - v1.12.1 - 2016-09-14
* http://jqueryui.com
* Includes: widget.js, position.js, data.js, disable-selection.js, effect.js, effects/effect-blind.js, effects/effect-bounce.js, effects/effect-clip.js, effects/effect-drop.js, effects/effect-explode.js, effects/effect-fade.js, effects/effect-fold.js, effects/effect-highlight.js, effects/effect-puff.js, effects/effect-pulsate.js, effects/effect-scale.js, effects/effect-shake.js, effects/effect-size.js, effects/effect-slide.js, effects/effect-transfer.js, focusable.js, form-reset-mixin.js, jquery-1-7.js, keycode.js, labels.js, scroll-parent.js, tabbable.js, unique-id.js, widgets/accordion.js, widgets/autocomplete.js, widgets/button.js, widgets/checkboxradio.js, widgets/controlgroup.js, widgets/datepicker.js, widgets/dialog.js, widgets/draggable.js, widgets/droppable.js, widgets/menu.js, widgets/mouse.js, widgets/progressbar.js, widgets/resizable.js, widgets/selectable.js, widgets/selectmenu.js, widgets/slider.js, widgets/sortable.js, widgets/spinner.js, widgets/tabs.js, widgets/tooltip.js
* Copyright jQuery Foundation and other contributors; Licensed MIT */

(function(t){"function"==typeof define&&define.amd?define(["jquery"],t):t(jQuery)})(function(t){function e(t){for(var e=t.css("visibility");"inherit"===e;)t=t.parent(),e=t.css("visibility");return"hidden"!==e}function i(t){for(var e,i;t.length&&t[0]!==document;){if(e=t.css("position"),("absolute"===e||"relative"===e||"fixed"===e)&&(i=parseInt(t.css("zIndex"),10),!isNaN(i)&&0!==i))return i;t=t.parent()}return 0}function s(){this._curInst=null,this._keyEvent=!1,this._disabledInputs=[],this._datepickerShowing=!1,this._inDialog=!1,this._mainDivId="ui-datepicker-div",this._inlineClass="ui-datepicker-inline",this._appendClass="ui-datepicker-append",this._triggerClass="ui-datepicker-trigger",this._dialogClass="ui-datepicker-dialog",this._disableClass="ui-datepicker-disabled",this._unselectableClass="ui-datepicker-unselectable",this._currentClass="ui-datepicker-current-day",this._dayOverClass="ui-datepicker-days-cell-over",this.regional=[],this.regional[""]={closeText:"Done",prevText:"Prev",nextText:"Next",currentText:"Today",monthNames:["January","February","March","April","May","June","July","August","September","October","November","December"],monthNamesShort:["Jan","Feb","Mar","Apr","May","Jun","Jul","Aug","Sep","Oct","Nov","Dec"],dayNames:["Sunday","Monday","Tuesday","Wednesday","Thursday","Friday","Saturday"],dayNamesShort:["Sun","Mon","Tue","Wed","Thu","Fri","Sat"],dayNamesMin:["Su","Mo","Tu","We","Th","Fr","Sa"],weekHeader:"Wk",dateFormat:"mm/dd/yy",firstDay:0,isRTL:!1,showMonthAfterYear:!1,yearSuffix:""},this._defaults={showOn:"focus",showAnim:"fadeIn",showOptions:{},defaultDate:null,appendText:"",buttonText:"...",buttonImage:"",buttonImageOnly:!1,hideIfNoPrevNext:!1,navigationAsDateFormat:!1,gotoCurrent:!1,changeMonth:!1,changeYear:!1,yearRange:"c-10:c+10",showOtherMonths:!1,selectOtherMonths:!1,showWeek:!1,calculateWeek:this.iso8601Week,shortYearCutoff:"+10",minDate:null,maxDate:null,duration:"fast",beforeShowDay:null,beforeShow:null,onSelect:null,onChangeMonthYear:null,onClose:null,numberOfMonths:1,showCurrentAtPos:0,stepMonths:1,stepBigMonths:12,altField:"",altFormat:"",constrainInput:!0,showButtonPanel:!1,autoSize:!1,disabled:!1},t.extend(this._defaults,this.regional[""]),this.regional.en=t.extend(!0,{},this.regional[""]),this.regional["en-US"]=t.extend(!0,{},this.regional.en),this.dpDiv=n(t("<div id='"+this._mainDivId+"' class='ui-datepicker ui-widget ui-widget-content ui-helper-clearfix ui-corner-all'></div>"))}function n(e){var i="button, .ui-datepicker-prev, .ui-datepicker-next, .ui-datepicker-calendar td a";return e.on("mouseout",i,function(){t(this).removeClass("ui-state-hover"),-1!==this.className.indexOf("ui-datepicker-prev")&&t(this).removeClass("ui-datepicker-prev-hover"),-1!==this.className.indexOf("ui-datepicker-next")&&t(this).removeClass("ui-datepicker-next-hover")}).on("mouseover",i,o)}function o(){t.datepicker._isDisabledDatepicker(m.inline?m.dpDiv.parent()[0]:m.input[0])||(t(this).parents(".ui-datepicker-calendar").find("a").removeClass("ui-state-hover"),t(this).addClass("ui-state-hover"),-1!==this.className.indexOf("ui-datepicker-prev")&&t(this).addClass("ui-datepicker-prev-hover"),-1!==this.className.indexOf("ui-datepicker-next")&&t(this).addClass("ui-datepicker-next-hover"))}function a(e,i){t.extend(e,i);for(var s in i)null==i[s]&&(e[s]=i[s]);return e}function r(t){return function(){var e=this.element.val();t.apply(this,arguments),this._refresh(),e!==this.element.val()&&this._trigger("change")}}t.ui=t.ui||{},t.ui.version="1.12.1";var h=0,l=Array.prototype.slice;t.cleanData=function(e){return function(i){var s,n,o;for(o=0;null!=(n=i[o]);o++)try{s=t._data(n,"events"),s&&s.remove&&t(n).triggerHandler("remove")}catch(a){}e(i)}}(t.cleanData),t.widget=function(e,i,s){var n,o,a,r={},h=e.split(".")[0];e=e.split(".")[1];var l=h+"-"+e;return s||(s=i,i=t.Widget),t.isArray(s)&&(s=t.extend.apply(null,[{}].concat(s))),t.expr[":"][l.toLowerCase()]=function(e){return!!t.data(e,l)},t[h]=t[h]||{},n=t[h][e],o=t[h][e]=function(t,e){return this._createWidget?(arguments.length&&this._createWidget(t,e),void 0):new o(t,e)},t.extend(o,n,{version:s.version,_proto:t.extend({},s),_childConstructors:[]}),a=new i,a.options=t.widget.extend({},a.options),t.each(s,function(e,s){return t.isFunction(s)?(r[e]=function(){function t(){return i.prototype[e].apply(this,arguments)}function n(t){return i.prototype[e].apply(this,t)}return function(){var e,i=this._super,o=this._superApply;return this._super=t,this._superApply=n,e=s.apply(this,arguments),this._super=i,this._superApply=o,e}}(),void 0):(r[e]=s,void 0)}),o.prototype=t.widget.extend(a,{widgetEventPrefix:n?a.widgetEventPrefix||e:e},r,{constructor:o,namespace:h,widgetName:e,widgetFullName:l}),n?(t.each(n._childConstructors,function(e,i){var s=i.prototype;t.widget(s.namespace+"."+s.widgetName,o,i._proto)}),delete n._childConstructors):i._childConstructors.push(o),t.widget.bridge(e,o),o},t.widget.extend=function(e){for(var i,s,n=l.call(arguments,1),o=0,a=n.length;a>o;o++)for(i in n[o])s=n[o][i],n[o].hasOwnProperty(i)&&void 0!==s&&(e[i]=t.isPlainObject(s)?t.isPlainObject(e[i])?t.widget.extend({},e[i],s):t.widget.extend({},s):s);return e},t.widget.bridge=function(e,i){var s=i.prototype.widgetFullName||e;t.fn[e]=function(n){var o="string"==typeof n,a=l.call(arguments,1),r=this;return o?this.length||"instance"!==n?this.each(function(){var i,o=t.data(this,s);return"instance"===n?(r=o,!1):o?t.isFunction(o[n])&&"_"!==n.charAt(0)?(i=o[n].apply(o,a),i!==o&&void 0!==i?(r=i&&i.jquery?r.pushStack(i.get()):i,!1):void 0):t.error("no such method '"+n+"' for "+e+" widget instance"):t.error("cannot call methods on "+e+" prior to initialization; "+"attempted to call method '"+n+"'")}):r=void 0:(a.length&&(n=t.widget.extend.apply(null,[n].concat(a))),this.each(function(){var e=t.data(this,s);e?(e.option(n||{}),e._init&&e._init()):t.data(this,s,new i(n,this))})),r}},t.Widget=function(){},t.Widget._childConstructors=[],t.Widget.prototype={widgetName:"widget",widgetEventPrefix:"",defaultElement:"<div>",options:{classes:{},disabled:!1,create:null},_createWidget:function(e,i){i=t(i||this.defaultElement||this)[0],this.element=t(i),this.uuid=h++,this.eventNamespace="."+this.widgetName+this.uuid,this.bindings=t(),this.hoverable=t(),this.focusable=t(),this.classesElementLookup={},i!==this&&(t.data(i,this.widgetFullName,this),this._on(!0,this.element,{remove:function(t){t.target===i&&this.destroy()}}),this.document=t(i.style?i.ownerDocument:i.document||i),this.window=t(this.document[0].defaultView||this.document[0].parentWindow)),this.options=t.widget.extend({},this.options,this._getCreateOptions(),e),this._create(),this.options.disabled&&this._setOptionDisabled(this.options.disabled),this._trigger("create",null,this._getCreateEventData()),this._init()},_getCreateOptions:function(){return{}},_getCreateEventData:t.noop,_create:t.noop,_init:t.noop,destroy:function(){var e=this;this._destroy(),t.each(this.classesElementLookup,function(t,i){e._removeClass(i,t)}),this.element.off(this.eventNamespace).removeData(this.widgetFullName),this.widget().off(this.eventNamespace).removeAttr("aria-disabled"),this.bindings.off(this.eventNamespace)},_destroy:t.noop,widget:function(){return this.element},option:function(e,i){var s,n,o,a=e;if(0===arguments.length)return t.widget.extend({},this.options);if("string"==typeof e)if(a={},s=e.split("."),e=s.shift(),s.length){for(n=a[e]=t.widget.extend({},this.options[e]),o=0;s.length-1>o;o++)n[s[o]]=n[s[o]]||{},n=n[s[o]];if(e=s.pop(),1===arguments.length)return void 0===n[e]?null:n[e];n[e]=i}else{if(1===arguments.length)return void 0===this.options[e]?null:this.options[e];a[e]=i}return this._setOptions(a),this},_setOptions:function(t){var e;for(e in t)this._setOption(e,t[e]);return this},_setOption:function(t,e){return"classes"===t&&this._setOptionClasses(e),this.options[t]=e,"disabled"===t&&this._setOptionDisabled(e),this},_setOptionClasses:function(e){var i,s,n;for(i in e)n=this.classesElementLookup[i],e[i]!==this.options.classes[i]&&n&&n.length&&(s=t(n.get()),this._removeClass(n,i),s.addClass(this._classes({element:s,keys:i,classes:e,add:!0})))},_setOptionDisabled:function(t){this._toggleClass(this.widget(),this.widgetFullName+"-disabled",null,!!t),t&&(this._removeClass(this.hoverable,null,"ui-state-hover"),this._removeClass(this.focusable,null,"ui-state-focus"))},enable:function(){return this._setOptions({disabled:!1})},disable:function(){return this._setOptions({disabled:!0})},_classes:function(e){function i(i,o){var a,r;for(r=0;i.length>r;r++)a=n.classesElementLookup[i[r]]||t(),a=e.add?t(t.unique(a.get().concat(e.element.get()))):t(a.not(e.element).get()),n.classesElementLookup[i[r]]=a,s.push(i[r]),o&&e.classes[i[r]]&&s.push(e.classes[i[r]])}var s=[],n=this;return e=t.extend({element:this.element,classes:this.options.classes||{}},e),this._on(e.element,{remove:"_untrackClassesElement"}),e.keys&&i(e.keys.match(/\S+/g)||[],!0),e.extra&&i(e.extra.match(/\S+/g)||[]),s.join(" ")},_untrackClassesElement:function(e){var i=this;t.each(i.classesElementLookup,function(s,n){-1!==t.inArray(e.target,n)&&(i.classesElementLookup[s]=t(n.not(e.target).get()))})},_removeClass:function(t,e,i){return this._toggleClass(t,e,i,!1)},_addClass:function(t,e,i){return this._toggleClass(t,e,i,!0)},_toggleClass:function(t,e,i,s){s="boolean"==typeof s?s:i;var n="string"==typeof t||null===t,o={extra:n?e:i,keys:n?t:e,element:n?this.element:t,add:s};return o.element.toggleClass(this._classes(o),s),this},_on:function(e,i,s){var n,o=this;"boolean"!=typeof e&&(s=i,i=e,e=!1),s?(i=n=t(i),this.bindings=this.bindings.add(i)):(s=i,i=this.element,n=this.widget()),t.each(s,function(s,a){function r(){return e||o.options.disabled!==!0&&!t(this).hasClass("ui-state-disabled")?("string"==typeof a?o[a]:a).apply(o,arguments):void 0}"string"!=typeof a&&(r.guid=a.guid=a.guid||r.guid||t.guid++);var h=s.match(/^([\w:-]*)\s*(.*)$/),l=h[1]+o.eventNamespace,c=h[2];c?n.on(l,c,r):i.on(l,r)})},_off:function(e,i){i=(i||"").split(" ").join(this.eventNamespace+" ")+this.eventNamespace,e.off(i).off(i),this.bindings=t(this.bindings.not(e).get()),this.focusable=t(this.focusable.not(e).get()),this.hoverable=t(this.hoverable.not(e).get())},_delay:function(t,e){function i(){return("string"==typeof t?s[t]:t).apply(s,arguments)}var s=this;return setTimeout(i,e||0)},_hoverable:function(e){this.hoverable=this.hoverable.add(e),this._on(e,{mouseenter:function(e){this._addClass(t(e.currentTarget),null,"ui-state-hover")},mouseleave:function(e){this._removeClass(t(e.currentTarget),null,"ui-state-hover")}})},_focusable:function(e){this.focusable=this.focusable.add(e),this._on(e,{focusin:function(e){this._addClass(t(e.currentTarget),null,"ui-state-focus")},focusout:function(e){this._removeClass(t(e.currentTarget),null,"ui-state-focus")}})},_trigger:function(e,i,s){var n,o,a=this.options[e];if(s=s||{},i=t.Event(i),i.type=(e===this.widgetEventPrefix?e:this.widgetEventPrefix+e).toLowerCase(),i.target=this.element[0],o=i.originalEvent)for(n in o)n in i||(i[n]=o[n]);return this.element.trigger(i,s),!(t.isFunction(a)&&a.apply(this.element[0],[i].concat(s))===!1||i.isDefaultPrevented())}},t.each({show:"fadeIn",hide:"fadeOut"},function(e,i){t.Widget.prototype["_"+e]=function(s,n,o){"string"==typeof n&&(n={effect:n});var a,r=n?n===!0||"number"==typeof n?i:n.effect||i:e;n=n||{},"number"==typeof n&&(n={duration:n}),a=!t.isEmptyObject(n),n.complete=o,n.delay&&s.delay(n.delay),a&&t.effects&&t.effects.effect[r]?s[e](n):r!==e&&s[r]?s[r](n.duration,n.easing,o):s.queue(function(i){t(this)[e](),o&&o.call(s[0]),i()})}}),t.widget,function(){function e(t,e,i){return[parseFloat(t[0])*(u.test(t[0])?e/100:1),parseFloat(t[1])*(u.test(t[1])?i/100:1)]}function i(e,i){return parseInt(t.css(e,i),10)||0}function s(e){var i=e[0];return 9===i.nodeType?{width:e.width(),height:e.height(),offset:{top:0,left:0}}:t.isWindow(i)?{width:e.width(),height:e.height(),offset:{top:e.scrollTop(),left:e.scrollLeft()}}:i.preventDefault?{width:0,height:0,offset:{top:i.pageY,left:i.pageX}}:{width:e.outerWidth(),height:e.outerHeight(),offset:e.offset()}}var n,o=Math.max,a=Math.abs,r=/left|center|right/,h=/top|center|bottom/,l=/[\+\-]\d+(\.[\d]+)?%?/,c=/^\w+/,u=/%$/,d=t.fn.position;t.position={scrollbarWidth:function(){if(void 0!==n)return n;var e,i,s=t("<div style='display:block;position:absolute;width:50px;height:50px;overflow:hidden;'><div style='height:100px;width:auto;'></div></div>"),o=s.children()[0];return t("body").append(s),e=o.offsetWidth,s.css("overflow","scroll"),i=o.offsetWidth,e===i&&(i=s[0].clientWidth),s.remove(),n=e-i},getScrollInfo:function(e){var i=e.isWindow||e.isDocument?"":e.element.css("overflow-x"),s=e.isWindow||e.isDocument?"":e.element.css("overflow-y"),n="scroll"===i||"auto"===i&&e.width<e.element[0].scrollWidth,o="scroll"===s||"auto"===s&&e.height<e.element[0].scrollHeight;return{width:o?t.position.scrollbarWidth():0,height:n?t.position.scrollbarWidth():0}},getWithinInfo:function(e){var i=t(e||window),s=t.isWindow(i[0]),n=!!i[0]&&9===i[0].nodeType,o=!s&&!n;return{element:i,isWindow:s,isDocument:n,offset:o?t(e).offset():{left:0,top:0},scrollLeft:i.scrollLeft(),scrollTop:i.scrollTop(),width:i.outerWidth(),height:i.outerHeight()}}},t.fn.position=function(n){if(!n||!n.of)return d.apply(this,arguments);n=t.extend({},n);var u,p,f,g,m,_,v=t(n.of),b=t.position.getWithinInfo(n.within),y=t.position.getScrollInfo(b),w=(n.collision||"flip").split(" "),k={};return _=s(v),v[0].preventDefault&&(n.at="left top"),p=_.width,f=_.height,g=_.offset,m=t.extend({},g),t.each(["my","at"],function(){var t,e,i=(n[this]||"").split(" ");1===i.length&&(i=r.test(i[0])?i.concat(["center"]):h.test(i[0])?["center"].concat(i):["center","center"]),i[0]=r.test(i[0])?i[0]:"center",i[1]=h.test(i[1])?i[1]:"center",t=l.exec(i[0]),e=l.exec(i[1]),k[this]=[t?t[0]:0,e?e[0]:0],n[this]=[c.exec(i[0])[0],c.exec(i[1])[0]]}),1===w.length&&(w[1]=w[0]),"right"===n.at[0]?m.left+=p:"center"===n.at[0]&&(m.left+=p/2),"bottom"===n.at[1]?m.top+=f:"center"===n.at[1]&&(m.top+=f/2),u=e(k.at,p,f),m.left+=u[0],m.top+=u[1],this.each(function(){var s,r,h=t(this),l=h.outerWidth(),c=h.outerHeight(),d=i(this,"marginLeft"),_=i(this,"marginTop"),x=l+d+i(this,"marginRight")+y.width,C=c+_+i(this,"marginBottom")+y.height,D=t.extend({},m),I=e(k.my,h.outerWidth(),h.outerHeight());"right"===n.my[0]?D.left-=l:"center"===n.my[0]&&(D.left-=l/2),"bottom"===n.my[1]?D.top-=c:"center"===n.my[1]&&(D.top-=c/2),D.left+=I[0],D.top+=I[1],s={marginLeft:d,marginTop:_},t.each(["left","top"],function(e,i){t.ui.position[w[e]]&&t.ui.position[w[e]][i](D,{targetWidth:p,targetHeight:f,elemWidth:l,elemHeight:c,collisionPosition:s,collisionWidth:x,collisionHeight:C,offset:[u[0]+I[0],u[1]+I[1]],my:n.my,at:n.at,within:b,elem:h})}),n.using&&(r=function(t){var e=g.left-D.left,i=e+p-l,s=g.top-D.top,r=s+f-c,u={target:{element:v,left:g.left,top:g.top,width:p,height:f},element:{element:h,left:D.left,top:D.top,width:l,height:c},horizontal:0>i?"left":e>0?"right":"center",vertical:0>r?"top":s>0?"bottom":"middle"};l>p&&p>a(e+i)&&(u.horizontal="center"),c>f&&f>a(s+r)&&(u.vertical="middle"),u.important=o(a(e),a(i))>o(a(s),a(r))?"horizontal":"vertical",n.using.call(this,t,u)}),h.offset(t.extend(D,{using:r}))})},t.ui.position={fit:{left:function(t,e){var i,s=e.within,n=s.isWindow?s.scrollLeft:s.offset.left,a=s.width,r=t.left-e.collisionPosition.marginLeft,h=n-r,l=r+e.collisionWidth-a-n;e.collisionWidth>a?h>0&&0>=l?(i=t.left+h+e.collisionWidth-a-n,t.left+=h-i):t.left=l>0&&0>=h?n:h>l?n+a-e.collisionWidth:n:h>0?t.left+=h:l>0?t.left-=l:t.left=o(t.left-r,t.left)},top:function(t,e){var i,s=e.within,n=s.isWindow?s.scrollTop:s.offset.top,a=e.within.height,r=t.top-e.collisionPosition.marginTop,h=n-r,l=r+e.collisionHeight-a-n;e.collisionHeight>a?h>0&&0>=l?(i=t.top+h+e.collisionHeight-a-n,t.top+=h-i):t.top=l>0&&0>=h?n:h>l?n+a-e.collisionHeight:n:h>0?t.top+=h:l>0?t.top-=l:t.top=o(t.top-r,t.top)}},flip:{left:function(t,e){var i,s,n=e.within,o=n.offset.left+n.scrollLeft,r=n.width,h=n.isWindow?n.scrollLeft:n.offset.left,l=t.left-e.collisionPosition.marginLeft,c=l-h,u=l+e.collisionWidth-r-h,d="left"===e.my[0]?-e.elemWidth:"right"===e.my[0]?e.elemWidth:0,p="left"===e.at[0]?e.targetWidth:"right"===e.at[0]?-e.targetWidth:0,f=-2*e.offset[0];0>c?(i=t.left+d+p+f+e.collisionWidth-r-o,(0>i||a(c)>i)&&(t.left+=d+p+f)):u>0&&(s=t.left-e.collisionPosition.marginLeft+d+p+f-h,(s>0||u>a(s))&&(t.left+=d+p+f))},top:function(t,e){var i,s,n=e.within,o=n.offset.top+n.scrollTop,r=n.height,h=n.isWindow?n.scrollTop:n.offset.top,l=t.top-e.collisionPosition.marginTop,c=l-h,u=l+e.collisionHeight-r-h,d="top"===e.my[1],p=d?-e.elemHeight:"bottom"===e.my[1]?e.elemHeight:0,f="top"===e.at[1]?e.targetHeight:"bottom"===e.at[1]?-e.targetHeight:0,g=-2*e.offset[1];0>c?(s=t.top+p+f+g+e.collisionHeight-r-o,(0>s||a(c)>s)&&(t.top+=p+f+g)):u>0&&(i=t.top-e.collisionPosition.marginTop+p+f+g-h,(i>0||u>a(i))&&(t.top+=p+f+g))}},flipfit:{left:function(){t.ui.position.flip.left.apply(this,arguments),t.ui.position.fit.left.apply(this,arguments)},top:function(){t.ui.position.flip.top.apply(this,arguments),t.ui.position.fit.top.apply(this,arguments)}}}}(),t.ui.position,t.extend(t.expr[":"],{data:t.expr.createPseudo?t.expr.createPseudo(function(e){return function(i){return!!t.data(i,e)}}):function(e,i,s){return!!t.data(e,s[3])}}),t.fn.extend({disableSelection:function(){var t="onselectstart"in document.createElement("div")?"selectstart":"mousedown";return function(){return this.on(t+".ui-disableSelection",function(t){t.preventDefault()})}}(),enableSelection:function(){return this.off(".ui-disableSelection")}});var c="ui-effects-",u="ui-effects-style",d="ui-effects-animated",p=t;t.effects={effect:{}},function(t,e){function i(t,e,i){var s=u[e.type]||{};return null==t?i||!e.def?null:e.def:(t=s.floor?~~t:parseFloat(t),isNaN(t)?e.def:s.mod?(t+s.mod)%s.mod:0>t?0:t>s.max?s.max:t)}function s(i){var s=l(),n=s._rgba=[];return i=i.toLowerCase(),f(h,function(t,o){var a,r=o.re.exec(i),h=r&&o.parse(r),l=o.space||"rgba";return h?(a=s[l](h),s[c[l].cache]=a[c[l].cache],n=s._rgba=a._rgba,!1):e}),n.length?("0,0,0,0"===n.join()&&t.extend(n,o.transparent),s):o[i]}function n(t,e,i){return i=(i+1)%1,1>6*i?t+6*(e-t)*i:1>2*i?e:2>3*i?t+6*(e-t)*(2/3-i):t}var o,a="backgroundColor borderBottomColor borderLeftColor borderRightColor borderTopColor color columnRuleColor outlineColor textDecorationColor textEmphasisColor",r=/^([\-+])=\s*(\d+\.?\d*)/,h=[{re:/rgba?\(\s*(\d{1,3})\s*,\s*(\d{1,3})\s*,\s*(\d{1,3})\s*(?:,\s*(\d?(?:\.\d+)?)\s*)?\)/,parse:function(t){return[t[1],t[2],t[3],t[4]]}},{re:/rgba?\(\s*(\d+(?:\.\d+)?)\%\s*,\s*(\d+(?:\.\d+)?)\%\s*,\s*(\d+(?:\.\d+)?)\%\s*(?:,\s*(\d?(?:\.\d+)?)\s*)?\)/,parse:function(t){return[2.55*t[1],2.55*t[2],2.55*t[3],t[4]]}},{re:/#([a-f0-9]{2})([a-f0-9]{2})([a-f0-9]{2})/,parse:function(t){return[parseInt(t[1],16),parseInt(t[2],16),parseInt(t[3],16)]}},{re:/#([a-f0-9])([a-f0-9])([a-f0-9])/,parse:function(t){return[parseInt(t[1]+t[1],16),parseInt(t[2]+t[2],16),parseInt(t[3]+t[3],16)]}},{re:/hsla?\(\s*(\d+(?:\.\d+)?)\s*,\s*(\d+(?:\.\d+)?)\%\s*,\s*(\d+(?:\.\d+)?)\%\s*(?:,\s*(\d?(?:\.\d+)?)\s*)?\)/,space:"hsla",parse:function(t){return[t[1],t[2]/100,t[3]/100,t[4]]}}],l=t.Color=function(e,i,s,n){return new t.Color.fn.parse(e,i,s,n)},c={rgba:{props:{red:{idx:0,type:"byte"},green:{idx:1,type:"byte"},blue:{idx:2,type:"byte"}}},hsla:{props:{hue:{idx:0,type:"degrees"},saturation:{idx:1,type:"percent"},lightness:{idx:2,type:"percent"}}}},u={"byte":{floor:!0,max:255},percent:{max:1},degrees:{mod:360,floor:!0}},d=l.support={},p=t("<p>")[0],f=t.each;p.style.cssText="background-color:rgba(1,1,1,.5)",d.rgba=p.style.backgroundColor.indexOf("rgba")>-1,f(c,function(t,e){e.cache="_"+t,e.props.alpha={idx:3,type:"percent",def:1}}),l.fn=t.extend(l.prototype,{parse:function(n,a,r,h){if(n===e)return this._rgba=[null,null,null,null],this;(n.jquery||n.nodeType)&&(n=t(n).css(a),a=e);var u=this,d=t.type(n),p=this._rgba=[];return a!==e&&(n=[n,a,r,h],d="array"),"string"===d?this.parse(s(n)||o._default):"array"===d?(f(c.rgba.props,function(t,e){p[e.idx]=i(n[e.idx],e)}),this):"object"===d?(n instanceof l?f(c,function(t,e){n[e.cache]&&(u[e.cache]=n[e.cache].slice())}):f(c,function(e,s){var o=s.cache;f(s.props,function(t,e){if(!u[o]&&s.to){if("alpha"===t||null==n[t])return;u[o]=s.to(u._rgba)}u[o][e.idx]=i(n[t],e,!0)}),u[o]&&0>t.inArray(null,u[o].slice(0,3))&&(u[o][3]=1,s.from&&(u._rgba=s.from(u[o])))}),this):e},is:function(t){var i=l(t),s=!0,n=this;return f(c,function(t,o){var a,r=i[o.cache];return r&&(a=n[o.cache]||o.to&&o.to(n._rgba)||[],f(o.props,function(t,i){return null!=r[i.idx]?s=r[i.idx]===a[i.idx]:e})),s}),s},_space:function(){var t=[],e=this;return f(c,function(i,s){e[s.cache]&&t.push(i)}),t.pop()},transition:function(t,e){var s=l(t),n=s._space(),o=c[n],a=0===this.alpha()?l("transparent"):this,r=a[o.cache]||o.to(a._rgba),h=r.slice();return s=s[o.cache],f(o.props,function(t,n){var o=n.idx,a=r[o],l=s[o],c=u[n.type]||{};null!==l&&(null===a?h[o]=l:(c.mod&&(l-a>c.mod/2?a+=c.mod:a-l>c.mod/2&&(a-=c.mod)),h[o]=i((l-a)*e+a,n)))}),this[n](h)},blend:function(e){if(1===this._rgba[3])return this;var i=this._rgba.slice(),s=i.pop(),n=l(e)._rgba;return l(t.map(i,function(t,e){return(1-s)*n[e]+s*t}))},toRgbaString:function(){var e="rgba(",i=t.map(this._rgba,function(t,e){return null==t?e>2?1:0:t});return 1===i[3]&&(i.pop(),e="rgb("),e+i.join()+")"},toHslaString:function(){var e="hsla(",i=t.map(this.hsla(),function(t,e){return null==t&&(t=e>2?1:0),e&&3>e&&(t=Math.round(100*t)+"%"),t});return 1===i[3]&&(i.pop(),e="hsl("),e+i.join()+")"},toHexString:function(e){var i=this._rgba.slice(),s=i.pop();return e&&i.push(~~(255*s)),"#"+t.map(i,function(t){return t=(t||0).toString(16),1===t.length?"0"+t:t}).join("")},toString:function(){return 0===this._rgba[3]?"transparent":this.toRgbaString()}}),l.fn.parse.prototype=l.fn,c.hsla.to=function(t){if(null==t[0]||null==t[1]||null==t[2])return[null,null,null,t[3]];var e,i,s=t[0]/255,n=t[1]/255,o=t[2]/255,a=t[3],r=Math.max(s,n,o),h=Math.min(s,n,o),l=r-h,c=r+h,u=.5*c;return e=h===r?0:s===r?60*(n-o)/l+360:n===r?60*(o-s)/l+120:60*(s-n)/l+240,i=0===l?0:.5>=u?l/c:l/(2-c),[Math.round(e)%360,i,u,null==a?1:a]},c.hsla.from=function(t){if(null==t[0]||null==t[1]||null==t[2])return[null,null,null,t[3]];var e=t[0]/360,i=t[1],s=t[2],o=t[3],a=.5>=s?s*(1+i):s+i-s*i,r=2*s-a;return[Math.round(255*n(r,a,e+1/3)),Math.round(255*n(r,a,e)),Math.round(255*n(r,a,e-1/3)),o]},f(c,function(s,n){var o=n.props,a=n.cache,h=n.to,c=n.from;l.fn[s]=function(s){if(h&&!this[a]&&(this[a]=h(this._rgba)),s===e)return this[a].slice();var n,r=t.type(s),u="array"===r||"object"===r?s:arguments,d=this[a].slice();return f(o,function(t,e){var s=u["object"===r?t:e.idx];null==s&&(s=d[e.idx]),d[e.idx]=i(s,e)}),c?(n=l(c(d)),n[a]=d,n):l(d)},f(o,function(e,i){l.fn[e]||(l.fn[e]=function(n){var o,a=t.type(n),h="alpha"===e?this._hsla?"hsla":"rgba":s,l=this[h](),c=l[i.idx];return"undefined"===a?c:("function"===a&&(n=n.call(this,c),a=t.type(n)),null==n&&i.empty?this:("string"===a&&(o=r.exec(n),o&&(n=c+parseFloat(o[2])*("+"===o[1]?1:-1))),l[i.idx]=n,this[h](l)))})})}),l.hook=function(e){var i=e.split(" ");f(i,function(e,i){t.cssHooks[i]={set:function(e,n){var o,a,r="";if("transparent"!==n&&("string"!==t.type(n)||(o=s(n)))){if(n=l(o||n),!d.rgba&&1!==n._rgba[3]){for(a="backgroundColor"===i?e.parentNode:e;(""===r||"transparent"===r)&&a&&a.style;)try{r=t.css(a,"backgroundColor"),a=a.parentNode}catch(h){}n=n.blend(r&&"transparent"!==r?r:"_default")}n=n.toRgbaString()}try{e.style[i]=n}catch(h){}}},t.fx.step[i]=function(e){e.colorInit||(e.start=l(e.elem,i),e.end=l(e.end),e.colorInit=!0),t.cssHooks[i].set(e.elem,e.start.transition(e.end,e.pos))}})},l.hook(a),t.cssHooks.borderColor={expand:function(t){var e={};return f(["Top","Right","Bottom","Left"],function(i,s){e["border"+s+"Color"]=t}),e}},o=t.Color.names={aqua:"#00ffff",black:"#000000",blue:"#0000ff",fuchsia:"#ff00ff",gray:"#808080",green:"#008000",lime:"#00ff00",maroon:"#800000",navy:"#000080",olive:"#808000",purple:"#800080",red:"#ff0000",silver:"#c0c0c0",teal:"#008080",white:"#ffffff",yellow:"#ffff00",transparent:[null,null,null,0],_default:"#ffffff"}}(p),function(){function e(e){var i,s,n=e.ownerDocument.defaultView?e.ownerDocument.defaultView.getComputedStyle(e,null):e.currentStyle,o={};if(n&&n.length&&n[0]&&n[n[0]])for(s=n.length;s--;)i=n[s],"string"==typeof n[i]&&(o[t.camelCase(i)]=n[i]);else for(i in n)"string"==typeof n[i]&&(o[i]=n[i]);return o}function i(e,i){var s,o,a={};for(s in i)o=i[s],e[s]!==o&&(n[s]||(t.fx.step[s]||!isNaN(parseFloat(o)))&&(a[s]=o));return a}var s=["add","remove","toggle"],n={border:1,borderBottom:1,borderColor:1,borderLeft:1,borderRight:1,borderTop:1,borderWidth:1,margin:1,padding:1};t.each(["borderLeftStyle","borderRightStyle","borderBottomStyle","borderTopStyle"],function(e,i){t.fx.step[i]=function(t){("none"!==t.end&&!t.setAttr||1===t.pos&&!t.setAttr)&&(p.style(t.elem,i,t.end),t.setAttr=!0)}}),t.fn.addBack||(t.fn.addBack=function(t){return this.add(null==t?this.prevObject:this.prevObject.filter(t))}),t.effects.animateClass=function(n,o,a,r){var h=t.speed(o,a,r);return this.queue(function(){var o,a=t(this),r=a.attr("class")||"",l=h.children?a.find("*").addBack():a;l=l.map(function(){var i=t(this);return{el:i,start:e(this)}}),o=function(){t.each(s,function(t,e){n[e]&&a[e+"Class"](n[e])})},o(),l=l.map(function(){return this.end=e(this.el[0]),this.diff=i(this.start,this.end),this}),a.attr("class",r),l=l.map(function(){var e=this,i=t.Deferred(),s=t.extend({},h,{queue:!1,complete:function(){i.resolve(e)}});return this.el.animate(this.diff,s),i.promise()}),t.when.apply(t,l.get()).done(function(){o(),t.each(arguments,function(){var e=this.el;t.each(this.diff,function(t){e.css(t,"")})}),h.complete.call(a[0])})})},t.fn.extend({addClass:function(e){return function(i,s,n,o){return s?t.effects.animateClass.call(this,{add:i},s,n,o):e.apply(this,arguments)}}(t.fn.addClass),removeClass:function(e){return function(i,s,n,o){return arguments.length>1?t.effects.animateClass.call(this,{remove:i},s,n,o):e.apply(this,arguments)}}(t.fn.removeClass),toggleClass:function(e){return function(i,s,n,o,a){return"boolean"==typeof s||void 0===s?n?t.effects.animateClass.call(this,s?{add:i}:{remove:i},n,o,a):e.apply(this,arguments):t.effects.animateClass.call(this,{toggle:i},s,n,o)}}(t.fn.toggleClass),switchClass:function(e,i,s,n,o){return t.effects.animateClass.call(this,{add:i,remove:e},s,n,o)}})}(),function(){function e(e,i,s,n){return t.isPlainObject(e)&&(i=e,e=e.effect),e={effect:e},null==i&&(i={}),t.isFunction(i)&&(n=i,s=null,i={}),("number"==typeof i||t.fx.speeds[i])&&(n=s,s=i,i={}),t.isFunction(s)&&(n=s,s=null),i&&t.extend(e,i),s=s||i.duration,e.duration=t.fx.off?0:"number"==typeof s?s:s in t.fx.speeds?t.fx.speeds[s]:t.fx.speeds._default,e.complete=n||i.complete,e}function i(e){return!e||"number"==typeof e||t.fx.speeds[e]?!0:"string"!=typeof e||t.effects.effect[e]?t.isFunction(e)?!0:"object"!=typeof e||e.effect?!1:!0:!0}function s(t,e){var i=e.outerWidth(),s=e.outerHeight(),n=/^rect\((-?\d*\.?\d*px|-?\d+%|auto),?\s*(-?\d*\.?\d*px|-?\d+%|auto),?\s*(-?\d*\.?\d*px|-?\d+%|auto),?\s*(-?\d*\.?\d*px|-?\d+%|auto)\)$/,o=n.exec(t)||["",0,i,s,0];return{top:parseFloat(o[1])||0,right:"auto"===o[2]?i:parseFloat(o[2]),bottom:"auto"===o[3]?s:parseFloat(o[3]),left:parseFloat(o[4])||0}}t.expr&&t.expr.filters&&t.expr.filters.animated&&(t.expr.filters.animated=function(e){return function(i){return!!t(i).data(d)||e(i)}}(t.expr.filters.animated)),t.uiBackCompat!==!1&&t.extend(t.effects,{save:function(t,e){for(var i=0,s=e.length;s>i;i++)null!==e[i]&&t.data(c+e[i],t[0].style[e[i]])},restore:function(t,e){for(var i,s=0,n=e.length;n>s;s++)null!==e[s]&&(i=t.data(c+e[s]),t.css(e[s],i))},setMode:function(t,e){return"toggle"===e&&(e=t.is(":hidden")?"show":"hide"),e},createWrapper:function(e){if(e.parent().is(".ui-effects-wrapper"))return e.parent();var i={width:e.outerWidth(!0),height:e.outerHeight(!0),"float":e.css("float")},s=t("<div></div>").addClass("ui-effects-wrapper").css({fontSize:"100%",background:"transparent",border:"none",margin:0,padding:0}),n={width:e.width(),height:e.height()},o=document.activeElement;try{o.id}catch(a){o=document.body}return e.wrap(s),(e[0]===o||t.contains(e[0],o))&&t(o).trigger("focus"),s=e.parent(),"static"===e.css("position")?(s.css({position:"relative"}),e.css({position:"relative"})):(t.extend(i,{position:e.css("position"),zIndex:e.css("z-index")}),t.each(["top","left","bottom","right"],function(t,s){i[s]=e.css(s),isNaN(parseInt(i[s],10))&&(i[s]="auto")}),e.css({position:"relative",top:0,left:0,right:"auto",bottom:"auto"})),e.css(n),s.css(i).show()},removeWrapper:function(e){var i=document.activeElement;return e.parent().is(".ui-effects-wrapper")&&(e.parent().replaceWith(e),(e[0]===i||t.contains(e[0],i))&&t(i).trigger("focus")),e}}),t.extend(t.effects,{version:"1.12.1",define:function(e,i,s){return s||(s=i,i="effect"),t.effects.effect[e]=s,t.effects.effect[e].mode=i,s},scaledDimensions:function(t,e,i){if(0===e)return{height:0,width:0,outerHeight:0,outerWidth:0};var s="horizontal"!==i?(e||100)/100:1,n="vertical"!==i?(e||100)/100:1;return{height:t.height()*n,width:t.width()*s,outerHeight:t.outerHeight()*n,outerWidth:t.outerWidth()*s}},clipToBox:function(t){return{width:t.clip.right-t.clip.left,height:t.clip.bottom-t.clip.top,left:t.clip.left,top:t.clip.top}},unshift:function(t,e,i){var s=t.queue();e>1&&s.splice.apply(s,[1,0].concat(s.splice(e,i))),t.dequeue()},saveStyle:function(t){t.data(u,t[0].style.cssText)},restoreStyle:function(t){t[0].style.cssText=t.data(u)||"",t.removeData(u)},mode:function(t,e){var i=t.is(":hidden");return"toggle"===e&&(e=i?"show":"hide"),(i?"hide"===e:"show"===e)&&(e="none"),e},getBaseline:function(t,e){var i,s;switch(t[0]){case"top":i=0;break;case"middle":i=.5;break;case"bottom":i=1;break;default:i=t[0]/e.height}switch(t[1]){case"left":s=0;break;case"center":s=.5;break;case"right":s=1;break;default:s=t[1]/e.width}return{x:s,y:i}},createPlaceholder:function(e){var i,s=e.css("position"),n=e.position();return e.css({marginTop:e.css("marginTop"),marginBottom:e.css("marginBottom"),marginLeft:e.css("marginLeft"),marginRight:e.css("marginRight")}).outerWidth(e.outerWidth()).outerHeight(e.outerHeight()),/^(static|relative)/.test(s)&&(s="absolute",i=t("<"+e[0].nodeName+">").insertAfter(e).css({display:/^(inline|ruby)/.test(e.css("display"))?"inline-block":"block",visibility:"hidden",marginTop:e.css("marginTop"),marginBottom:e.css("marginBottom"),marginLeft:e.css("marginLeft"),marginRight:e.css("marginRight"),"float":e.css("float")}).outerWidth(e.outerWidth()).outerHeight(e.outerHeight()).addClass("ui-effects-placeholder"),e.data(c+"placeholder",i)),e.css({position:s,left:n.left,top:n.top}),i},removePlaceholder:function(t){var e=c+"placeholder",i=t.data(e);i&&(i.remove(),t.removeData(e))},cleanUp:function(e){t.effects.restoreStyle(e),t.effects.removePlaceholder(e)},setTransition:function(e,i,s,n){return n=n||{},t.each(i,function(t,i){var o=e.cssUnit(i);o[0]>0&&(n[i]=o[0]*s+o[1])}),n}}),t.fn.extend({effect:function(){function i(e){function i(){r.removeData(d),t.effects.cleanUp(r),"hide"===s.mode&&r.hide(),a()}function a(){t.isFunction(h)&&h.call(r[0]),t.isFunction(e)&&e()}var r=t(this);s.mode=c.shift(),t.uiBackCompat===!1||o?"none"===s.mode?(r[l](),a()):n.call(r[0],s,i):(r.is(":hidden")?"hide"===l:"show"===l)?(r[l](),a()):n.call(r[0],s,a)}var s=e.apply(this,arguments),n=t.effects.effect[s.effect],o=n.mode,a=s.queue,r=a||"fx",h=s.complete,l=s.mode,c=[],u=function(e){var i=t(this),s=t.effects.mode(i,l)||o;i.data(d,!0),c.push(s),o&&("show"===s||s===o&&"hide"===s)&&i.show(),o&&"none"===s||t.effects.saveStyle(i),t.isFunction(e)&&e()};return t.fx.off||!n?l?this[l](s.duration,h):this.each(function(){h&&h.call(this)}):a===!1?this.each(u).each(i):this.queue(r,u).queue(r,i)},show:function(t){return function(s){if(i(s))return t.apply(this,arguments);var n=e.apply(this,arguments);return n.mode="show",this.effect.call(this,n)
}}(t.fn.show),hide:function(t){return function(s){if(i(s))return t.apply(this,arguments);var n=e.apply(this,arguments);return n.mode="hide",this.effect.call(this,n)}}(t.fn.hide),toggle:function(t){return function(s){if(i(s)||"boolean"==typeof s)return t.apply(this,arguments);var n=e.apply(this,arguments);return n.mode="toggle",this.effect.call(this,n)}}(t.fn.toggle),cssUnit:function(e){var i=this.css(e),s=[];return t.each(["em","px","%","pt"],function(t,e){i.indexOf(e)>0&&(s=[parseFloat(i),e])}),s},cssClip:function(t){return t?this.css("clip","rect("+t.top+"px "+t.right+"px "+t.bottom+"px "+t.left+"px)"):s(this.css("clip"),this)},transfer:function(e,i){var s=t(this),n=t(e.to),o="fixed"===n.css("position"),a=t("body"),r=o?a.scrollTop():0,h=o?a.scrollLeft():0,l=n.offset(),c={top:l.top-r,left:l.left-h,height:n.innerHeight(),width:n.innerWidth()},u=s.offset(),d=t("<div class='ui-effects-transfer'></div>").appendTo("body").addClass(e.className).css({top:u.top-r,left:u.left-h,height:s.innerHeight(),width:s.innerWidth(),position:o?"fixed":"absolute"}).animate(c,e.duration,e.easing,function(){d.remove(),t.isFunction(i)&&i()})}}),t.fx.step.clip=function(e){e.clipInit||(e.start=t(e.elem).cssClip(),"string"==typeof e.end&&(e.end=s(e.end,e.elem)),e.clipInit=!0),t(e.elem).cssClip({top:e.pos*(e.end.top-e.start.top)+e.start.top,right:e.pos*(e.end.right-e.start.right)+e.start.right,bottom:e.pos*(e.end.bottom-e.start.bottom)+e.start.bottom,left:e.pos*(e.end.left-e.start.left)+e.start.left})}}(),function(){var e={};t.each(["Quad","Cubic","Quart","Quint","Expo"],function(t,i){e[i]=function(e){return Math.pow(e,t+2)}}),t.extend(e,{Sine:function(t){return 1-Math.cos(t*Math.PI/2)},Circ:function(t){return 1-Math.sqrt(1-t*t)},Elastic:function(t){return 0===t||1===t?t:-Math.pow(2,8*(t-1))*Math.sin((80*(t-1)-7.5)*Math.PI/15)},Back:function(t){return t*t*(3*t-2)},Bounce:function(t){for(var e,i=4;((e=Math.pow(2,--i))-1)/11>t;);return 1/Math.pow(4,3-i)-7.5625*Math.pow((3*e-2)/22-t,2)}}),t.each(e,function(e,i){t.easing["easeIn"+e]=i,t.easing["easeOut"+e]=function(t){return 1-i(1-t)},t.easing["easeInOut"+e]=function(t){return.5>t?i(2*t)/2:1-i(-2*t+2)/2}})}();var f=t.effects;t.effects.define("blind","hide",function(e,i){var s={up:["bottom","top"],vertical:["bottom","top"],down:["top","bottom"],left:["right","left"],horizontal:["right","left"],right:["left","right"]},n=t(this),o=e.direction||"up",a=n.cssClip(),r={clip:t.extend({},a)},h=t.effects.createPlaceholder(n);r.clip[s[o][0]]=r.clip[s[o][1]],"show"===e.mode&&(n.cssClip(r.clip),h&&h.css(t.effects.clipToBox(r)),r.clip=a),h&&h.animate(t.effects.clipToBox(r),e.duration,e.easing),n.animate(r,{queue:!1,duration:e.duration,easing:e.easing,complete:i})}),t.effects.define("bounce",function(e,i){var s,n,o,a=t(this),r=e.mode,h="hide"===r,l="show"===r,c=e.direction||"up",u=e.distance,d=e.times||5,p=2*d+(l||h?1:0),f=e.duration/p,g=e.easing,m="up"===c||"down"===c?"top":"left",_="up"===c||"left"===c,v=0,b=a.queue().length;for(t.effects.createPlaceholder(a),o=a.css(m),u||(u=a["top"===m?"outerHeight":"outerWidth"]()/3),l&&(n={opacity:1},n[m]=o,a.css("opacity",0).css(m,_?2*-u:2*u).animate(n,f,g)),h&&(u/=Math.pow(2,d-1)),n={},n[m]=o;d>v;v++)s={},s[m]=(_?"-=":"+=")+u,a.animate(s,f,g).animate(n,f,g),u=h?2*u:u/2;h&&(s={opacity:0},s[m]=(_?"-=":"+=")+u,a.animate(s,f,g)),a.queue(i),t.effects.unshift(a,b,p+1)}),t.effects.define("clip","hide",function(e,i){var s,n={},o=t(this),a=e.direction||"vertical",r="both"===a,h=r||"horizontal"===a,l=r||"vertical"===a;s=o.cssClip(),n.clip={top:l?(s.bottom-s.top)/2:s.top,right:h?(s.right-s.left)/2:s.right,bottom:l?(s.bottom-s.top)/2:s.bottom,left:h?(s.right-s.left)/2:s.left},t.effects.createPlaceholder(o),"show"===e.mode&&(o.cssClip(n.clip),n.clip=s),o.animate(n,{queue:!1,duration:e.duration,easing:e.easing,complete:i})}),t.effects.define("drop","hide",function(e,i){var s,n=t(this),o=e.mode,a="show"===o,r=e.direction||"left",h="up"===r||"down"===r?"top":"left",l="up"===r||"left"===r?"-=":"+=",c="+="===l?"-=":"+=",u={opacity:0};t.effects.createPlaceholder(n),s=e.distance||n["top"===h?"outerHeight":"outerWidth"](!0)/2,u[h]=l+s,a&&(n.css(u),u[h]=c+s,u.opacity=1),n.animate(u,{queue:!1,duration:e.duration,easing:e.easing,complete:i})}),t.effects.define("explode","hide",function(e,i){function s(){b.push(this),b.length===u*d&&n()}function n(){p.css({visibility:"visible"}),t(b).remove(),i()}var o,a,r,h,l,c,u=e.pieces?Math.round(Math.sqrt(e.pieces)):3,d=u,p=t(this),f=e.mode,g="show"===f,m=p.show().css("visibility","hidden").offset(),_=Math.ceil(p.outerWidth()/d),v=Math.ceil(p.outerHeight()/u),b=[];for(o=0;u>o;o++)for(h=m.top+o*v,c=o-(u-1)/2,a=0;d>a;a++)r=m.left+a*_,l=a-(d-1)/2,p.clone().appendTo("body").wrap("<div></div>").css({position:"absolute",visibility:"visible",left:-a*_,top:-o*v}).parent().addClass("ui-effects-explode").css({position:"absolute",overflow:"hidden",width:_,height:v,left:r+(g?l*_:0),top:h+(g?c*v:0),opacity:g?0:1}).animate({left:r+(g?0:l*_),top:h+(g?0:c*v),opacity:g?1:0},e.duration||500,e.easing,s)}),t.effects.define("fade","toggle",function(e,i){var s="show"===e.mode;t(this).css("opacity",s?0:1).animate({opacity:s?1:0},{queue:!1,duration:e.duration,easing:e.easing,complete:i})}),t.effects.define("fold","hide",function(e,i){var s=t(this),n=e.mode,o="show"===n,a="hide"===n,r=e.size||15,h=/([0-9]+)%/.exec(r),l=!!e.horizFirst,c=l?["right","bottom"]:["bottom","right"],u=e.duration/2,d=t.effects.createPlaceholder(s),p=s.cssClip(),f={clip:t.extend({},p)},g={clip:t.extend({},p)},m=[p[c[0]],p[c[1]]],_=s.queue().length;h&&(r=parseInt(h[1],10)/100*m[a?0:1]),f.clip[c[0]]=r,g.clip[c[0]]=r,g.clip[c[1]]=0,o&&(s.cssClip(g.clip),d&&d.css(t.effects.clipToBox(g)),g.clip=p),s.queue(function(i){d&&d.animate(t.effects.clipToBox(f),u,e.easing).animate(t.effects.clipToBox(g),u,e.easing),i()}).animate(f,u,e.easing).animate(g,u,e.easing).queue(i),t.effects.unshift(s,_,4)}),t.effects.define("highlight","show",function(e,i){var s=t(this),n={backgroundColor:s.css("backgroundColor")};"hide"===e.mode&&(n.opacity=0),t.effects.saveStyle(s),s.css({backgroundImage:"none",backgroundColor:e.color||"#ffff99"}).animate(n,{queue:!1,duration:e.duration,easing:e.easing,complete:i})}),t.effects.define("size",function(e,i){var s,n,o,a=t(this),r=["fontSize"],h=["borderTopWidth","borderBottomWidth","paddingTop","paddingBottom"],l=["borderLeftWidth","borderRightWidth","paddingLeft","paddingRight"],c=e.mode,u="effect"!==c,d=e.scale||"both",p=e.origin||["middle","center"],f=a.css("position"),g=a.position(),m=t.effects.scaledDimensions(a),_=e.from||m,v=e.to||t.effects.scaledDimensions(a,0);t.effects.createPlaceholder(a),"show"===c&&(o=_,_=v,v=o),n={from:{y:_.height/m.height,x:_.width/m.width},to:{y:v.height/m.height,x:v.width/m.width}},("box"===d||"both"===d)&&(n.from.y!==n.to.y&&(_=t.effects.setTransition(a,h,n.from.y,_),v=t.effects.setTransition(a,h,n.to.y,v)),n.from.x!==n.to.x&&(_=t.effects.setTransition(a,l,n.from.x,_),v=t.effects.setTransition(a,l,n.to.x,v))),("content"===d||"both"===d)&&n.from.y!==n.to.y&&(_=t.effects.setTransition(a,r,n.from.y,_),v=t.effects.setTransition(a,r,n.to.y,v)),p&&(s=t.effects.getBaseline(p,m),_.top=(m.outerHeight-_.outerHeight)*s.y+g.top,_.left=(m.outerWidth-_.outerWidth)*s.x+g.left,v.top=(m.outerHeight-v.outerHeight)*s.y+g.top,v.left=(m.outerWidth-v.outerWidth)*s.x+g.left),a.css(_),("content"===d||"both"===d)&&(h=h.concat(["marginTop","marginBottom"]).concat(r),l=l.concat(["marginLeft","marginRight"]),a.find("*[width]").each(function(){var i=t(this),s=t.effects.scaledDimensions(i),o={height:s.height*n.from.y,width:s.width*n.from.x,outerHeight:s.outerHeight*n.from.y,outerWidth:s.outerWidth*n.from.x},a={height:s.height*n.to.y,width:s.width*n.to.x,outerHeight:s.height*n.to.y,outerWidth:s.width*n.to.x};n.from.y!==n.to.y&&(o=t.effects.setTransition(i,h,n.from.y,o),a=t.effects.setTransition(i,h,n.to.y,a)),n.from.x!==n.to.x&&(o=t.effects.setTransition(i,l,n.from.x,o),a=t.effects.setTransition(i,l,n.to.x,a)),u&&t.effects.saveStyle(i),i.css(o),i.animate(a,e.duration,e.easing,function(){u&&t.effects.restoreStyle(i)})})),a.animate(v,{queue:!1,duration:e.duration,easing:e.easing,complete:function(){var e=a.offset();0===v.opacity&&a.css("opacity",_.opacity),u||(a.css("position","static"===f?"relative":f).offset(e),t.effects.saveStyle(a)),i()}})}),t.effects.define("scale",function(e,i){var s=t(this),n=e.mode,o=parseInt(e.percent,10)||(0===parseInt(e.percent,10)?0:"effect"!==n?0:100),a=t.extend(!0,{from:t.effects.scaledDimensions(s),to:t.effects.scaledDimensions(s,o,e.direction||"both"),origin:e.origin||["middle","center"]},e);e.fade&&(a.from.opacity=1,a.to.opacity=0),t.effects.effect.size.call(this,a,i)}),t.effects.define("puff","hide",function(e,i){var s=t.extend(!0,{},e,{fade:!0,percent:parseInt(e.percent,10)||150});t.effects.effect.scale.call(this,s,i)}),t.effects.define("pulsate","show",function(e,i){var s=t(this),n=e.mode,o="show"===n,a="hide"===n,r=o||a,h=2*(e.times||5)+(r?1:0),l=e.duration/h,c=0,u=1,d=s.queue().length;for((o||!s.is(":visible"))&&(s.css("opacity",0).show(),c=1);h>u;u++)s.animate({opacity:c},l,e.easing),c=1-c;s.animate({opacity:c},l,e.easing),s.queue(i),t.effects.unshift(s,d,h+1)}),t.effects.define("shake",function(e,i){var s=1,n=t(this),o=e.direction||"left",a=e.distance||20,r=e.times||3,h=2*r+1,l=Math.round(e.duration/h),c="up"===o||"down"===o?"top":"left",u="up"===o||"left"===o,d={},p={},f={},g=n.queue().length;for(t.effects.createPlaceholder(n),d[c]=(u?"-=":"+=")+a,p[c]=(u?"+=":"-=")+2*a,f[c]=(u?"-=":"+=")+2*a,n.animate(d,l,e.easing);r>s;s++)n.animate(p,l,e.easing).animate(f,l,e.easing);n.animate(p,l,e.easing).animate(d,l/2,e.easing).queue(i),t.effects.unshift(n,g,h+1)}),t.effects.define("slide","show",function(e,i){var s,n,o=t(this),a={up:["bottom","top"],down:["top","bottom"],left:["right","left"],right:["left","right"]},r=e.mode,h=e.direction||"left",l="up"===h||"down"===h?"top":"left",c="up"===h||"left"===h,u=e.distance||o["top"===l?"outerHeight":"outerWidth"](!0),d={};t.effects.createPlaceholder(o),s=o.cssClip(),n=o.position()[l],d[l]=(c?-1:1)*u+n,d.clip=o.cssClip(),d.clip[a[h][1]]=d.clip[a[h][0]],"show"===r&&(o.cssClip(d.clip),o.css(l,d[l]),d.clip=s,d[l]=n),o.animate(d,{queue:!1,duration:e.duration,easing:e.easing,complete:i})});var f;t.uiBackCompat!==!1&&(f=t.effects.define("transfer",function(e,i){t(this).transfer(e,i)})),t.ui.focusable=function(i,s){var n,o,a,r,h,l=i.nodeName.toLowerCase();return"area"===l?(n=i.parentNode,o=n.name,i.href&&o&&"map"===n.nodeName.toLowerCase()?(a=t("img[usemap='#"+o+"']"),a.length>0&&a.is(":visible")):!1):(/^(input|select|textarea|button|object)$/.test(l)?(r=!i.disabled,r&&(h=t(i).closest("fieldset")[0],h&&(r=!h.disabled))):r="a"===l?i.href||s:s,r&&t(i).is(":visible")&&e(t(i)))},t.extend(t.expr[":"],{focusable:function(e){return t.ui.focusable(e,null!=t.attr(e,"tabindex"))}}),t.ui.focusable,t.fn.form=function(){return"string"==typeof this[0].form?this.closest("form"):t(this[0].form)},t.ui.formResetMixin={_formResetHandler:function(){var e=t(this);setTimeout(function(){var i=e.data("ui-form-reset-instances");t.each(i,function(){this.refresh()})})},_bindFormResetHandler:function(){if(this.form=this.element.form(),this.form.length){var t=this.form.data("ui-form-reset-instances")||[];t.length||this.form.on("reset.ui-form-reset",this._formResetHandler),t.push(this),this.form.data("ui-form-reset-instances",t)}},_unbindFormResetHandler:function(){if(this.form.length){var e=this.form.data("ui-form-reset-instances");e.splice(t.inArray(this,e),1),e.length?this.form.data("ui-form-reset-instances",e):this.form.removeData("ui-form-reset-instances").off("reset.ui-form-reset")}}},"1.7"===t.fn.jquery.substring(0,3)&&(t.each(["Width","Height"],function(e,i){function s(e,i,s,o){return t.each(n,function(){i-=parseFloat(t.css(e,"padding"+this))||0,s&&(i-=parseFloat(t.css(e,"border"+this+"Width"))||0),o&&(i-=parseFloat(t.css(e,"margin"+this))||0)}),i}var n="Width"===i?["Left","Right"]:["Top","Bottom"],o=i.toLowerCase(),a={innerWidth:t.fn.innerWidth,innerHeight:t.fn.innerHeight,outerWidth:t.fn.outerWidth,outerHeight:t.fn.outerHeight};t.fn["inner"+i]=function(e){return void 0===e?a["inner"+i].call(this):this.each(function(){t(this).css(o,s(this,e)+"px")})},t.fn["outer"+i]=function(e,n){return"number"!=typeof e?a["outer"+i].call(this,e):this.each(function(){t(this).css(o,s(this,e,!0,n)+"px")})}}),t.fn.addBack=function(t){return this.add(null==t?this.prevObject:this.prevObject.filter(t))}),t.ui.keyCode={BACKSPACE:8,COMMA:188,DELETE:46,DOWN:40,END:35,ENTER:13,ESCAPE:27,HOME:36,LEFT:37,PAGE_DOWN:34,PAGE_UP:33,PERIOD:190,RIGHT:39,SPACE:32,TAB:9,UP:38},t.ui.escapeSelector=function(){var t=/([!"#$%&'()*+,.\/:;<=>?@[\]^`{|}~])/g;return function(e){return e.replace(t,"\\$1")}}(),t.fn.labels=function(){var e,i,s,n,o;return this[0].labels&&this[0].labels.length?this.pushStack(this[0].labels):(n=this.eq(0).parents("label"),s=this.attr("id"),s&&(e=this.eq(0).parents().last(),o=e.add(e.length?e.siblings():this.siblings()),i="label[for='"+t.ui.escapeSelector(s)+"']",n=n.add(o.find(i).addBack(i))),this.pushStack(n))},t.fn.scrollParent=function(e){var i=this.css("position"),s="absolute"===i,n=e?/(auto|scroll|hidden)/:/(auto|scroll)/,o=this.parents().filter(function(){var e=t(this);return s&&"static"===e.css("position")?!1:n.test(e.css("overflow")+e.css("overflow-y")+e.css("overflow-x"))}).eq(0);return"fixed"!==i&&o.length?o:t(this[0].ownerDocument||document)},t.extend(t.expr[":"],{tabbable:function(e){var i=t.attr(e,"tabindex"),s=null!=i;return(!s||i>=0)&&t.ui.focusable(e,s)}}),t.fn.extend({uniqueId:function(){var t=0;return function(){return this.each(function(){this.id||(this.id="ui-id-"+ ++t)})}}(),removeUniqueId:function(){return this.each(function(){/^ui-id-\d+$/.test(this.id)&&t(this).removeAttr("id")})}}),t.widget("ui.accordion",{version:"1.12.1",options:{active:0,animate:{},classes:{"ui-accordion-header":"ui-corner-top","ui-accordion-header-collapsed":"ui-corner-all","ui-accordion-content":"ui-corner-bottom"},collapsible:!1,event:"click",header:"> li > :first-child, > :not(li):even",heightStyle:"auto",icons:{activeHeader:"ui-icon-triangle-1-s",header:"ui-icon-triangle-1-e"},activate:null,beforeActivate:null},hideProps:{borderTopWidth:"hide",borderBottomWidth:"hide",paddingTop:"hide",paddingBottom:"hide",height:"hide"},showProps:{borderTopWidth:"show",borderBottomWidth:"show",paddingTop:"show",paddingBottom:"show",height:"show"},_create:function(){var e=this.options;this.prevShow=this.prevHide=t(),this._addClass("ui-accordion","ui-widget ui-helper-reset"),this.element.attr("role","tablist"),e.collapsible||e.active!==!1&&null!=e.active||(e.active=0),this._processPanels(),0>e.active&&(e.active+=this.headers.length),this._refresh()},_getCreateEventData:function(){return{header:this.active,panel:this.active.length?this.active.next():t()}},_createIcons:function(){var e,i,s=this.options.icons;s&&(e=t("<span>"),this._addClass(e,"ui-accordion-header-icon","ui-icon "+s.header),e.prependTo(this.headers),i=this.active.children(".ui-accordion-header-icon"),this._removeClass(i,s.header)._addClass(i,null,s.activeHeader)._addClass(this.headers,"ui-accordion-icons"))},_destroyIcons:function(){this._removeClass(this.headers,"ui-accordion-icons"),this.headers.children(".ui-accordion-header-icon").remove()},_destroy:function(){var t;this.element.removeAttr("role"),this.headers.removeAttr("role aria-expanded aria-selected aria-controls tabIndex").removeUniqueId(),this._destroyIcons(),t=this.headers.next().css("display","").removeAttr("role aria-hidden aria-labelledby").removeUniqueId(),"content"!==this.options.heightStyle&&t.css("height","")},_setOption:function(t,e){return"active"===t?(this._activate(e),void 0):("event"===t&&(this.options.event&&this._off(this.headers,this.options.event),this._setupEvents(e)),this._super(t,e),"collapsible"!==t||e||this.options.active!==!1||this._activate(0),"icons"===t&&(this._destroyIcons(),e&&this._createIcons()),void 0)},_setOptionDisabled:function(t){this._super(t),this.element.attr("aria-disabled",t),this._toggleClass(null,"ui-state-disabled",!!t),this._toggleClass(this.headers.add(this.headers.next()),null,"ui-state-disabled",!!t)},_keydown:function(e){if(!e.altKey&&!e.ctrlKey){var i=t.ui.keyCode,s=this.headers.length,n=this.headers.index(e.target),o=!1;switch(e.keyCode){case i.RIGHT:case i.DOWN:o=this.headers[(n+1)%s];break;case i.LEFT:case i.UP:o=this.headers[(n-1+s)%s];break;case i.SPACE:case i.ENTER:this._eventHandler(e);break;case i.HOME:o=this.headers[0];break;case i.END:o=this.headers[s-1]}o&&(t(e.target).attr("tabIndex",-1),t(o).attr("tabIndex",0),t(o).trigger("focus"),e.preventDefault())}},_panelKeyDown:function(e){e.keyCode===t.ui.keyCode.UP&&e.ctrlKey&&t(e.currentTarget).prev().trigger("focus")},refresh:function(){var e=this.options;this._processPanels(),e.active===!1&&e.collapsible===!0||!this.headers.length?(e.active=!1,this.active=t()):e.active===!1?this._activate(0):this.active.length&&!t.contains(this.element[0],this.active[0])?this.headers.length===this.headers.find(".ui-state-disabled").length?(e.active=!1,this.active=t()):this._activate(Math.max(0,e.active-1)):e.active=this.headers.index(this.active),this._destroyIcons(),this._refresh()},_processPanels:function(){var t=this.headers,e=this.panels;this.headers=this.element.find(this.options.header),this._addClass(this.headers,"ui-accordion-header ui-accordion-header-collapsed","ui-state-default"),this.panels=this.headers.next().filter(":not(.ui-accordion-content-active)").hide(),this._addClass(this.panels,"ui-accordion-content","ui-helper-reset ui-widget-content"),e&&(this._off(t.not(this.headers)),this._off(e.not(this.panels)))},_refresh:function(){var e,i=this.options,s=i.heightStyle,n=this.element.parent();this.active=this._findActive(i.active),this._addClass(this.active,"ui-accordion-header-active","ui-state-active")._removeClass(this.active,"ui-accordion-header-collapsed"),this._addClass(this.active.next(),"ui-accordion-content-active"),this.active.next().show(),this.headers.attr("role","tab").each(function(){var e=t(this),i=e.uniqueId().attr("id"),s=e.next(),n=s.uniqueId().attr("id");e.attr("aria-controls",n),s.attr("aria-labelledby",i)}).next().attr("role","tabpanel"),this.headers.not(this.active).attr({"aria-selected":"false","aria-expanded":"false",tabIndex:-1}).next().attr({"aria-hidden":"true"}).hide(),this.active.length?this.active.attr({"aria-selected":"true","aria-expanded":"true",tabIndex:0}).next().attr({"aria-hidden":"false"}):this.headers.eq(0).attr("tabIndex",0),this._createIcons(),this._setupEvents(i.event),"fill"===s?(e=n.height(),this.element.siblings(":visible").each(function(){var i=t(this),s=i.css("position");"absolute"!==s&&"fixed"!==s&&(e-=i.outerHeight(!0))}),this.headers.each(function(){e-=t(this).outerHeight(!0)}),this.headers.next().each(function(){t(this).height(Math.max(0,e-t(this).innerHeight()+t(this).height()))}).css("overflow","auto")):"auto"===s&&(e=0,this.headers.next().each(function(){var i=t(this).is(":visible");i||t(this).show(),e=Math.max(e,t(this).css("height","").height()),i||t(this).hide()}).height(e))},_activate:function(e){var i=this._findActive(e)[0];i!==this.active[0]&&(i=i||this.active[0],this._eventHandler({target:i,currentTarget:i,preventDefault:t.noop}))},_findActive:function(e){return"number"==typeof e?this.headers.eq(e):t()},_setupEvents:function(e){var i={keydown:"_keydown"};e&&t.each(e.split(" "),function(t,e){i[e]="_eventHandler"}),this._off(this.headers.add(this.headers.next())),this._on(this.headers,i),this._on(this.headers.next(),{keydown:"_panelKeyDown"}),this._hoverable(this.headers),this._focusable(this.headers)},_eventHandler:function(e){var i,s,n=this.options,o=this.active,a=t(e.currentTarget),r=a[0]===o[0],h=r&&n.collapsible,l=h?t():a.next(),c=o.next(),u={oldHeader:o,oldPanel:c,newHeader:h?t():a,newPanel:l};e.preventDefault(),r&&!n.collapsible||this._trigger("beforeActivate",e,u)===!1||(n.active=h?!1:this.headers.index(a),this.active=r?t():a,this._toggle(u),this._removeClass(o,"ui-accordion-header-active","ui-state-active"),n.icons&&(i=o.children(".ui-accordion-header-icon"),this._removeClass(i,null,n.icons.activeHeader)._addClass(i,null,n.icons.header)),r||(this._removeClass(a,"ui-accordion-header-collapsed")._addClass(a,"ui-accordion-header-active","ui-state-active"),n.icons&&(s=a.children(".ui-accordion-header-icon"),this._removeClass(s,null,n.icons.header)._addClass(s,null,n.icons.activeHeader)),this._addClass(a.next(),"ui-accordion-content-active")))},_toggle:function(e){var i=e.newPanel,s=this.prevShow.length?this.prevShow:e.oldPanel;this.prevShow.add(this.prevHide).stop(!0,!0),this.prevShow=i,this.prevHide=s,this.options.animate?this._animate(i,s,e):(s.hide(),i.show(),this._toggleComplete(e)),s.attr({"aria-hidden":"true"}),s.prev().attr({"aria-selected":"false","aria-expanded":"false"}),i.length&&s.length?s.prev().attr({tabIndex:-1,"aria-expanded":"false"}):i.length&&this.headers.filter(function(){return 0===parseInt(t(this).attr("tabIndex"),10)}).attr("tabIndex",-1),i.attr("aria-hidden","false").prev().attr({"aria-selected":"true","aria-expanded":"true",tabIndex:0})},_animate:function(t,e,i){var s,n,o,a=this,r=0,h=t.css("box-sizing"),l=t.length&&(!e.length||t.index()<e.index()),c=this.options.animate||{},u=l&&c.down||c,d=function(){a._toggleComplete(i)};return"number"==typeof u&&(o=u),"string"==typeof u&&(n=u),n=n||u.easing||c.easing,o=o||u.duration||c.duration,e.length?t.length?(s=t.show().outerHeight(),e.animate(this.hideProps,{duration:o,easing:n,step:function(t,e){e.now=Math.round(t)}}),t.hide().animate(this.showProps,{duration:o,easing:n,complete:d,step:function(t,i){i.now=Math.round(t),"height"!==i.prop?"content-box"===h&&(r+=i.now):"content"!==a.options.heightStyle&&(i.now=Math.round(s-e.outerHeight()-r),r=0)}}),void 0):e.animate(this.hideProps,o,n,d):t.animate(this.showProps,o,n,d)},_toggleComplete:function(t){var e=t.oldPanel,i=e.prev();this._removeClass(e,"ui-accordion-content-active"),this._removeClass(i,"ui-accordion-header-active")._addClass(i,"ui-accordion-header-collapsed"),e.length&&(e.parent()[0].className=e.parent()[0].className),this._trigger("activate",null,t)}}),t.ui.safeActiveElement=function(t){var e;try{e=t.activeElement}catch(i){e=t.body}return e||(e=t.body),e.nodeName||(e=t.body),e},t.widget("ui.menu",{version:"1.12.1",defaultElement:"<ul>",delay:300,options:{icons:{submenu:"ui-icon-caret-1-e"},items:"> *",menus:"ul",position:{my:"left top",at:"right top"},role:"menu",blur:null,focus:null,select:null},_create:function(){this.activeMenu=this.element,this.mouseHandled=!1,this.element.uniqueId().attr({role:this.options.role,tabIndex:0}),this._addClass("ui-menu","ui-widget ui-widget-content"),this._on({"mousedown .ui-menu-item":function(t){t.preventDefault()},"click .ui-menu-item":function(e){var i=t(e.target),s=t(t.ui.safeActiveElement(this.document[0]));!this.mouseHandled&&i.not(".ui-state-disabled").length&&(this.select(e),e.isPropagationStopped()||(this.mouseHandled=!0),i.has(".ui-menu").length?this.expand(e):!this.element.is(":focus")&&s.closest(".ui-menu").length&&(this.element.trigger("focus",[!0]),this.active&&1===this.active.parents(".ui-menu").length&&clearTimeout(this.timer)))},"mouseenter .ui-menu-item":function(e){if(!this.previousFilter){var i=t(e.target).closest(".ui-menu-item"),s=t(e.currentTarget);i[0]===s[0]&&(this._removeClass(s.siblings().children(".ui-state-active"),null,"ui-state-active"),this.focus(e,s))}},mouseleave:"collapseAll","mouseleave .ui-menu":"collapseAll",focus:function(t,e){var i=this.active||this.element.find(this.options.items).eq(0);e||this.focus(t,i)},blur:function(e){this._delay(function(){var i=!t.contains(this.element[0],t.ui.safeActiveElement(this.document[0]));i&&this.collapseAll(e)})},keydown:"_keydown"}),this.refresh(),this._on(this.document,{click:function(t){this._closeOnDocumentClick(t)&&this.collapseAll(t),this.mouseHandled=!1}})},_destroy:function(){var e=this.element.find(".ui-menu-item").removeAttr("role aria-disabled"),i=e.children(".ui-menu-item-wrapper").removeUniqueId().removeAttr("tabIndex role aria-haspopup");this.element.removeAttr("aria-activedescendant").find(".ui-menu").addBack().removeAttr("role aria-labelledby aria-expanded aria-hidden aria-disabled tabIndex").removeUniqueId().show(),i.children().each(function(){var e=t(this);e.data("ui-menu-submenu-caret")&&e.remove()})},_keydown:function(e){var i,s,n,o,a=!0;switch(e.keyCode){case t.ui.keyCode.PAGE_UP:this.previousPage(e);break;case t.ui.keyCode.PAGE_DOWN:this.nextPage(e);break;case t.ui.keyCode.HOME:this._move("first","first",e);break;case t.ui.keyCode.END:this._move("last","last",e);break;case t.ui.keyCode.UP:this.previous(e);break;case t.ui.keyCode.DOWN:this.next(e);break;case t.ui.keyCode.LEFT:this.collapse(e);break;case t.ui.keyCode.RIGHT:this.active&&!this.active.is(".ui-state-disabled")&&this.expand(e);break;case t.ui.keyCode.ENTER:case t.ui.keyCode.SPACE:this._activate(e);break;case t.ui.keyCode.ESCAPE:this.collapse(e);break;default:a=!1,s=this.previousFilter||"",o=!1,n=e.keyCode>=96&&105>=e.keyCode?""+(e.keyCode-96):String.fromCharCode(e.keyCode),clearTimeout(this.filterTimer),n===s?o=!0:n=s+n,i=this._filterMenuItems(n),i=o&&-1!==i.index(this.active.next())?this.active.nextAll(".ui-menu-item"):i,i.length||(n=String.fromCharCode(e.keyCode),i=this._filterMenuItems(n)),i.length?(this.focus(e,i),this.previousFilter=n,this.filterTimer=this._delay(function(){delete this.previousFilter},1e3)):delete this.previousFilter}a&&e.preventDefault()},_activate:function(t){this.active&&!this.active.is(".ui-state-disabled")&&(this.active.children("[aria-haspopup='true']").length?this.expand(t):this.select(t))},refresh:function(){var e,i,s,n,o,a=this,r=this.options.icons.submenu,h=this.element.find(this.options.menus);this._toggleClass("ui-menu-icons",null,!!this.element.find(".ui-icon").length),s=h.filter(":not(.ui-menu)").hide().attr({role:this.options.role,"aria-hidden":"true","aria-expanded":"false"}).each(function(){var e=t(this),i=e.prev(),s=t("<span>").data("ui-menu-submenu-caret",!0);a._addClass(s,"ui-menu-icon","ui-icon "+r),i.attr("aria-haspopup","true").prepend(s),e.attr("aria-labelledby",i.attr("id"))}),this._addClass(s,"ui-menu","ui-widget ui-widget-content ui-front"),e=h.add(this.element),i=e.find(this.options.items),i.not(".ui-menu-item").each(function(){var e=t(this);a._isDivider(e)&&a._addClass(e,"ui-menu-divider","ui-widget-content")}),n=i.not(".ui-menu-item, .ui-menu-divider"),o=n.children().not(".ui-menu").uniqueId().attr({tabIndex:-1,role:this._itemRole()}),this._addClass(n,"ui-menu-item")._addClass(o,"ui-menu-item-wrapper"),i.filter(".ui-state-disabled").attr("aria-disabled","true"),this.active&&!t.contains(this.element[0],this.active[0])&&this.blur()},_itemRole:function(){return{menu:"menuitem",listbox:"option"}[this.options.role]},_setOption:function(t,e){if("icons"===t){var i=this.element.find(".ui-menu-icon");this._removeClass(i,null,this.options.icons.submenu)._addClass(i,null,e.submenu)}this._super(t,e)},_setOptionDisabled:function(t){this._super(t),this.element.attr("aria-disabled",t+""),this._toggleClass(null,"ui-state-disabled",!!t)},focus:function(t,e){var i,s,n;this.blur(t,t&&"focus"===t.type),this._scrollIntoView(e),this.active=e.first(),s=this.active.children(".ui-menu-item-wrapper"),this._addClass(s,null,"ui-state-active"),this.options.role&&this.element.attr("aria-activedescendant",s.attr("id")),n=this.active.parent().closest(".ui-menu-item").children(".ui-menu-item-wrapper"),this._addClass(n,null,"ui-state-active"),t&&"keydown"===t.type?this._close():this.timer=this._delay(function(){this._close()},this.delay),i=e.children(".ui-menu"),i.length&&t&&/^mouse/.test(t.type)&&this._startOpening(i),this.activeMenu=e.parent(),this._trigger("focus",t,{item:e})},_scrollIntoView:function(e){var i,s,n,o,a,r;this._hasScroll()&&(i=parseFloat(t.css(this.activeMenu[0],"borderTopWidth"))||0,s=parseFloat(t.css(this.activeMenu[0],"paddingTop"))||0,n=e.offset().top-this.activeMenu.offset().top-i-s,o=this.activeMenu.scrollTop(),a=this.activeMenu.height(),r=e.outerHeight(),0>n?this.activeMenu.scrollTop(o+n):n+r>a&&this.activeMenu.scrollTop(o+n-a+r))},blur:function(t,e){e||clearTimeout(this.timer),this.active&&(this._removeClass(this.active.children(".ui-menu-item-wrapper"),null,"ui-state-active"),this._trigger("blur",t,{item:this.active}),this.active=null)},_startOpening:function(t){clearTimeout(this.timer),"true"===t.attr("aria-hidden")&&(this.timer=this._delay(function(){this._close(),this._open(t)},this.delay))},_open:function(e){var i=t.extend({of:this.active},this.options.position);clearTimeout(this.timer),this.element.find(".ui-menu").not(e.parents(".ui-menu")).hide().attr("aria-hidden","true"),e.show().removeAttr("aria-hidden").attr("aria-expanded","true").position(i)},collapseAll:function(e,i){clearTimeout(this.timer),this.timer=this._delay(function(){var s=i?this.element:t(e&&e.target).closest(this.element.find(".ui-menu"));s.length||(s=this.element),this._close(s),this.blur(e),this._removeClass(s.find(".ui-state-active"),null,"ui-state-active"),this.activeMenu=s},this.delay)},_close:function(t){t||(t=this.active?this.active.parent():this.element),t.find(".ui-menu").hide().attr("aria-hidden","true").attr("aria-expanded","false")},_closeOnDocumentClick:function(e){return!t(e.target).closest(".ui-menu").length},_isDivider:function(t){return!/[^\-\u2014\u2013\s]/.test(t.text())},collapse:function(t){var e=this.active&&this.active.parent().closest(".ui-menu-item",this.element);e&&e.length&&(this._close(),this.focus(t,e))},expand:function(t){var e=this.active&&this.active.children(".ui-menu ").find(this.options.items).first();e&&e.length&&(this._open(e.parent()),this._delay(function(){this.focus(t,e)}))},next:function(t){this._move("next","first",t)},previous:function(t){this._move("prev","last",t)},isFirstItem:function(){return this.active&&!this.active.prevAll(".ui-menu-item").length},isLastItem:function(){return this.active&&!this.active.nextAll(".ui-menu-item").length},_move:function(t,e,i){var s;this.active&&(s="first"===t||"last"===t?this.active["first"===t?"prevAll":"nextAll"](".ui-menu-item").eq(-1):this.active[t+"All"](".ui-menu-item").eq(0)),s&&s.length&&this.active||(s=this.activeMenu.find(this.options.items)[e]()),this.focus(i,s)},nextPage:function(e){var i,s,n;return this.active?(this.isLastItem()||(this._hasScroll()?(s=this.active.offset().top,n=this.element.height(),this.active.nextAll(".ui-menu-item").each(function(){return i=t(this),0>i.offset().top-s-n}),this.focus(e,i)):this.focus(e,this.activeMenu.find(this.options.items)[this.active?"last":"first"]())),void 0):(this.next(e),void 0)},previousPage:function(e){var i,s,n;return this.active?(this.isFirstItem()||(this._hasScroll()?(s=this.active.offset().top,n=this.element.height(),this.active.prevAll(".ui-menu-item").each(function(){return i=t(this),i.offset().top-s+n>0}),this.focus(e,i)):this.focus(e,this.activeMenu.find(this.options.items).first())),void 0):(this.next(e),void 0)},_hasScroll:function(){return this.element.outerHeight()<this.element.prop("scrollHeight")},select:function(e){this.active=this.active||t(e.target).closest(".ui-menu-item");var i={item:this.active};this.active.has(".ui-menu").length||this.collapseAll(e,!0),this._trigger("select",e,i)},_filterMenuItems:function(e){var i=e.replace(/[\-\[\]{}()*+?.,\\\^$|#\s]/g,"\\$&"),s=RegExp("^"+i,"i");return this.activeMenu.find(this.options.items).filter(".ui-menu-item").filter(function(){return s.test(t.trim(t(this).children(".ui-menu-item-wrapper").text()))})}}),t.widget("ui.autocomplete",{version:"1.12.1",defaultElement:"<input>",options:{appendTo:null,autoFocus:!1,delay:300,minLength:1,position:{my:"left top",at:"left bottom",collision:"none"},source:null,change:null,close:null,focus:null,open:null,response:null,search:null,select:null},requestIndex:0,pending:0,_create:function(){var e,i,s,n=this.element[0].nodeName.toLowerCase(),o="textarea"===n,a="input"===n;
this.isMultiLine=o||!a&&this._isContentEditable(this.element),this.valueMethod=this.element[o||a?"val":"text"],this.isNewMenu=!0,this._addClass("ui-autocomplete-input"),this.element.attr("autocomplete","off"),this._on(this.element,{keydown:function(n){if(this.element.prop("readOnly"))return e=!0,s=!0,i=!0,void 0;e=!1,s=!1,i=!1;var o=t.ui.keyCode;switch(n.keyCode){case o.PAGE_UP:e=!0,this._move("previousPage",n);break;case o.PAGE_DOWN:e=!0,this._move("nextPage",n);break;case o.UP:e=!0,this._keyEvent("previous",n);break;case o.DOWN:e=!0,this._keyEvent("next",n);break;case o.ENTER:this.menu.active&&(e=!0,n.preventDefault(),this.menu.select(n));break;case o.TAB:this.menu.active&&this.menu.select(n);break;case o.ESCAPE:this.menu.element.is(":visible")&&(this.isMultiLine||this._value(this.term),this.close(n),n.preventDefault());break;default:i=!0,this._searchTimeout(n)}},keypress:function(s){if(e)return e=!1,(!this.isMultiLine||this.menu.element.is(":visible"))&&s.preventDefault(),void 0;if(!i){var n=t.ui.keyCode;switch(s.keyCode){case n.PAGE_UP:this._move("previousPage",s);break;case n.PAGE_DOWN:this._move("nextPage",s);break;case n.UP:this._keyEvent("previous",s);break;case n.DOWN:this._keyEvent("next",s)}}},input:function(t){return s?(s=!1,t.preventDefault(),void 0):(this._searchTimeout(t),void 0)},focus:function(){this.selectedItem=null,this.previous=this._value()},blur:function(t){return this.cancelBlur?(delete this.cancelBlur,void 0):(clearTimeout(this.searching),this.close(t),this._change(t),void 0)}}),this._initSource(),this.menu=t("<ul>").appendTo(this._appendTo()).menu({role:null}).hide().menu("instance"),this._addClass(this.menu.element,"ui-autocomplete","ui-front"),this._on(this.menu.element,{mousedown:function(e){e.preventDefault(),this.cancelBlur=!0,this._delay(function(){delete this.cancelBlur,this.element[0]!==t.ui.safeActiveElement(this.document[0])&&this.element.trigger("focus")})},menufocus:function(e,i){var s,n;return this.isNewMenu&&(this.isNewMenu=!1,e.originalEvent&&/^mouse/.test(e.originalEvent.type))?(this.menu.blur(),this.document.one("mousemove",function(){t(e.target).trigger(e.originalEvent)}),void 0):(n=i.item.data("ui-autocomplete-item"),!1!==this._trigger("focus",e,{item:n})&&e.originalEvent&&/^key/.test(e.originalEvent.type)&&this._value(n.value),s=i.item.attr("aria-label")||n.value,s&&t.trim(s).length&&(this.liveRegion.children().hide(),t("<div>").text(s).appendTo(this.liveRegion)),void 0)},menuselect:function(e,i){var s=i.item.data("ui-autocomplete-item"),n=this.previous;this.element[0]!==t.ui.safeActiveElement(this.document[0])&&(this.element.trigger("focus"),this.previous=n,this._delay(function(){this.previous=n,this.selectedItem=s})),!1!==this._trigger("select",e,{item:s})&&this._value(s.value),this.term=this._value(),this.close(e),this.selectedItem=s}}),this.liveRegion=t("<div>",{role:"status","aria-live":"assertive","aria-relevant":"additions"}).appendTo(this.document[0].body),this._addClass(this.liveRegion,null,"ui-helper-hidden-accessible"),this._on(this.window,{beforeunload:function(){this.element.removeAttr("autocomplete")}})},_destroy:function(){clearTimeout(this.searching),this.element.removeAttr("autocomplete"),this.menu.element.remove(),this.liveRegion.remove()},_setOption:function(t,e){this._super(t,e),"source"===t&&this._initSource(),"appendTo"===t&&this.menu.element.appendTo(this._appendTo()),"disabled"===t&&e&&this.xhr&&this.xhr.abort()},_isEventTargetInWidget:function(e){var i=this.menu.element[0];return e.target===this.element[0]||e.target===i||t.contains(i,e.target)},_closeOnClickOutside:function(t){this._isEventTargetInWidget(t)||this.close()},_appendTo:function(){var e=this.options.appendTo;return e&&(e=e.jquery||e.nodeType?t(e):this.document.find(e).eq(0)),e&&e[0]||(e=this.element.closest(".ui-front, dialog")),e.length||(e=this.document[0].body),e},_initSource:function(){var e,i,s=this;t.isArray(this.options.source)?(e=this.options.source,this.source=function(i,s){s(t.ui.autocomplete.filter(e,i.term))}):"string"==typeof this.options.source?(i=this.options.source,this.source=function(e,n){s.xhr&&s.xhr.abort(),s.xhr=t.ajax({url:i,data:e,dataType:"json",success:function(t){n(t)},error:function(){n([])}})}):this.source=this.options.source},_searchTimeout:function(t){clearTimeout(this.searching),this.searching=this._delay(function(){var e=this.term===this._value(),i=this.menu.element.is(":visible"),s=t.altKey||t.ctrlKey||t.metaKey||t.shiftKey;(!e||e&&!i&&!s)&&(this.selectedItem=null,this.search(null,t))},this.options.delay)},search:function(t,e){return t=null!=t?t:this._value(),this.term=this._value(),t.length<this.options.minLength?this.close(e):this._trigger("search",e)!==!1?this._search(t):void 0},_search:function(t){this.pending++,this._addClass("ui-autocomplete-loading"),this.cancelSearch=!1,this.source({term:t},this._response())},_response:function(){var e=++this.requestIndex;return t.proxy(function(t){e===this.requestIndex&&this.__response(t),this.pending--,this.pending||this._removeClass("ui-autocomplete-loading")},this)},__response:function(t){t&&(t=this._normalize(t)),this._trigger("response",null,{content:t}),!this.options.disabled&&t&&t.length&&!this.cancelSearch?(this._suggest(t),this._trigger("open")):this._close()},close:function(t){this.cancelSearch=!0,this._close(t)},_close:function(t){this._off(this.document,"mousedown"),this.menu.element.is(":visible")&&(this.menu.element.hide(),this.menu.blur(),this.isNewMenu=!0,this._trigger("close",t))},_change:function(t){this.previous!==this._value()&&this._trigger("change",t,{item:this.selectedItem})},_normalize:function(e){return e.length&&e[0].label&&e[0].value?e:t.map(e,function(e){return"string"==typeof e?{label:e,value:e}:t.extend({},e,{label:e.label||e.value,value:e.value||e.label})})},_suggest:function(e){var i=this.menu.element.empty();this._renderMenu(i,e),this.isNewMenu=!0,this.menu.refresh(),i.show(),this._resizeMenu(),i.position(t.extend({of:this.element},this.options.position)),this.options.autoFocus&&this.menu.next(),this._on(this.document,{mousedown:"_closeOnClickOutside"})},_resizeMenu:function(){var t=this.menu.element;t.outerWidth(Math.max(t.width("").outerWidth()+1,this.element.outerWidth()))},_renderMenu:function(e,i){var s=this;t.each(i,function(t,i){s._renderItemData(e,i)})},_renderItemData:function(t,e){return this._renderItem(t,e).data("ui-autocomplete-item",e)},_renderItem:function(e,i){return t("<li>").append(t("<div>").text(i.label)).appendTo(e)},_move:function(t,e){return this.menu.element.is(":visible")?this.menu.isFirstItem()&&/^previous/.test(t)||this.menu.isLastItem()&&/^next/.test(t)?(this.isMultiLine||this._value(this.term),this.menu.blur(),void 0):(this.menu[t](e),void 0):(this.search(null,e),void 0)},widget:function(){return this.menu.element},_value:function(){return this.valueMethod.apply(this.element,arguments)},_keyEvent:function(t,e){(!this.isMultiLine||this.menu.element.is(":visible"))&&(this._move(t,e),e.preventDefault())},_isContentEditable:function(t){if(!t.length)return!1;var e=t.prop("contentEditable");return"inherit"===e?this._isContentEditable(t.parent()):"true"===e}}),t.extend(t.ui.autocomplete,{escapeRegex:function(t){return t.replace(/[\-\[\]{}()*+?.,\\\^$|#\s]/g,"\\$&")},filter:function(e,i){var s=RegExp(t.ui.autocomplete.escapeRegex(i),"i");return t.grep(e,function(t){return s.test(t.label||t.value||t)})}}),t.widget("ui.autocomplete",t.ui.autocomplete,{options:{messages:{noResults:"No search results.",results:function(t){return t+(t>1?" results are":" result is")+" available, use up and down arrow keys to navigate."}}},__response:function(e){var i;this._superApply(arguments),this.options.disabled||this.cancelSearch||(i=e&&e.length?this.options.messages.results(e.length):this.options.messages.noResults,this.liveRegion.children().hide(),t("<div>").text(i).appendTo(this.liveRegion))}}),t.ui.autocomplete;var g=/ui-corner-([a-z]){2,6}/g;t.widget("ui.controlgroup",{version:"1.12.1",defaultElement:"<div>",options:{direction:"horizontal",disabled:null,onlyVisible:!0,items:{button:"input[type=button], input[type=submit], input[type=reset], button, a",controlgroupLabel:".ui-controlgroup-label",checkboxradio:"input[type='checkbox'], input[type='radio']",selectmenu:"select",spinner:".ui-spinner-input"}},_create:function(){this._enhance()},_enhance:function(){this.element.attr("role","toolbar"),this.refresh()},_destroy:function(){this._callChildMethod("destroy"),this.childWidgets.removeData("ui-controlgroup-data"),this.element.removeAttr("role"),this.options.items.controlgroupLabel&&this.element.find(this.options.items.controlgroupLabel).find(".ui-controlgroup-label-contents").contents().unwrap()},_initWidgets:function(){var e=this,i=[];t.each(this.options.items,function(s,n){var o,a={};return n?"controlgroupLabel"===s?(o=e.element.find(n),o.each(function(){var e=t(this);e.children(".ui-controlgroup-label-contents").length||e.contents().wrapAll("<span class='ui-controlgroup-label-contents'></span>")}),e._addClass(o,null,"ui-widget ui-widget-content ui-state-default"),i=i.concat(o.get()),void 0):(t.fn[s]&&(a=e["_"+s+"Options"]?e["_"+s+"Options"]("middle"):{classes:{}},e.element.find(n).each(function(){var n=t(this),o=n[s]("instance"),r=t.widget.extend({},a);if("button"!==s||!n.parent(".ui-spinner").length){o||(o=n[s]()[s]("instance")),o&&(r.classes=e._resolveClassesValues(r.classes,o)),n[s](r);var h=n[s]("widget");t.data(h[0],"ui-controlgroup-data",o?o:n[s]("instance")),i.push(h[0])}})),void 0):void 0}),this.childWidgets=t(t.unique(i)),this._addClass(this.childWidgets,"ui-controlgroup-item")},_callChildMethod:function(e){this.childWidgets.each(function(){var i=t(this),s=i.data("ui-controlgroup-data");s&&s[e]&&s[e]()})},_updateCornerClass:function(t,e){var i="ui-corner-top ui-corner-bottom ui-corner-left ui-corner-right ui-corner-all",s=this._buildSimpleOptions(e,"label").classes.label;this._removeClass(t,null,i),this._addClass(t,null,s)},_buildSimpleOptions:function(t,e){var i="vertical"===this.options.direction,s={classes:{}};return s.classes[e]={middle:"",first:"ui-corner-"+(i?"top":"left"),last:"ui-corner-"+(i?"bottom":"right"),only:"ui-corner-all"}[t],s},_spinnerOptions:function(t){var e=this._buildSimpleOptions(t,"ui-spinner");return e.classes["ui-spinner-up"]="",e.classes["ui-spinner-down"]="",e},_buttonOptions:function(t){return this._buildSimpleOptions(t,"ui-button")},_checkboxradioOptions:function(t){return this._buildSimpleOptions(t,"ui-checkboxradio-label")},_selectmenuOptions:function(t){var e="vertical"===this.options.direction;return{width:e?"auto":!1,classes:{middle:{"ui-selectmenu-button-open":"","ui-selectmenu-button-closed":""},first:{"ui-selectmenu-button-open":"ui-corner-"+(e?"top":"tl"),"ui-selectmenu-button-closed":"ui-corner-"+(e?"top":"left")},last:{"ui-selectmenu-button-open":e?"":"ui-corner-tr","ui-selectmenu-button-closed":"ui-corner-"+(e?"bottom":"right")},only:{"ui-selectmenu-button-open":"ui-corner-top","ui-selectmenu-button-closed":"ui-corner-all"}}[t]}},_resolveClassesValues:function(e,i){var s={};return t.each(e,function(n){var o=i.options.classes[n]||"";o=t.trim(o.replace(g,"")),s[n]=(o+" "+e[n]).replace(/\s+/g," ")}),s},_setOption:function(t,e){return"direction"===t&&this._removeClass("ui-controlgroup-"+this.options.direction),this._super(t,e),"disabled"===t?(this._callChildMethod(e?"disable":"enable"),void 0):(this.refresh(),void 0)},refresh:function(){var e,i=this;this._addClass("ui-controlgroup ui-controlgroup-"+this.options.direction),"horizontal"===this.options.direction&&this._addClass(null,"ui-helper-clearfix"),this._initWidgets(),e=this.childWidgets,this.options.onlyVisible&&(e=e.filter(":visible")),e.length&&(t.each(["first","last"],function(t,s){var n=e[s]().data("ui-controlgroup-data");if(n&&i["_"+n.widgetName+"Options"]){var o=i["_"+n.widgetName+"Options"](1===e.length?"only":s);o.classes=i._resolveClassesValues(o.classes,n),n.element[n.widgetName](o)}else i._updateCornerClass(e[s](),s)}),this._callChildMethod("refresh"))}}),t.widget("ui.checkboxradio",[t.ui.formResetMixin,{version:"1.12.1",options:{disabled:null,label:null,icon:!0,classes:{"ui-checkboxradio-label":"ui-corner-all","ui-checkboxradio-icon":"ui-corner-all"}},_getCreateOptions:function(){var e,i,s=this,n=this._super()||{};return this._readType(),i=this.element.labels(),this.label=t(i[i.length-1]),this.label.length||t.error("No label found for checkboxradio widget"),this.originalLabel="",this.label.contents().not(this.element[0]).each(function(){s.originalLabel+=3===this.nodeType?t(this).text():this.outerHTML}),this.originalLabel&&(n.label=this.originalLabel),e=this.element[0].disabled,null!=e&&(n.disabled=e),n},_create:function(){var t=this.element[0].checked;this._bindFormResetHandler(),null==this.options.disabled&&(this.options.disabled=this.element[0].disabled),this._setOption("disabled",this.options.disabled),this._addClass("ui-checkboxradio","ui-helper-hidden-accessible"),this._addClass(this.label,"ui-checkboxradio-label","ui-button ui-widget"),"radio"===this.type&&this._addClass(this.label,"ui-checkboxradio-radio-label"),this.options.label&&this.options.label!==this.originalLabel?this._updateLabel():this.originalLabel&&(this.options.label=this.originalLabel),this._enhance(),t&&(this._addClass(this.label,"ui-checkboxradio-checked","ui-state-active"),this.icon&&this._addClass(this.icon,null,"ui-state-hover")),this._on({change:"_toggleClasses",focus:function(){this._addClass(this.label,null,"ui-state-focus ui-visual-focus")},blur:function(){this._removeClass(this.label,null,"ui-state-focus ui-visual-focus")}})},_readType:function(){var e=this.element[0].nodeName.toLowerCase();this.type=this.element[0].type,"input"===e&&/radio|checkbox/.test(this.type)||t.error("Can't create checkboxradio on element.nodeName="+e+" and element.type="+this.type)},_enhance:function(){this._updateIcon(this.element[0].checked)},widget:function(){return this.label},_getRadioGroup:function(){var e,i=this.element[0].name,s="input[name='"+t.ui.escapeSelector(i)+"']";return i?(e=this.form.length?t(this.form[0].elements).filter(s):t(s).filter(function(){return 0===t(this).form().length}),e.not(this.element)):t([])},_toggleClasses:function(){var e=this.element[0].checked;this._toggleClass(this.label,"ui-checkboxradio-checked","ui-state-active",e),this.options.icon&&"checkbox"===this.type&&this._toggleClass(this.icon,null,"ui-icon-check ui-state-checked",e)._toggleClass(this.icon,null,"ui-icon-blank",!e),"radio"===this.type&&this._getRadioGroup().each(function(){var e=t(this).checkboxradio("instance");e&&e._removeClass(e.label,"ui-checkboxradio-checked","ui-state-active")})},_destroy:function(){this._unbindFormResetHandler(),this.icon&&(this.icon.remove(),this.iconSpace.remove())},_setOption:function(t,e){return"label"!==t||e?(this._super(t,e),"disabled"===t?(this._toggleClass(this.label,null,"ui-state-disabled",e),this.element[0].disabled=e,void 0):(this.refresh(),void 0)):void 0},_updateIcon:function(e){var i="ui-icon ui-icon-background ";this.options.icon?(this.icon||(this.icon=t("<span>"),this.iconSpace=t("<span> </span>"),this._addClass(this.iconSpace,"ui-checkboxradio-icon-space")),"checkbox"===this.type?(i+=e?"ui-icon-check ui-state-checked":"ui-icon-blank",this._removeClass(this.icon,null,e?"ui-icon-blank":"ui-icon-check")):i+="ui-icon-blank",this._addClass(this.icon,"ui-checkboxradio-icon",i),e||this._removeClass(this.icon,null,"ui-icon-check ui-state-checked"),this.icon.prependTo(this.label).after(this.iconSpace)):void 0!==this.icon&&(this.icon.remove(),this.iconSpace.remove(),delete this.icon)},_updateLabel:function(){var t=this.label.contents().not(this.element[0]);this.icon&&(t=t.not(this.icon[0])),this.iconSpace&&(t=t.not(this.iconSpace[0])),t.remove(),this.label.append(this.options.label)},refresh:function(){var t=this.element[0].checked,e=this.element[0].disabled;this._updateIcon(t),this._toggleClass(this.label,"ui-checkboxradio-checked","ui-state-active",t),null!==this.options.label&&this._updateLabel(),e!==this.options.disabled&&this._setOptions({disabled:e})}}]),t.ui.checkboxradio,t.widget("ui.button",{version:"1.12.1",defaultElement:"<button>",options:{classes:{"ui-button":"ui-corner-all"},disabled:null,icon:null,iconPosition:"beginning",label:null,showLabel:!0},_getCreateOptions:function(){var t,e=this._super()||{};return this.isInput=this.element.is("input"),t=this.element[0].disabled,null!=t&&(e.disabled=t),this.originalLabel=this.isInput?this.element.val():this.element.html(),this.originalLabel&&(e.label=this.originalLabel),e},_create:function(){!this.option.showLabel&!this.options.icon&&(this.options.showLabel=!0),null==this.options.disabled&&(this.options.disabled=this.element[0].disabled||!1),this.hasTitle=!!this.element.attr("title"),this.options.label&&this.options.label!==this.originalLabel&&(this.isInput?this.element.val(this.options.label):this.element.html(this.options.label)),this._addClass("ui-button","ui-widget"),this._setOption("disabled",this.options.disabled),this._enhance(),this.element.is("a")&&this._on({keyup:function(e){e.keyCode===t.ui.keyCode.SPACE&&(e.preventDefault(),this.element[0].click?this.element[0].click():this.element.trigger("click"))}})},_enhance:function(){this.element.is("button")||this.element.attr("role","button"),this.options.icon&&(this._updateIcon("icon",this.options.icon),this._updateTooltip())},_updateTooltip:function(){this.title=this.element.attr("title"),this.options.showLabel||this.title||this.element.attr("title",this.options.label)},_updateIcon:function(e,i){var s="iconPosition"!==e,n=s?this.options.iconPosition:i,o="top"===n||"bottom"===n;this.icon?s&&this._removeClass(this.icon,null,this.options.icon):(this.icon=t("<span>"),this._addClass(this.icon,"ui-button-icon","ui-icon"),this.options.showLabel||this._addClass("ui-button-icon-only")),s&&this._addClass(this.icon,null,i),this._attachIcon(n),o?(this._addClass(this.icon,null,"ui-widget-icon-block"),this.iconSpace&&this.iconSpace.remove()):(this.iconSpace||(this.iconSpace=t("<span> </span>"),this._addClass(this.iconSpace,"ui-button-icon-space")),this._removeClass(this.icon,null,"ui-wiget-icon-block"),this._attachIconSpace(n))},_destroy:function(){this.element.removeAttr("role"),this.icon&&this.icon.remove(),this.iconSpace&&this.iconSpace.remove(),this.hasTitle||this.element.removeAttr("title")},_attachIconSpace:function(t){this.icon[/^(?:end|bottom)/.test(t)?"before":"after"](this.iconSpace)},_attachIcon:function(t){this.element[/^(?:end|bottom)/.test(t)?"append":"prepend"](this.icon)},_setOptions:function(t){var e=void 0===t.showLabel?this.options.showLabel:t.showLabel,i=void 0===t.icon?this.options.icon:t.icon;e||i||(t.showLabel=!0),this._super(t)},_setOption:function(t,e){"icon"===t&&(e?this._updateIcon(t,e):this.icon&&(this.icon.remove(),this.iconSpace&&this.iconSpace.remove())),"iconPosition"===t&&this._updateIcon(t,e),"showLabel"===t&&(this._toggleClass("ui-button-icon-only",null,!e),this._updateTooltip()),"label"===t&&(this.isInput?this.element.val(e):(this.element.html(e),this.icon&&(this._attachIcon(this.options.iconPosition),this._attachIconSpace(this.options.iconPosition)))),this._super(t,e),"disabled"===t&&(this._toggleClass(null,"ui-state-disabled",e),this.element[0].disabled=e,e&&this.element.blur())},refresh:function(){var t=this.element.is("input, button")?this.element[0].disabled:this.element.hasClass("ui-button-disabled");t!==this.options.disabled&&this._setOptions({disabled:t}),this._updateTooltip()}}),t.uiBackCompat!==!1&&(t.widget("ui.button",t.ui.button,{options:{text:!0,icons:{primary:null,secondary:null}},_create:function(){this.options.showLabel&&!this.options.text&&(this.options.showLabel=this.options.text),!this.options.showLabel&&this.options.text&&(this.options.text=this.options.showLabel),this.options.icon||!this.options.icons.primary&&!this.options.icons.secondary?this.options.icon&&(this.options.icons.primary=this.options.icon):this.options.icons.primary?this.options.icon=this.options.icons.primary:(this.options.icon=this.options.icons.secondary,this.options.iconPosition="end"),this._super()},_setOption:function(t,e){return"text"===t?(this._super("showLabel",e),void 0):("showLabel"===t&&(this.options.text=e),"icon"===t&&(this.options.icons.primary=e),"icons"===t&&(e.primary?(this._super("icon",e.primary),this._super("iconPosition","beginning")):e.secondary&&(this._super("icon",e.secondary),this._super("iconPosition","end"))),this._superApply(arguments),void 0)}}),t.fn.button=function(e){return function(){return!this.length||this.length&&"INPUT"!==this[0].tagName||this.length&&"INPUT"===this[0].tagName&&"checkbox"!==this.attr("type")&&"radio"!==this.attr("type")?e.apply(this,arguments):(t.ui.checkboxradio||t.error("Checkboxradio widget missing"),0===arguments.length?this.checkboxradio({icon:!1}):this.checkboxradio.apply(this,arguments))}}(t.fn.button),t.fn.buttonset=function(){return t.ui.controlgroup||t.error("Controlgroup widget missing"),"option"===arguments[0]&&"items"===arguments[1]&&arguments[2]?this.controlgroup.apply(this,[arguments[0],"items.button",arguments[2]]):"option"===arguments[0]&&"items"===arguments[1]?this.controlgroup.apply(this,[arguments[0],"items.button"]):("object"==typeof arguments[0]&&arguments[0].items&&(arguments[0].items={button:arguments[0].items}),this.controlgroup.apply(this,arguments))}),t.ui.button,t.extend(t.ui,{datepicker:{version:"1.12.1"}});var m;t.extend(s.prototype,{markerClassName:"hasDatepicker",maxRows:4,_widgetDatepicker:function(){return this.dpDiv},setDefaults:function(t){return a(this._defaults,t||{}),this},_attachDatepicker:function(e,i){var s,n,o;s=e.nodeName.toLowerCase(),n="div"===s||"span"===s,e.id||(this.uuid+=1,e.id="dp"+this.uuid),o=this._newInst(t(e),n),o.settings=t.extend({},i||{}),"input"===s?this._connectDatepicker(e,o):n&&this._inlineDatepicker(e,o)},_newInst:function(e,i){var s=e[0].id.replace(/([^A-Za-z0-9_\-])/g,"\\\\$1");return{id:s,input:e,selectedDay:0,selectedMonth:0,selectedYear:0,drawMonth:0,drawYear:0,inline:i,dpDiv:i?n(t("<div class='"+this._inlineClass+" ui-datepicker ui-widget ui-widget-content ui-helper-clearfix ui-corner-all'></div>")):this.dpDiv}},_connectDatepicker:function(e,i){var s=t(e);i.append=t([]),i.trigger=t([]),s.hasClass(this.markerClassName)||(this._attachments(s,i),s.addClass(this.markerClassName).on("keydown",this._doKeyDown).on("keypress",this._doKeyPress).on("keyup",this._doKeyUp),this._autoSize(i),t.data(e,"datepicker",i),i.settings.disabled&&this._disableDatepicker(e))},_attachments:function(e,i){var s,n,o,a=this._get(i,"appendText"),r=this._get(i,"isRTL");i.append&&i.append.remove(),a&&(i.append=t("<span class='"+this._appendClass+"'>"+a+"</span>"),e[r?"before":"after"](i.append)),e.off("focus",this._showDatepicker),i.trigger&&i.trigger.remove(),s=this._get(i,"showOn"),("focus"===s||"both"===s)&&e.on("focus",this._showDatepicker),("button"===s||"both"===s)&&(n=this._get(i,"buttonText"),o=this._get(i,"buttonImage"),i.trigger=t(this._get(i,"buttonImageOnly")?t("<img/>").addClass(this._triggerClass).attr({src:o,alt:n,title:n}):t("<button type='button'></button>").addClass(this._triggerClass).html(o?t("<img/>").attr({src:o,alt:n,title:n}):n)),e[r?"before":"after"](i.trigger),i.trigger.on("click",function(){return t.datepicker._datepickerShowing&&t.datepicker._lastInput===e[0]?t.datepicker._hideDatepicker():t.datepicker._datepickerShowing&&t.datepicker._lastInput!==e[0]?(t.datepicker._hideDatepicker(),t.datepicker._showDatepicker(e[0])):t.datepicker._showDatepicker(e[0]),!1}))},_autoSize:function(t){if(this._get(t,"autoSize")&&!t.inline){var e,i,s,n,o=new Date(2009,11,20),a=this._get(t,"dateFormat");a.match(/[DM]/)&&(e=function(t){for(i=0,s=0,n=0;t.length>n;n++)t[n].length>i&&(i=t[n].length,s=n);return s},o.setMonth(e(this._get(t,a.match(/MM/)?"monthNames":"monthNamesShort"))),o.setDate(e(this._get(t,a.match(/DD/)?"dayNames":"dayNamesShort"))+20-o.getDay())),t.input.attr("size",this._formatDate(t,o).length)}},_inlineDatepicker:function(e,i){var s=t(e);s.hasClass(this.markerClassName)||(s.addClass(this.markerClassName).append(i.dpDiv),t.data(e,"datepicker",i),this._setDate(i,this._getDefaultDate(i),!0),this._updateDatepicker(i),this._updateAlternate(i),i.settings.disabled&&this._disableDatepicker(e),i.dpDiv.css("display","block"))},_dialogDatepicker:function(e,i,s,n,o){var r,h,l,c,u,d=this._dialogInst;return d||(this.uuid+=1,r="dp"+this.uuid,this._dialogInput=t("<input type='text' id='"+r+"' style='position: absolute; top: -100px; width: 0px;'/>"),this._dialogInput.on("keydown",this._doKeyDown),t("body").append(this._dialogInput),d=this._dialogInst=this._newInst(this._dialogInput,!1),d.settings={},t.data(this._dialogInput[0],"datepicker",d)),a(d.settings,n||{}),i=i&&i.constructor===Date?this._formatDate(d,i):i,this._dialogInput.val(i),this._pos=o?o.length?o:[o.pageX,o.pageY]:null,this._pos||(h=document.documentElement.clientWidth,l=document.documentElement.clientHeight,c=document.documentElement.scrollLeft||document.body.scrollLeft,u=document.documentElement.scrollTop||document.body.scrollTop,this._pos=[h/2-100+c,l/2-150+u]),this._dialogInput.css("left",this._pos[0]+20+"px").css("top",this._pos[1]+"px"),d.settings.onSelect=s,this._inDialog=!0,this.dpDiv.addClass(this._dialogClass),this._showDatepicker(this._dialogInput[0]),t.blockUI&&t.blockUI(this.dpDiv),t.data(this._dialogInput[0],"datepicker",d),this},_destroyDatepicker:function(e){var i,s=t(e),n=t.data(e,"datepicker");s.hasClass(this.markerClassName)&&(i=e.nodeName.toLowerCase(),t.removeData(e,"datepicker"),"input"===i?(n.append.remove(),n.trigger.remove(),s.removeClass(this.markerClassName).off("focus",this._showDatepicker).off("keydown",this._doKeyDown).off("keypress",this._doKeyPress).off("keyup",this._doKeyUp)):("div"===i||"span"===i)&&s.removeClass(this.markerClassName).empty(),m===n&&(m=null))},_enableDatepicker:function(e){var i,s,n=t(e),o=t.data(e,"datepicker");n.hasClass(this.markerClassName)&&(i=e.nodeName.toLowerCase(),"input"===i?(e.disabled=!1,o.trigger.filter("button").each(function(){this.disabled=!1}).end().filter("img").css({opacity:"1.0",cursor:""})):("div"===i||"span"===i)&&(s=n.children("."+this._inlineClass),s.children().removeClass("ui-state-disabled"),s.find("select.ui-datepicker-month, select.ui-datepicker-year").prop("disabled",!1)),this._disabledInputs=t.map(this._disabledInputs,function(t){return t===e?null:t}))},_disableDatepicker:function(e){var i,s,n=t(e),o=t.data(e,"datepicker");n.hasClass(this.markerClassName)&&(i=e.nodeName.toLowerCase(),"input"===i?(e.disabled=!0,o.trigger.filter("button").each(function(){this.disabled=!0}).end().filter("img").css({opacity:"0.5",cursor:"default"})):("div"===i||"span"===i)&&(s=n.children("."+this._inlineClass),s.children().addClass("ui-state-disabled"),s.find("select.ui-datepicker-month, select.ui-datepicker-year").prop("disabled",!0)),this._disabledInputs=t.map(this._disabledInputs,function(t){return t===e?null:t}),this._disabledInputs[this._disabledInputs.length]=e)},_isDisabledDatepicker:function(t){if(!t)return!1;for(var e=0;this._disabledInputs.length>e;e++)if(this._disabledInputs[e]===t)return!0;return!1},_getInst:function(e){try{return t.data(e,"datepicker")}catch(i){throw"Missing instance data for this datepicker"}},_optionDatepicker:function(e,i,s){var n,o,r,h,l=this._getInst(e);return 2===arguments.length&&"string"==typeof i?"defaults"===i?t.extend({},t.datepicker._defaults):l?"all"===i?t.extend({},l.settings):this._get(l,i):null:(n=i||{},"string"==typeof i&&(n={},n[i]=s),l&&(this._curInst===l&&this._hideDatepicker(),o=this._getDateDatepicker(e,!0),r=this._getMinMaxDate(l,"min"),h=this._getMinMaxDate(l,"max"),a(l.settings,n),null!==r&&void 0!==n.dateFormat&&void 0===n.minDate&&(l.settings.minDate=this._formatDate(l,r)),null!==h&&void 0!==n.dateFormat&&void 0===n.maxDate&&(l.settings.maxDate=this._formatDate(l,h)),"disabled"in n&&(n.disabled?this._disableDatepicker(e):this._enableDatepicker(e)),this._attachments(t(e),l),this._autoSize(l),this._setDate(l,o),this._updateAlternate(l),this._updateDatepicker(l)),void 0)},_changeDatepicker:function(t,e,i){this._optionDatepicker(t,e,i)},_refreshDatepicker:function(t){var e=this._getInst(t);e&&this._updateDatepicker(e)},_setDateDatepicker:function(t,e){var i=this._getInst(t);i&&(this._setDate(i,e),this._updateDatepicker(i),this._updateAlternate(i))},_getDateDatepicker:function(t,e){var i=this._getInst(t);return i&&!i.inline&&this._setDateFromField(i,e),i?this._getDate(i):null},_doKeyDown:function(e){var i,s,n,o=t.datepicker._getInst(e.target),a=!0,r=o.dpDiv.is(".ui-datepicker-rtl");if(o._keyEvent=!0,t.datepicker._datepickerShowing)switch(e.keyCode){case 9:t.datepicker._hideDatepicker(),a=!1;break;case 13:return n=t("td."+t.datepicker._dayOverClass+":not(."+t.datepicker._currentClass+")",o.dpDiv),n[0]&&t.datepicker._selectDay(e.target,o.selectedMonth,o.selectedYear,n[0]),i=t.datepicker._get(o,"onSelect"),i?(s=t.datepicker._formatDate(o),i.apply(o.input?o.input[0]:null,[s,o])):t.datepicker._hideDatepicker(),!1;case 27:t.datepicker._hideDatepicker();break;case 33:t.datepicker._adjustDate(e.target,e.ctrlKey?-t.datepicker._get(o,"stepBigMonths"):-t.datepicker._get(o,"stepMonths"),"M");break;case 34:t.datepicker._adjustDate(e.target,e.ctrlKey?+t.datepicker._get(o,"stepBigMonths"):+t.datepicker._get(o,"stepMonths"),"M");break;case 35:(e.ctrlKey||e.metaKey)&&t.datepicker._clearDate(e.target),a=e.ctrlKey||e.metaKey;break;case 36:(e.ctrlKey||e.metaKey)&&t.datepicker._gotoToday(e.target),a=e.ctrlKey||e.metaKey;break;case 37:(e.ctrlKey||e.metaKey)&&t.datepicker._adjustDate(e.target,r?1:-1,"D"),a=e.ctrlKey||e.metaKey,e.originalEvent.altKey&&t.datepicker._adjustDate(e.target,e.ctrlKey?-t.datepicker._get(o,"stepBigMonths"):-t.datepicker._get(o,"stepMonths"),"M");break;case 38:(e.ctrlKey||e.metaKey)&&t.datepicker._adjustDate(e.target,-7,"D"),a=e.ctrlKey||e.metaKey;break;case 39:(e.ctrlKey||e.metaKey)&&t.datepicker._adjustDate(e.target,r?-1:1,"D"),a=e.ctrlKey||e.metaKey,e.originalEvent.altKey&&t.datepicker._adjustDate(e.target,e.ctrlKey?+t.datepicker._get(o,"stepBigMonths"):+t.datepicker._get(o,"stepMonths"),"M");break;case 40:(e.ctrlKey||e.metaKey)&&t.datepicker._adjustDate(e.target,7,"D"),a=e.ctrlKey||e.metaKey;break;default:a=!1}else 36===e.keyCode&&e.ctrlKey?t.datepicker._showDatepicker(this):a=!1;a&&(e.preventDefault(),e.stopPropagation())},_doKeyPress:function(e){var i,s,n=t.datepicker._getInst(e.target);return t.datepicker._get(n,"constrainInput")?(i=t.datepicker._possibleChars(t.datepicker._get(n,"dateFormat")),s=String.fromCharCode(null==e.charCode?e.keyCode:e.charCode),e.ctrlKey||e.metaKey||" ">s||!i||i.indexOf(s)>-1):void 0},_doKeyUp:function(e){var i,s=t.datepicker._getInst(e.target);if(s.input.val()!==s.lastVal)try{i=t.datepicker.parseDate(t.datepicker._get(s,"dateFormat"),s.input?s.input.val():null,t.datepicker._getFormatConfig(s)),i&&(t.datepicker._setDateFromField(s),t.datepicker._updateAlternate(s),t.datepicker._updateDatepicker(s))}catch(n){}return!0},_showDatepicker:function(e){if(e=e.target||e,"input"!==e.nodeName.toLowerCase()&&(e=t("input",e.parentNode)[0]),!t.datepicker._isDisabledDatepicker(e)&&t.datepicker._lastInput!==e){var s,n,o,r,h,l,c;s=t.datepicker._getInst(e),t.datepicker._curInst&&t.datepicker._curInst!==s&&(t.datepicker._curInst.dpDiv.stop(!0,!0),s&&t.datepicker._datepickerShowing&&t.datepicker._hideDatepicker(t.datepicker._curInst.input[0])),n=t.datepicker._get(s,"beforeShow"),o=n?n.apply(e,[e,s]):{},o!==!1&&(a(s.settings,o),s.lastVal=null,t.datepicker._lastInput=e,t.datepicker._setDateFromField(s),t.datepicker._inDialog&&(e.value=""),t.datepicker._pos||(t.datepicker._pos=t.datepicker._findPos(e),t.datepicker._pos[1]+=e.offsetHeight),r=!1,t(e).parents().each(function(){return r|="fixed"===t(this).css("position"),!r}),h={left:t.datepicker._pos[0],top:t.datepicker._pos[1]},t.datepicker._pos=null,s.dpDiv.empty(),s.dpDiv.css({position:"absolute",display:"block",top:"-1000px"}),t.datepicker._updateDatepicker(s),h=t.datepicker._checkOffset(s,h,r),s.dpDiv.css({position:t.datepicker._inDialog&&t.blockUI?"static":r?"fixed":"absolute",display:"none",left:h.left+"px",top:h.top+"px"}),s.inline||(l=t.datepicker._get(s,"showAnim"),c=t.datepicker._get(s,"duration"),s.dpDiv.css("z-index",i(t(e))+1),t.datepicker._datepickerShowing=!0,t.effects&&t.effects.effect[l]?s.dpDiv.show(l,t.datepicker._get(s,"showOptions"),c):s.dpDiv[l||"show"](l?c:null),t.datepicker._shouldFocusInput(s)&&s.input.trigger("focus"),t.datepicker._curInst=s))
}},_updateDatepicker:function(e){this.maxRows=4,m=e,e.dpDiv.empty().append(this._generateHTML(e)),this._attachHandlers(e);var i,s=this._getNumberOfMonths(e),n=s[1],a=17,r=e.dpDiv.find("."+this._dayOverClass+" a");r.length>0&&o.apply(r.get(0)),e.dpDiv.removeClass("ui-datepicker-multi-2 ui-datepicker-multi-3 ui-datepicker-multi-4").width(""),n>1&&e.dpDiv.addClass("ui-datepicker-multi-"+n).css("width",a*n+"em"),e.dpDiv[(1!==s[0]||1!==s[1]?"add":"remove")+"Class"]("ui-datepicker-multi"),e.dpDiv[(this._get(e,"isRTL")?"add":"remove")+"Class"]("ui-datepicker-rtl"),e===t.datepicker._curInst&&t.datepicker._datepickerShowing&&t.datepicker._shouldFocusInput(e)&&e.input.trigger("focus"),e.yearshtml&&(i=e.yearshtml,setTimeout(function(){i===e.yearshtml&&e.yearshtml&&e.dpDiv.find("select.ui-datepicker-year:first").replaceWith(e.yearshtml),i=e.yearshtml=null},0))},_shouldFocusInput:function(t){return t.input&&t.input.is(":visible")&&!t.input.is(":disabled")&&!t.input.is(":focus")},_checkOffset:function(e,i,s){var n=e.dpDiv.outerWidth(),o=e.dpDiv.outerHeight(),a=e.input?e.input.outerWidth():0,r=e.input?e.input.outerHeight():0,h=document.documentElement.clientWidth+(s?0:t(document).scrollLeft()),l=document.documentElement.clientHeight+(s?0:t(document).scrollTop());return i.left-=this._get(e,"isRTL")?n-a:0,i.left-=s&&i.left===e.input.offset().left?t(document).scrollLeft():0,i.top-=s&&i.top===e.input.offset().top+r?t(document).scrollTop():0,i.left-=Math.min(i.left,i.left+n>h&&h>n?Math.abs(i.left+n-h):0),i.top-=Math.min(i.top,i.top+o>l&&l>o?Math.abs(o+r):0),i},_findPos:function(e){for(var i,s=this._getInst(e),n=this._get(s,"isRTL");e&&("hidden"===e.type||1!==e.nodeType||t.expr.filters.hidden(e));)e=e[n?"previousSibling":"nextSibling"];return i=t(e).offset(),[i.left,i.top]},_hideDatepicker:function(e){var i,s,n,o,a=this._curInst;!a||e&&a!==t.data(e,"datepicker")||this._datepickerShowing&&(i=this._get(a,"showAnim"),s=this._get(a,"duration"),n=function(){t.datepicker._tidyDialog(a)},t.effects&&(t.effects.effect[i]||t.effects[i])?a.dpDiv.hide(i,t.datepicker._get(a,"showOptions"),s,n):a.dpDiv["slideDown"===i?"slideUp":"fadeIn"===i?"fadeOut":"hide"](i?s:null,n),i||n(),this._datepickerShowing=!1,o=this._get(a,"onClose"),o&&o.apply(a.input?a.input[0]:null,[a.input?a.input.val():"",a]),this._lastInput=null,this._inDialog&&(this._dialogInput.css({position:"absolute",left:"0",top:"-100px"}),t.blockUI&&(t.unblockUI(),t("body").append(this.dpDiv))),this._inDialog=!1)},_tidyDialog:function(t){t.dpDiv.removeClass(this._dialogClass).off(".ui-datepicker-calendar")},_checkExternalClick:function(e){if(t.datepicker._curInst){var i=t(e.target),s=t.datepicker._getInst(i[0]);(i[0].id!==t.datepicker._mainDivId&&0===i.parents("#"+t.datepicker._mainDivId).length&&!i.hasClass(t.datepicker.markerClassName)&&!i.closest("."+t.datepicker._triggerClass).length&&t.datepicker._datepickerShowing&&(!t.datepicker._inDialog||!t.blockUI)||i.hasClass(t.datepicker.markerClassName)&&t.datepicker._curInst!==s)&&t.datepicker._hideDatepicker()}},_adjustDate:function(e,i,s){var n=t(e),o=this._getInst(n[0]);this._isDisabledDatepicker(n[0])||(this._adjustInstDate(o,i+("M"===s?this._get(o,"showCurrentAtPos"):0),s),this._updateDatepicker(o))},_gotoToday:function(e){var i,s=t(e),n=this._getInst(s[0]);this._get(n,"gotoCurrent")&&n.currentDay?(n.selectedDay=n.currentDay,n.drawMonth=n.selectedMonth=n.currentMonth,n.drawYear=n.selectedYear=n.currentYear):(i=new Date,n.selectedDay=i.getDate(),n.drawMonth=n.selectedMonth=i.getMonth(),n.drawYear=n.selectedYear=i.getFullYear()),this._notifyChange(n),this._adjustDate(s)},_selectMonthYear:function(e,i,s){var n=t(e),o=this._getInst(n[0]);o["selected"+("M"===s?"Month":"Year")]=o["draw"+("M"===s?"Month":"Year")]=parseInt(i.options[i.selectedIndex].value,10),this._notifyChange(o),this._adjustDate(n)},_selectDay:function(e,i,s,n){var o,a=t(e);t(n).hasClass(this._unselectableClass)||this._isDisabledDatepicker(a[0])||(o=this._getInst(a[0]),o.selectedDay=o.currentDay=t("a",n).html(),o.selectedMonth=o.currentMonth=i,o.selectedYear=o.currentYear=s,this._selectDate(e,this._formatDate(o,o.currentDay,o.currentMonth,o.currentYear)))},_clearDate:function(e){var i=t(e);this._selectDate(i,"")},_selectDate:function(e,i){var s,n=t(e),o=this._getInst(n[0]);i=null!=i?i:this._formatDate(o),o.input&&o.input.val(i),this._updateAlternate(o),s=this._get(o,"onSelect"),s?s.apply(o.input?o.input[0]:null,[i,o]):o.input&&o.input.trigger("change"),o.inline?this._updateDatepicker(o):(this._hideDatepicker(),this._lastInput=o.input[0],"object"!=typeof o.input[0]&&o.input.trigger("focus"),this._lastInput=null)},_updateAlternate:function(e){var i,s,n,o=this._get(e,"altField");o&&(i=this._get(e,"altFormat")||this._get(e,"dateFormat"),s=this._getDate(e),n=this.formatDate(i,s,this._getFormatConfig(e)),t(o).val(n))},noWeekends:function(t){var e=t.getDay();return[e>0&&6>e,""]},iso8601Week:function(t){var e,i=new Date(t.getTime());return i.setDate(i.getDate()+4-(i.getDay()||7)),e=i.getTime(),i.setMonth(0),i.setDate(1),Math.floor(Math.round((e-i)/864e5)/7)+1},parseDate:function(e,i,s){if(null==e||null==i)throw"Invalid arguments";if(i="object"==typeof i?""+i:i+"",""===i)return null;var n,o,a,r,h=0,l=(s?s.shortYearCutoff:null)||this._defaults.shortYearCutoff,c="string"!=typeof l?l:(new Date).getFullYear()%100+parseInt(l,10),u=(s?s.dayNamesShort:null)||this._defaults.dayNamesShort,d=(s?s.dayNames:null)||this._defaults.dayNames,p=(s?s.monthNamesShort:null)||this._defaults.monthNamesShort,f=(s?s.monthNames:null)||this._defaults.monthNames,g=-1,m=-1,_=-1,v=-1,b=!1,y=function(t){var i=e.length>n+1&&e.charAt(n+1)===t;return i&&n++,i},w=function(t){var e=y(t),s="@"===t?14:"!"===t?20:"y"===t&&e?4:"o"===t?3:2,n="y"===t?s:1,o=RegExp("^\\d{"+n+","+s+"}"),a=i.substring(h).match(o);if(!a)throw"Missing number at position "+h;return h+=a[0].length,parseInt(a[0],10)},k=function(e,s,n){var o=-1,a=t.map(y(e)?n:s,function(t,e){return[[e,t]]}).sort(function(t,e){return-(t[1].length-e[1].length)});if(t.each(a,function(t,e){var s=e[1];return i.substr(h,s.length).toLowerCase()===s.toLowerCase()?(o=e[0],h+=s.length,!1):void 0}),-1!==o)return o+1;throw"Unknown name at position "+h},x=function(){if(i.charAt(h)!==e.charAt(n))throw"Unexpected literal at position "+h;h++};for(n=0;e.length>n;n++)if(b)"'"!==e.charAt(n)||y("'")?x():b=!1;else switch(e.charAt(n)){case"d":_=w("d");break;case"D":k("D",u,d);break;case"o":v=w("o");break;case"m":m=w("m");break;case"M":m=k("M",p,f);break;case"y":g=w("y");break;case"@":r=new Date(w("@")),g=r.getFullYear(),m=r.getMonth()+1,_=r.getDate();break;case"!":r=new Date((w("!")-this._ticksTo1970)/1e4),g=r.getFullYear(),m=r.getMonth()+1,_=r.getDate();break;case"'":y("'")?x():b=!0;break;default:x()}if(i.length>h&&(a=i.substr(h),!/^\s+/.test(a)))throw"Extra/unparsed characters found in date: "+a;if(-1===g?g=(new Date).getFullYear():100>g&&(g+=(new Date).getFullYear()-(new Date).getFullYear()%100+(c>=g?0:-100)),v>-1)for(m=1,_=v;;){if(o=this._getDaysInMonth(g,m-1),o>=_)break;m++,_-=o}if(r=this._daylightSavingAdjust(new Date(g,m-1,_)),r.getFullYear()!==g||r.getMonth()+1!==m||r.getDate()!==_)throw"Invalid date";return r},ATOM:"yy-mm-dd",COOKIE:"D, dd M yy",ISO_8601:"yy-mm-dd",RFC_822:"D, d M y",RFC_850:"DD, dd-M-y",RFC_1036:"D, d M y",RFC_1123:"D, d M yy",RFC_2822:"D, d M yy",RSS:"D, d M y",TICKS:"!",TIMESTAMP:"@",W3C:"yy-mm-dd",_ticksTo1970:1e7*60*60*24*(718685+Math.floor(492.5)-Math.floor(19.7)+Math.floor(4.925)),formatDate:function(t,e,i){if(!e)return"";var s,n=(i?i.dayNamesShort:null)||this._defaults.dayNamesShort,o=(i?i.dayNames:null)||this._defaults.dayNames,a=(i?i.monthNamesShort:null)||this._defaults.monthNamesShort,r=(i?i.monthNames:null)||this._defaults.monthNames,h=function(e){var i=t.length>s+1&&t.charAt(s+1)===e;return i&&s++,i},l=function(t,e,i){var s=""+e;if(h(t))for(;i>s.length;)s="0"+s;return s},c=function(t,e,i,s){return h(t)?s[e]:i[e]},u="",d=!1;if(e)for(s=0;t.length>s;s++)if(d)"'"!==t.charAt(s)||h("'")?u+=t.charAt(s):d=!1;else switch(t.charAt(s)){case"d":u+=l("d",e.getDate(),2);break;case"D":u+=c("D",e.getDay(),n,o);break;case"o":u+=l("o",Math.round((new Date(e.getFullYear(),e.getMonth(),e.getDate()).getTime()-new Date(e.getFullYear(),0,0).getTime())/864e5),3);break;case"m":u+=l("m",e.getMonth()+1,2);break;case"M":u+=c("M",e.getMonth(),a,r);break;case"y":u+=h("y")?e.getFullYear():(10>e.getFullYear()%100?"0":"")+e.getFullYear()%100;break;case"@":u+=e.getTime();break;case"!":u+=1e4*e.getTime()+this._ticksTo1970;break;case"'":h("'")?u+="'":d=!0;break;default:u+=t.charAt(s)}return u},_possibleChars:function(t){var e,i="",s=!1,n=function(i){var s=t.length>e+1&&t.charAt(e+1)===i;return s&&e++,s};for(e=0;t.length>e;e++)if(s)"'"!==t.charAt(e)||n("'")?i+=t.charAt(e):s=!1;else switch(t.charAt(e)){case"d":case"m":case"y":case"@":i+="0123456789";break;case"D":case"M":return null;case"'":n("'")?i+="'":s=!0;break;default:i+=t.charAt(e)}return i},_get:function(t,e){return void 0!==t.settings[e]?t.settings[e]:this._defaults[e]},_setDateFromField:function(t,e){if(t.input.val()!==t.lastVal){var i=this._get(t,"dateFormat"),s=t.lastVal=t.input?t.input.val():null,n=this._getDefaultDate(t),o=n,a=this._getFormatConfig(t);try{o=this.parseDate(i,s,a)||n}catch(r){s=e?"":s}t.selectedDay=o.getDate(),t.drawMonth=t.selectedMonth=o.getMonth(),t.drawYear=t.selectedYear=o.getFullYear(),t.currentDay=s?o.getDate():0,t.currentMonth=s?o.getMonth():0,t.currentYear=s?o.getFullYear():0,this._adjustInstDate(t)}},_getDefaultDate:function(t){return this._restrictMinMax(t,this._determineDate(t,this._get(t,"defaultDate"),new Date))},_determineDate:function(e,i,s){var n=function(t){var e=new Date;return e.setDate(e.getDate()+t),e},o=function(i){try{return t.datepicker.parseDate(t.datepicker._get(e,"dateFormat"),i,t.datepicker._getFormatConfig(e))}catch(s){}for(var n=(i.toLowerCase().match(/^c/)?t.datepicker._getDate(e):null)||new Date,o=n.getFullYear(),a=n.getMonth(),r=n.getDate(),h=/([+\-]?[0-9]+)\s*(d|D|w|W|m|M|y|Y)?/g,l=h.exec(i);l;){switch(l[2]||"d"){case"d":case"D":r+=parseInt(l[1],10);break;case"w":case"W":r+=7*parseInt(l[1],10);break;case"m":case"M":a+=parseInt(l[1],10),r=Math.min(r,t.datepicker._getDaysInMonth(o,a));break;case"y":case"Y":o+=parseInt(l[1],10),r=Math.min(r,t.datepicker._getDaysInMonth(o,a))}l=h.exec(i)}return new Date(o,a,r)},a=null==i||""===i?s:"string"==typeof i?o(i):"number"==typeof i?isNaN(i)?s:n(i):new Date(i.getTime());return a=a&&"Invalid Date"==""+a?s:a,a&&(a.setHours(0),a.setMinutes(0),a.setSeconds(0),a.setMilliseconds(0)),this._daylightSavingAdjust(a)},_daylightSavingAdjust:function(t){return t?(t.setHours(t.getHours()>12?t.getHours()+2:0),t):null},_setDate:function(t,e,i){var s=!e,n=t.selectedMonth,o=t.selectedYear,a=this._restrictMinMax(t,this._determineDate(t,e,new Date));t.selectedDay=t.currentDay=a.getDate(),t.drawMonth=t.selectedMonth=t.currentMonth=a.getMonth(),t.drawYear=t.selectedYear=t.currentYear=a.getFullYear(),n===t.selectedMonth&&o===t.selectedYear||i||this._notifyChange(t),this._adjustInstDate(t),t.input&&t.input.val(s?"":this._formatDate(t))},_getDate:function(t){var e=!t.currentYear||t.input&&""===t.input.val()?null:this._daylightSavingAdjust(new Date(t.currentYear,t.currentMonth,t.currentDay));return e},_attachHandlers:function(e){var i=this._get(e,"stepMonths"),s="#"+e.id.replace(/\\\\/g,"\\");e.dpDiv.find("[data-handler]").map(function(){var e={prev:function(){t.datepicker._adjustDate(s,-i,"M")},next:function(){t.datepicker._adjustDate(s,+i,"M")},hide:function(){t.datepicker._hideDatepicker()},today:function(){t.datepicker._gotoToday(s)},selectDay:function(){return t.datepicker._selectDay(s,+this.getAttribute("data-month"),+this.getAttribute("data-year"),this),!1},selectMonth:function(){return t.datepicker._selectMonthYear(s,this,"M"),!1},selectYear:function(){return t.datepicker._selectMonthYear(s,this,"Y"),!1}};t(this).on(this.getAttribute("data-event"),e[this.getAttribute("data-handler")])})},_generateHTML:function(t){var e,i,s,n,o,a,r,h,l,c,u,d,p,f,g,m,_,v,b,y,w,k,x,C,D,I,T,P,M,S,H,z,O,A,N,W,E,F,L,R=new Date,B=this._daylightSavingAdjust(new Date(R.getFullYear(),R.getMonth(),R.getDate())),Y=this._get(t,"isRTL"),j=this._get(t,"showButtonPanel"),q=this._get(t,"hideIfNoPrevNext"),K=this._get(t,"navigationAsDateFormat"),U=this._getNumberOfMonths(t),V=this._get(t,"showCurrentAtPos"),$=this._get(t,"stepMonths"),X=1!==U[0]||1!==U[1],G=this._daylightSavingAdjust(t.currentDay?new Date(t.currentYear,t.currentMonth,t.currentDay):new Date(9999,9,9)),Q=this._getMinMaxDate(t,"min"),J=this._getMinMaxDate(t,"max"),Z=t.drawMonth-V,te=t.drawYear;if(0>Z&&(Z+=12,te--),J)for(e=this._daylightSavingAdjust(new Date(J.getFullYear(),J.getMonth()-U[0]*U[1]+1,J.getDate())),e=Q&&Q>e?Q:e;this._daylightSavingAdjust(new Date(te,Z,1))>e;)Z--,0>Z&&(Z=11,te--);for(t.drawMonth=Z,t.drawYear=te,i=this._get(t,"prevText"),i=K?this.formatDate(i,this._daylightSavingAdjust(new Date(te,Z-$,1)),this._getFormatConfig(t)):i,s=this._canAdjustMonth(t,-1,te,Z)?"<a class='ui-datepicker-prev ui-corner-all' data-handler='prev' data-event='click' title='"+i+"'><span class='ui-icon ui-icon-circle-triangle-"+(Y?"e":"w")+"'>"+i+"</span></a>":q?"":"<a class='ui-datepicker-prev ui-corner-all ui-state-disabled' title='"+i+"'><span class='ui-icon ui-icon-circle-triangle-"+(Y?"e":"w")+"'>"+i+"</span></a>",n=this._get(t,"nextText"),n=K?this.formatDate(n,this._daylightSavingAdjust(new Date(te,Z+$,1)),this._getFormatConfig(t)):n,o=this._canAdjustMonth(t,1,te,Z)?"<a class='ui-datepicker-next ui-corner-all' data-handler='next' data-event='click' title='"+n+"'><span class='ui-icon ui-icon-circle-triangle-"+(Y?"w":"e")+"'>"+n+"</span></a>":q?"":"<a class='ui-datepicker-next ui-corner-all ui-state-disabled' title='"+n+"'><span class='ui-icon ui-icon-circle-triangle-"+(Y?"w":"e")+"'>"+n+"</span></a>",a=this._get(t,"currentText"),r=this._get(t,"gotoCurrent")&&t.currentDay?G:B,a=K?this.formatDate(a,r,this._getFormatConfig(t)):a,h=t.inline?"":"<button type='button' class='ui-datepicker-close ui-state-default ui-priority-primary ui-corner-all' data-handler='hide' data-event='click'>"+this._get(t,"closeText")+"</button>",l=j?"<div class='ui-datepicker-buttonpane ui-widget-content'>"+(Y?h:"")+(this._isInRange(t,r)?"<button type='button' class='ui-datepicker-current ui-state-default ui-priority-secondary ui-corner-all' data-handler='today' data-event='click'>"+a+"</button>":"")+(Y?"":h)+"</div>":"",c=parseInt(this._get(t,"firstDay"),10),c=isNaN(c)?0:c,u=this._get(t,"showWeek"),d=this._get(t,"dayNames"),p=this._get(t,"dayNamesMin"),f=this._get(t,"monthNames"),g=this._get(t,"monthNamesShort"),m=this._get(t,"beforeShowDay"),_=this._get(t,"showOtherMonths"),v=this._get(t,"selectOtherMonths"),b=this._getDefaultDate(t),y="",k=0;U[0]>k;k++){for(x="",this.maxRows=4,C=0;U[1]>C;C++){if(D=this._daylightSavingAdjust(new Date(te,Z,t.selectedDay)),I=" ui-corner-all",T="",X){if(T+="<div class='ui-datepicker-group",U[1]>1)switch(C){case 0:T+=" ui-datepicker-group-first",I=" ui-corner-"+(Y?"right":"left");break;case U[1]-1:T+=" ui-datepicker-group-last",I=" ui-corner-"+(Y?"left":"right");break;default:T+=" ui-datepicker-group-middle",I=""}T+="'>"}for(T+="<div class='ui-datepicker-header ui-widget-header ui-helper-clearfix"+I+"'>"+(/all|left/.test(I)&&0===k?Y?o:s:"")+(/all|right/.test(I)&&0===k?Y?s:o:"")+this._generateMonthYearHeader(t,Z,te,Q,J,k>0||C>0,f,g)+"</div><table class='ui-datepicker-calendar'><thead>"+"<tr>",P=u?"<th class='ui-datepicker-week-col'>"+this._get(t,"weekHeader")+"</th>":"",w=0;7>w;w++)M=(w+c)%7,P+="<th scope='col'"+((w+c+6)%7>=5?" class='ui-datepicker-week-end'":"")+">"+"<span title='"+d[M]+"'>"+p[M]+"</span></th>";for(T+=P+"</tr></thead><tbody>",S=this._getDaysInMonth(te,Z),te===t.selectedYear&&Z===t.selectedMonth&&(t.selectedDay=Math.min(t.selectedDay,S)),H=(this._getFirstDayOfMonth(te,Z)-c+7)%7,z=Math.ceil((H+S)/7),O=X?this.maxRows>z?this.maxRows:z:z,this.maxRows=O,A=this._daylightSavingAdjust(new Date(te,Z,1-H)),N=0;O>N;N++){for(T+="<tr>",W=u?"<td class='ui-datepicker-week-col'>"+this._get(t,"calculateWeek")(A)+"</td>":"",w=0;7>w;w++)E=m?m.apply(t.input?t.input[0]:null,[A]):[!0,""],F=A.getMonth()!==Z,L=F&&!v||!E[0]||Q&&Q>A||J&&A>J,W+="<td class='"+((w+c+6)%7>=5?" ui-datepicker-week-end":"")+(F?" ui-datepicker-other-month":"")+(A.getTime()===D.getTime()&&Z===t.selectedMonth&&t._keyEvent||b.getTime()===A.getTime()&&b.getTime()===D.getTime()?" "+this._dayOverClass:"")+(L?" "+this._unselectableClass+" ui-state-disabled":"")+(F&&!_?"":" "+E[1]+(A.getTime()===G.getTime()?" "+this._currentClass:"")+(A.getTime()===B.getTime()?" ui-datepicker-today":""))+"'"+(F&&!_||!E[2]?"":" title='"+E[2].replace(/'/g,"&#39;")+"'")+(L?"":" data-handler='selectDay' data-event='click' data-month='"+A.getMonth()+"' data-year='"+A.getFullYear()+"'")+">"+(F&&!_?"&#xa0;":L?"<span class='ui-state-default'>"+A.getDate()+"</span>":"<a class='ui-state-default"+(A.getTime()===B.getTime()?" ui-state-highlight":"")+(A.getTime()===G.getTime()?" ui-state-active":"")+(F?" ui-priority-secondary":"")+"' href='#'>"+A.getDate()+"</a>")+"</td>",A.setDate(A.getDate()+1),A=this._daylightSavingAdjust(A);T+=W+"</tr>"}Z++,Z>11&&(Z=0,te++),T+="</tbody></table>"+(X?"</div>"+(U[0]>0&&C===U[1]-1?"<div class='ui-datepicker-row-break'></div>":""):""),x+=T}y+=x}return y+=l,t._keyEvent=!1,y},_generateMonthYearHeader:function(t,e,i,s,n,o,a,r){var h,l,c,u,d,p,f,g,m=this._get(t,"changeMonth"),_=this._get(t,"changeYear"),v=this._get(t,"showMonthAfterYear"),b="<div class='ui-datepicker-title'>",y="";if(o||!m)y+="<span class='ui-datepicker-month'>"+a[e]+"</span>";else{for(h=s&&s.getFullYear()===i,l=n&&n.getFullYear()===i,y+="<select class='ui-datepicker-month' data-handler='selectMonth' data-event='change'>",c=0;12>c;c++)(!h||c>=s.getMonth())&&(!l||n.getMonth()>=c)&&(y+="<option value='"+c+"'"+(c===e?" selected='selected'":"")+">"+r[c]+"</option>");y+="</select>"}if(v||(b+=y+(!o&&m&&_?"":"&#xa0;")),!t.yearshtml)if(t.yearshtml="",o||!_)b+="<span class='ui-datepicker-year'>"+i+"</span>";else{for(u=this._get(t,"yearRange").split(":"),d=(new Date).getFullYear(),p=function(t){var e=t.match(/c[+\-].*/)?i+parseInt(t.substring(1),10):t.match(/[+\-].*/)?d+parseInt(t,10):parseInt(t,10);return isNaN(e)?d:e},f=p(u[0]),g=Math.max(f,p(u[1]||"")),f=s?Math.max(f,s.getFullYear()):f,g=n?Math.min(g,n.getFullYear()):g,t.yearshtml+="<select class='ui-datepicker-year' data-handler='selectYear' data-event='change'>";g>=f;f++)t.yearshtml+="<option value='"+f+"'"+(f===i?" selected='selected'":"")+">"+f+"</option>";t.yearshtml+="</select>",b+=t.yearshtml,t.yearshtml=null}return b+=this._get(t,"yearSuffix"),v&&(b+=(!o&&m&&_?"":"&#xa0;")+y),b+="</div>"},_adjustInstDate:function(t,e,i){var s=t.selectedYear+("Y"===i?e:0),n=t.selectedMonth+("M"===i?e:0),o=Math.min(t.selectedDay,this._getDaysInMonth(s,n))+("D"===i?e:0),a=this._restrictMinMax(t,this._daylightSavingAdjust(new Date(s,n,o)));t.selectedDay=a.getDate(),t.drawMonth=t.selectedMonth=a.getMonth(),t.drawYear=t.selectedYear=a.getFullYear(),("M"===i||"Y"===i)&&this._notifyChange(t)},_restrictMinMax:function(t,e){var i=this._getMinMaxDate(t,"min"),s=this._getMinMaxDate(t,"max"),n=i&&i>e?i:e;return s&&n>s?s:n},_notifyChange:function(t){var e=this._get(t,"onChangeMonthYear");e&&e.apply(t.input?t.input[0]:null,[t.selectedYear,t.selectedMonth+1,t])},_getNumberOfMonths:function(t){var e=this._get(t,"numberOfMonths");return null==e?[1,1]:"number"==typeof e?[1,e]:e},_getMinMaxDate:function(t,e){return this._determineDate(t,this._get(t,e+"Date"),null)},_getDaysInMonth:function(t,e){return 32-this._daylightSavingAdjust(new Date(t,e,32)).getDate()},_getFirstDayOfMonth:function(t,e){return new Date(t,e,1).getDay()},_canAdjustMonth:function(t,e,i,s){var n=this._getNumberOfMonths(t),o=this._daylightSavingAdjust(new Date(i,s+(0>e?e:n[0]*n[1]),1));return 0>e&&o.setDate(this._getDaysInMonth(o.getFullYear(),o.getMonth())),this._isInRange(t,o)},_isInRange:function(t,e){var i,s,n=this._getMinMaxDate(t,"min"),o=this._getMinMaxDate(t,"max"),a=null,r=null,h=this._get(t,"yearRange");return h&&(i=h.split(":"),s=(new Date).getFullYear(),a=parseInt(i[0],10),r=parseInt(i[1],10),i[0].match(/[+\-].*/)&&(a+=s),i[1].match(/[+\-].*/)&&(r+=s)),(!n||e.getTime()>=n.getTime())&&(!o||e.getTime()<=o.getTime())&&(!a||e.getFullYear()>=a)&&(!r||r>=e.getFullYear())},_getFormatConfig:function(t){var e=this._get(t,"shortYearCutoff");return e="string"!=typeof e?e:(new Date).getFullYear()%100+parseInt(e,10),{shortYearCutoff:e,dayNamesShort:this._get(t,"dayNamesShort"),dayNames:this._get(t,"dayNames"),monthNamesShort:this._get(t,"monthNamesShort"),monthNames:this._get(t,"monthNames")}},_formatDate:function(t,e,i,s){e||(t.currentDay=t.selectedDay,t.currentMonth=t.selectedMonth,t.currentYear=t.selectedYear);var n=e?"object"==typeof e?e:this._daylightSavingAdjust(new Date(s,i,e)):this._daylightSavingAdjust(new Date(t.currentYear,t.currentMonth,t.currentDay));return this.formatDate(this._get(t,"dateFormat"),n,this._getFormatConfig(t))}}),t.fn.datepicker=function(e){if(!this.length)return this;t.datepicker.initialized||(t(document).on("mousedown",t.datepicker._checkExternalClick),t.datepicker.initialized=!0),0===t("#"+t.datepicker._mainDivId).length&&t("body").append(t.datepicker.dpDiv);var i=Array.prototype.slice.call(arguments,1);return"string"!=typeof e||"isDisabled"!==e&&"getDate"!==e&&"widget"!==e?"option"===e&&2===arguments.length&&"string"==typeof arguments[1]?t.datepicker["_"+e+"Datepicker"].apply(t.datepicker,[this[0]].concat(i)):this.each(function(){"string"==typeof e?t.datepicker["_"+e+"Datepicker"].apply(t.datepicker,[this].concat(i)):t.datepicker._attachDatepicker(this,e)}):t.datepicker["_"+e+"Datepicker"].apply(t.datepicker,[this[0]].concat(i))},t.datepicker=new s,t.datepicker.initialized=!1,t.datepicker.uuid=(new Date).getTime(),t.datepicker.version="1.12.1",t.datepicker,t.ui.ie=!!/msie [\w.]+/.exec(navigator.userAgent.toLowerCase());var _=!1;t(document).on("mouseup",function(){_=!1}),t.widget("ui.mouse",{version:"1.12.1",options:{cancel:"input, textarea, button, select, option",distance:1,delay:0},_mouseInit:function(){var e=this;this.element.on("mousedown."+this.widgetName,function(t){return e._mouseDown(t)}).on("click."+this.widgetName,function(i){return!0===t.data(i.target,e.widgetName+".preventClickEvent")?(t.removeData(i.target,e.widgetName+".preventClickEvent"),i.stopImmediatePropagation(),!1):void 0}),this.started=!1},_mouseDestroy:function(){this.element.off("."+this.widgetName),this._mouseMoveDelegate&&this.document.off("mousemove."+this.widgetName,this._mouseMoveDelegate).off("mouseup."+this.widgetName,this._mouseUpDelegate)},_mouseDown:function(e){if(!_){this._mouseMoved=!1,this._mouseStarted&&this._mouseUp(e),this._mouseDownEvent=e;var i=this,s=1===e.which,n="string"==typeof this.options.cancel&&e.target.nodeName?t(e.target).closest(this.options.cancel).length:!1;return s&&!n&&this._mouseCapture(e)?(this.mouseDelayMet=!this.options.delay,this.mouseDelayMet||(this._mouseDelayTimer=setTimeout(function(){i.mouseDelayMet=!0},this.options.delay)),this._mouseDistanceMet(e)&&this._mouseDelayMet(e)&&(this._mouseStarted=this._mouseStart(e)!==!1,!this._mouseStarted)?(e.preventDefault(),!0):(!0===t.data(e.target,this.widgetName+".preventClickEvent")&&t.removeData(e.target,this.widgetName+".preventClickEvent"),this._mouseMoveDelegate=function(t){return i._mouseMove(t)},this._mouseUpDelegate=function(t){return i._mouseUp(t)},this.document.on("mousemove."+this.widgetName,this._mouseMoveDelegate).on("mouseup."+this.widgetName,this._mouseUpDelegate),e.preventDefault(),_=!0,!0)):!0}},_mouseMove:function(e){if(this._mouseMoved){if(t.ui.ie&&(!document.documentMode||9>document.documentMode)&&!e.button)return this._mouseUp(e);if(!e.which)if(e.originalEvent.altKey||e.originalEvent.ctrlKey||e.originalEvent.metaKey||e.originalEvent.shiftKey)this.ignoreMissingWhich=!0;else if(!this.ignoreMissingWhich)return this._mouseUp(e)}return(e.which||e.button)&&(this._mouseMoved=!0),this._mouseStarted?(this._mouseDrag(e),e.preventDefault()):(this._mouseDistanceMet(e)&&this._mouseDelayMet(e)&&(this._mouseStarted=this._mouseStart(this._mouseDownEvent,e)!==!1,this._mouseStarted?this._mouseDrag(e):this._mouseUp(e)),!this._mouseStarted)},_mouseUp:function(e){this.document.off("mousemove."+this.widgetName,this._mouseMoveDelegate).off("mouseup."+this.widgetName,this._mouseUpDelegate),this._mouseStarted&&(this._mouseStarted=!1,e.target===this._mouseDownEvent.target&&t.data(e.target,this.widgetName+".preventClickEvent",!0),this._mouseStop(e)),this._mouseDelayTimer&&(clearTimeout(this._mouseDelayTimer),delete this._mouseDelayTimer),this.ignoreMissingWhich=!1,_=!1,e.preventDefault()},_mouseDistanceMet:function(t){return Math.max(Math.abs(this._mouseDownEvent.pageX-t.pageX),Math.abs(this._mouseDownEvent.pageY-t.pageY))>=this.options.distance},_mouseDelayMet:function(){return this.mouseDelayMet},_mouseStart:function(){},_mouseDrag:function(){},_mouseStop:function(){},_mouseCapture:function(){return!0}}),t.ui.plugin={add:function(e,i,s){var n,o=t.ui[e].prototype;for(n in s)o.plugins[n]=o.plugins[n]||[],o.plugins[n].push([i,s[n]])},call:function(t,e,i,s){var n,o=t.plugins[e];if(o&&(s||t.element[0].parentNode&&11!==t.element[0].parentNode.nodeType))for(n=0;o.length>n;n++)t.options[o[n][0]]&&o[n][1].apply(t.element,i)}},t.ui.safeBlur=function(e){e&&"body"!==e.nodeName.toLowerCase()&&t(e).trigger("blur")},t.widget("ui.draggable",t.ui.mouse,{version:"1.12.1",widgetEventPrefix:"drag",options:{addClasses:!0,appendTo:"parent",axis:!1,connectToSortable:!1,containment:!1,cursor:"auto",cursorAt:!1,grid:!1,handle:!1,helper:"original",iframeFix:!1,opacity:!1,refreshPositions:!1,revert:!1,revertDuration:500,scope:"default",scroll:!0,scrollSensitivity:20,scrollSpeed:20,snap:!1,snapMode:"both",snapTolerance:20,stack:!1,zIndex:!1,drag:null,start:null,stop:null},_create:function(){"original"===this.options.helper&&this._setPositionRelative(),this.options.addClasses&&this._addClass("ui-draggable"),this._setHandleClassName(),this._mouseInit()},_setOption:function(t,e){this._super(t,e),"handle"===t&&(this._removeHandleClassName(),this._setHandleClassName())},_destroy:function(){return(this.helper||this.element).is(".ui-draggable-dragging")?(this.destroyOnClear=!0,void 0):(this._removeHandleClassName(),this._mouseDestroy(),void 0)},_mouseCapture:function(e){var i=this.options;return this.helper||i.disabled||t(e.target).closest(".ui-resizable-handle").length>0?!1:(this.handle=this._getHandle(e),this.handle?(this._blurActiveElement(e),this._blockFrames(i.iframeFix===!0?"iframe":i.iframeFix),!0):!1)},_blockFrames:function(e){this.iframeBlocks=this.document.find(e).map(function(){var e=t(this);return t("<div>").css("position","absolute").appendTo(e.parent()).outerWidth(e.outerWidth()).outerHeight(e.outerHeight()).offset(e.offset())[0]})},_unblockFrames:function(){this.iframeBlocks&&(this.iframeBlocks.remove(),delete this.iframeBlocks)},_blurActiveElement:function(e){var i=t.ui.safeActiveElement(this.document[0]),s=t(e.target);s.closest(i).length||t.ui.safeBlur(i)},_mouseStart:function(e){var i=this.options;return this.helper=this._createHelper(e),this._addClass(this.helper,"ui-draggable-dragging"),this._cacheHelperProportions(),t.ui.ddmanager&&(t.ui.ddmanager.current=this),this._cacheMargins(),this.cssPosition=this.helper.css("position"),this.scrollParent=this.helper.scrollParent(!0),this.offsetParent=this.helper.offsetParent(),this.hasFixedAncestor=this.helper.parents().filter(function(){return"fixed"===t(this).css("position")}).length>0,this.positionAbs=this.element.offset(),this._refreshOffsets(e),this.originalPosition=this.position=this._generatePosition(e,!1),this.originalPageX=e.pageX,this.originalPageY=e.pageY,i.cursorAt&&this._adjustOffsetFromHelper(i.cursorAt),this._setContainment(),this._trigger("start",e)===!1?(this._clear(),!1):(this._cacheHelperProportions(),t.ui.ddmanager&&!i.dropBehaviour&&t.ui.ddmanager.prepareOffsets(this,e),this._mouseDrag(e,!0),t.ui.ddmanager&&t.ui.ddmanager.dragStart(this,e),!0)},_refreshOffsets:function(t){this.offset={top:this.positionAbs.top-this.margins.top,left:this.positionAbs.left-this.margins.left,scroll:!1,parent:this._getParentOffset(),relative:this._getRelativeOffset()},this.offset.click={left:t.pageX-this.offset.left,top:t.pageY-this.offset.top}},_mouseDrag:function(e,i){if(this.hasFixedAncestor&&(this.offset.parent=this._getParentOffset()),this.position=this._generatePosition(e,!0),this.positionAbs=this._convertPositionTo("absolute"),!i){var s=this._uiHash();if(this._trigger("drag",e,s)===!1)return this._mouseUp(new t.Event("mouseup",e)),!1;this.position=s.position}return this.helper[0].style.left=this.position.left+"px",this.helper[0].style.top=this.position.top+"px",t.ui.ddmanager&&t.ui.ddmanager.drag(this,e),!1},_mouseStop:function(e){var i=this,s=!1;return t.ui.ddmanager&&!this.options.dropBehaviour&&(s=t.ui.ddmanager.drop(this,e)),this.dropped&&(s=this.dropped,this.dropped=!1),"invalid"===this.options.revert&&!s||"valid"===this.options.revert&&s||this.options.revert===!0||t.isFunction(this.options.revert)&&this.options.revert.call(this.element,s)?t(this.helper).animate(this.originalPosition,parseInt(this.options.revertDuration,10),function(){i._trigger("stop",e)!==!1&&i._clear()}):this._trigger("stop",e)!==!1&&this._clear(),!1},_mouseUp:function(e){return this._unblockFrames(),t.ui.ddmanager&&t.ui.ddmanager.dragStop(this,e),this.handleElement.is(e.target)&&this.element.trigger("focus"),t.ui.mouse.prototype._mouseUp.call(this,e)},cancel:function(){return this.helper.is(".ui-draggable-dragging")?this._mouseUp(new t.Event("mouseup",{target:this.element[0]})):this._clear(),this},_getHandle:function(e){return this.options.handle?!!t(e.target).closest(this.element.find(this.options.handle)).length:!0},_setHandleClassName:function(){this.handleElement=this.options.handle?this.element.find(this.options.handle):this.element,this._addClass(this.handleElement,"ui-draggable-handle")},_removeHandleClassName:function(){this._removeClass(this.handleElement,"ui-draggable-handle")},_createHelper:function(e){var i=this.options,s=t.isFunction(i.helper),n=s?t(i.helper.apply(this.element[0],[e])):"clone"===i.helper?this.element.clone().removeAttr("id"):this.element;return n.parents("body").length||n.appendTo("parent"===i.appendTo?this.element[0].parentNode:i.appendTo),s&&n[0]===this.element[0]&&this._setPositionRelative(),n[0]===this.element[0]||/(fixed|absolute)/.test(n.css("position"))||n.css("position","absolute"),n},_setPositionRelative:function(){/^(?:r|a|f)/.test(this.element.css("position"))||(this.element[0].style.position="relative")},_adjustOffsetFromHelper:function(e){"string"==typeof e&&(e=e.split(" ")),t.isArray(e)&&(e={left:+e[0],top:+e[1]||0}),"left"in e&&(this.offset.click.left=e.left+this.margins.left),"right"in e&&(this.offset.click.left=this.helperProportions.width-e.right+this.margins.left),"top"in e&&(this.offset.click.top=e.top+this.margins.top),"bottom"in e&&(this.offset.click.top=this.helperProportions.height-e.bottom+this.margins.top)},_isRootNode:function(t){return/(html|body)/i.test(t.tagName)||t===this.document[0]},_getParentOffset:function(){var e=this.offsetParent.offset(),i=this.document[0];return"absolute"===this.cssPosition&&this.scrollParent[0]!==i&&t.contains(this.scrollParent[0],this.offsetParent[0])&&(e.left+=this.scrollParent.scrollLeft(),e.top+=this.scrollParent.scrollTop()),this._isRootNode(this.offsetParent[0])&&(e={top:0,left:0}),{top:e.top+(parseInt(this.offsetParent.css("borderTopWidth"),10)||0),left:e.left+(parseInt(this.offsetParent.css("borderLeftWidth"),10)||0)}},_getRelativeOffset:function(){if("relative"!==this.cssPosition)return{top:0,left:0};var t=this.element.position(),e=this._isRootNode(this.scrollParent[0]);return{top:t.top-(parseInt(this.helper.css("top"),10)||0)+(e?0:this.scrollParent.scrollTop()),left:t.left-(parseInt(this.helper.css("left"),10)||0)+(e?0:this.scrollParent.scrollLeft())}
},_cacheMargins:function(){this.margins={left:parseInt(this.element.css("marginLeft"),10)||0,top:parseInt(this.element.css("marginTop"),10)||0,right:parseInt(this.element.css("marginRight"),10)||0,bottom:parseInt(this.element.css("marginBottom"),10)||0}},_cacheHelperProportions:function(){this.helperProportions={width:this.helper.outerWidth(),height:this.helper.outerHeight()}},_setContainment:function(){var e,i,s,n=this.options,o=this.document[0];return this.relativeContainer=null,n.containment?"window"===n.containment?(this.containment=[t(window).scrollLeft()-this.offset.relative.left-this.offset.parent.left,t(window).scrollTop()-this.offset.relative.top-this.offset.parent.top,t(window).scrollLeft()+t(window).width()-this.helperProportions.width-this.margins.left,t(window).scrollTop()+(t(window).height()||o.body.parentNode.scrollHeight)-this.helperProportions.height-this.margins.top],void 0):"document"===n.containment?(this.containment=[0,0,t(o).width()-this.helperProportions.width-this.margins.left,(t(o).height()||o.body.parentNode.scrollHeight)-this.helperProportions.height-this.margins.top],void 0):n.containment.constructor===Array?(this.containment=n.containment,void 0):("parent"===n.containment&&(n.containment=this.helper[0].parentNode),i=t(n.containment),s=i[0],s&&(e=/(scroll|auto)/.test(i.css("overflow")),this.containment=[(parseInt(i.css("borderLeftWidth"),10)||0)+(parseInt(i.css("paddingLeft"),10)||0),(parseInt(i.css("borderTopWidth"),10)||0)+(parseInt(i.css("paddingTop"),10)||0),(e?Math.max(s.scrollWidth,s.offsetWidth):s.offsetWidth)-(parseInt(i.css("borderRightWidth"),10)||0)-(parseInt(i.css("paddingRight"),10)||0)-this.helperProportions.width-this.margins.left-this.margins.right,(e?Math.max(s.scrollHeight,s.offsetHeight):s.offsetHeight)-(parseInt(i.css("borderBottomWidth"),10)||0)-(parseInt(i.css("paddingBottom"),10)||0)-this.helperProportions.height-this.margins.top-this.margins.bottom],this.relativeContainer=i),void 0):(this.containment=null,void 0)},_convertPositionTo:function(t,e){e||(e=this.position);var i="absolute"===t?1:-1,s=this._isRootNode(this.scrollParent[0]);return{top:e.top+this.offset.relative.top*i+this.offset.parent.top*i-("fixed"===this.cssPosition?-this.offset.scroll.top:s?0:this.offset.scroll.top)*i,left:e.left+this.offset.relative.left*i+this.offset.parent.left*i-("fixed"===this.cssPosition?-this.offset.scroll.left:s?0:this.offset.scroll.left)*i}},_generatePosition:function(t,e){var i,s,n,o,a=this.options,r=this._isRootNode(this.scrollParent[0]),h=t.pageX,l=t.pageY;return r&&this.offset.scroll||(this.offset.scroll={top:this.scrollParent.scrollTop(),left:this.scrollParent.scrollLeft()}),e&&(this.containment&&(this.relativeContainer?(s=this.relativeContainer.offset(),i=[this.containment[0]+s.left,this.containment[1]+s.top,this.containment[2]+s.left,this.containment[3]+s.top]):i=this.containment,t.pageX-this.offset.click.left<i[0]&&(h=i[0]+this.offset.click.left),t.pageY-this.offset.click.top<i[1]&&(l=i[1]+this.offset.click.top),t.pageX-this.offset.click.left>i[2]&&(h=i[2]+this.offset.click.left),t.pageY-this.offset.click.top>i[3]&&(l=i[3]+this.offset.click.top)),a.grid&&(n=a.grid[1]?this.originalPageY+Math.round((l-this.originalPageY)/a.grid[1])*a.grid[1]:this.originalPageY,l=i?n-this.offset.click.top>=i[1]||n-this.offset.click.top>i[3]?n:n-this.offset.click.top>=i[1]?n-a.grid[1]:n+a.grid[1]:n,o=a.grid[0]?this.originalPageX+Math.round((h-this.originalPageX)/a.grid[0])*a.grid[0]:this.originalPageX,h=i?o-this.offset.click.left>=i[0]||o-this.offset.click.left>i[2]?o:o-this.offset.click.left>=i[0]?o-a.grid[0]:o+a.grid[0]:o),"y"===a.axis&&(h=this.originalPageX),"x"===a.axis&&(l=this.originalPageY)),{top:l-this.offset.click.top-this.offset.relative.top-this.offset.parent.top+("fixed"===this.cssPosition?-this.offset.scroll.top:r?0:this.offset.scroll.top),left:h-this.offset.click.left-this.offset.relative.left-this.offset.parent.left+("fixed"===this.cssPosition?-this.offset.scroll.left:r?0:this.offset.scroll.left)}},_clear:function(){this._removeClass(this.helper,"ui-draggable-dragging"),this.helper[0]===this.element[0]||this.cancelHelperRemoval||this.helper.remove(),this.helper=null,this.cancelHelperRemoval=!1,this.destroyOnClear&&this.destroy()},_trigger:function(e,i,s){return s=s||this._uiHash(),t.ui.plugin.call(this,e,[i,s,this],!0),/^(drag|start|stop)/.test(e)&&(this.positionAbs=this._convertPositionTo("absolute"),s.offset=this.positionAbs),t.Widget.prototype._trigger.call(this,e,i,s)},plugins:{},_uiHash:function(){return{helper:this.helper,position:this.position,originalPosition:this.originalPosition,offset:this.positionAbs}}}),t.ui.plugin.add("draggable","connectToSortable",{start:function(e,i,s){var n=t.extend({},i,{item:s.element});s.sortables=[],t(s.options.connectToSortable).each(function(){var i=t(this).sortable("instance");i&&!i.options.disabled&&(s.sortables.push(i),i.refreshPositions(),i._trigger("activate",e,n))})},stop:function(e,i,s){var n=t.extend({},i,{item:s.element});s.cancelHelperRemoval=!1,t.each(s.sortables,function(){var t=this;t.isOver?(t.isOver=0,s.cancelHelperRemoval=!0,t.cancelHelperRemoval=!1,t._storedCSS={position:t.placeholder.css("position"),top:t.placeholder.css("top"),left:t.placeholder.css("left")},t._mouseStop(e),t.options.helper=t.options._helper):(t.cancelHelperRemoval=!0,t._trigger("deactivate",e,n))})},drag:function(e,i,s){t.each(s.sortables,function(){var n=!1,o=this;o.positionAbs=s.positionAbs,o.helperProportions=s.helperProportions,o.offset.click=s.offset.click,o._intersectsWith(o.containerCache)&&(n=!0,t.each(s.sortables,function(){return this.positionAbs=s.positionAbs,this.helperProportions=s.helperProportions,this.offset.click=s.offset.click,this!==o&&this._intersectsWith(this.containerCache)&&t.contains(o.element[0],this.element[0])&&(n=!1),n})),n?(o.isOver||(o.isOver=1,s._parent=i.helper.parent(),o.currentItem=i.helper.appendTo(o.element).data("ui-sortable-item",!0),o.options._helper=o.options.helper,o.options.helper=function(){return i.helper[0]},e.target=o.currentItem[0],o._mouseCapture(e,!0),o._mouseStart(e,!0,!0),o.offset.click.top=s.offset.click.top,o.offset.click.left=s.offset.click.left,o.offset.parent.left-=s.offset.parent.left-o.offset.parent.left,o.offset.parent.top-=s.offset.parent.top-o.offset.parent.top,s._trigger("toSortable",e),s.dropped=o.element,t.each(s.sortables,function(){this.refreshPositions()}),s.currentItem=s.element,o.fromOutside=s),o.currentItem&&(o._mouseDrag(e),i.position=o.position)):o.isOver&&(o.isOver=0,o.cancelHelperRemoval=!0,o.options._revert=o.options.revert,o.options.revert=!1,o._trigger("out",e,o._uiHash(o)),o._mouseStop(e,!0),o.options.revert=o.options._revert,o.options.helper=o.options._helper,o.placeholder&&o.placeholder.remove(),i.helper.appendTo(s._parent),s._refreshOffsets(e),i.position=s._generatePosition(e,!0),s._trigger("fromSortable",e),s.dropped=!1,t.each(s.sortables,function(){this.refreshPositions()}))})}}),t.ui.plugin.add("draggable","cursor",{start:function(e,i,s){var n=t("body"),o=s.options;n.css("cursor")&&(o._cursor=n.css("cursor")),n.css("cursor",o.cursor)},stop:function(e,i,s){var n=s.options;n._cursor&&t("body").css("cursor",n._cursor)}}),t.ui.plugin.add("draggable","opacity",{start:function(e,i,s){var n=t(i.helper),o=s.options;n.css("opacity")&&(o._opacity=n.css("opacity")),n.css("opacity",o.opacity)},stop:function(e,i,s){var n=s.options;n._opacity&&t(i.helper).css("opacity",n._opacity)}}),t.ui.plugin.add("draggable","scroll",{start:function(t,e,i){i.scrollParentNotHidden||(i.scrollParentNotHidden=i.helper.scrollParent(!1)),i.scrollParentNotHidden[0]!==i.document[0]&&"HTML"!==i.scrollParentNotHidden[0].tagName&&(i.overflowOffset=i.scrollParentNotHidden.offset())},drag:function(e,i,s){var n=s.options,o=!1,a=s.scrollParentNotHidden[0],r=s.document[0];a!==r&&"HTML"!==a.tagName?(n.axis&&"x"===n.axis||(s.overflowOffset.top+a.offsetHeight-e.pageY<n.scrollSensitivity?a.scrollTop=o=a.scrollTop+n.scrollSpeed:e.pageY-s.overflowOffset.top<n.scrollSensitivity&&(a.scrollTop=o=a.scrollTop-n.scrollSpeed)),n.axis&&"y"===n.axis||(s.overflowOffset.left+a.offsetWidth-e.pageX<n.scrollSensitivity?a.scrollLeft=o=a.scrollLeft+n.scrollSpeed:e.pageX-s.overflowOffset.left<n.scrollSensitivity&&(a.scrollLeft=o=a.scrollLeft-n.scrollSpeed))):(n.axis&&"x"===n.axis||(e.pageY-t(r).scrollTop()<n.scrollSensitivity?o=t(r).scrollTop(t(r).scrollTop()-n.scrollSpeed):t(window).height()-(e.pageY-t(r).scrollTop())<n.scrollSensitivity&&(o=t(r).scrollTop(t(r).scrollTop()+n.scrollSpeed))),n.axis&&"y"===n.axis||(e.pageX-t(r).scrollLeft()<n.scrollSensitivity?o=t(r).scrollLeft(t(r).scrollLeft()-n.scrollSpeed):t(window).width()-(e.pageX-t(r).scrollLeft())<n.scrollSensitivity&&(o=t(r).scrollLeft(t(r).scrollLeft()+n.scrollSpeed)))),o!==!1&&t.ui.ddmanager&&!n.dropBehaviour&&t.ui.ddmanager.prepareOffsets(s,e)}}),t.ui.plugin.add("draggable","snap",{start:function(e,i,s){var n=s.options;s.snapElements=[],t(n.snap.constructor!==String?n.snap.items||":data(ui-draggable)":n.snap).each(function(){var e=t(this),i=e.offset();this!==s.element[0]&&s.snapElements.push({item:this,width:e.outerWidth(),height:e.outerHeight(),top:i.top,left:i.left})})},drag:function(e,i,s){var n,o,a,r,h,l,c,u,d,p,f=s.options,g=f.snapTolerance,m=i.offset.left,_=m+s.helperProportions.width,v=i.offset.top,b=v+s.helperProportions.height;for(d=s.snapElements.length-1;d>=0;d--)h=s.snapElements[d].left-s.margins.left,l=h+s.snapElements[d].width,c=s.snapElements[d].top-s.margins.top,u=c+s.snapElements[d].height,h-g>_||m>l+g||c-g>b||v>u+g||!t.contains(s.snapElements[d].item.ownerDocument,s.snapElements[d].item)?(s.snapElements[d].snapping&&s.options.snap.release&&s.options.snap.release.call(s.element,e,t.extend(s._uiHash(),{snapItem:s.snapElements[d].item})),s.snapElements[d].snapping=!1):("inner"!==f.snapMode&&(n=g>=Math.abs(c-b),o=g>=Math.abs(u-v),a=g>=Math.abs(h-_),r=g>=Math.abs(l-m),n&&(i.position.top=s._convertPositionTo("relative",{top:c-s.helperProportions.height,left:0}).top),o&&(i.position.top=s._convertPositionTo("relative",{top:u,left:0}).top),a&&(i.position.left=s._convertPositionTo("relative",{top:0,left:h-s.helperProportions.width}).left),r&&(i.position.left=s._convertPositionTo("relative",{top:0,left:l}).left)),p=n||o||a||r,"outer"!==f.snapMode&&(n=g>=Math.abs(c-v),o=g>=Math.abs(u-b),a=g>=Math.abs(h-m),r=g>=Math.abs(l-_),n&&(i.position.top=s._convertPositionTo("relative",{top:c,left:0}).top),o&&(i.position.top=s._convertPositionTo("relative",{top:u-s.helperProportions.height,left:0}).top),a&&(i.position.left=s._convertPositionTo("relative",{top:0,left:h}).left),r&&(i.position.left=s._convertPositionTo("relative",{top:0,left:l-s.helperProportions.width}).left)),!s.snapElements[d].snapping&&(n||o||a||r||p)&&s.options.snap.snap&&s.options.snap.snap.call(s.element,e,t.extend(s._uiHash(),{snapItem:s.snapElements[d].item})),s.snapElements[d].snapping=n||o||a||r||p)}}),t.ui.plugin.add("draggable","stack",{start:function(e,i,s){var n,o=s.options,a=t.makeArray(t(o.stack)).sort(function(e,i){return(parseInt(t(e).css("zIndex"),10)||0)-(parseInt(t(i).css("zIndex"),10)||0)});a.length&&(n=parseInt(t(a[0]).css("zIndex"),10)||0,t(a).each(function(e){t(this).css("zIndex",n+e)}),this.css("zIndex",n+a.length))}}),t.ui.plugin.add("draggable","zIndex",{start:function(e,i,s){var n=t(i.helper),o=s.options;n.css("zIndex")&&(o._zIndex=n.css("zIndex")),n.css("zIndex",o.zIndex)},stop:function(e,i,s){var n=s.options;n._zIndex&&t(i.helper).css("zIndex",n._zIndex)}}),t.ui.draggable,t.widget("ui.resizable",t.ui.mouse,{version:"1.12.1",widgetEventPrefix:"resize",options:{alsoResize:!1,animate:!1,animateDuration:"slow",animateEasing:"swing",aspectRatio:!1,autoHide:!1,classes:{"ui-resizable-se":"ui-icon ui-icon-gripsmall-diagonal-se"},containment:!1,ghost:!1,grid:!1,handles:"e,s,se",helper:!1,maxHeight:null,maxWidth:null,minHeight:10,minWidth:10,zIndex:90,resize:null,start:null,stop:null},_num:function(t){return parseFloat(t)||0},_isNumber:function(t){return!isNaN(parseFloat(t))},_hasScroll:function(e,i){if("hidden"===t(e).css("overflow"))return!1;var s=i&&"left"===i?"scrollLeft":"scrollTop",n=!1;return e[s]>0?!0:(e[s]=1,n=e[s]>0,e[s]=0,n)},_create:function(){var e,i=this.options,s=this;this._addClass("ui-resizable"),t.extend(this,{_aspectRatio:!!i.aspectRatio,aspectRatio:i.aspectRatio,originalElement:this.element,_proportionallyResizeElements:[],_helper:i.helper||i.ghost||i.animate?i.helper||"ui-resizable-helper":null}),this.element[0].nodeName.match(/^(canvas|textarea|input|select|button|img)$/i)&&(this.element.wrap(t("<div class='ui-wrapper' style='overflow: hidden;'></div>").css({position:this.element.css("position"),width:this.element.outerWidth(),height:this.element.outerHeight(),top:this.element.css("top"),left:this.element.css("left")})),this.element=this.element.parent().data("ui-resizable",this.element.resizable("instance")),this.elementIsWrapper=!0,e={marginTop:this.originalElement.css("marginTop"),marginRight:this.originalElement.css("marginRight"),marginBottom:this.originalElement.css("marginBottom"),marginLeft:this.originalElement.css("marginLeft")},this.element.css(e),this.originalElement.css("margin",0),this.originalResizeStyle=this.originalElement.css("resize"),this.originalElement.css("resize","none"),this._proportionallyResizeElements.push(this.originalElement.css({position:"static",zoom:1,display:"block"})),this.originalElement.css(e),this._proportionallyResize()),this._setupHandles(),i.autoHide&&t(this.element).on("mouseenter",function(){i.disabled||(s._removeClass("ui-resizable-autohide"),s._handles.show())}).on("mouseleave",function(){i.disabled||s.resizing||(s._addClass("ui-resizable-autohide"),s._handles.hide())}),this._mouseInit()},_destroy:function(){this._mouseDestroy();var e,i=function(e){t(e).removeData("resizable").removeData("ui-resizable").off(".resizable").find(".ui-resizable-handle").remove()};return this.elementIsWrapper&&(i(this.element),e=this.element,this.originalElement.css({position:e.css("position"),width:e.outerWidth(),height:e.outerHeight(),top:e.css("top"),left:e.css("left")}).insertAfter(e),e.remove()),this.originalElement.css("resize",this.originalResizeStyle),i(this.originalElement),this},_setOption:function(t,e){switch(this._super(t,e),t){case"handles":this._removeHandles(),this._setupHandles();break;default:}},_setupHandles:function(){var e,i,s,n,o,a=this.options,r=this;if(this.handles=a.handles||(t(".ui-resizable-handle",this.element).length?{n:".ui-resizable-n",e:".ui-resizable-e",s:".ui-resizable-s",w:".ui-resizable-w",se:".ui-resizable-se",sw:".ui-resizable-sw",ne:".ui-resizable-ne",nw:".ui-resizable-nw"}:"e,s,se"),this._handles=t(),this.handles.constructor===String)for("all"===this.handles&&(this.handles="n,e,s,w,se,sw,ne,nw"),s=this.handles.split(","),this.handles={},i=0;s.length>i;i++)e=t.trim(s[i]),n="ui-resizable-"+e,o=t("<div>"),this._addClass(o,"ui-resizable-handle "+n),o.css({zIndex:a.zIndex}),this.handles[e]=".ui-resizable-"+e,this.element.append(o);this._renderAxis=function(e){var i,s,n,o;e=e||this.element;for(i in this.handles)this.handles[i].constructor===String?this.handles[i]=this.element.children(this.handles[i]).first().show():(this.handles[i].jquery||this.handles[i].nodeType)&&(this.handles[i]=t(this.handles[i]),this._on(this.handles[i],{mousedown:r._mouseDown})),this.elementIsWrapper&&this.originalElement[0].nodeName.match(/^(textarea|input|select|button)$/i)&&(s=t(this.handles[i],this.element),o=/sw|ne|nw|se|n|s/.test(i)?s.outerHeight():s.outerWidth(),n=["padding",/ne|nw|n/.test(i)?"Top":/se|sw|s/.test(i)?"Bottom":/^e$/.test(i)?"Right":"Left"].join(""),e.css(n,o),this._proportionallyResize()),this._handles=this._handles.add(this.handles[i])},this._renderAxis(this.element),this._handles=this._handles.add(this.element.find(".ui-resizable-handle")),this._handles.disableSelection(),this._handles.on("mouseover",function(){r.resizing||(this.className&&(o=this.className.match(/ui-resizable-(se|sw|ne|nw|n|e|s|w)/i)),r.axis=o&&o[1]?o[1]:"se")}),a.autoHide&&(this._handles.hide(),this._addClass("ui-resizable-autohide"))},_removeHandles:function(){this._handles.remove()},_mouseCapture:function(e){var i,s,n=!1;for(i in this.handles)s=t(this.handles[i])[0],(s===e.target||t.contains(s,e.target))&&(n=!0);return!this.options.disabled&&n},_mouseStart:function(e){var i,s,n,o=this.options,a=this.element;return this.resizing=!0,this._renderProxy(),i=this._num(this.helper.css("left")),s=this._num(this.helper.css("top")),o.containment&&(i+=t(o.containment).scrollLeft()||0,s+=t(o.containment).scrollTop()||0),this.offset=this.helper.offset(),this.position={left:i,top:s},this.size=this._helper?{width:this.helper.width(),height:this.helper.height()}:{width:a.width(),height:a.height()},this.originalSize=this._helper?{width:a.outerWidth(),height:a.outerHeight()}:{width:a.width(),height:a.height()},this.sizeDiff={width:a.outerWidth()-a.width(),height:a.outerHeight()-a.height()},this.originalPosition={left:i,top:s},this.originalMousePosition={left:e.pageX,top:e.pageY},this.aspectRatio="number"==typeof o.aspectRatio?o.aspectRatio:this.originalSize.width/this.originalSize.height||1,n=t(".ui-resizable-"+this.axis).css("cursor"),t("body").css("cursor","auto"===n?this.axis+"-resize":n),this._addClass("ui-resizable-resizing"),this._propagate("start",e),!0},_mouseDrag:function(e){var i,s,n=this.originalMousePosition,o=this.axis,a=e.pageX-n.left||0,r=e.pageY-n.top||0,h=this._change[o];return this._updatePrevProperties(),h?(i=h.apply(this,[e,a,r]),this._updateVirtualBoundaries(e.shiftKey),(this._aspectRatio||e.shiftKey)&&(i=this._updateRatio(i,e)),i=this._respectSize(i,e),this._updateCache(i),this._propagate("resize",e),s=this._applyChanges(),!this._helper&&this._proportionallyResizeElements.length&&this._proportionallyResize(),t.isEmptyObject(s)||(this._updatePrevProperties(),this._trigger("resize",e,this.ui()),this._applyChanges()),!1):!1},_mouseStop:function(e){this.resizing=!1;var i,s,n,o,a,r,h,l=this.options,c=this;return this._helper&&(i=this._proportionallyResizeElements,s=i.length&&/textarea/i.test(i[0].nodeName),n=s&&this._hasScroll(i[0],"left")?0:c.sizeDiff.height,o=s?0:c.sizeDiff.width,a={width:c.helper.width()-o,height:c.helper.height()-n},r=parseFloat(c.element.css("left"))+(c.position.left-c.originalPosition.left)||null,h=parseFloat(c.element.css("top"))+(c.position.top-c.originalPosition.top)||null,l.animate||this.element.css(t.extend(a,{top:h,left:r})),c.helper.height(c.size.height),c.helper.width(c.size.width),this._helper&&!l.animate&&this._proportionallyResize()),t("body").css("cursor","auto"),this._removeClass("ui-resizable-resizing"),this._propagate("stop",e),this._helper&&this.helper.remove(),!1},_updatePrevProperties:function(){this.prevPosition={top:this.position.top,left:this.position.left},this.prevSize={width:this.size.width,height:this.size.height}},_applyChanges:function(){var t={};return this.position.top!==this.prevPosition.top&&(t.top=this.position.top+"px"),this.position.left!==this.prevPosition.left&&(t.left=this.position.left+"px"),this.size.width!==this.prevSize.width&&(t.width=this.size.width+"px"),this.size.height!==this.prevSize.height&&(t.height=this.size.height+"px"),this.helper.css(t),t},_updateVirtualBoundaries:function(t){var e,i,s,n,o,a=this.options;o={minWidth:this._isNumber(a.minWidth)?a.minWidth:0,maxWidth:this._isNumber(a.maxWidth)?a.maxWidth:1/0,minHeight:this._isNumber(a.minHeight)?a.minHeight:0,maxHeight:this._isNumber(a.maxHeight)?a.maxHeight:1/0},(this._aspectRatio||t)&&(e=o.minHeight*this.aspectRatio,s=o.minWidth/this.aspectRatio,i=o.maxHeight*this.aspectRatio,n=o.maxWidth/this.aspectRatio,e>o.minWidth&&(o.minWidth=e),s>o.minHeight&&(o.minHeight=s),o.maxWidth>i&&(o.maxWidth=i),o.maxHeight>n&&(o.maxHeight=n)),this._vBoundaries=o},_updateCache:function(t){this.offset=this.helper.offset(),this._isNumber(t.left)&&(this.position.left=t.left),this._isNumber(t.top)&&(this.position.top=t.top),this._isNumber(t.height)&&(this.size.height=t.height),this._isNumber(t.width)&&(this.size.width=t.width)},_updateRatio:function(t){var e=this.position,i=this.size,s=this.axis;return this._isNumber(t.height)?t.width=t.height*this.aspectRatio:this._isNumber(t.width)&&(t.height=t.width/this.aspectRatio),"sw"===s&&(t.left=e.left+(i.width-t.width),t.top=null),"nw"===s&&(t.top=e.top+(i.height-t.height),t.left=e.left+(i.width-t.width)),t},_respectSize:function(t){var e=this._vBoundaries,i=this.axis,s=this._isNumber(t.width)&&e.maxWidth&&e.maxWidth<t.width,n=this._isNumber(t.height)&&e.maxHeight&&e.maxHeight<t.height,o=this._isNumber(t.width)&&e.minWidth&&e.minWidth>t.width,a=this._isNumber(t.height)&&e.minHeight&&e.minHeight>t.height,r=this.originalPosition.left+this.originalSize.width,h=this.originalPosition.top+this.originalSize.height,l=/sw|nw|w/.test(i),c=/nw|ne|n/.test(i);return o&&(t.width=e.minWidth),a&&(t.height=e.minHeight),s&&(t.width=e.maxWidth),n&&(t.height=e.maxHeight),o&&l&&(t.left=r-e.minWidth),s&&l&&(t.left=r-e.maxWidth),a&&c&&(t.top=h-e.minHeight),n&&c&&(t.top=h-e.maxHeight),t.width||t.height||t.left||!t.top?t.width||t.height||t.top||!t.left||(t.left=null):t.top=null,t},_getPaddingPlusBorderDimensions:function(t){for(var e=0,i=[],s=[t.css("borderTopWidth"),t.css("borderRightWidth"),t.css("borderBottomWidth"),t.css("borderLeftWidth")],n=[t.css("paddingTop"),t.css("paddingRight"),t.css("paddingBottom"),t.css("paddingLeft")];4>e;e++)i[e]=parseFloat(s[e])||0,i[e]+=parseFloat(n[e])||0;return{height:i[0]+i[2],width:i[1]+i[3]}},_proportionallyResize:function(){if(this._proportionallyResizeElements.length)for(var t,e=0,i=this.helper||this.element;this._proportionallyResizeElements.length>e;e++)t=this._proportionallyResizeElements[e],this.outerDimensions||(this.outerDimensions=this._getPaddingPlusBorderDimensions(t)),t.css({height:i.height()-this.outerDimensions.height||0,width:i.width()-this.outerDimensions.width||0})},_renderProxy:function(){var e=this.element,i=this.options;this.elementOffset=e.offset(),this._helper?(this.helper=this.helper||t("<div style='overflow:hidden;'></div>"),this._addClass(this.helper,this._helper),this.helper.css({width:this.element.outerWidth(),height:this.element.outerHeight(),position:"absolute",left:this.elementOffset.left+"px",top:this.elementOffset.top+"px",zIndex:++i.zIndex}),this.helper.appendTo("body").disableSelection()):this.helper=this.element},_change:{e:function(t,e){return{width:this.originalSize.width+e}},w:function(t,e){var i=this.originalSize,s=this.originalPosition;return{left:s.left+e,width:i.width-e}},n:function(t,e,i){var s=this.originalSize,n=this.originalPosition;return{top:n.top+i,height:s.height-i}},s:function(t,e,i){return{height:this.originalSize.height+i}},se:function(e,i,s){return t.extend(this._change.s.apply(this,arguments),this._change.e.apply(this,[e,i,s]))},sw:function(e,i,s){return t.extend(this._change.s.apply(this,arguments),this._change.w.apply(this,[e,i,s]))},ne:function(e,i,s){return t.extend(this._change.n.apply(this,arguments),this._change.e.apply(this,[e,i,s]))},nw:function(e,i,s){return t.extend(this._change.n.apply(this,arguments),this._change.w.apply(this,[e,i,s]))}},_propagate:function(e,i){t.ui.plugin.call(this,e,[i,this.ui()]),"resize"!==e&&this._trigger(e,i,this.ui())},plugins:{},ui:function(){return{originalElement:this.originalElement,element:this.element,helper:this.helper,position:this.position,size:this.size,originalSize:this.originalSize,originalPosition:this.originalPosition}}}),t.ui.plugin.add("resizable","animate",{stop:function(e){var i=t(this).resizable("instance"),s=i.options,n=i._proportionallyResizeElements,o=n.length&&/textarea/i.test(n[0].nodeName),a=o&&i._hasScroll(n[0],"left")?0:i.sizeDiff.height,r=o?0:i.sizeDiff.width,h={width:i.size.width-r,height:i.size.height-a},l=parseFloat(i.element.css("left"))+(i.position.left-i.originalPosition.left)||null,c=parseFloat(i.element.css("top"))+(i.position.top-i.originalPosition.top)||null;i.element.animate(t.extend(h,c&&l?{top:c,left:l}:{}),{duration:s.animateDuration,easing:s.animateEasing,step:function(){var s={width:parseFloat(i.element.css("width")),height:parseFloat(i.element.css("height")),top:parseFloat(i.element.css("top")),left:parseFloat(i.element.css("left"))};n&&n.length&&t(n[0]).css({width:s.width,height:s.height}),i._updateCache(s),i._propagate("resize",e)}})}}),t.ui.plugin.add("resizable","containment",{start:function(){var e,i,s,n,o,a,r,h=t(this).resizable("instance"),l=h.options,c=h.element,u=l.containment,d=u instanceof t?u.get(0):/parent/.test(u)?c.parent().get(0):u;d&&(h.containerElement=t(d),/document/.test(u)||u===document?(h.containerOffset={left:0,top:0},h.containerPosition={left:0,top:0},h.parentData={element:t(document),left:0,top:0,width:t(document).width(),height:t(document).height()||document.body.parentNode.scrollHeight}):(e=t(d),i=[],t(["Top","Right","Left","Bottom"]).each(function(t,s){i[t]=h._num(e.css("padding"+s))}),h.containerOffset=e.offset(),h.containerPosition=e.position(),h.containerSize={height:e.innerHeight()-i[3],width:e.innerWidth()-i[1]},s=h.containerOffset,n=h.containerSize.height,o=h.containerSize.width,a=h._hasScroll(d,"left")?d.scrollWidth:o,r=h._hasScroll(d)?d.scrollHeight:n,h.parentData={element:d,left:s.left,top:s.top,width:a,height:r}))},resize:function(e){var i,s,n,o,a=t(this).resizable("instance"),r=a.options,h=a.containerOffset,l=a.position,c=a._aspectRatio||e.shiftKey,u={top:0,left:0},d=a.containerElement,p=!0;d[0]!==document&&/static/.test(d.css("position"))&&(u=h),l.left<(a._helper?h.left:0)&&(a.size.width=a.size.width+(a._helper?a.position.left-h.left:a.position.left-u.left),c&&(a.size.height=a.size.width/a.aspectRatio,p=!1),a.position.left=r.helper?h.left:0),l.top<(a._helper?h.top:0)&&(a.size.height=a.size.height+(a._helper?a.position.top-h.top:a.position.top),c&&(a.size.width=a.size.height*a.aspectRatio,p=!1),a.position.top=a._helper?h.top:0),n=a.containerElement.get(0)===a.element.parent().get(0),o=/relative|absolute/.test(a.containerElement.css("position")),n&&o?(a.offset.left=a.parentData.left+a.position.left,a.offset.top=a.parentData.top+a.position.top):(a.offset.left=a.element.offset().left,a.offset.top=a.element.offset().top),i=Math.abs(a.sizeDiff.width+(a._helper?a.offset.left-u.left:a.offset.left-h.left)),s=Math.abs(a.sizeDiff.height+(a._helper?a.offset.top-u.top:a.offset.top-h.top)),i+a.size.width>=a.parentData.width&&(a.size.width=a.parentData.width-i,c&&(a.size.height=a.size.width/a.aspectRatio,p=!1)),s+a.size.height>=a.parentData.height&&(a.size.height=a.parentData.height-s,c&&(a.size.width=a.size.height*a.aspectRatio,p=!1)),p||(a.position.left=a.prevPosition.left,a.position.top=a.prevPosition.top,a.size.width=a.prevSize.width,a.size.height=a.prevSize.height)},stop:function(){var e=t(this).resizable("instance"),i=e.options,s=e.containerOffset,n=e.containerPosition,o=e.containerElement,a=t(e.helper),r=a.offset(),h=a.outerWidth()-e.sizeDiff.width,l=a.outerHeight()-e.sizeDiff.height;e._helper&&!i.animate&&/relative/.test(o.css("position"))&&t(this).css({left:r.left-n.left-s.left,width:h,height:l}),e._helper&&!i.animate&&/static/.test(o.css("position"))&&t(this).css({left:r.left-n.left-s.left,width:h,height:l})}}),t.ui.plugin.add("resizable","alsoResize",{start:function(){var e=t(this).resizable("instance"),i=e.options;t(i.alsoResize).each(function(){var e=t(this);e.data("ui-resizable-alsoresize",{width:parseFloat(e.width()),height:parseFloat(e.height()),left:parseFloat(e.css("left")),top:parseFloat(e.css("top"))})})},resize:function(e,i){var s=t(this).resizable("instance"),n=s.options,o=s.originalSize,a=s.originalPosition,r={height:s.size.height-o.height||0,width:s.size.width-o.width||0,top:s.position.top-a.top||0,left:s.position.left-a.left||0};t(n.alsoResize).each(function(){var e=t(this),s=t(this).data("ui-resizable-alsoresize"),n={},o=e.parents(i.originalElement[0]).length?["width","height"]:["width","height","top","left"];t.each(o,function(t,e){var i=(s[e]||0)+(r[e]||0);i&&i>=0&&(n[e]=i||null)}),e.css(n)})},stop:function(){t(this).removeData("ui-resizable-alsoresize")}}),t.ui.plugin.add("resizable","ghost",{start:function(){var e=t(this).resizable("instance"),i=e.size;e.ghost=e.originalElement.clone(),e.ghost.css({opacity:.25,display:"block",position:"relative",height:i.height,width:i.width,margin:0,left:0,top:0}),e._addClass(e.ghost,"ui-resizable-ghost"),t.uiBackCompat!==!1&&"string"==typeof e.options.ghost&&e.ghost.addClass(this.options.ghost),e.ghost.appendTo(e.helper)},resize:function(){var e=t(this).resizable("instance");e.ghost&&e.ghost.css({position:"relative",height:e.size.height,width:e.size.width})},stop:function(){var e=t(this).resizable("instance");e.ghost&&e.helper&&e.helper.get(0).removeChild(e.ghost.get(0))}}),t.ui.plugin.add("resizable","grid",{resize:function(){var e,i=t(this).resizable("instance"),s=i.options,n=i.size,o=i.originalSize,a=i.originalPosition,r=i.axis,h="number"==typeof s.grid?[s.grid,s.grid]:s.grid,l=h[0]||1,c=h[1]||1,u=Math.round((n.width-o.width)/l)*l,d=Math.round((n.height-o.height)/c)*c,p=o.width+u,f=o.height+d,g=s.maxWidth&&p>s.maxWidth,m=s.maxHeight&&f>s.maxHeight,_=s.minWidth&&s.minWidth>p,v=s.minHeight&&s.minHeight>f;s.grid=h,_&&(p+=l),v&&(f+=c),g&&(p-=l),m&&(f-=c),/^(se|s|e)$/.test(r)?(i.size.width=p,i.size.height=f):/^(ne)$/.test(r)?(i.size.width=p,i.size.height=f,i.position.top=a.top-d):/^(sw)$/.test(r)?(i.size.width=p,i.size.height=f,i.position.left=a.left-u):((0>=f-c||0>=p-l)&&(e=i._getPaddingPlusBorderDimensions(this)),f-c>0?(i.size.height=f,i.position.top=a.top-d):(f=c-e.height,i.size.height=f,i.position.top=a.top+o.height-f),p-l>0?(i.size.width=p,i.position.left=a.left-u):(p=l-e.width,i.size.width=p,i.position.left=a.left+o.width-p))}}),t.ui.resizable,t.widget("ui.dialog",{version:"1.12.1",options:{appendTo:"body",autoOpen:!0,buttons:[],classes:{"ui-dialog":"ui-corner-all","ui-dialog-titlebar":"ui-corner-all"},closeOnEscape:!0,closeText:"Close",draggable:!0,hide:null,height:"auto",maxHeight:null,maxWidth:null,minHeight:150,minWidth:150,modal:!1,position:{my:"center",at:"center",of:window,collision:"fit",using:function(e){var i=t(this).css(e).offset().top;0>i&&t(this).css("top",e.top-i)}},resizable:!0,show:null,title:null,width:300,beforeClose:null,close:null,drag:null,dragStart:null,dragStop:null,focus:null,open:null,resize:null,resizeStart:null,resizeStop:null},sizeRelatedOptions:{buttons:!0,height:!0,maxHeight:!0,maxWidth:!0,minHeight:!0,minWidth:!0,width:!0},resizableRelatedOptions:{maxHeight:!0,maxWidth:!0,minHeight:!0,minWidth:!0},_create:function(){this.originalCss={display:this.element[0].style.display,width:this.element[0].style.width,minHeight:this.element[0].style.minHeight,maxHeight:this.element[0].style.maxHeight,height:this.element[0].style.height},this.originalPosition={parent:this.element.parent(),index:this.element.parent().children().index(this.element)},this.originalTitle=this.element.attr("title"),null==this.options.title&&null!=this.originalTitle&&(this.options.title=this.originalTitle),this.options.disabled&&(this.options.disabled=!1),this._createWrapper(),this.element.show().removeAttr("title").appendTo(this.uiDialog),this._addClass("ui-dialog-content","ui-widget-content"),this._createTitlebar(),this._createButtonPane(),this.options.draggable&&t.fn.draggable&&this._makeDraggable(),this.options.resizable&&t.fn.resizable&&this._makeResizable(),this._isOpen=!1,this._trackFocus()},_init:function(){this.options.autoOpen&&this.open()},_appendTo:function(){var e=this.options.appendTo;return e&&(e.jquery||e.nodeType)?t(e):this.document.find(e||"body").eq(0)},_destroy:function(){var t,e=this.originalPosition;this._untrackInstance(),this._destroyOverlay(),this.element.removeUniqueId().css(this.originalCss).detach(),this.uiDialog.remove(),this.originalTitle&&this.element.attr("title",this.originalTitle),t=e.parent.children().eq(e.index),t.length&&t[0]!==this.element[0]?t.before(this.element):e.parent.append(this.element)},widget:function(){return this.uiDialog
},disable:t.noop,enable:t.noop,close:function(e){var i=this;this._isOpen&&this._trigger("beforeClose",e)!==!1&&(this._isOpen=!1,this._focusedElement=null,this._destroyOverlay(),this._untrackInstance(),this.opener.filter(":focusable").trigger("focus").length||t.ui.safeBlur(t.ui.safeActiveElement(this.document[0])),this._hide(this.uiDialog,this.options.hide,function(){i._trigger("close",e)}))},isOpen:function(){return this._isOpen},moveToTop:function(){this._moveToTop()},_moveToTop:function(e,i){var s=!1,n=this.uiDialog.siblings(".ui-front:visible").map(function(){return+t(this).css("z-index")}).get(),o=Math.max.apply(null,n);return o>=+this.uiDialog.css("z-index")&&(this.uiDialog.css("z-index",o+1),s=!0),s&&!i&&this._trigger("focus",e),s},open:function(){var e=this;return this._isOpen?(this._moveToTop()&&this._focusTabbable(),void 0):(this._isOpen=!0,this.opener=t(t.ui.safeActiveElement(this.document[0])),this._size(),this._position(),this._createOverlay(),this._moveToTop(null,!0),this.overlay&&this.overlay.css("z-index",this.uiDialog.css("z-index")-1),this._show(this.uiDialog,this.options.show,function(){e._focusTabbable(),e._trigger("focus")}),this._makeFocusTarget(),this._trigger("open"),void 0)},_focusTabbable:function(){var t=this._focusedElement;t||(t=this.element.find("[autofocus]")),t.length||(t=this.element.find(":tabbable")),t.length||(t=this.uiDialogButtonPane.find(":tabbable")),t.length||(t=this.uiDialogTitlebarClose.filter(":tabbable")),t.length||(t=this.uiDialog),t.eq(0).trigger("focus")},_keepFocus:function(e){function i(){var e=t.ui.safeActiveElement(this.document[0]),i=this.uiDialog[0]===e||t.contains(this.uiDialog[0],e);i||this._focusTabbable()}e.preventDefault(),i.call(this),this._delay(i)},_createWrapper:function(){this.uiDialog=t("<div>").hide().attr({tabIndex:-1,role:"dialog"}).appendTo(this._appendTo()),this._addClass(this.uiDialog,"ui-dialog","ui-widget ui-widget-content ui-front"),this._on(this.uiDialog,{keydown:function(e){if(this.options.closeOnEscape&&!e.isDefaultPrevented()&&e.keyCode&&e.keyCode===t.ui.keyCode.ESCAPE)return e.preventDefault(),this.close(e),void 0;if(e.keyCode===t.ui.keyCode.TAB&&!e.isDefaultPrevented()){var i=this.uiDialog.find(":tabbable"),s=i.filter(":first"),n=i.filter(":last");e.target!==n[0]&&e.target!==this.uiDialog[0]||e.shiftKey?e.target!==s[0]&&e.target!==this.uiDialog[0]||!e.shiftKey||(this._delay(function(){n.trigger("focus")}),e.preventDefault()):(this._delay(function(){s.trigger("focus")}),e.preventDefault())}},mousedown:function(t){this._moveToTop(t)&&this._focusTabbable()}}),this.element.find("[aria-describedby]").length||this.uiDialog.attr({"aria-describedby":this.element.uniqueId().attr("id")})},_createTitlebar:function(){var e;this.uiDialogTitlebar=t("<div>"),this._addClass(this.uiDialogTitlebar,"ui-dialog-titlebar","ui-widget-header ui-helper-clearfix"),this._on(this.uiDialogTitlebar,{mousedown:function(e){t(e.target).closest(".ui-dialog-titlebar-close")||this.uiDialog.trigger("focus")}}),this.uiDialogTitlebarClose=t("<button type='button'></button>").button({label:t("<a>").text(this.options.closeText).html(),icon:"ui-icon-closethick",showLabel:!1}).appendTo(this.uiDialogTitlebar),this._addClass(this.uiDialogTitlebarClose,"ui-dialog-titlebar-close"),this._on(this.uiDialogTitlebarClose,{click:function(t){t.preventDefault(),this.close(t)}}),e=t("<span>").uniqueId().prependTo(this.uiDialogTitlebar),this._addClass(e,"ui-dialog-title"),this._title(e),this.uiDialogTitlebar.prependTo(this.uiDialog),this.uiDialog.attr({"aria-labelledby":e.attr("id")})},_title:function(t){this.options.title?t.text(this.options.title):t.html("&#160;")},_createButtonPane:function(){this.uiDialogButtonPane=t("<div>"),this._addClass(this.uiDialogButtonPane,"ui-dialog-buttonpane","ui-widget-content ui-helper-clearfix"),this.uiButtonSet=t("<div>").appendTo(this.uiDialogButtonPane),this._addClass(this.uiButtonSet,"ui-dialog-buttonset"),this._createButtons()},_createButtons:function(){var e=this,i=this.options.buttons;return this.uiDialogButtonPane.remove(),this.uiButtonSet.empty(),t.isEmptyObject(i)||t.isArray(i)&&!i.length?(this._removeClass(this.uiDialog,"ui-dialog-buttons"),void 0):(t.each(i,function(i,s){var n,o;s=t.isFunction(s)?{click:s,text:i}:s,s=t.extend({type:"button"},s),n=s.click,o={icon:s.icon,iconPosition:s.iconPosition,showLabel:s.showLabel,icons:s.icons,text:s.text},delete s.click,delete s.icon,delete s.iconPosition,delete s.showLabel,delete s.icons,"boolean"==typeof s.text&&delete s.text,t("<button></button>",s).button(o).appendTo(e.uiButtonSet).on("click",function(){n.apply(e.element[0],arguments)})}),this._addClass(this.uiDialog,"ui-dialog-buttons"),this.uiDialogButtonPane.appendTo(this.uiDialog),void 0)},_makeDraggable:function(){function e(t){return{position:t.position,offset:t.offset}}var i=this,s=this.options;this.uiDialog.draggable({cancel:".ui-dialog-content, .ui-dialog-titlebar-close",handle:".ui-dialog-titlebar",containment:"document",start:function(s,n){i._addClass(t(this),"ui-dialog-dragging"),i._blockFrames(),i._trigger("dragStart",s,e(n))},drag:function(t,s){i._trigger("drag",t,e(s))},stop:function(n,o){var a=o.offset.left-i.document.scrollLeft(),r=o.offset.top-i.document.scrollTop();s.position={my:"left top",at:"left"+(a>=0?"+":"")+a+" "+"top"+(r>=0?"+":"")+r,of:i.window},i._removeClass(t(this),"ui-dialog-dragging"),i._unblockFrames(),i._trigger("dragStop",n,e(o))}})},_makeResizable:function(){function e(t){return{originalPosition:t.originalPosition,originalSize:t.originalSize,position:t.position,size:t.size}}var i=this,s=this.options,n=s.resizable,o=this.uiDialog.css("position"),a="string"==typeof n?n:"n,e,s,w,se,sw,ne,nw";this.uiDialog.resizable({cancel:".ui-dialog-content",containment:"document",alsoResize:this.element,maxWidth:s.maxWidth,maxHeight:s.maxHeight,minWidth:s.minWidth,minHeight:this._minHeight(),handles:a,start:function(s,n){i._addClass(t(this),"ui-dialog-resizing"),i._blockFrames(),i._trigger("resizeStart",s,e(n))},resize:function(t,s){i._trigger("resize",t,e(s))},stop:function(n,o){var a=i.uiDialog.offset(),r=a.left-i.document.scrollLeft(),h=a.top-i.document.scrollTop();s.height=i.uiDialog.height(),s.width=i.uiDialog.width(),s.position={my:"left top",at:"left"+(r>=0?"+":"")+r+" "+"top"+(h>=0?"+":"")+h,of:i.window},i._removeClass(t(this),"ui-dialog-resizing"),i._unblockFrames(),i._trigger("resizeStop",n,e(o))}}).css("position",o)},_trackFocus:function(){this._on(this.widget(),{focusin:function(e){this._makeFocusTarget(),this._focusedElement=t(e.target)}})},_makeFocusTarget:function(){this._untrackInstance(),this._trackingInstances().unshift(this)},_untrackInstance:function(){var e=this._trackingInstances(),i=t.inArray(this,e);-1!==i&&e.splice(i,1)},_trackingInstances:function(){var t=this.document.data("ui-dialog-instances");return t||(t=[],this.document.data("ui-dialog-instances",t)),t},_minHeight:function(){var t=this.options;return"auto"===t.height?t.minHeight:Math.min(t.minHeight,t.height)},_position:function(){var t=this.uiDialog.is(":visible");t||this.uiDialog.show(),this.uiDialog.position(this.options.position),t||this.uiDialog.hide()},_setOptions:function(e){var i=this,s=!1,n={};t.each(e,function(t,e){i._setOption(t,e),t in i.sizeRelatedOptions&&(s=!0),t in i.resizableRelatedOptions&&(n[t]=e)}),s&&(this._size(),this._position()),this.uiDialog.is(":data(ui-resizable)")&&this.uiDialog.resizable("option",n)},_setOption:function(e,i){var s,n,o=this.uiDialog;"disabled"!==e&&(this._super(e,i),"appendTo"===e&&this.uiDialog.appendTo(this._appendTo()),"buttons"===e&&this._createButtons(),"closeText"===e&&this.uiDialogTitlebarClose.button({label:t("<a>").text(""+this.options.closeText).html()}),"draggable"===e&&(s=o.is(":data(ui-draggable)"),s&&!i&&o.draggable("destroy"),!s&&i&&this._makeDraggable()),"position"===e&&this._position(),"resizable"===e&&(n=o.is(":data(ui-resizable)"),n&&!i&&o.resizable("destroy"),n&&"string"==typeof i&&o.resizable("option","handles",i),n||i===!1||this._makeResizable()),"title"===e&&this._title(this.uiDialogTitlebar.find(".ui-dialog-title")))},_size:function(){var t,e,i,s=this.options;this.element.show().css({width:"auto",minHeight:0,maxHeight:"none",height:0}),s.minWidth>s.width&&(s.width=s.minWidth),t=this.uiDialog.css({height:"auto",width:s.width}).outerHeight(),e=Math.max(0,s.minHeight-t),i="number"==typeof s.maxHeight?Math.max(0,s.maxHeight-t):"none","auto"===s.height?this.element.css({minHeight:e,maxHeight:i,height:"auto"}):this.element.height(Math.max(0,s.height-t)),this.uiDialog.is(":data(ui-resizable)")&&this.uiDialog.resizable("option","minHeight",this._minHeight())},_blockFrames:function(){this.iframeBlocks=this.document.find("iframe").map(function(){var e=t(this);return t("<div>").css({position:"absolute",width:e.outerWidth(),height:e.outerHeight()}).appendTo(e.parent()).offset(e.offset())[0]})},_unblockFrames:function(){this.iframeBlocks&&(this.iframeBlocks.remove(),delete this.iframeBlocks)},_allowInteraction:function(e){return t(e.target).closest(".ui-dialog").length?!0:!!t(e.target).closest(".ui-datepicker").length},_createOverlay:function(){if(this.options.modal){var e=!0;this._delay(function(){e=!1}),this.document.data("ui-dialog-overlays")||this._on(this.document,{focusin:function(t){e||this._allowInteraction(t)||(t.preventDefault(),this._trackingInstances()[0]._focusTabbable())}}),this.overlay=t("<div>").appendTo(this._appendTo()),this._addClass(this.overlay,null,"ui-widget-overlay ui-front"),this._on(this.overlay,{mousedown:"_keepFocus"}),this.document.data("ui-dialog-overlays",(this.document.data("ui-dialog-overlays")||0)+1)}},_destroyOverlay:function(){if(this.options.modal&&this.overlay){var t=this.document.data("ui-dialog-overlays")-1;t?this.document.data("ui-dialog-overlays",t):(this._off(this.document,"focusin"),this.document.removeData("ui-dialog-overlays")),this.overlay.remove(),this.overlay=null}}}),t.uiBackCompat!==!1&&t.widget("ui.dialog",t.ui.dialog,{options:{dialogClass:""},_createWrapper:function(){this._super(),this.uiDialog.addClass(this.options.dialogClass)},_setOption:function(t,e){"dialogClass"===t&&this.uiDialog.removeClass(this.options.dialogClass).addClass(e),this._superApply(arguments)}}),t.ui.dialog,t.widget("ui.droppable",{version:"1.12.1",widgetEventPrefix:"drop",options:{accept:"*",addClasses:!0,greedy:!1,scope:"default",tolerance:"intersect",activate:null,deactivate:null,drop:null,out:null,over:null},_create:function(){var e,i=this.options,s=i.accept;this.isover=!1,this.isout=!0,this.accept=t.isFunction(s)?s:function(t){return t.is(s)},this.proportions=function(){return arguments.length?(e=arguments[0],void 0):e?e:e={width:this.element[0].offsetWidth,height:this.element[0].offsetHeight}},this._addToManager(i.scope),i.addClasses&&this._addClass("ui-droppable")},_addToManager:function(e){t.ui.ddmanager.droppables[e]=t.ui.ddmanager.droppables[e]||[],t.ui.ddmanager.droppables[e].push(this)},_splice:function(t){for(var e=0;t.length>e;e++)t[e]===this&&t.splice(e,1)},_destroy:function(){var e=t.ui.ddmanager.droppables[this.options.scope];this._splice(e)},_setOption:function(e,i){if("accept"===e)this.accept=t.isFunction(i)?i:function(t){return t.is(i)};else if("scope"===e){var s=t.ui.ddmanager.droppables[this.options.scope];this._splice(s),this._addToManager(i)}this._super(e,i)},_activate:function(e){var i=t.ui.ddmanager.current;this._addActiveClass(),i&&this._trigger("activate",e,this.ui(i))},_deactivate:function(e){var i=t.ui.ddmanager.current;this._removeActiveClass(),i&&this._trigger("deactivate",e,this.ui(i))},_over:function(e){var i=t.ui.ddmanager.current;i&&(i.currentItem||i.element)[0]!==this.element[0]&&this.accept.call(this.element[0],i.currentItem||i.element)&&(this._addHoverClass(),this._trigger("over",e,this.ui(i)))},_out:function(e){var i=t.ui.ddmanager.current;i&&(i.currentItem||i.element)[0]!==this.element[0]&&this.accept.call(this.element[0],i.currentItem||i.element)&&(this._removeHoverClass(),this._trigger("out",e,this.ui(i)))},_drop:function(e,i){var s=i||t.ui.ddmanager.current,n=!1;return s&&(s.currentItem||s.element)[0]!==this.element[0]?(this.element.find(":data(ui-droppable)").not(".ui-draggable-dragging").each(function(){var i=t(this).droppable("instance");return i.options.greedy&&!i.options.disabled&&i.options.scope===s.options.scope&&i.accept.call(i.element[0],s.currentItem||s.element)&&v(s,t.extend(i,{offset:i.element.offset()}),i.options.tolerance,e)?(n=!0,!1):void 0}),n?!1:this.accept.call(this.element[0],s.currentItem||s.element)?(this._removeActiveClass(),this._removeHoverClass(),this._trigger("drop",e,this.ui(s)),this.element):!1):!1},ui:function(t){return{draggable:t.currentItem||t.element,helper:t.helper,position:t.position,offset:t.positionAbs}},_addHoverClass:function(){this._addClass("ui-droppable-hover")},_removeHoverClass:function(){this._removeClass("ui-droppable-hover")},_addActiveClass:function(){this._addClass("ui-droppable-active")},_removeActiveClass:function(){this._removeClass("ui-droppable-active")}});var v=t.ui.intersect=function(){function t(t,e,i){return t>=e&&e+i>t}return function(e,i,s,n){if(!i.offset)return!1;var o=(e.positionAbs||e.position.absolute).left+e.margins.left,a=(e.positionAbs||e.position.absolute).top+e.margins.top,r=o+e.helperProportions.width,h=a+e.helperProportions.height,l=i.offset.left,c=i.offset.top,u=l+i.proportions().width,d=c+i.proportions().height;switch(s){case"fit":return o>=l&&u>=r&&a>=c&&d>=h;case"intersect":return o+e.helperProportions.width/2>l&&u>r-e.helperProportions.width/2&&a+e.helperProportions.height/2>c&&d>h-e.helperProportions.height/2;case"pointer":return t(n.pageY,c,i.proportions().height)&&t(n.pageX,l,i.proportions().width);case"touch":return(a>=c&&d>=a||h>=c&&d>=h||c>a&&h>d)&&(o>=l&&u>=o||r>=l&&u>=r||l>o&&r>u);default:return!1}}}();t.ui.ddmanager={current:null,droppables:{"default":[]},prepareOffsets:function(e,i){var s,n,o=t.ui.ddmanager.droppables[e.options.scope]||[],a=i?i.type:null,r=(e.currentItem||e.element).find(":data(ui-droppable)").addBack();t:for(s=0;o.length>s;s++)if(!(o[s].options.disabled||e&&!o[s].accept.call(o[s].element[0],e.currentItem||e.element))){for(n=0;r.length>n;n++)if(r[n]===o[s].element[0]){o[s].proportions().height=0;continue t}o[s].visible="none"!==o[s].element.css("display"),o[s].visible&&("mousedown"===a&&o[s]._activate.call(o[s],i),o[s].offset=o[s].element.offset(),o[s].proportions({width:o[s].element[0].offsetWidth,height:o[s].element[0].offsetHeight}))}},drop:function(e,i){var s=!1;return t.each((t.ui.ddmanager.droppables[e.options.scope]||[]).slice(),function(){this.options&&(!this.options.disabled&&this.visible&&v(e,this,this.options.tolerance,i)&&(s=this._drop.call(this,i)||s),!this.options.disabled&&this.visible&&this.accept.call(this.element[0],e.currentItem||e.element)&&(this.isout=!0,this.isover=!1,this._deactivate.call(this,i)))}),s},dragStart:function(e,i){e.element.parentsUntil("body").on("scroll.droppable",function(){e.options.refreshPositions||t.ui.ddmanager.prepareOffsets(e,i)})},drag:function(e,i){e.options.refreshPositions&&t.ui.ddmanager.prepareOffsets(e,i),t.each(t.ui.ddmanager.droppables[e.options.scope]||[],function(){if(!this.options.disabled&&!this.greedyChild&&this.visible){var s,n,o,a=v(e,this,this.options.tolerance,i),r=!a&&this.isover?"isout":a&&!this.isover?"isover":null;r&&(this.options.greedy&&(n=this.options.scope,o=this.element.parents(":data(ui-droppable)").filter(function(){return t(this).droppable("instance").options.scope===n}),o.length&&(s=t(o[0]).droppable("instance"),s.greedyChild="isover"===r)),s&&"isover"===r&&(s.isover=!1,s.isout=!0,s._out.call(s,i)),this[r]=!0,this["isout"===r?"isover":"isout"]=!1,this["isover"===r?"_over":"_out"].call(this,i),s&&"isout"===r&&(s.isout=!1,s.isover=!0,s._over.call(s,i)))}})},dragStop:function(e,i){e.element.parentsUntil("body").off("scroll.droppable"),e.options.refreshPositions||t.ui.ddmanager.prepareOffsets(e,i)}},t.uiBackCompat!==!1&&t.widget("ui.droppable",t.ui.droppable,{options:{hoverClass:!1,activeClass:!1},_addActiveClass:function(){this._super(),this.options.activeClass&&this.element.addClass(this.options.activeClass)},_removeActiveClass:function(){this._super(),this.options.activeClass&&this.element.removeClass(this.options.activeClass)},_addHoverClass:function(){this._super(),this.options.hoverClass&&this.element.addClass(this.options.hoverClass)},_removeHoverClass:function(){this._super(),this.options.hoverClass&&this.element.removeClass(this.options.hoverClass)}}),t.ui.droppable,t.widget("ui.progressbar",{version:"1.12.1",options:{classes:{"ui-progressbar":"ui-corner-all","ui-progressbar-value":"ui-corner-left","ui-progressbar-complete":"ui-corner-right"},max:100,value:0,change:null,complete:null},min:0,_create:function(){this.oldValue=this.options.value=this._constrainedValue(),this.element.attr({role:"progressbar","aria-valuemin":this.min}),this._addClass("ui-progressbar","ui-widget ui-widget-content"),this.valueDiv=t("<div>").appendTo(this.element),this._addClass(this.valueDiv,"ui-progressbar-value","ui-widget-header"),this._refreshValue()},_destroy:function(){this.element.removeAttr("role aria-valuemin aria-valuemax aria-valuenow"),this.valueDiv.remove()},value:function(t){return void 0===t?this.options.value:(this.options.value=this._constrainedValue(t),this._refreshValue(),void 0)},_constrainedValue:function(t){return void 0===t&&(t=this.options.value),this.indeterminate=t===!1,"number"!=typeof t&&(t=0),this.indeterminate?!1:Math.min(this.options.max,Math.max(this.min,t))},_setOptions:function(t){var e=t.value;delete t.value,this._super(t),this.options.value=this._constrainedValue(e),this._refreshValue()},_setOption:function(t,e){"max"===t&&(e=Math.max(this.min,e)),this._super(t,e)},_setOptionDisabled:function(t){this._super(t),this.element.attr("aria-disabled",t),this._toggleClass(null,"ui-state-disabled",!!t)},_percentage:function(){return this.indeterminate?100:100*(this.options.value-this.min)/(this.options.max-this.min)},_refreshValue:function(){var e=this.options.value,i=this._percentage();this.valueDiv.toggle(this.indeterminate||e>this.min).width(i.toFixed(0)+"%"),this._toggleClass(this.valueDiv,"ui-progressbar-complete",null,e===this.options.max)._toggleClass("ui-progressbar-indeterminate",null,this.indeterminate),this.indeterminate?(this.element.removeAttr("aria-valuenow"),this.overlayDiv||(this.overlayDiv=t("<div>").appendTo(this.valueDiv),this._addClass(this.overlayDiv,"ui-progressbar-overlay"))):(this.element.attr({"aria-valuemax":this.options.max,"aria-valuenow":e}),this.overlayDiv&&(this.overlayDiv.remove(),this.overlayDiv=null)),this.oldValue!==e&&(this.oldValue=e,this._trigger("change")),e===this.options.max&&this._trigger("complete")}}),t.widget("ui.selectable",t.ui.mouse,{version:"1.12.1",options:{appendTo:"body",autoRefresh:!0,distance:0,filter:"*",tolerance:"touch",selected:null,selecting:null,start:null,stop:null,unselected:null,unselecting:null},_create:function(){var e=this;this._addClass("ui-selectable"),this.dragged=!1,this.refresh=function(){e.elementPos=t(e.element[0]).offset(),e.selectees=t(e.options.filter,e.element[0]),e._addClass(e.selectees,"ui-selectee"),e.selectees.each(function(){var i=t(this),s=i.offset(),n={left:s.left-e.elementPos.left,top:s.top-e.elementPos.top};t.data(this,"selectable-item",{element:this,$element:i,left:n.left,top:n.top,right:n.left+i.outerWidth(),bottom:n.top+i.outerHeight(),startselected:!1,selected:i.hasClass("ui-selected"),selecting:i.hasClass("ui-selecting"),unselecting:i.hasClass("ui-unselecting")})})},this.refresh(),this._mouseInit(),this.helper=t("<div>"),this._addClass(this.helper,"ui-selectable-helper")},_destroy:function(){this.selectees.removeData("selectable-item"),this._mouseDestroy()},_mouseStart:function(e){var i=this,s=this.options;this.opos=[e.pageX,e.pageY],this.elementPos=t(this.element[0]).offset(),this.options.disabled||(this.selectees=t(s.filter,this.element[0]),this._trigger("start",e),t(s.appendTo).append(this.helper),this.helper.css({left:e.pageX,top:e.pageY,width:0,height:0}),s.autoRefresh&&this.refresh(),this.selectees.filter(".ui-selected").each(function(){var s=t.data(this,"selectable-item");s.startselected=!0,e.metaKey||e.ctrlKey||(i._removeClass(s.$element,"ui-selected"),s.selected=!1,i._addClass(s.$element,"ui-unselecting"),s.unselecting=!0,i._trigger("unselecting",e,{unselecting:s.element}))}),t(e.target).parents().addBack().each(function(){var s,n=t.data(this,"selectable-item");return n?(s=!e.metaKey&&!e.ctrlKey||!n.$element.hasClass("ui-selected"),i._removeClass(n.$element,s?"ui-unselecting":"ui-selected")._addClass(n.$element,s?"ui-selecting":"ui-unselecting"),n.unselecting=!s,n.selecting=s,n.selected=s,s?i._trigger("selecting",e,{selecting:n.element}):i._trigger("unselecting",e,{unselecting:n.element}),!1):void 0}))},_mouseDrag:function(e){if(this.dragged=!0,!this.options.disabled){var i,s=this,n=this.options,o=this.opos[0],a=this.opos[1],r=e.pageX,h=e.pageY;return o>r&&(i=r,r=o,o=i),a>h&&(i=h,h=a,a=i),this.helper.css({left:o,top:a,width:r-o,height:h-a}),this.selectees.each(function(){var i=t.data(this,"selectable-item"),l=!1,c={};i&&i.element!==s.element[0]&&(c.left=i.left+s.elementPos.left,c.right=i.right+s.elementPos.left,c.top=i.top+s.elementPos.top,c.bottom=i.bottom+s.elementPos.top,"touch"===n.tolerance?l=!(c.left>r||o>c.right||c.top>h||a>c.bottom):"fit"===n.tolerance&&(l=c.left>o&&r>c.right&&c.top>a&&h>c.bottom),l?(i.selected&&(s._removeClass(i.$element,"ui-selected"),i.selected=!1),i.unselecting&&(s._removeClass(i.$element,"ui-unselecting"),i.unselecting=!1),i.selecting||(s._addClass(i.$element,"ui-selecting"),i.selecting=!0,s._trigger("selecting",e,{selecting:i.element}))):(i.selecting&&((e.metaKey||e.ctrlKey)&&i.startselected?(s._removeClass(i.$element,"ui-selecting"),i.selecting=!1,s._addClass(i.$element,"ui-selected"),i.selected=!0):(s._removeClass(i.$element,"ui-selecting"),i.selecting=!1,i.startselected&&(s._addClass(i.$element,"ui-unselecting"),i.unselecting=!0),s._trigger("unselecting",e,{unselecting:i.element}))),i.selected&&(e.metaKey||e.ctrlKey||i.startselected||(s._removeClass(i.$element,"ui-selected"),i.selected=!1,s._addClass(i.$element,"ui-unselecting"),i.unselecting=!0,s._trigger("unselecting",e,{unselecting:i.element})))))}),!1}},_mouseStop:function(e){var i=this;return this.dragged=!1,t(".ui-unselecting",this.element[0]).each(function(){var s=t.data(this,"selectable-item");i._removeClass(s.$element,"ui-unselecting"),s.unselecting=!1,s.startselected=!1,i._trigger("unselected",e,{unselected:s.element})}),t(".ui-selecting",this.element[0]).each(function(){var s=t.data(this,"selectable-item");i._removeClass(s.$element,"ui-selecting")._addClass(s.$element,"ui-selected"),s.selecting=!1,s.selected=!0,s.startselected=!0,i._trigger("selected",e,{selected:s.element})}),this._trigger("stop",e),this.helper.remove(),!1}}),t.widget("ui.selectmenu",[t.ui.formResetMixin,{version:"1.12.1",defaultElement:"<select>",options:{appendTo:null,classes:{"ui-selectmenu-button-open":"ui-corner-top","ui-selectmenu-button-closed":"ui-corner-all"},disabled:null,icons:{button:"ui-icon-triangle-1-s"},position:{my:"left top",at:"left bottom",collision:"none"},width:!1,change:null,close:null,focus:null,open:null,select:null},_create:function(){var e=this.element.uniqueId().attr("id");this.ids={element:e,button:e+"-button",menu:e+"-menu"},this._drawButton(),this._drawMenu(),this._bindFormResetHandler(),this._rendered=!1,this.menuItems=t()},_drawButton:function(){var e,i=this,s=this._parseOption(this.element.find("option:selected"),this.element[0].selectedIndex);this.labels=this.element.labels().attr("for",this.ids.button),this._on(this.labels,{click:function(t){this.button.focus(),t.preventDefault()}}),this.element.hide(),this.button=t("<span>",{tabindex:this.options.disabled?-1:0,id:this.ids.button,role:"combobox","aria-expanded":"false","aria-autocomplete":"list","aria-owns":this.ids.menu,"aria-haspopup":"true",title:this.element.attr("title")}).insertAfter(this.element),this._addClass(this.button,"ui-selectmenu-button ui-selectmenu-button-closed","ui-button ui-widget"),e=t("<span>").appendTo(this.button),this._addClass(e,"ui-selectmenu-icon","ui-icon "+this.options.icons.button),this.buttonItem=this._renderButtonItem(s).appendTo(this.button),this.options.width!==!1&&this._resizeButton(),this._on(this.button,this._buttonEvents),this.button.one("focusin",function(){i._rendered||i._refreshMenu()})},_drawMenu:function(){var e=this;this.menu=t("<ul>",{"aria-hidden":"true","aria-labelledby":this.ids.button,id:this.ids.menu}),this.menuWrap=t("<div>").append(this.menu),this._addClass(this.menuWrap,"ui-selectmenu-menu","ui-front"),this.menuWrap.appendTo(this._appendTo()),this.menuInstance=this.menu.menu({classes:{"ui-menu":"ui-corner-bottom"},role:"listbox",select:function(t,i){t.preventDefault(),e._setSelection(),e._select(i.item.data("ui-selectmenu-item"),t)},focus:function(t,i){var s=i.item.data("ui-selectmenu-item");null!=e.focusIndex&&s.index!==e.focusIndex&&(e._trigger("focus",t,{item:s}),e.isOpen||e._select(s,t)),e.focusIndex=s.index,e.button.attr("aria-activedescendant",e.menuItems.eq(s.index).attr("id"))}}).menu("instance"),this.menuInstance._off(this.menu,"mouseleave"),this.menuInstance._closeOnDocumentClick=function(){return!1},this.menuInstance._isDivider=function(){return!1}},refresh:function(){this._refreshMenu(),this.buttonItem.replaceWith(this.buttonItem=this._renderButtonItem(this._getSelectedItem().data("ui-selectmenu-item")||{})),null===this.options.width&&this._resizeButton()},_refreshMenu:function(){var t,e=this.element.find("option");this.menu.empty(),this._parseOptions(e),this._renderMenu(this.menu,this.items),this.menuInstance.refresh(),this.menuItems=this.menu.find("li").not(".ui-selectmenu-optgroup").find(".ui-menu-item-wrapper"),this._rendered=!0,e.length&&(t=this._getSelectedItem(),this.menuInstance.focus(null,t),this._setAria(t.data("ui-selectmenu-item")),this._setOption("disabled",this.element.prop("disabled")))},open:function(t){this.options.disabled||(this._rendered?(this._removeClass(this.menu.find(".ui-state-active"),null,"ui-state-active"),this.menuInstance.focus(null,this._getSelectedItem())):this._refreshMenu(),this.menuItems.length&&(this.isOpen=!0,this._toggleAttr(),this._resizeMenu(),this._position(),this._on(this.document,this._documentClick),this._trigger("open",t)))},_position:function(){this.menuWrap.position(t.extend({of:this.button},this.options.position))},close:function(t){this.isOpen&&(this.isOpen=!1,this._toggleAttr(),this.range=null,this._off(this.document),this._trigger("close",t))},widget:function(){return this.button},menuWidget:function(){return this.menu},_renderButtonItem:function(e){var i=t("<span>");return this._setText(i,e.label),this._addClass(i,"ui-selectmenu-text"),i},_renderMenu:function(e,i){var s=this,n="";t.each(i,function(i,o){var a;o.optgroup!==n&&(a=t("<li>",{text:o.optgroup}),s._addClass(a,"ui-selectmenu-optgroup","ui-menu-divider"+(o.element.parent("optgroup").prop("disabled")?" ui-state-disabled":"")),a.appendTo(e),n=o.optgroup),s._renderItemData(e,o)})},_renderItemData:function(t,e){return this._renderItem(t,e).data("ui-selectmenu-item",e)},_renderItem:function(e,i){var s=t("<li>"),n=t("<div>",{title:i.element.attr("title")});return i.disabled&&this._addClass(s,null,"ui-state-disabled"),this._setText(n,i.label),s.append(n).appendTo(e)},_setText:function(t,e){e?t.text(e):t.html("&#160;")},_move:function(t,e){var i,s,n=".ui-menu-item";this.isOpen?i=this.menuItems.eq(this.focusIndex).parent("li"):(i=this.menuItems.eq(this.element[0].selectedIndex).parent("li"),n+=":not(.ui-state-disabled)"),s="first"===t||"last"===t?i["first"===t?"prevAll":"nextAll"](n).eq(-1):i[t+"All"](n).eq(0),s.length&&this.menuInstance.focus(e,s)},_getSelectedItem:function(){return this.menuItems.eq(this.element[0].selectedIndex).parent("li")},_toggle:function(t){this[this.isOpen?"close":"open"](t)},_setSelection:function(){var t;this.range&&(window.getSelection?(t=window.getSelection(),t.removeAllRanges(),t.addRange(this.range)):this.range.select(),this.button.focus())},_documentClick:{mousedown:function(e){this.isOpen&&(t(e.target).closest(".ui-selectmenu-menu, #"+t.ui.escapeSelector(this.ids.button)).length||this.close(e))}},_buttonEvents:{mousedown:function(){var t;window.getSelection?(t=window.getSelection(),t.rangeCount&&(this.range=t.getRangeAt(0))):this.range=document.selection.createRange()},click:function(t){this._setSelection(),this._toggle(t)},keydown:function(e){var i=!0;switch(e.keyCode){case t.ui.keyCode.TAB:case t.ui.keyCode.ESCAPE:this.close(e),i=!1;break;case t.ui.keyCode.ENTER:this.isOpen&&this._selectFocusedItem(e);break;case t.ui.keyCode.UP:e.altKey?this._toggle(e):this._move("prev",e);break;case t.ui.keyCode.DOWN:e.altKey?this._toggle(e):this._move("next",e);break;case t.ui.keyCode.SPACE:this.isOpen?this._selectFocusedItem(e):this._toggle(e);break;case t.ui.keyCode.LEFT:this._move("prev",e);break;case t.ui.keyCode.RIGHT:this._move("next",e);break;case t.ui.keyCode.HOME:case t.ui.keyCode.PAGE_UP:this._move("first",e);break;case t.ui.keyCode.END:case t.ui.keyCode.PAGE_DOWN:this._move("last",e);break;default:this.menu.trigger(e),i=!1}i&&e.preventDefault()}},_selectFocusedItem:function(t){var e=this.menuItems.eq(this.focusIndex).parent("li");e.hasClass("ui-state-disabled")||this._select(e.data("ui-selectmenu-item"),t)},_select:function(t,e){var i=this.element[0].selectedIndex;this.element[0].selectedIndex=t.index,this.buttonItem.replaceWith(this.buttonItem=this._renderButtonItem(t)),this._setAria(t),this._trigger("select",e,{item:t}),t.index!==i&&this._trigger("change",e,{item:t}),this.close(e)},_setAria:function(t){var e=this.menuItems.eq(t.index).attr("id");this.button.attr({"aria-labelledby":e,"aria-activedescendant":e}),this.menu.attr("aria-activedescendant",e)},_setOption:function(t,e){if("icons"===t){var i=this.button.find("span.ui-icon");this._removeClass(i,null,this.options.icons.button)._addClass(i,null,e.button)}this._super(t,e),"appendTo"===t&&this.menuWrap.appendTo(this._appendTo()),"width"===t&&this._resizeButton()},_setOptionDisabled:function(t){this._super(t),this.menuInstance.option("disabled",t),this.button.attr("aria-disabled",t),this._toggleClass(this.button,null,"ui-state-disabled",t),this.element.prop("disabled",t),t?(this.button.attr("tabindex",-1),this.close()):this.button.attr("tabindex",0)},_appendTo:function(){var e=this.options.appendTo;return e&&(e=e.jquery||e.nodeType?t(e):this.document.find(e).eq(0)),e&&e[0]||(e=this.element.closest(".ui-front, dialog")),e.length||(e=this.document[0].body),e},_toggleAttr:function(){this.button.attr("aria-expanded",this.isOpen),this._removeClass(this.button,"ui-selectmenu-button-"+(this.isOpen?"closed":"open"))._addClass(this.button,"ui-selectmenu-button-"+(this.isOpen?"open":"closed"))._toggleClass(this.menuWrap,"ui-selectmenu-open",null,this.isOpen),this.menu.attr("aria-hidden",!this.isOpen)},_resizeButton:function(){var t=this.options.width;return t===!1?(this.button.css("width",""),void 0):(null===t&&(t=this.element.show().outerWidth(),this.element.hide()),this.button.outerWidth(t),void 0)},_resizeMenu:function(){this.menu.outerWidth(Math.max(this.button.outerWidth(),this.menu.width("").outerWidth()+1))},_getCreateOptions:function(){var t=this._super();return t.disabled=this.element.prop("disabled"),t},_parseOptions:function(e){var i=this,s=[];e.each(function(e,n){s.push(i._parseOption(t(n),e))}),this.items=s},_parseOption:function(t,e){var i=t.parent("optgroup");return{element:t,index:e,value:t.val(),label:t.text(),optgroup:i.attr("label")||"",disabled:i.prop("disabled")||t.prop("disabled")}},_destroy:function(){this._unbindFormResetHandler(),this.menuWrap.remove(),this.button.remove(),this.element.show(),this.element.removeUniqueId(),this.labels.attr("for",this.ids.element)}}]),t.widget("ui.slider",t.ui.mouse,{version:"1.12.1",widgetEventPrefix:"slide",options:{animate:!1,classes:{"ui-slider":"ui-corner-all","ui-slider-handle":"ui-corner-all","ui-slider-range":"ui-corner-all ui-widget-header"},distance:0,max:100,min:0,orientation:"horizontal",range:!1,step:1,value:0,values:null,change:null,slide:null,start:null,stop:null},numPages:5,_create:function(){this._keySliding=!1,this._mouseSliding=!1,this._animateOff=!0,this._handleIndex=null,this._detectOrientation(),this._mouseInit(),this._calculateNewMax(),this._addClass("ui-slider ui-slider-"+this.orientation,"ui-widget ui-widget-content"),this._refresh(),this._animateOff=!1
},_refresh:function(){this._createRange(),this._createHandles(),this._setupEvents(),this._refreshValue()},_createHandles:function(){var e,i,s=this.options,n=this.element.find(".ui-slider-handle"),o="<span tabindex='0'></span>",a=[];for(i=s.values&&s.values.length||1,n.length>i&&(n.slice(i).remove(),n=n.slice(0,i)),e=n.length;i>e;e++)a.push(o);this.handles=n.add(t(a.join("")).appendTo(this.element)),this._addClass(this.handles,"ui-slider-handle","ui-state-default"),this.handle=this.handles.eq(0),this.handles.each(function(e){t(this).data("ui-slider-handle-index",e).attr("tabIndex",0)})},_createRange:function(){var e=this.options;e.range?(e.range===!0&&(e.values?e.values.length&&2!==e.values.length?e.values=[e.values[0],e.values[0]]:t.isArray(e.values)&&(e.values=e.values.slice(0)):e.values=[this._valueMin(),this._valueMin()]),this.range&&this.range.length?(this._removeClass(this.range,"ui-slider-range-min ui-slider-range-max"),this.range.css({left:"",bottom:""})):(this.range=t("<div>").appendTo(this.element),this._addClass(this.range,"ui-slider-range")),("min"===e.range||"max"===e.range)&&this._addClass(this.range,"ui-slider-range-"+e.range)):(this.range&&this.range.remove(),this.range=null)},_setupEvents:function(){this._off(this.handles),this._on(this.handles,this._handleEvents),this._hoverable(this.handles),this._focusable(this.handles)},_destroy:function(){this.handles.remove(),this.range&&this.range.remove(),this._mouseDestroy()},_mouseCapture:function(e){var i,s,n,o,a,r,h,l,c=this,u=this.options;return u.disabled?!1:(this.elementSize={width:this.element.outerWidth(),height:this.element.outerHeight()},this.elementOffset=this.element.offset(),i={x:e.pageX,y:e.pageY},s=this._normValueFromMouse(i),n=this._valueMax()-this._valueMin()+1,this.handles.each(function(e){var i=Math.abs(s-c.values(e));(n>i||n===i&&(e===c._lastChangedValue||c.values(e)===u.min))&&(n=i,o=t(this),a=e)}),r=this._start(e,a),r===!1?!1:(this._mouseSliding=!0,this._handleIndex=a,this._addClass(o,null,"ui-state-active"),o.trigger("focus"),h=o.offset(),l=!t(e.target).parents().addBack().is(".ui-slider-handle"),this._clickOffset=l?{left:0,top:0}:{left:e.pageX-h.left-o.width()/2,top:e.pageY-h.top-o.height()/2-(parseInt(o.css("borderTopWidth"),10)||0)-(parseInt(o.css("borderBottomWidth"),10)||0)+(parseInt(o.css("marginTop"),10)||0)},this.handles.hasClass("ui-state-hover")||this._slide(e,a,s),this._animateOff=!0,!0))},_mouseStart:function(){return!0},_mouseDrag:function(t){var e={x:t.pageX,y:t.pageY},i=this._normValueFromMouse(e);return this._slide(t,this._handleIndex,i),!1},_mouseStop:function(t){return this._removeClass(this.handles,null,"ui-state-active"),this._mouseSliding=!1,this._stop(t,this._handleIndex),this._change(t,this._handleIndex),this._handleIndex=null,this._clickOffset=null,this._animateOff=!1,!1},_detectOrientation:function(){this.orientation="vertical"===this.options.orientation?"vertical":"horizontal"},_normValueFromMouse:function(t){var e,i,s,n,o;return"horizontal"===this.orientation?(e=this.elementSize.width,i=t.x-this.elementOffset.left-(this._clickOffset?this._clickOffset.left:0)):(e=this.elementSize.height,i=t.y-this.elementOffset.top-(this._clickOffset?this._clickOffset.top:0)),s=i/e,s>1&&(s=1),0>s&&(s=0),"vertical"===this.orientation&&(s=1-s),n=this._valueMax()-this._valueMin(),o=this._valueMin()+s*n,this._trimAlignValue(o)},_uiHash:function(t,e,i){var s={handle:this.handles[t],handleIndex:t,value:void 0!==e?e:this.value()};return this._hasMultipleValues()&&(s.value=void 0!==e?e:this.values(t),s.values=i||this.values()),s},_hasMultipleValues:function(){return this.options.values&&this.options.values.length},_start:function(t,e){return this._trigger("start",t,this._uiHash(e))},_slide:function(t,e,i){var s,n,o=this.value(),a=this.values();this._hasMultipleValues()&&(n=this.values(e?0:1),o=this.values(e),2===this.options.values.length&&this.options.range===!0&&(i=0===e?Math.min(n,i):Math.max(n,i)),a[e]=i),i!==o&&(s=this._trigger("slide",t,this._uiHash(e,i,a)),s!==!1&&(this._hasMultipleValues()?this.values(e,i):this.value(i)))},_stop:function(t,e){this._trigger("stop",t,this._uiHash(e))},_change:function(t,e){this._keySliding||this._mouseSliding||(this._lastChangedValue=e,this._trigger("change",t,this._uiHash(e)))},value:function(t){return arguments.length?(this.options.value=this._trimAlignValue(t),this._refreshValue(),this._change(null,0),void 0):this._value()},values:function(e,i){var s,n,o;if(arguments.length>1)return this.options.values[e]=this._trimAlignValue(i),this._refreshValue(),this._change(null,e),void 0;if(!arguments.length)return this._values();if(!t.isArray(arguments[0]))return this._hasMultipleValues()?this._values(e):this.value();for(s=this.options.values,n=arguments[0],o=0;s.length>o;o+=1)s[o]=this._trimAlignValue(n[o]),this._change(null,o);this._refreshValue()},_setOption:function(e,i){var s,n=0;switch("range"===e&&this.options.range===!0&&("min"===i?(this.options.value=this._values(0),this.options.values=null):"max"===i&&(this.options.value=this._values(this.options.values.length-1),this.options.values=null)),t.isArray(this.options.values)&&(n=this.options.values.length),this._super(e,i),e){case"orientation":this._detectOrientation(),this._removeClass("ui-slider-horizontal ui-slider-vertical")._addClass("ui-slider-"+this.orientation),this._refreshValue(),this.options.range&&this._refreshRange(i),this.handles.css("horizontal"===i?"bottom":"left","");break;case"value":this._animateOff=!0,this._refreshValue(),this._change(null,0),this._animateOff=!1;break;case"values":for(this._animateOff=!0,this._refreshValue(),s=n-1;s>=0;s--)this._change(null,s);this._animateOff=!1;break;case"step":case"min":case"max":this._animateOff=!0,this._calculateNewMax(),this._refreshValue(),this._animateOff=!1;break;case"range":this._animateOff=!0,this._refresh(),this._animateOff=!1}},_setOptionDisabled:function(t){this._super(t),this._toggleClass(null,"ui-state-disabled",!!t)},_value:function(){var t=this.options.value;return t=this._trimAlignValue(t)},_values:function(t){var e,i,s;if(arguments.length)return e=this.options.values[t],e=this._trimAlignValue(e);if(this._hasMultipleValues()){for(i=this.options.values.slice(),s=0;i.length>s;s+=1)i[s]=this._trimAlignValue(i[s]);return i}return[]},_trimAlignValue:function(t){if(this._valueMin()>=t)return this._valueMin();if(t>=this._valueMax())return this._valueMax();var e=this.options.step>0?this.options.step:1,i=(t-this._valueMin())%e,s=t-i;return 2*Math.abs(i)>=e&&(s+=i>0?e:-e),parseFloat(s.toFixed(5))},_calculateNewMax:function(){var t=this.options.max,e=this._valueMin(),i=this.options.step,s=Math.round((t-e)/i)*i;t=s+e,t>this.options.max&&(t-=i),this.max=parseFloat(t.toFixed(this._precision()))},_precision:function(){var t=this._precisionOf(this.options.step);return null!==this.options.min&&(t=Math.max(t,this._precisionOf(this.options.min))),t},_precisionOf:function(t){var e=""+t,i=e.indexOf(".");return-1===i?0:e.length-i-1},_valueMin:function(){return this.options.min},_valueMax:function(){return this.max},_refreshRange:function(t){"vertical"===t&&this.range.css({width:"",left:""}),"horizontal"===t&&this.range.css({height:"",bottom:""})},_refreshValue:function(){var e,i,s,n,o,a=this.options.range,r=this.options,h=this,l=this._animateOff?!1:r.animate,c={};this._hasMultipleValues()?this.handles.each(function(s){i=100*((h.values(s)-h._valueMin())/(h._valueMax()-h._valueMin())),c["horizontal"===h.orientation?"left":"bottom"]=i+"%",t(this).stop(1,1)[l?"animate":"css"](c,r.animate),h.options.range===!0&&("horizontal"===h.orientation?(0===s&&h.range.stop(1,1)[l?"animate":"css"]({left:i+"%"},r.animate),1===s&&h.range[l?"animate":"css"]({width:i-e+"%"},{queue:!1,duration:r.animate})):(0===s&&h.range.stop(1,1)[l?"animate":"css"]({bottom:i+"%"},r.animate),1===s&&h.range[l?"animate":"css"]({height:i-e+"%"},{queue:!1,duration:r.animate}))),e=i}):(s=this.value(),n=this._valueMin(),o=this._valueMax(),i=o!==n?100*((s-n)/(o-n)):0,c["horizontal"===this.orientation?"left":"bottom"]=i+"%",this.handle.stop(1,1)[l?"animate":"css"](c,r.animate),"min"===a&&"horizontal"===this.orientation&&this.range.stop(1,1)[l?"animate":"css"]({width:i+"%"},r.animate),"max"===a&&"horizontal"===this.orientation&&this.range.stop(1,1)[l?"animate":"css"]({width:100-i+"%"},r.animate),"min"===a&&"vertical"===this.orientation&&this.range.stop(1,1)[l?"animate":"css"]({height:i+"%"},r.animate),"max"===a&&"vertical"===this.orientation&&this.range.stop(1,1)[l?"animate":"css"]({height:100-i+"%"},r.animate))},_handleEvents:{keydown:function(e){var i,s,n,o,a=t(e.target).data("ui-slider-handle-index");switch(e.keyCode){case t.ui.keyCode.HOME:case t.ui.keyCode.END:case t.ui.keyCode.PAGE_UP:case t.ui.keyCode.PAGE_DOWN:case t.ui.keyCode.UP:case t.ui.keyCode.RIGHT:case t.ui.keyCode.DOWN:case t.ui.keyCode.LEFT:if(e.preventDefault(),!this._keySliding&&(this._keySliding=!0,this._addClass(t(e.target),null,"ui-state-active"),i=this._start(e,a),i===!1))return}switch(o=this.options.step,s=n=this._hasMultipleValues()?this.values(a):this.value(),e.keyCode){case t.ui.keyCode.HOME:n=this._valueMin();break;case t.ui.keyCode.END:n=this._valueMax();break;case t.ui.keyCode.PAGE_UP:n=this._trimAlignValue(s+(this._valueMax()-this._valueMin())/this.numPages);break;case t.ui.keyCode.PAGE_DOWN:n=this._trimAlignValue(s-(this._valueMax()-this._valueMin())/this.numPages);break;case t.ui.keyCode.UP:case t.ui.keyCode.RIGHT:if(s===this._valueMax())return;n=this._trimAlignValue(s+o);break;case t.ui.keyCode.DOWN:case t.ui.keyCode.LEFT:if(s===this._valueMin())return;n=this._trimAlignValue(s-o)}this._slide(e,a,n)},keyup:function(e){var i=t(e.target).data("ui-slider-handle-index");this._keySliding&&(this._keySliding=!1,this._stop(e,i),this._change(e,i),this._removeClass(t(e.target),null,"ui-state-active"))}}}),t.widget("ui.sortable",t.ui.mouse,{version:"1.12.1",widgetEventPrefix:"sort",ready:!1,options:{appendTo:"parent",axis:!1,connectWith:!1,containment:!1,cursor:"auto",cursorAt:!1,dropOnEmpty:!0,forcePlaceholderSize:!1,forceHelperSize:!1,grid:!1,handle:!1,helper:"original",items:"> *",opacity:!1,placeholder:!1,revert:!1,scroll:!0,scrollSensitivity:20,scrollSpeed:20,scope:"default",tolerance:"intersect",zIndex:1e3,activate:null,beforeStop:null,change:null,deactivate:null,out:null,over:null,receive:null,remove:null,sort:null,start:null,stop:null,update:null},_isOverAxis:function(t,e,i){return t>=e&&e+i>t},_isFloating:function(t){return/left|right/.test(t.css("float"))||/inline|table-cell/.test(t.css("display"))},_create:function(){this.containerCache={},this._addClass("ui-sortable"),this.refresh(),this.offset=this.element.offset(),this._mouseInit(),this._setHandleClassName(),this.ready=!0},_setOption:function(t,e){this._super(t,e),"handle"===t&&this._setHandleClassName()},_setHandleClassName:function(){var e=this;this._removeClass(this.element.find(".ui-sortable-handle"),"ui-sortable-handle"),t.each(this.items,function(){e._addClass(this.instance.options.handle?this.item.find(this.instance.options.handle):this.item,"ui-sortable-handle")})},_destroy:function(){this._mouseDestroy();for(var t=this.items.length-1;t>=0;t--)this.items[t].item.removeData(this.widgetName+"-item");return this},_mouseCapture:function(e,i){var s=null,n=!1,o=this;return this.reverting?!1:this.options.disabled||"static"===this.options.type?!1:(this._refreshItems(e),t(e.target).parents().each(function(){return t.data(this,o.widgetName+"-item")===o?(s=t(this),!1):void 0}),t.data(e.target,o.widgetName+"-item")===o&&(s=t(e.target)),s?!this.options.handle||i||(t(this.options.handle,s).find("*").addBack().each(function(){this===e.target&&(n=!0)}),n)?(this.currentItem=s,this._removeCurrentsFromItems(),!0):!1:!1)},_mouseStart:function(e,i,s){var n,o,a=this.options;if(this.currentContainer=this,this.refreshPositions(),this.helper=this._createHelper(e),this._cacheHelperProportions(),this._cacheMargins(),this.scrollParent=this.helper.scrollParent(),this.offset=this.currentItem.offset(),this.offset={top:this.offset.top-this.margins.top,left:this.offset.left-this.margins.left},t.extend(this.offset,{click:{left:e.pageX-this.offset.left,top:e.pageY-this.offset.top},parent:this._getParentOffset(),relative:this._getRelativeOffset()}),this.helper.css("position","absolute"),this.cssPosition=this.helper.css("position"),this.originalPosition=this._generatePosition(e),this.originalPageX=e.pageX,this.originalPageY=e.pageY,a.cursorAt&&this._adjustOffsetFromHelper(a.cursorAt),this.domPosition={prev:this.currentItem.prev()[0],parent:this.currentItem.parent()[0]},this.helper[0]!==this.currentItem[0]&&this.currentItem.hide(),this._createPlaceholder(),a.containment&&this._setContainment(),a.cursor&&"auto"!==a.cursor&&(o=this.document.find("body"),this.storedCursor=o.css("cursor"),o.css("cursor",a.cursor),this.storedStylesheet=t("<style>*{ cursor: "+a.cursor+" !important; }</style>").appendTo(o)),a.opacity&&(this.helper.css("opacity")&&(this._storedOpacity=this.helper.css("opacity")),this.helper.css("opacity",a.opacity)),a.zIndex&&(this.helper.css("zIndex")&&(this._storedZIndex=this.helper.css("zIndex")),this.helper.css("zIndex",a.zIndex)),this.scrollParent[0]!==this.document[0]&&"HTML"!==this.scrollParent[0].tagName&&(this.overflowOffset=this.scrollParent.offset()),this._trigger("start",e,this._uiHash()),this._preserveHelperProportions||this._cacheHelperProportions(),!s)for(n=this.containers.length-1;n>=0;n--)this.containers[n]._trigger("activate",e,this._uiHash(this));return t.ui.ddmanager&&(t.ui.ddmanager.current=this),t.ui.ddmanager&&!a.dropBehaviour&&t.ui.ddmanager.prepareOffsets(this,e),this.dragging=!0,this._addClass(this.helper,"ui-sortable-helper"),this._mouseDrag(e),!0},_mouseDrag:function(e){var i,s,n,o,a=this.options,r=!1;for(this.position=this._generatePosition(e),this.positionAbs=this._convertPositionTo("absolute"),this.lastPositionAbs||(this.lastPositionAbs=this.positionAbs),this.options.scroll&&(this.scrollParent[0]!==this.document[0]&&"HTML"!==this.scrollParent[0].tagName?(this.overflowOffset.top+this.scrollParent[0].offsetHeight-e.pageY<a.scrollSensitivity?this.scrollParent[0].scrollTop=r=this.scrollParent[0].scrollTop+a.scrollSpeed:e.pageY-this.overflowOffset.top<a.scrollSensitivity&&(this.scrollParent[0].scrollTop=r=this.scrollParent[0].scrollTop-a.scrollSpeed),this.overflowOffset.left+this.scrollParent[0].offsetWidth-e.pageX<a.scrollSensitivity?this.scrollParent[0].scrollLeft=r=this.scrollParent[0].scrollLeft+a.scrollSpeed:e.pageX-this.overflowOffset.left<a.scrollSensitivity&&(this.scrollParent[0].scrollLeft=r=this.scrollParent[0].scrollLeft-a.scrollSpeed)):(e.pageY-this.document.scrollTop()<a.scrollSensitivity?r=this.document.scrollTop(this.document.scrollTop()-a.scrollSpeed):this.window.height()-(e.pageY-this.document.scrollTop())<a.scrollSensitivity&&(r=this.document.scrollTop(this.document.scrollTop()+a.scrollSpeed)),e.pageX-this.document.scrollLeft()<a.scrollSensitivity?r=this.document.scrollLeft(this.document.scrollLeft()-a.scrollSpeed):this.window.width()-(e.pageX-this.document.scrollLeft())<a.scrollSensitivity&&(r=this.document.scrollLeft(this.document.scrollLeft()+a.scrollSpeed))),r!==!1&&t.ui.ddmanager&&!a.dropBehaviour&&t.ui.ddmanager.prepareOffsets(this,e)),this.positionAbs=this._convertPositionTo("absolute"),this.options.axis&&"y"===this.options.axis||(this.helper[0].style.left=this.position.left+"px"),this.options.axis&&"x"===this.options.axis||(this.helper[0].style.top=this.position.top+"px"),i=this.items.length-1;i>=0;i--)if(s=this.items[i],n=s.item[0],o=this._intersectsWithPointer(s),o&&s.instance===this.currentContainer&&n!==this.currentItem[0]&&this.placeholder[1===o?"next":"prev"]()[0]!==n&&!t.contains(this.placeholder[0],n)&&("semi-dynamic"===this.options.type?!t.contains(this.element[0],n):!0)){if(this.direction=1===o?"down":"up","pointer"!==this.options.tolerance&&!this._intersectsWithSides(s))break;this._rearrange(e,s),this._trigger("change",e,this._uiHash());break}return this._contactContainers(e),t.ui.ddmanager&&t.ui.ddmanager.drag(this,e),this._trigger("sort",e,this._uiHash()),this.lastPositionAbs=this.positionAbs,!1},_mouseStop:function(e,i){if(e){if(t.ui.ddmanager&&!this.options.dropBehaviour&&t.ui.ddmanager.drop(this,e),this.options.revert){var s=this,n=this.placeholder.offset(),o=this.options.axis,a={};o&&"x"!==o||(a.left=n.left-this.offset.parent.left-this.margins.left+(this.offsetParent[0]===this.document[0].body?0:this.offsetParent[0].scrollLeft)),o&&"y"!==o||(a.top=n.top-this.offset.parent.top-this.margins.top+(this.offsetParent[0]===this.document[0].body?0:this.offsetParent[0].scrollTop)),this.reverting=!0,t(this.helper).animate(a,parseInt(this.options.revert,10)||500,function(){s._clear(e)})}else this._clear(e,i);return!1}},cancel:function(){if(this.dragging){this._mouseUp(new t.Event("mouseup",{target:null})),"original"===this.options.helper?(this.currentItem.css(this._storedCSS),this._removeClass(this.currentItem,"ui-sortable-helper")):this.currentItem.show();for(var e=this.containers.length-1;e>=0;e--)this.containers[e]._trigger("deactivate",null,this._uiHash(this)),this.containers[e].containerCache.over&&(this.containers[e]._trigger("out",null,this._uiHash(this)),this.containers[e].containerCache.over=0)}return this.placeholder&&(this.placeholder[0].parentNode&&this.placeholder[0].parentNode.removeChild(this.placeholder[0]),"original"!==this.options.helper&&this.helper&&this.helper[0].parentNode&&this.helper.remove(),t.extend(this,{helper:null,dragging:!1,reverting:!1,_noFinalSort:null}),this.domPosition.prev?t(this.domPosition.prev).after(this.currentItem):t(this.domPosition.parent).prepend(this.currentItem)),this},serialize:function(e){var i=this._getItemsAsjQuery(e&&e.connected),s=[];return e=e||{},t(i).each(function(){var i=(t(e.item||this).attr(e.attribute||"id")||"").match(e.expression||/(.+)[\-=_](.+)/);i&&s.push((e.key||i[1]+"[]")+"="+(e.key&&e.expression?i[1]:i[2]))}),!s.length&&e.key&&s.push(e.key+"="),s.join("&")},toArray:function(e){var i=this._getItemsAsjQuery(e&&e.connected),s=[];return e=e||{},i.each(function(){s.push(t(e.item||this).attr(e.attribute||"id")||"")}),s},_intersectsWith:function(t){var e=this.positionAbs.left,i=e+this.helperProportions.width,s=this.positionAbs.top,n=s+this.helperProportions.height,o=t.left,a=o+t.width,r=t.top,h=r+t.height,l=this.offset.click.top,c=this.offset.click.left,u="x"===this.options.axis||s+l>r&&h>s+l,d="y"===this.options.axis||e+c>o&&a>e+c,p=u&&d;return"pointer"===this.options.tolerance||this.options.forcePointerForContainers||"pointer"!==this.options.tolerance&&this.helperProportions[this.floating?"width":"height"]>t[this.floating?"width":"height"]?p:e+this.helperProportions.width/2>o&&a>i-this.helperProportions.width/2&&s+this.helperProportions.height/2>r&&h>n-this.helperProportions.height/2},_intersectsWithPointer:function(t){var e,i,s="x"===this.options.axis||this._isOverAxis(this.positionAbs.top+this.offset.click.top,t.top,t.height),n="y"===this.options.axis||this._isOverAxis(this.positionAbs.left+this.offset.click.left,t.left,t.width),o=s&&n;return o?(e=this._getDragVerticalDirection(),i=this._getDragHorizontalDirection(),this.floating?"right"===i||"down"===e?2:1:e&&("down"===e?2:1)):!1},_intersectsWithSides:function(t){var e=this._isOverAxis(this.positionAbs.top+this.offset.click.top,t.top+t.height/2,t.height),i=this._isOverAxis(this.positionAbs.left+this.offset.click.left,t.left+t.width/2,t.width),s=this._getDragVerticalDirection(),n=this._getDragHorizontalDirection();return this.floating&&n?"right"===n&&i||"left"===n&&!i:s&&("down"===s&&e||"up"===s&&!e)},_getDragVerticalDirection:function(){var t=this.positionAbs.top-this.lastPositionAbs.top;return 0!==t&&(t>0?"down":"up")},_getDragHorizontalDirection:function(){var t=this.positionAbs.left-this.lastPositionAbs.left;return 0!==t&&(t>0?"right":"left")},refresh:function(t){return this._refreshItems(t),this._setHandleClassName(),this.refreshPositions(),this},_connectWith:function(){var t=this.options;return t.connectWith.constructor===String?[t.connectWith]:t.connectWith},_getItemsAsjQuery:function(e){function i(){r.push(this)}var s,n,o,a,r=[],h=[],l=this._connectWith();if(l&&e)for(s=l.length-1;s>=0;s--)for(o=t(l[s],this.document[0]),n=o.length-1;n>=0;n--)a=t.data(o[n],this.widgetFullName),a&&a!==this&&!a.options.disabled&&h.push([t.isFunction(a.options.items)?a.options.items.call(a.element):t(a.options.items,a.element).not(".ui-sortable-helper").not(".ui-sortable-placeholder"),a]);for(h.push([t.isFunction(this.options.items)?this.options.items.call(this.element,null,{options:this.options,item:this.currentItem}):t(this.options.items,this.element).not(".ui-sortable-helper").not(".ui-sortable-placeholder"),this]),s=h.length-1;s>=0;s--)h[s][0].each(i);return t(r)},_removeCurrentsFromItems:function(){var e=this.currentItem.find(":data("+this.widgetName+"-item)");this.items=t.grep(this.items,function(t){for(var i=0;e.length>i;i++)if(e[i]===t.item[0])return!1;return!0})},_refreshItems:function(e){this.items=[],this.containers=[this];var i,s,n,o,a,r,h,l,c=this.items,u=[[t.isFunction(this.options.items)?this.options.items.call(this.element[0],e,{item:this.currentItem}):t(this.options.items,this.element),this]],d=this._connectWith();if(d&&this.ready)for(i=d.length-1;i>=0;i--)for(n=t(d[i],this.document[0]),s=n.length-1;s>=0;s--)o=t.data(n[s],this.widgetFullName),o&&o!==this&&!o.options.disabled&&(u.push([t.isFunction(o.options.items)?o.options.items.call(o.element[0],e,{item:this.currentItem}):t(o.options.items,o.element),o]),this.containers.push(o));for(i=u.length-1;i>=0;i--)for(a=u[i][1],r=u[i][0],s=0,l=r.length;l>s;s++)h=t(r[s]),h.data(this.widgetName+"-item",a),c.push({item:h,instance:a,width:0,height:0,left:0,top:0})},refreshPositions:function(e){this.floating=this.items.length?"x"===this.options.axis||this._isFloating(this.items[0].item):!1,this.offsetParent&&this.helper&&(this.offset.parent=this._getParentOffset());var i,s,n,o;for(i=this.items.length-1;i>=0;i--)s=this.items[i],s.instance!==this.currentContainer&&this.currentContainer&&s.item[0]!==this.currentItem[0]||(n=this.options.toleranceElement?t(this.options.toleranceElement,s.item):s.item,e||(s.width=n.outerWidth(),s.height=n.outerHeight()),o=n.offset(),s.left=o.left,s.top=o.top);if(this.options.custom&&this.options.custom.refreshContainers)this.options.custom.refreshContainers.call(this);else for(i=this.containers.length-1;i>=0;i--)o=this.containers[i].element.offset(),this.containers[i].containerCache.left=o.left,this.containers[i].containerCache.top=o.top,this.containers[i].containerCache.width=this.containers[i].element.outerWidth(),this.containers[i].containerCache.height=this.containers[i].element.outerHeight();return this},_createPlaceholder:function(e){e=e||this;var i,s=e.options;s.placeholder&&s.placeholder.constructor!==String||(i=s.placeholder,s.placeholder={element:function(){var s=e.currentItem[0].nodeName.toLowerCase(),n=t("<"+s+">",e.document[0]);return e._addClass(n,"ui-sortable-placeholder",i||e.currentItem[0].className)._removeClass(n,"ui-sortable-helper"),"tbody"===s?e._createTrPlaceholder(e.currentItem.find("tr").eq(0),t("<tr>",e.document[0]).appendTo(n)):"tr"===s?e._createTrPlaceholder(e.currentItem,n):"img"===s&&n.attr("src",e.currentItem.attr("src")),i||n.css("visibility","hidden"),n},update:function(t,n){(!i||s.forcePlaceholderSize)&&(n.height()||n.height(e.currentItem.innerHeight()-parseInt(e.currentItem.css("paddingTop")||0,10)-parseInt(e.currentItem.css("paddingBottom")||0,10)),n.width()||n.width(e.currentItem.innerWidth()-parseInt(e.currentItem.css("paddingLeft")||0,10)-parseInt(e.currentItem.css("paddingRight")||0,10)))}}),e.placeholder=t(s.placeholder.element.call(e.element,e.currentItem)),e.currentItem.after(e.placeholder),s.placeholder.update(e,e.placeholder)},_createTrPlaceholder:function(e,i){var s=this;e.children().each(function(){t("<td>&#160;</td>",s.document[0]).attr("colspan",t(this).attr("colspan")||1).appendTo(i)})},_contactContainers:function(e){var i,s,n,o,a,r,h,l,c,u,d=null,p=null;for(i=this.containers.length-1;i>=0;i--)if(!t.contains(this.currentItem[0],this.containers[i].element[0]))if(this._intersectsWith(this.containers[i].containerCache)){if(d&&t.contains(this.containers[i].element[0],d.element[0]))continue;d=this.containers[i],p=i}else this.containers[i].containerCache.over&&(this.containers[i]._trigger("out",e,this._uiHash(this)),this.containers[i].containerCache.over=0);if(d)if(1===this.containers.length)this.containers[p].containerCache.over||(this.containers[p]._trigger("over",e,this._uiHash(this)),this.containers[p].containerCache.over=1);else{for(n=1e4,o=null,c=d.floating||this._isFloating(this.currentItem),a=c?"left":"top",r=c?"width":"height",u=c?"pageX":"pageY",s=this.items.length-1;s>=0;s--)t.contains(this.containers[p].element[0],this.items[s].item[0])&&this.items[s].item[0]!==this.currentItem[0]&&(h=this.items[s].item.offset()[a],l=!1,e[u]-h>this.items[s][r]/2&&(l=!0),n>Math.abs(e[u]-h)&&(n=Math.abs(e[u]-h),o=this.items[s],this.direction=l?"up":"down"));if(!o&&!this.options.dropOnEmpty)return;if(this.currentContainer===this.containers[p])return this.currentContainer.containerCache.over||(this.containers[p]._trigger("over",e,this._uiHash()),this.currentContainer.containerCache.over=1),void 0;o?this._rearrange(e,o,null,!0):this._rearrange(e,null,this.containers[p].element,!0),this._trigger("change",e,this._uiHash()),this.containers[p]._trigger("change",e,this._uiHash(this)),this.currentContainer=this.containers[p],this.options.placeholder.update(this.currentContainer,this.placeholder),this.containers[p]._trigger("over",e,this._uiHash(this)),this.containers[p].containerCache.over=1}},_createHelper:function(e){var i=this.options,s=t.isFunction(i.helper)?t(i.helper.apply(this.element[0],[e,this.currentItem])):"clone"===i.helper?this.currentItem.clone():this.currentItem;return s.parents("body").length||t("parent"!==i.appendTo?i.appendTo:this.currentItem[0].parentNode)[0].appendChild(s[0]),s[0]===this.currentItem[0]&&(this._storedCSS={width:this.currentItem[0].style.width,height:this.currentItem[0].style.height,position:this.currentItem.css("position"),top:this.currentItem.css("top"),left:this.currentItem.css("left")}),(!s[0].style.width||i.forceHelperSize)&&s.width(this.currentItem.width()),(!s[0].style.height||i.forceHelperSize)&&s.height(this.currentItem.height()),s},_adjustOffsetFromHelper:function(e){"string"==typeof e&&(e=e.split(" ")),t.isArray(e)&&(e={left:+e[0],top:+e[1]||0}),"left"in e&&(this.offset.click.left=e.left+this.margins.left),"right"in e&&(this.offset.click.left=this.helperProportions.width-e.right+this.margins.left),"top"in e&&(this.offset.click.top=e.top+this.margins.top),"bottom"in e&&(this.offset.click.top=this.helperProportions.height-e.bottom+this.margins.top)},_getParentOffset:function(){this.offsetParent=this.helper.offsetParent();var e=this.offsetParent.offset();return"absolute"===this.cssPosition&&this.scrollParent[0]!==this.document[0]&&t.contains(this.scrollParent[0],this.offsetParent[0])&&(e.left+=this.scrollParent.scrollLeft(),e.top+=this.scrollParent.scrollTop()),(this.offsetParent[0]===this.document[0].body||this.offsetParent[0].tagName&&"html"===this.offsetParent[0].tagName.toLowerCase()&&t.ui.ie)&&(e={top:0,left:0}),{top:e.top+(parseInt(this.offsetParent.css("borderTopWidth"),10)||0),left:e.left+(parseInt(this.offsetParent.css("borderLeftWidth"),10)||0)}},_getRelativeOffset:function(){if("relative"===this.cssPosition){var t=this.currentItem.position();return{top:t.top-(parseInt(this.helper.css("top"),10)||0)+this.scrollParent.scrollTop(),left:t.left-(parseInt(this.helper.css("left"),10)||0)+this.scrollParent.scrollLeft()}}return{top:0,left:0}},_cacheMargins:function(){this.margins={left:parseInt(this.currentItem.css("marginLeft"),10)||0,top:parseInt(this.currentItem.css("marginTop"),10)||0}},_cacheHelperProportions:function(){this.helperProportions={width:this.helper.outerWidth(),height:this.helper.outerHeight()}},_setContainment:function(){var e,i,s,n=this.options;"parent"===n.containment&&(n.containment=this.helper[0].parentNode),("document"===n.containment||"window"===n.containment)&&(this.containment=[0-this.offset.relative.left-this.offset.parent.left,0-this.offset.relative.top-this.offset.parent.top,"document"===n.containment?this.document.width():this.window.width()-this.helperProportions.width-this.margins.left,("document"===n.containment?this.document.height()||document.body.parentNode.scrollHeight:this.window.height()||this.document[0].body.parentNode.scrollHeight)-this.helperProportions.height-this.margins.top]),/^(document|window|parent)$/.test(n.containment)||(e=t(n.containment)[0],i=t(n.containment).offset(),s="hidden"!==t(e).css("overflow"),this.containment=[i.left+(parseInt(t(e).css("borderLeftWidth"),10)||0)+(parseInt(t(e).css("paddingLeft"),10)||0)-this.margins.left,i.top+(parseInt(t(e).css("borderTopWidth"),10)||0)+(parseInt(t(e).css("paddingTop"),10)||0)-this.margins.top,i.left+(s?Math.max(e.scrollWidth,e.offsetWidth):e.offsetWidth)-(parseInt(t(e).css("borderLeftWidth"),10)||0)-(parseInt(t(e).css("paddingRight"),10)||0)-this.helperProportions.width-this.margins.left,i.top+(s?Math.max(e.scrollHeight,e.offsetHeight):e.offsetHeight)-(parseInt(t(e).css("borderTopWidth"),10)||0)-(parseInt(t(e).css("paddingBottom"),10)||0)-this.helperProportions.height-this.margins.top])},_convertPositionTo:function(e,i){i||(i=this.position);var s="absolute"===e?1:-1,n="absolute"!==this.cssPosition||this.scrollParent[0]!==this.document[0]&&t.contains(this.scrollParent[0],this.offsetParent[0])?this.scrollParent:this.offsetParent,o=/(html|body)/i.test(n[0].tagName);return{top:i.top+this.offset.relative.top*s+this.offset.parent.top*s-("fixed"===this.cssPosition?-this.scrollParent.scrollTop():o?0:n.scrollTop())*s,left:i.left+this.offset.relative.left*s+this.offset.parent.left*s-("fixed"===this.cssPosition?-this.scrollParent.scrollLeft():o?0:n.scrollLeft())*s}},_generatePosition:function(e){var i,s,n=this.options,o=e.pageX,a=e.pageY,r="absolute"!==this.cssPosition||this.scrollParent[0]!==this.document[0]&&t.contains(this.scrollParent[0],this.offsetParent[0])?this.scrollParent:this.offsetParent,h=/(html|body)/i.test(r[0].tagName);return"relative"!==this.cssPosition||this.scrollParent[0]!==this.document[0]&&this.scrollParent[0]!==this.offsetParent[0]||(this.offset.relative=this._getRelativeOffset()),this.originalPosition&&(this.containment&&(e.pageX-this.offset.click.left<this.containment[0]&&(o=this.containment[0]+this.offset.click.left),e.pageY-this.offset.click.top<this.containment[1]&&(a=this.containment[1]+this.offset.click.top),e.pageX-this.offset.click.left>this.containment[2]&&(o=this.containment[2]+this.offset.click.left),e.pageY-this.offset.click.top>this.containment[3]&&(a=this.containment[3]+this.offset.click.top)),n.grid&&(i=this.originalPageY+Math.round((a-this.originalPageY)/n.grid[1])*n.grid[1],a=this.containment?i-this.offset.click.top>=this.containment[1]&&i-this.offset.click.top<=this.containment[3]?i:i-this.offset.click.top>=this.containment[1]?i-n.grid[1]:i+n.grid[1]:i,s=this.originalPageX+Math.round((o-this.originalPageX)/n.grid[0])*n.grid[0],o=this.containment?s-this.offset.click.left>=this.containment[0]&&s-this.offset.click.left<=this.containment[2]?s:s-this.offset.click.left>=this.containment[0]?s-n.grid[0]:s+n.grid[0]:s)),{top:a-this.offset.click.top-this.offset.relative.top-this.offset.parent.top+("fixed"===this.cssPosition?-this.scrollParent.scrollTop():h?0:r.scrollTop()),left:o-this.offset.click.left-this.offset.relative.left-this.offset.parent.left+("fixed"===this.cssPosition?-this.scrollParent.scrollLeft():h?0:r.scrollLeft())}},_rearrange:function(t,e,i,s){i?i[0].appendChild(this.placeholder[0]):e.item[0].parentNode.insertBefore(this.placeholder[0],"down"===this.direction?e.item[0]:e.item[0].nextSibling),this.counter=this.counter?++this.counter:1;var n=this.counter;
this._delay(function(){n===this.counter&&this.refreshPositions(!s)})},_clear:function(t,e){function i(t,e,i){return function(s){i._trigger(t,s,e._uiHash(e))}}this.reverting=!1;var s,n=[];if(!this._noFinalSort&&this.currentItem.parent().length&&this.placeholder.before(this.currentItem),this._noFinalSort=null,this.helper[0]===this.currentItem[0]){for(s in this._storedCSS)("auto"===this._storedCSS[s]||"static"===this._storedCSS[s])&&(this._storedCSS[s]="");this.currentItem.css(this._storedCSS),this._removeClass(this.currentItem,"ui-sortable-helper")}else this.currentItem.show();for(this.fromOutside&&!e&&n.push(function(t){this._trigger("receive",t,this._uiHash(this.fromOutside))}),!this.fromOutside&&this.domPosition.prev===this.currentItem.prev().not(".ui-sortable-helper")[0]&&this.domPosition.parent===this.currentItem.parent()[0]||e||n.push(function(t){this._trigger("update",t,this._uiHash())}),this!==this.currentContainer&&(e||(n.push(function(t){this._trigger("remove",t,this._uiHash())}),n.push(function(t){return function(e){t._trigger("receive",e,this._uiHash(this))}}.call(this,this.currentContainer)),n.push(function(t){return function(e){t._trigger("update",e,this._uiHash(this))}}.call(this,this.currentContainer)))),s=this.containers.length-1;s>=0;s--)e||n.push(i("deactivate",this,this.containers[s])),this.containers[s].containerCache.over&&(n.push(i("out",this,this.containers[s])),this.containers[s].containerCache.over=0);if(this.storedCursor&&(this.document.find("body").css("cursor",this.storedCursor),this.storedStylesheet.remove()),this._storedOpacity&&this.helper.css("opacity",this._storedOpacity),this._storedZIndex&&this.helper.css("zIndex","auto"===this._storedZIndex?"":this._storedZIndex),this.dragging=!1,e||this._trigger("beforeStop",t,this._uiHash()),this.placeholder[0].parentNode.removeChild(this.placeholder[0]),this.cancelHelperRemoval||(this.helper[0]!==this.currentItem[0]&&this.helper.remove(),this.helper=null),!e){for(s=0;n.length>s;s++)n[s].call(this,t);this._trigger("stop",t,this._uiHash())}return this.fromOutside=!1,!this.cancelHelperRemoval},_trigger:function(){t.Widget.prototype._trigger.apply(this,arguments)===!1&&this.cancel()},_uiHash:function(e){var i=e||this;return{helper:i.helper,placeholder:i.placeholder||t([]),position:i.position,originalPosition:i.originalPosition,offset:i.positionAbs,item:i.currentItem,sender:e?e.element:null}}}),t.widget("ui.spinner",{version:"1.12.1",defaultElement:"<input>",widgetEventPrefix:"spin",options:{classes:{"ui-spinner":"ui-corner-all","ui-spinner-down":"ui-corner-br","ui-spinner-up":"ui-corner-tr"},culture:null,icons:{down:"ui-icon-triangle-1-s",up:"ui-icon-triangle-1-n"},incremental:!0,max:null,min:null,numberFormat:null,page:10,step:1,change:null,spin:null,start:null,stop:null},_create:function(){this._setOption("max",this.options.max),this._setOption("min",this.options.min),this._setOption("step",this.options.step),""!==this.value()&&this._value(this.element.val(),!0),this._draw(),this._on(this._events),this._refresh(),this._on(this.window,{beforeunload:function(){this.element.removeAttr("autocomplete")}})},_getCreateOptions:function(){var e=this._super(),i=this.element;return t.each(["min","max","step"],function(t,s){var n=i.attr(s);null!=n&&n.length&&(e[s]=n)}),e},_events:{keydown:function(t){this._start(t)&&this._keydown(t)&&t.preventDefault()},keyup:"_stop",focus:function(){this.previous=this.element.val()},blur:function(t){return this.cancelBlur?(delete this.cancelBlur,void 0):(this._stop(),this._refresh(),this.previous!==this.element.val()&&this._trigger("change",t),void 0)},mousewheel:function(t,e){if(e){if(!this.spinning&&!this._start(t))return!1;this._spin((e>0?1:-1)*this.options.step,t),clearTimeout(this.mousewheelTimer),this.mousewheelTimer=this._delay(function(){this.spinning&&this._stop(t)},100),t.preventDefault()}},"mousedown .ui-spinner-button":function(e){function i(){var e=this.element[0]===t.ui.safeActiveElement(this.document[0]);e||(this.element.trigger("focus"),this.previous=s,this._delay(function(){this.previous=s}))}var s;s=this.element[0]===t.ui.safeActiveElement(this.document[0])?this.previous:this.element.val(),e.preventDefault(),i.call(this),this.cancelBlur=!0,this._delay(function(){delete this.cancelBlur,i.call(this)}),this._start(e)!==!1&&this._repeat(null,t(e.currentTarget).hasClass("ui-spinner-up")?1:-1,e)},"mouseup .ui-spinner-button":"_stop","mouseenter .ui-spinner-button":function(e){return t(e.currentTarget).hasClass("ui-state-active")?this._start(e)===!1?!1:(this._repeat(null,t(e.currentTarget).hasClass("ui-spinner-up")?1:-1,e),void 0):void 0},"mouseleave .ui-spinner-button":"_stop"},_enhance:function(){this.uiSpinner=this.element.attr("autocomplete","off").wrap("<span>").parent().append("<a></a><a></a>")},_draw:function(){this._enhance(),this._addClass(this.uiSpinner,"ui-spinner","ui-widget ui-widget-content"),this._addClass("ui-spinner-input"),this.element.attr("role","spinbutton"),this.buttons=this.uiSpinner.children("a").attr("tabIndex",-1).attr("aria-hidden",!0).button({classes:{"ui-button":""}}),this._removeClass(this.buttons,"ui-corner-all"),this._addClass(this.buttons.first(),"ui-spinner-button ui-spinner-up"),this._addClass(this.buttons.last(),"ui-spinner-button ui-spinner-down"),this.buttons.first().button({icon:this.options.icons.up,showLabel:!1}),this.buttons.last().button({icon:this.options.icons.down,showLabel:!1}),this.buttons.height()>Math.ceil(.5*this.uiSpinner.height())&&this.uiSpinner.height()>0&&this.uiSpinner.height(this.uiSpinner.height())},_keydown:function(e){var i=this.options,s=t.ui.keyCode;switch(e.keyCode){case s.UP:return this._repeat(null,1,e),!0;case s.DOWN:return this._repeat(null,-1,e),!0;case s.PAGE_UP:return this._repeat(null,i.page,e),!0;case s.PAGE_DOWN:return this._repeat(null,-i.page,e),!0}return!1},_start:function(t){return this.spinning||this._trigger("start",t)!==!1?(this.counter||(this.counter=1),this.spinning=!0,!0):!1},_repeat:function(t,e,i){t=t||500,clearTimeout(this.timer),this.timer=this._delay(function(){this._repeat(40,e,i)},t),this._spin(e*this.options.step,i)},_spin:function(t,e){var i=this.value()||0;this.counter||(this.counter=1),i=this._adjustValue(i+t*this._increment(this.counter)),this.spinning&&this._trigger("spin",e,{value:i})===!1||(this._value(i),this.counter++)},_increment:function(e){var i=this.options.incremental;return i?t.isFunction(i)?i(e):Math.floor(e*e*e/5e4-e*e/500+17*e/200+1):1},_precision:function(){var t=this._precisionOf(this.options.step);return null!==this.options.min&&(t=Math.max(t,this._precisionOf(this.options.min))),t},_precisionOf:function(t){var e=""+t,i=e.indexOf(".");return-1===i?0:e.length-i-1},_adjustValue:function(t){var e,i,s=this.options;return e=null!==s.min?s.min:0,i=t-e,i=Math.round(i/s.step)*s.step,t=e+i,t=parseFloat(t.toFixed(this._precision())),null!==s.max&&t>s.max?s.max:null!==s.min&&s.min>t?s.min:t},_stop:function(t){this.spinning&&(clearTimeout(this.timer),clearTimeout(this.mousewheelTimer),this.counter=0,this.spinning=!1,this._trigger("stop",t))},_setOption:function(t,e){var i,s,n;return"culture"===t||"numberFormat"===t?(i=this._parse(this.element.val()),this.options[t]=e,this.element.val(this._format(i)),void 0):(("max"===t||"min"===t||"step"===t)&&"string"==typeof e&&(e=this._parse(e)),"icons"===t&&(s=this.buttons.first().find(".ui-icon"),this._removeClass(s,null,this.options.icons.up),this._addClass(s,null,e.up),n=this.buttons.last().find(".ui-icon"),this._removeClass(n,null,this.options.icons.down),this._addClass(n,null,e.down)),this._super(t,e),void 0)},_setOptionDisabled:function(t){this._super(t),this._toggleClass(this.uiSpinner,null,"ui-state-disabled",!!t),this.element.prop("disabled",!!t),this.buttons.button(t?"disable":"enable")},_setOptions:r(function(t){this._super(t)}),_parse:function(t){return"string"==typeof t&&""!==t&&(t=window.Globalize&&this.options.numberFormat?Globalize.parseFloat(t,10,this.options.culture):+t),""===t||isNaN(t)?null:t},_format:function(t){return""===t?"":window.Globalize&&this.options.numberFormat?Globalize.format(t,this.options.numberFormat,this.options.culture):t},_refresh:function(){this.element.attr({"aria-valuemin":this.options.min,"aria-valuemax":this.options.max,"aria-valuenow":this._parse(this.element.val())})},isValid:function(){var t=this.value();return null===t?!1:t===this._adjustValue(t)},_value:function(t,e){var i;""!==t&&(i=this._parse(t),null!==i&&(e||(i=this._adjustValue(i)),t=this._format(i))),this.element.val(t),this._refresh()},_destroy:function(){this.element.prop("disabled",!1).removeAttr("autocomplete role aria-valuemin aria-valuemax aria-valuenow"),this.uiSpinner.replaceWith(this.element)},stepUp:r(function(t){this._stepUp(t)}),_stepUp:function(t){this._start()&&(this._spin((t||1)*this.options.step),this._stop())},stepDown:r(function(t){this._stepDown(t)}),_stepDown:function(t){this._start()&&(this._spin((t||1)*-this.options.step),this._stop())},pageUp:r(function(t){this._stepUp((t||1)*this.options.page)}),pageDown:r(function(t){this._stepDown((t||1)*this.options.page)}),value:function(t){return arguments.length?(r(this._value).call(this,t),void 0):this._parse(this.element.val())},widget:function(){return this.uiSpinner}}),t.uiBackCompat!==!1&&t.widget("ui.spinner",t.ui.spinner,{_enhance:function(){this.uiSpinner=this.element.attr("autocomplete","off").wrap(this._uiSpinnerHtml()).parent().append(this._buttonHtml())},_uiSpinnerHtml:function(){return"<span>"},_buttonHtml:function(){return"<a></a><a></a>"}}),t.ui.spinner,t.widget("ui.tabs",{version:"1.12.1",delay:300,options:{active:null,classes:{"ui-tabs":"ui-corner-all","ui-tabs-nav":"ui-corner-all","ui-tabs-panel":"ui-corner-bottom","ui-tabs-tab":"ui-corner-top"},collapsible:!1,event:"click",heightStyle:"content",hide:null,show:null,activate:null,beforeActivate:null,beforeLoad:null,load:null},_isLocal:function(){var t=/#.*$/;return function(e){var i,s;i=e.href.replace(t,""),s=location.href.replace(t,"");try{i=decodeURIComponent(i)}catch(n){}try{s=decodeURIComponent(s)}catch(n){}return e.hash.length>1&&i===s}}(),_create:function(){var e=this,i=this.options;this.running=!1,this._addClass("ui-tabs","ui-widget ui-widget-content"),this._toggleClass("ui-tabs-collapsible",null,i.collapsible),this._processTabs(),i.active=this._initialActive(),t.isArray(i.disabled)&&(i.disabled=t.unique(i.disabled.concat(t.map(this.tabs.filter(".ui-state-disabled"),function(t){return e.tabs.index(t)}))).sort()),this.active=this.options.active!==!1&&this.anchors.length?this._findActive(i.active):t(),this._refresh(),this.active.length&&this.load(i.active)},_initialActive:function(){var e=this.options.active,i=this.options.collapsible,s=location.hash.substring(1);return null===e&&(s&&this.tabs.each(function(i,n){return t(n).attr("aria-controls")===s?(e=i,!1):void 0}),null===e&&(e=this.tabs.index(this.tabs.filter(".ui-tabs-active"))),(null===e||-1===e)&&(e=this.tabs.length?0:!1)),e!==!1&&(e=this.tabs.index(this.tabs.eq(e)),-1===e&&(e=i?!1:0)),!i&&e===!1&&this.anchors.length&&(e=0),e},_getCreateEventData:function(){return{tab:this.active,panel:this.active.length?this._getPanelForTab(this.active):t()}},_tabKeydown:function(e){var i=t(t.ui.safeActiveElement(this.document[0])).closest("li"),s=this.tabs.index(i),n=!0;if(!this._handlePageNav(e)){switch(e.keyCode){case t.ui.keyCode.RIGHT:case t.ui.keyCode.DOWN:s++;break;case t.ui.keyCode.UP:case t.ui.keyCode.LEFT:n=!1,s--;break;case t.ui.keyCode.END:s=this.anchors.length-1;break;case t.ui.keyCode.HOME:s=0;break;case t.ui.keyCode.SPACE:return e.preventDefault(),clearTimeout(this.activating),this._activate(s),void 0;case t.ui.keyCode.ENTER:return e.preventDefault(),clearTimeout(this.activating),this._activate(s===this.options.active?!1:s),void 0;default:return}e.preventDefault(),clearTimeout(this.activating),s=this._focusNextTab(s,n),e.ctrlKey||e.metaKey||(i.attr("aria-selected","false"),this.tabs.eq(s).attr("aria-selected","true"),this.activating=this._delay(function(){this.option("active",s)},this.delay))}},_panelKeydown:function(e){this._handlePageNav(e)||e.ctrlKey&&e.keyCode===t.ui.keyCode.UP&&(e.preventDefault(),this.active.trigger("focus"))},_handlePageNav:function(e){return e.altKey&&e.keyCode===t.ui.keyCode.PAGE_UP?(this._activate(this._focusNextTab(this.options.active-1,!1)),!0):e.altKey&&e.keyCode===t.ui.keyCode.PAGE_DOWN?(this._activate(this._focusNextTab(this.options.active+1,!0)),!0):void 0},_findNextTab:function(e,i){function s(){return e>n&&(e=0),0>e&&(e=n),e}for(var n=this.tabs.length-1;-1!==t.inArray(s(),this.options.disabled);)e=i?e+1:e-1;return e},_focusNextTab:function(t,e){return t=this._findNextTab(t,e),this.tabs.eq(t).trigger("focus"),t},_setOption:function(t,e){return"active"===t?(this._activate(e),void 0):(this._super(t,e),"collapsible"===t&&(this._toggleClass("ui-tabs-collapsible",null,e),e||this.options.active!==!1||this._activate(0)),"event"===t&&this._setupEvents(e),"heightStyle"===t&&this._setupHeightStyle(e),void 0)},_sanitizeSelector:function(t){return t?t.replace(/[!"$%&'()*+,.\/:;<=>?@\[\]\^`{|}~]/g,"\\$&"):""},refresh:function(){var e=this.options,i=this.tablist.children(":has(a[href])");e.disabled=t.map(i.filter(".ui-state-disabled"),function(t){return i.index(t)}),this._processTabs(),e.active!==!1&&this.anchors.length?this.active.length&&!t.contains(this.tablist[0],this.active[0])?this.tabs.length===e.disabled.length?(e.active=!1,this.active=t()):this._activate(this._findNextTab(Math.max(0,e.active-1),!1)):e.active=this.tabs.index(this.active):(e.active=!1,this.active=t()),this._refresh()},_refresh:function(){this._setOptionDisabled(this.options.disabled),this._setupEvents(this.options.event),this._setupHeightStyle(this.options.heightStyle),this.tabs.not(this.active).attr({"aria-selected":"false","aria-expanded":"false",tabIndex:-1}),this.panels.not(this._getPanelForTab(this.active)).hide().attr({"aria-hidden":"true"}),this.active.length?(this.active.attr({"aria-selected":"true","aria-expanded":"true",tabIndex:0}),this._addClass(this.active,"ui-tabs-active","ui-state-active"),this._getPanelForTab(this.active).show().attr({"aria-hidden":"false"})):this.tabs.eq(0).attr("tabIndex",0)},_processTabs:function(){var e=this,i=this.tabs,s=this.anchors,n=this.panels;this.tablist=this._getList().attr("role","tablist"),this._addClass(this.tablist,"ui-tabs-nav","ui-helper-reset ui-helper-clearfix ui-widget-header"),this.tablist.on("mousedown"+this.eventNamespace,"> li",function(e){t(this).is(".ui-state-disabled")&&e.preventDefault()}).on("focus"+this.eventNamespace,".ui-tabs-anchor",function(){t(this).closest("li").is(".ui-state-disabled")&&this.blur()}),this.tabs=this.tablist.find("> li:has(a[href])").attr({role:"tab",tabIndex:-1}),this._addClass(this.tabs,"ui-tabs-tab","ui-state-default"),this.anchors=this.tabs.map(function(){return t("a",this)[0]}).attr({role:"presentation",tabIndex:-1}),this._addClass(this.anchors,"ui-tabs-anchor"),this.panels=t(),this.anchors.each(function(i,s){var n,o,a,r=t(s).uniqueId().attr("id"),h=t(s).closest("li"),l=h.attr("aria-controls");e._isLocal(s)?(n=s.hash,a=n.substring(1),o=e.element.find(e._sanitizeSelector(n))):(a=h.attr("aria-controls")||t({}).uniqueId()[0].id,n="#"+a,o=e.element.find(n),o.length||(o=e._createPanel(a),o.insertAfter(e.panels[i-1]||e.tablist)),o.attr("aria-live","polite")),o.length&&(e.panels=e.panels.add(o)),l&&h.data("ui-tabs-aria-controls",l),h.attr({"aria-controls":a,"aria-labelledby":r}),o.attr("aria-labelledby",r)}),this.panels.attr("role","tabpanel"),this._addClass(this.panels,"ui-tabs-panel","ui-widget-content"),i&&(this._off(i.not(this.tabs)),this._off(s.not(this.anchors)),this._off(n.not(this.panels)))},_getList:function(){return this.tablist||this.element.find("ol, ul").eq(0)},_createPanel:function(e){return t("<div>").attr("id",e).data("ui-tabs-destroy",!0)},_setOptionDisabled:function(e){var i,s,n;for(t.isArray(e)&&(e.length?e.length===this.anchors.length&&(e=!0):e=!1),n=0;s=this.tabs[n];n++)i=t(s),e===!0||-1!==t.inArray(n,e)?(i.attr("aria-disabled","true"),this._addClass(i,null,"ui-state-disabled")):(i.removeAttr("aria-disabled"),this._removeClass(i,null,"ui-state-disabled"));this.options.disabled=e,this._toggleClass(this.widget(),this.widgetFullName+"-disabled",null,e===!0)},_setupEvents:function(e){var i={};e&&t.each(e.split(" "),function(t,e){i[e]="_eventHandler"}),this._off(this.anchors.add(this.tabs).add(this.panels)),this._on(!0,this.anchors,{click:function(t){t.preventDefault()}}),this._on(this.anchors,i),this._on(this.tabs,{keydown:"_tabKeydown"}),this._on(this.panels,{keydown:"_panelKeydown"}),this._focusable(this.tabs),this._hoverable(this.tabs)},_setupHeightStyle:function(e){var i,s=this.element.parent();"fill"===e?(i=s.height(),i-=this.element.outerHeight()-this.element.height(),this.element.siblings(":visible").each(function(){var e=t(this),s=e.css("position");"absolute"!==s&&"fixed"!==s&&(i-=e.outerHeight(!0))}),this.element.children().not(this.panels).each(function(){i-=t(this).outerHeight(!0)}),this.panels.each(function(){t(this).height(Math.max(0,i-t(this).innerHeight()+t(this).height()))}).css("overflow","auto")):"auto"===e&&(i=0,this.panels.each(function(){i=Math.max(i,t(this).height("").height())}).height(i))},_eventHandler:function(e){var i=this.options,s=this.active,n=t(e.currentTarget),o=n.closest("li"),a=o[0]===s[0],r=a&&i.collapsible,h=r?t():this._getPanelForTab(o),l=s.length?this._getPanelForTab(s):t(),c={oldTab:s,oldPanel:l,newTab:r?t():o,newPanel:h};e.preventDefault(),o.hasClass("ui-state-disabled")||o.hasClass("ui-tabs-loading")||this.running||a&&!i.collapsible||this._trigger("beforeActivate",e,c)===!1||(i.active=r?!1:this.tabs.index(o),this.active=a?t():o,this.xhr&&this.xhr.abort(),l.length||h.length||t.error("jQuery UI Tabs: Mismatching fragment identifier."),h.length&&this.load(this.tabs.index(o),e),this._toggle(e,c))},_toggle:function(e,i){function s(){o.running=!1,o._trigger("activate",e,i)}function n(){o._addClass(i.newTab.closest("li"),"ui-tabs-active","ui-state-active"),a.length&&o.options.show?o._show(a,o.options.show,s):(a.show(),s())}var o=this,a=i.newPanel,r=i.oldPanel;this.running=!0,r.length&&this.options.hide?this._hide(r,this.options.hide,function(){o._removeClass(i.oldTab.closest("li"),"ui-tabs-active","ui-state-active"),n()}):(this._removeClass(i.oldTab.closest("li"),"ui-tabs-active","ui-state-active"),r.hide(),n()),r.attr("aria-hidden","true"),i.oldTab.attr({"aria-selected":"false","aria-expanded":"false"}),a.length&&r.length?i.oldTab.attr("tabIndex",-1):a.length&&this.tabs.filter(function(){return 0===t(this).attr("tabIndex")}).attr("tabIndex",-1),a.attr("aria-hidden","false"),i.newTab.attr({"aria-selected":"true","aria-expanded":"true",tabIndex:0})},_activate:function(e){var i,s=this._findActive(e);s[0]!==this.active[0]&&(s.length||(s=this.active),i=s.find(".ui-tabs-anchor")[0],this._eventHandler({target:i,currentTarget:i,preventDefault:t.noop}))},_findActive:function(e){return e===!1?t():this.tabs.eq(e)},_getIndex:function(e){return"string"==typeof e&&(e=this.anchors.index(this.anchors.filter("[href$='"+t.ui.escapeSelector(e)+"']"))),e},_destroy:function(){this.xhr&&this.xhr.abort(),this.tablist.removeAttr("role").off(this.eventNamespace),this.anchors.removeAttr("role tabIndex").removeUniqueId(),this.tabs.add(this.panels).each(function(){t.data(this,"ui-tabs-destroy")?t(this).remove():t(this).removeAttr("role tabIndex aria-live aria-busy aria-selected aria-labelledby aria-hidden aria-expanded")}),this.tabs.each(function(){var e=t(this),i=e.data("ui-tabs-aria-controls");i?e.attr("aria-controls",i).removeData("ui-tabs-aria-controls"):e.removeAttr("aria-controls")}),this.panels.show(),"content"!==this.options.heightStyle&&this.panels.css("height","")},enable:function(e){var i=this.options.disabled;i!==!1&&(void 0===e?i=!1:(e=this._getIndex(e),i=t.isArray(i)?t.map(i,function(t){return t!==e?t:null}):t.map(this.tabs,function(t,i){return i!==e?i:null})),this._setOptionDisabled(i))},disable:function(e){var i=this.options.disabled;if(i!==!0){if(void 0===e)i=!0;else{if(e=this._getIndex(e),-1!==t.inArray(e,i))return;i=t.isArray(i)?t.merge([e],i).sort():[e]}this._setOptionDisabled(i)}},load:function(e,i){e=this._getIndex(e);var s=this,n=this.tabs.eq(e),o=n.find(".ui-tabs-anchor"),a=this._getPanelForTab(n),r={tab:n,panel:a},h=function(t,e){"abort"===e&&s.panels.stop(!1,!0),s._removeClass(n,"ui-tabs-loading"),a.removeAttr("aria-busy"),t===s.xhr&&delete s.xhr};this._isLocal(o[0])||(this.xhr=t.ajax(this._ajaxSettings(o,i,r)),this.xhr&&"canceled"!==this.xhr.statusText&&(this._addClass(n,"ui-tabs-loading"),a.attr("aria-busy","true"),this.xhr.done(function(t,e,n){setTimeout(function(){a.html(t),s._trigger("load",i,r),h(n,e)},1)}).fail(function(t,e){setTimeout(function(){h(t,e)},1)})))},_ajaxSettings:function(e,i,s){var n=this;return{url:e.attr("href").replace(/#.*$/,""),beforeSend:function(e,o){return n._trigger("beforeLoad",i,t.extend({jqXHR:e,ajaxSettings:o},s))}}},_getPanelForTab:function(e){var i=t(e).attr("aria-controls");return this.element.find(this._sanitizeSelector("#"+i))}}),t.uiBackCompat!==!1&&t.widget("ui.tabs",t.ui.tabs,{_processTabs:function(){this._superApply(arguments),this._addClass(this.tabs,"ui-tab")}}),t.ui.tabs,t.widget("ui.tooltip",{version:"1.12.1",options:{classes:{"ui-tooltip":"ui-corner-all ui-widget-shadow"},content:function(){var e=t(this).attr("title")||"";return t("<a>").text(e).html()},hide:!0,items:"[title]:not([disabled])",position:{my:"left top+15",at:"left bottom",collision:"flipfit flip"},show:!0,track:!1,close:null,open:null},_addDescribedBy:function(e,i){var s=(e.attr("aria-describedby")||"").split(/\s+/);s.push(i),e.data("ui-tooltip-id",i).attr("aria-describedby",t.trim(s.join(" ")))},_removeDescribedBy:function(e){var i=e.data("ui-tooltip-id"),s=(e.attr("aria-describedby")||"").split(/\s+/),n=t.inArray(i,s);-1!==n&&s.splice(n,1),e.removeData("ui-tooltip-id"),s=t.trim(s.join(" ")),s?e.attr("aria-describedby",s):e.removeAttr("aria-describedby")},_create:function(){this._on({mouseover:"open",focusin:"open"}),this.tooltips={},this.parents={},this.liveRegion=t("<div>").attr({role:"log","aria-live":"assertive","aria-relevant":"additions"}).appendTo(this.document[0].body),this._addClass(this.liveRegion,null,"ui-helper-hidden-accessible"),this.disabledTitles=t([])},_setOption:function(e,i){var s=this;this._super(e,i),"content"===e&&t.each(this.tooltips,function(t,e){s._updateContent(e.element)})},_setOptionDisabled:function(t){this[t?"_disable":"_enable"]()},_disable:function(){var e=this;t.each(this.tooltips,function(i,s){var n=t.Event("blur");n.target=n.currentTarget=s.element[0],e.close(n,!0)}),this.disabledTitles=this.disabledTitles.add(this.element.find(this.options.items).addBack().filter(function(){var e=t(this);return e.is("[title]")?e.data("ui-tooltip-title",e.attr("title")).removeAttr("title"):void 0}))},_enable:function(){this.disabledTitles.each(function(){var e=t(this);e.data("ui-tooltip-title")&&e.attr("title",e.data("ui-tooltip-title"))}),this.disabledTitles=t([])},open:function(e){var i=this,s=t(e?e.target:this.element).closest(this.options.items);s.length&&!s.data("ui-tooltip-id")&&(s.attr("title")&&s.data("ui-tooltip-title",s.attr("title")),s.data("ui-tooltip-open",!0),e&&"mouseover"===e.type&&s.parents().each(function(){var e,s=t(this);s.data("ui-tooltip-open")&&(e=t.Event("blur"),e.target=e.currentTarget=this,i.close(e,!0)),s.attr("title")&&(s.uniqueId(),i.parents[this.id]={element:this,title:s.attr("title")},s.attr("title",""))}),this._registerCloseHandlers(e,s),this._updateContent(s,e))},_updateContent:function(t,e){var i,s=this.options.content,n=this,o=e?e.type:null;return"string"==typeof s||s.nodeType||s.jquery?this._open(e,t,s):(i=s.call(t[0],function(i){n._delay(function(){t.data("ui-tooltip-open")&&(e&&(e.type=o),this._open(e,t,i))})}),i&&this._open(e,t,i),void 0)},_open:function(e,i,s){function n(t){l.of=t,a.is(":hidden")||a.position(l)}var o,a,r,h,l=t.extend({},this.options.position);if(s){if(o=this._find(i))return o.tooltip.find(".ui-tooltip-content").html(s),void 0;i.is("[title]")&&(e&&"mouseover"===e.type?i.attr("title",""):i.removeAttr("title")),o=this._tooltip(i),a=o.tooltip,this._addDescribedBy(i,a.attr("id")),a.find(".ui-tooltip-content").html(s),this.liveRegion.children().hide(),h=t("<div>").html(a.find(".ui-tooltip-content").html()),h.removeAttr("name").find("[name]").removeAttr("name"),h.removeAttr("id").find("[id]").removeAttr("id"),h.appendTo(this.liveRegion),this.options.track&&e&&/^mouse/.test(e.type)?(this._on(this.document,{mousemove:n}),n(e)):a.position(t.extend({of:i},this.options.position)),a.hide(),this._show(a,this.options.show),this.options.track&&this.options.show&&this.options.show.delay&&(r=this.delayedShow=setInterval(function(){a.is(":visible")&&(n(l.of),clearInterval(r))},t.fx.interval)),this._trigger("open",e,{tooltip:a})}},_registerCloseHandlers:function(e,i){var s={keyup:function(e){if(e.keyCode===t.ui.keyCode.ESCAPE){var s=t.Event(e);s.currentTarget=i[0],this.close(s,!0)}}};i[0]!==this.element[0]&&(s.remove=function(){this._removeTooltip(this._find(i).tooltip)}),e&&"mouseover"!==e.type||(s.mouseleave="close"),e&&"focusin"!==e.type||(s.focusout="close"),this._on(!0,i,s)},close:function(e){var i,s=this,n=t(e?e.currentTarget:this.element),o=this._find(n);return o?(i=o.tooltip,o.closing||(clearInterval(this.delayedShow),n.data("ui-tooltip-title")&&!n.attr("title")&&n.attr("title",n.data("ui-tooltip-title")),this._removeDescribedBy(n),o.hiding=!0,i.stop(!0),this._hide(i,this.options.hide,function(){s._removeTooltip(t(this))}),n.removeData("ui-tooltip-open"),this._off(n,"mouseleave focusout keyup"),n[0]!==this.element[0]&&this._off(n,"remove"),this._off(this.document,"mousemove"),e&&"mouseleave"===e.type&&t.each(this.parents,function(e,i){t(i.element).attr("title",i.title),delete s.parents[e]}),o.closing=!0,this._trigger("close",e,{tooltip:i}),o.hiding||(o.closing=!1)),void 0):(n.removeData("ui-tooltip-open"),void 0)},_tooltip:function(e){var i=t("<div>").attr("role","tooltip"),s=t("<div>").appendTo(i),n=i.uniqueId().attr("id");return this._addClass(s,"ui-tooltip-content"),this._addClass(i,"ui-tooltip","ui-widget ui-widget-content"),i.appendTo(this._appendTo(e)),this.tooltips[n]={element:e,tooltip:i}},_find:function(t){var e=t.data("ui-tooltip-id");return e?this.tooltips[e]:null},_removeTooltip:function(t){t.remove(),delete this.tooltips[t.attr("id")]},_appendTo:function(t){var e=t.closest(".ui-front, dialog");return e.length||(e=this.document[0].body),e},_destroy:function(){var e=this;t.each(this.tooltips,function(i,s){var n=t.Event("blur"),o=s.element;n.target=n.currentTarget=o[0],e.close(n,!0),t("#"+i).remove(),o.data("ui-tooltip-title")&&(o.attr("title")||o.attr("title",o.data("ui-tooltip-title")),o.removeData("ui-tooltip-title"))}),this.liveRegion.remove()}}),t.uiBackCompat!==!1&&t.widget("ui.tooltip",t.ui.tooltip,{options:{tooltipClass:null},_tooltip:function(){var t=this._superApply(arguments);return this.options.tooltipClass&&t.tooltip.addClass(this.options.tooltipClass),t}}),t.ui.tooltip});
```
