## Supplemental Data Set 1 for "The *pos-1* 3ʹ untranslated region governs germline specification and proliferation to ensure reproductive robustness": jquery.tocify.min.js

```
/*! jquery.tocify - v1.9.0 - 2013-10-01 
* http://gregfranko.com/jquery.tocify.js/
* Copyright (c) 2013 Greg Franko; Licensed MIT*/
(function(e){"use strict";e(window.jQuery,window,document)})(function(e,t,s){"use strict";var i="tocify",o="tocify-focus",n="tocify-hover",a="tocify-hide",l="tocify-header",h="."+l,r="tocify-subheader",d="."+r,c="tocify-item",f="."+c,u="tocify-extend-page",p="."+u;e.widget("toc.tocify",{version:"1.9.0",options:{context:"body",ignoreSelector:null,selectors:"h1, h2, h3",showAndHide:!0,showEffect:"slideDown",showEffectSpeed:"medium",hideEffect:"slideUp",hideEffectSpeed:"medium",smoothScroll:!0,smoothScrollSpeed:"medium",scrollTo:0,showAndHideOnScroll:!0,highlightOnScroll:!0,highlightOffset:40,theme:"bootstrap",extendPage:!0,extendPageOffset:100,history:!0,scrollHistory:!1,hashGenerator:"compact",highlightDefault:!0},_create:function(){var s=this;s.extendPageScroll=!0,s.items=[],s._generateToc(),s._addCSSClasses(),s.webkit=function(){for(var e in t)if(e&&-1!==e.toLowerCase().indexOf("webkit"))return!0;return!1}(),s._setEventHandlers(),e(t).load(function(){s._setActiveElement(!0),e("html, body").promise().done(function(){setTimeout(function(){s.extendPageScroll=!1},0)})})},_generateToc:function(){var t,s,o=this,n=o.options.ignoreSelector;return t=-1!==this.options.selectors.indexOf(",")?e(this.options.context).find(this.options.selectors.replace(/ /g,"").substr(0,this.options.selectors.indexOf(","))):e(this.options.context).find(this.options.selectors.replace(/ /g,"")),t.length?(o.element.addClass(i),t.each(function(t){e(this).is(n)||(s=e("<ul/>",{id:l+t,"class":l}).append(o._nestElements(e(this),t)),o.element.append(s),e(this).nextUntil(this.nodeName.toLowerCase()).each(function(){0===e(this).find(o.options.selectors).length?e(this).filter(o.options.selectors).each(function(){e(this).is(n)||o._appendSubheaders.call(this,o,s)}):e(this).find(o.options.selectors).each(function(){e(this).is(n)||o._appendSubheaders.call(this,o,s)})}))}),undefined):(o.element.addClass(a),undefined)},_setActiveElement:function(e){var s=this,i=t.location.hash.substring(1),o=s.element.find('li[data-unique="'+i+'"]');return i.length?(s.element.find("."+s.focusClass).removeClass(s.focusClass),o.addClass(s.focusClass),s.options.showAndHide&&o.click()):(s.element.find("."+s.focusClass).removeClass(s.focusClass),!i.length&&e&&s.options.highlightDefault&&s.element.find(f).first().addClass(s.focusClass)),s},_nestElements:function(t,s){var i,o,n;return i=e.grep(this.items,function(e){return e===t.text()}),i.length?this.items.push(t.text()+s):this.items.push(t.text()),n=this._generateHashValue(i,t,s),o=e("<li/>",{"class":c,"data-unique":n}).append(e("<a/>",{text:t.text()})),t.before(e("<div/>",{name:n,"data-unique":n})),o},_generateHashValue:function(e,t,s){var i="",o=this.options.hashGenerator;if("pretty"===o){for(i=t.text().toLowerCase().replace(/\s/g,"-");i.indexOf("--")>-1;)i=i.replace(/--/g,"-");for(;i.indexOf(":-")>-1;)i=i.replace(/:-/g,"-")}else i="function"==typeof o?o(t.text(),t):t.text().replace(/\s/g,"");return e.length&&(i+=""+s),i},_appendSubheaders:function(t,s){var i=e(this).index(t.options.selectors),o=e(t.options.selectors).eq(i-1),n=+e(this).prop("tagName").charAt(1),a=+o.prop("tagName").charAt(1);a>n?t.element.find(d+"[data-tag="+n+"]").last().append(t._nestElements(e(this),i)):n===a?s.find(f).last().after(t._nestElements(e(this),i)):s.find(f).last().after(e("<ul/>",{"class":r,"data-tag":n})).next(d).append(t._nestElements(e(this),i))},_setEventHandlers:function(){var i=this;this.element.on("click.tocify","li",function(){if(i.options.history&&(t.location.hash=e(this).attr("data-unique")),i.element.find("."+i.focusClass).removeClass(i.focusClass),e(this).addClass(i.focusClass),i.options.showAndHide){var s=e('li[data-unique="'+e(this).attr("data-unique")+'"]');i._triggerShow(s)}i._scrollTo(e(this))}),this.element.find("li").on({"mouseenter.tocify":function(){e(this).addClass(i.hoverClass),e(this).css("cursor","pointer")},"mouseleave.tocify":function(){"bootstrap"!==i.options.theme&&e(this).removeClass(i.hoverClass)}}),(i.options.extendPage||i.options.highlightOnScroll||i.options.scrollHistory||i.options.showAndHideOnScroll)&&e(t).on("scroll.tocify",function(){e("html, body").promise().done(function(){var o,n,a,l,h=e(t).scrollTop(),r=e(t).height(),d=e(s).height(),c=e("body")[0].scrollHeight;if(i.options.extendPage&&(i.webkit&&h>=c-r-i.options.extendPageOffset||!i.webkit&&r+h>d-i.options.extendPageOffset)&&!e(p).length){if(n=e('div[data-unique="'+e(f).last().attr("data-unique")+'"]'),!n.length)return;a=n.offset().top,e(i.options.context).append(e("<div />",{"class":u,height:Math.abs(a-h)+"px","data-unique":u})),i.extendPageScroll&&(l=i.element.find("li.active"),i._scrollTo(e('div[data-unique="'+l.attr("data-unique")+'"]')))}setTimeout(function(){var s,n=null,a=null,l=e(i.options.context).find("div[data-unique]");l.each(function(t){var s=Math.abs((e(this).next().length?e(this).next():e(this)).offset().top-h-i.options.highlightOffset);return null==n||n>s?(n=s,a=t,undefined):!1}),s=e(l[a]).attr("data-unique"),o=e('li[data-unique="'+s+'"]'),i.options.highlightOnScroll&&o.length&&(i.element.find("."+i.focusClass).removeClass(i.focusClass),o.addClass(i.focusClass)),i.options.scrollHistory&&t.location.hash!=="#"+s&&t.location.replace("#"+s),i.options.showAndHideOnScroll&&i.options.showAndHide&&i._triggerShow(o,!0)},0)})})},show:function(t){var s=this;if(!t.is(":visible"))switch(t.find(d).length||t.parent().is(h)||t.parent().is(":visible")?t.children(d).length||t.parent().is(h)||(t=t.closest(d)):t=t.parents(d).add(t),s.options.showEffect){case"none":t.show();break;case"show":t.show(s.options.showEffectSpeed);break;case"slideDown":t.slideDown(s.options.showEffectSpeed);break;case"fadeIn":t.fadeIn(s.options.showEffectSpeed);break;default:t.show()}return t.parent().is(h)?s.hide(e(d).not(t)):s.hide(e(d).not(t.closest(h).find(d).not(t.siblings()))),s},hide:function(e){var t=this;switch(t.options.hideEffect){case"none":e.hide();break;case"hide":e.hide(t.options.hideEffectSpeed);break;case"slideUp":e.slideUp(t.options.hideEffectSpeed);break;case"fadeOut":e.fadeOut(t.options.hideEffectSpeed);break;default:e.hide()}return t},_triggerShow:function(e,t){var s=this;return e.parent().is(h)||e.next().is(d)?s.show(e.next(d),t):e.parent().is(d)&&s.show(e.parent(),t),s},_addCSSClasses:function(){return"jqueryui"===this.options.theme?(this.focusClass="ui-state-default",this.hoverClass="ui-state-hover",this.element.addClass("ui-widget").find(".toc-title").addClass("ui-widget-header").end().find("li").addClass("ui-widget-content")):"bootstrap"===this.options.theme?(this.element.find(h+","+d).addClass("nav nav-list"),this.focusClass="active"):(this.focusClass=o,this.hoverClass=n),this},setOption:function(){e.Widget.prototype._setOption.apply(this,arguments)},setOptions:function(){e.Widget.prototype._setOptions.apply(this,arguments)},_scrollTo:function(t){var s=this,i=s.options.smoothScroll||0,o=s.options.scrollTo,n=e('div[data-unique="'+t.attr("data-unique")+'"]');return n.length?(e("html, body").promise().done(function(){e("html, body").animate({scrollTop:n.offset().top-(e.isFunction(o)?o.call():o)+"px"},{duration:i})}),s):s}})});
```
