## Supplemental Data Set 1 for "The *pos-1* 3ʹ untranslated region governs germline specification and proliferation to ensure reproductive robustness": scrolltop.js

```
$(document).ready(function(){
	//1.首先将#goTopBtn隐藏
	$("#goTopBtn").hide();

	//2.给窗口添加事件：滚动时触发
	$(window).scroll(function(){
		//当滚动条的位置处于距顶部0像素以上时，返回顶部按钮出现，否则消失
		if ($(window).scrollTop()>0){
			$("#goTopBtn").fadeIn(1500);
			}
		else{
			$("#goTopBtn").fadeOut(1500);
			}
	});
	
	//3.当点击返回顶部按钮后，回到页面顶部位置
	$("#goTopBtn").click(function(){
		$('body,html').animate({scrollTop:0},1000);
	});
});
```
