## Supplemental Data Set 1 for "The *pos-1* 3ʹ untranslated region governs germline specification and proliferation to ensure reproductive robustness": materials_methods_gatk.SNPINDEL.CNV.v1.pdf

### 1. Experimental Procedure

#### 1.1 Sample Quality Control

Please refer to QC report for methods of sample quality control.

#### 1.2 Library Construction, Quality Control and Sequencing

The genomic DNA sample was fragmented into short fragments. These DNA fragments were then end-polished, A-tailed, and ligated with full-length adapters for Illumina sequencing before further size selection. PCR amplification was then conducted unless specified as PCR-free. Purification was then conducted through the AMPure XP system (Beverly). The resulting library was assessed on the Agilent Fragment Analyzer System (Agilent) and quantified to 1.5nM through Qubit (Thermo Fisher Scientific) and qPCR.

The qualified libraries were pooled and sequenced on Illumina platforms, according to the effective library concentration and data amount required.

### 2. Bioinformatics Analysis Pipeline

#### 2.1 Data Quality Control

##### 2.1.1 Raw data

The original fluorescence image files obtained from Illumina platform are transformed to short reads (Raw data) by base calling and these short reads are recorded in FASTQ (Cock et al., 2010) format, which contains sequence information and corresponding sequencing quality information.

##### 2.1.2 Evaluation of data (Data quality control)

Sequence artifacts, including reads containing adapter contamination, low-quality nucleotides and unrecognizable nucleotide (N), undoubtedly set the barrier for the subsequent reliable bioinformatics analysis. Hence quality control is an essential step and applied to guarantee the meaningful downstream analysis.

The steps of data processing were as follows:

(1) Discard a paired reads if either one read contains adapter contamination (>10 nucleotides aligned to the adapter, allowing ≤ 10% mismatches);
(2) Discard a paired reads if more than 10% of bases are uncertain in either one read;
(3) Discard a paired reads if the proportion of low quality (Phred quality <5) bases is over 50% in either one read.

#### 2.2 Reads Mapping to Reference Sequence

Valid sequencing data was mapped to the reference genome by Burrows-Wheeler Aligner (BWA) (Li et al., 2009a) software to get the original mapping results stored in BAM format (parameter: mem -t 4 -k 32 -M). Then, the results were dislodged duplication by SAMtools (Li et al., 2009b) (parameter: rmdup) and Picard (http://broadinstitute.github.io/picard/).

#### 2.3 Variant detection and annotation

##### 2.3.1 SNP/InDel

The raw SNP/InDel sets are called by GATK(DePristo et al., 2011) with the parameters as ' – gcpHMM 10 -stand_emit_conf 10 -stand_call_conf 30'. Then we filtered this sets using the following criteria:

SNP: QD < 2.0, FS > 60.0, MQ < 30.0, HaplotypeScore > 13.0, MappingQualityRankSum < -12.5, ReadPosRankSum < -8.0

INDEL: QD < 2.0, FS > 200.0, ReadPosRankSum < -20.0

##### 2.3.2 CNV

CNVs were detected by CNVnator (Abyzov et al., 2011) to acquire potential deletion and duplication (parameter: - call 100).

#### 2.4 Annotation

ANNOVAR (Wang et al., 2010) was used for functional annotation of variants. The UCSC known genes were used for gene and region annotations.
