## Supplemental Data Set 1 for "The *pos-1* 3ʹ untranslated region governs germline specification and proliferation to ensure reproductive robustness": materials_methods_samtools.SNPINDEL.SV.CNV_freec.v1.pdf

(1) The depth of the variate position >4;

(2) The mapping quality >20.

#### 2.3.2 CNV and SV

CNVs were detected by Control-FREEC(Barillot et al., 2012)to acquire potential deletion and duplication (parameter: window=100 step = 100).
SVs were detected by BreakDancer (Chen et al., 2009).

### 2.4 Annotation

ANNOVAR (Wang et al., 2010) was used for functional annotation of variants. The UCSC known genes were used for gene and region annotations.

Cock PJ, Fields CJ, Goto N, Heuer ML, Rice PM. The Sanger FASTQ file format for sequences with quality scores, and the Solexa/Illumina FASTQ variants.

Nucleic Acids Res. 2010;38(6):1767-1771. doi:10.1093/nar/gkp1137

Li H, Durbin R. Fast and accurate short read alignment with Burrows-Wheeler transform. Bioinformatics. 2009a;25(14):1754-1760. doi:10.1093/bioinformatics/btp324

Li H, Handsaker B, Wysoker A, et al. The Sequence Alignment/Map format and SAMtools. Bioinformatics. 2009b;25(16):2078-2079. doi:10.1093/bioinformatics/btp352

Wang K, Li M, Hakonarson H. ANNOVAR: functional annotation of genetic variants from high-throughput sequencing data. Nucleic Acids Res. 2010;38(16):e164. doi:10.1093/nar/gkq603
