## Supplemental Data Set 1 for "The *pos-1* 3ʹ untranslated region governs germline specification and proliferation to ensure reproductive robustness": mWGS-Method.pdf

### Methods

The method is only used as a reference for publication purpose.
Customers are responsible for the related risks of duplicate checking.

### 1. Experimental Procedure

##### 1.1 Sample Quality Control

Please refer to Novogene's QC report for methods of sample quality control.

##### 1.2 Library Construction, Quality Control and Sequencing

The genomic DNA sample was fragmented into short fragments. These DNAfragments were then end-polished, A-tailed, and ligated with full-length adapters forIllumina sequencing before further size selection. PCR amplification was thenconducted unless specified as PCR-free. Purification was then conducted throughthe AMPure XP system (Beverly). The resulting library was assessed on the AgilentFragment Analyzer System (Agilent) and quantified to 1.5 nM through Qubit (ThermoFisher Scientific) and qPCR.

The qualified libraries were pooled and sequenced on Illumina platforms, according tothe effective library concentration and data amount required.

### 2. Bioinformatic analysis Pipeline

##### 2.1 Data quality control

2.1.1 Raw data

The original fluorescence image files obtained from Illumina platformare transformed to short reads (Raw data) by base calling and these short reads arerecorded in FASTQ format (Chen et al., 2018), which contains sequence informationand corresponding sequencing quality information.

2.1.2 Evaluation of data (Data quality control)

Sequence artifacts, including reads containing adapter contamination,low-quality nucleotides and unrecognizable nucleotide (N), undoubtedly set thebarrier for the subsequent reliable bioinformatics analysis. Hence quality control is an essential step and applied to guarantee the meaningful downstream analysis. weused Fastp (version 0.23.1) (Chen et al., 2018) to perform basic statistics on thequality of the raw reads. The steps of data processing were as follows:

(1) Discard a paired reads if either one read contains adapter contamination( >10nucleotides aligned to the adapter, allowing ≤ 10% mismatches);

(2) Discard a paired reads if more than 10% of bases are uncertain in either one read;

(3) Discard a paired reads if the proportion of low quality (Phred quality <5) bases is over50% in either one read.

#### 2.2 Reads Mapping to Reference Sequence

Valid sequencing data was mapped to the reference genome by Burrows Wheeler Aligner (BWA) (Li et al., 2009a) software to get the original mapping results stored in BAM format (parameter: mem -t 4 -k 32 -M). Then, the results were dislodged duplication by SAMtools (Li et al., 2009b) (parameter: rmdup) and Picard (http://broadinstitute.github.io/picard/).

#### 2.3 Variant detection and annotation

##### 2.3.1 SNP/InDel

The raw SNP/InDel sets are called by GATK(DePristo et al., 2011) with the parameters as ' – gcpHMM 10 -stand_emit_conf 10 -stand_call_conf 30'. Then we filtered this sets using the following criteria:

SNP: QD < 2.0, FS > 60.0, MQ < 30.0, HaplotypeScore > 13.0,

MappingQualityRankSum < -12.5, ReadPosRankSum < -8.0

INDEL: QD < 2.0, FS > 200.0, ReadPosRankSum < -20.0

##### 2.3.2 CNV and SV

CNVs were detected by CNVnator (Abyzov et al., 2011) to acquire potential deletion and duplication (parameter: - call 100).

SVs were detected by BreakDancer(Chen et al., 2009).

#### 2.4 Annotation

ANNOVAR (Wang et al., 2010) was used for functional annotation of variants. The UCSC known genes were used for gene and region annotations.

KOG (Fungi)：Currently, the universal functional database used is the KOG database. The basic steps for functional annotation are as follows:

1) Perform a BLAST search between the predicted gene and various functional databases;
2) Filter the results of the BLAST search, selecting the one with the highest score for each sequence.

COG (Bacteria)：Currently, the universal functional database used is the COG database. The basic steps for functional annotation are as follows:

1) Perform a BLAST search between the predicted gene and various functional databases;

2) Filter the results of the BLAST search, selecting the one with the highest score for each sequence.
