## Supplemental Data Set 1 for "The *pos-1* 3ʹ untranslated region governs germline specification and proliferation to ensure reproductive robustness": result_tree.html

Directory Tree


### Directory Tree

.
  
 |-- 01.OriginalData
  
 |-- 02.QualityControl
  
 | |-- CleanData\_QCsummary  
 | |-- ErrorRate  
 | |-- QualityDistribution  
 | `-- ReadsClassification  
 |-- 03.Mapping
  
 | |-- MapStat  
 | `-- Reference  
 |-- 04.SNP\_VarDetect
  
 | |-- DG4222  
 | |-- WRM101  
 | |-- WRM102  
 | |-- WRM103  
 | `-- picture\_in\_reports  
 |-- 05.InDel\_VarDetect
  
 | |-- DG4222  
 | |-- WRM101  
 | |-- WRM102  
 | |-- WRM103  
 | `-- picture\_in\_reports  
 |-- 06.SV\_VarDetect
  
 | |-- DG4222  
 | |-- WRM101  
 | |-- WRM102  
 | |-- WRM103  
 | `-- picture\_in\_reports  
 |-- 07.CNV\_VarDetect
  
 | |-- DG4222  
 | |-- WRM101  
 | |-- WRM102  
 | |-- WRM103  
 | `-- picture\_in\_reports  
 `-- 08.VarDetect\_Visualization

34 directories

---

tree v1.5.3 (c) 1996 - 2009 by Steve Baker and Thomas Moore   
HTML output hacked and copyleft (c) 1998 by Francesc Rocher   
Charsets / OS/2 support (c) 2001 by Kyosuke Tokoro
