## Supplemental Data Set 1 for "The *pos-1* 3ʹ untranslated region governs germline specification and proliferation to ensure reproductive robustness": X202SC24112711-Z01-F001_C_elegans_Report.html

 H202SC24112711\_X202SC24112711-Z01-F001\_C\_elegans Plant and Animal Whole Genome Sequencing Standard Analysis Report 

### Plant and Animal Whole Genome Sequencing Standard Analysis Report

|  |  |
| --- | --- |
| Contract ID | H202SC24112711 |
| Contract Name | Davis-US-UMassMed-4-C. elegans-PAWGS-NovaseqXplus-3G-WBI-Advanced-NVUS2024103051 |
| Batch ID | X202SC24112711-Z01-F001 |
| Reference Genome and Version | https://ftp.ncbi.nlm.nih.gov/genomes/all/GCF/000/002/985/GCF\_000002985.6\_WBcel235/GCF\_000002985.6\_WBcel235\_genomic.fna.gz ncbi\_caenorhabditis\_elegans\_gcf\_000002985\_6\_wbcel235 |
| Report Time | 2024-11-26 |
 Reminder | Partial results are presented in this report, while full results will be delivered in data release.  Hyperlink of results in this report will be only valid in data release, after statement confirmation. |

  
  

---

  

#### 1 Introduction

Plant and animal whole genome resequencing via Illumina platforms, based on mechanism of SBS (sequencing by synthesis), has become the most rapid and effective method to identify the genetic variations in individuals of the same species or between related species. The variation information such as Single Nucleotide Polymorphism (SNP), Insertion and Deletion (InDel), Copy Number Variation (CNV), and structural variation (SV) obtained through resequencing is used in population genetics research and genome-wide association studies (GWAS) to investigate the causes of diseases, to select plants and animals for agricultural breeding programs, and to identify common genetic variations among populations. Plant and animal whole genome resequencing projects are carried out as follows:

Figure 1 Project workflow

---

  

#### 2 Library Construction and Sequencing

##### 2.1 Sample Quality Control

    Please refer to Novogene’s sample QC report.

##### 2.2 Library Construction , Quality Control and Sequencing

The genomic DNA was randomly sheared into shorter fragments. The obtained fragments were then end-repaired, A-tailed, and further ligated with Illumina adapters. The resulting fragments with adapters were size selected, PCR amplified unless otherwise specified as PCR-free, before proceeding for purification.
The experimental procedures of DNA library preparation are shown in Figure 2.

Figure 2 Workflow of library Construction

The library was quantified through Qubit and qPCR, and size distribution detected with fragment analyzer. Quantified libraries were pooled and sequenced on Illumina platforms according to the effective library concentration and required data amount.

---

  

#### 3 Bioinformatics Analysis Pipeline

The bioinformatic analysis workflow of raw sequencing data is shown as follows：

Figure 3 Bioinformatic analysis workflow

**Method details involved in the project are available at
methods**

---

#### 4 Analysis Results

##### 4.1 Raw Data

Original image data file from high-throughput sequencing platforms (like Illumina) is transformed to sequenced reads (called Raw Data or Raw Reads) by CASAVA base recognition (Base Calling). Raw data are stored in FASTQ (.fq) format files(Cock et al., 2010), which contain sequencing reads and corresponding base quality. Each read has four descriptive lines, as indicated below:

@K00124:82:H2MH5BBXX:1:1101:31389:1158 2:N:0:0  
TAGCCACATAGAAACCAACAGCCATATAACTGGTAGCTTTAAGCGGCTCACCTTTAGCATCAACAGGCCACAACCAACCAGAACGTGAAAAAGCGTCCTGCGTGTAGCGAACTGCGATGGGCATACAGATCGGAAGAGCGTCGTGTAGGG  
+  
AAFFFKKKKKKKKFKKKFFKKAAFKKKKKFKKKKFKKA,FKKKKKKKKKAKKFKKKKKKKAKKKKKKFFKKKKF<FFKKKKKKKKKKKKKFKKFKKF7FFFFFKFKKKFKKKKKKKKF<FFKKKKFKKKKKFKFKFKKFK<<F,A7,AFK

Line 1: begins with the at sign (@), followed by sequence identifiers and an optional description (such as FASTA header).  
Line 2: base sequences (raw read, A, G, C, and T)  
Line 3: begins with the plus sign (+), optionally followed by the same Illumina sequence identifiers and description information as in Line 1.  
Line 4: the quality values for each base, corresponding to the data in Line 2.

Table 4.1 Illumina sequence identifier details

|  |  |
| --- | --- |
| K00124 | Unique instrument name |
| 82 | Run ID |
| H2MH5BBXX | Flowcell ID |
| 1 | Flowcell lane |
| 1101 | Tile number within the flowcell lane |
| 31389 | 'x'-coordinate of the cluster within the tile |
| 1158 | 'y'-coordinate of the cluster within the tile |
| 2 | Member of a pair, 1 or 2 (paired-end or mate-pair reads only) |
| N | Y if the read fails filter (read is bad), N otherwise |
| 0 | 0 when none of the control bits are on, otherwise it is an even number |
| ATCACG | Index sequence |

Result：01.OriginalData

---

##### 4.2 Data Quality Control

###### 4.2.1 Sequencing Quality Distribution

If the sequencing error rate is represented by e, and Illumina sequencing quality by Q_phred_, the quality score of a base (Phred score) is calculated by the following equation: Q_phred_=-10log_10_(e). The correspondence relationship between Illumina sequencing quality and Phred score in base calling by Casava version 1.8 is listed as follows:

Table 4.2 Sequencing error rate and corresponding base quality value

| Phred score | Error Rate | Correct Rate | Q-score |
| --- | --- | --- | --- |
| 10 | 1/10 | 90% | Q10 |
| 20 | 1/100 | 99% | Q20 |
| 30 | 1/1000 | 99.9% | Q30 |
| 40 | 1/10000 | 99.99% | Q40 |

For next-generation sequencing (NGS), the sequencing platform, chemical reactants, and sample quality can influence sequencing quality and base error rate. Sequencing quality distribution is examined over the full length of all sequences, to detect any sites (base positions) with an unusually low sequencing quality, where incorrect bases may be incorporated at abnormally high levels. For detailed sequencing quality distribution, please refer to **Figure 4.1**.

DG4222\_CKDN240021609-1A\_22HTYHLT4\_L3
WRM101\_CKDN240021610-1A\_22HTVFLT4\_L1
WRM102\_CKDN240021611-1A\_22HVGTLT4\_L1
WRM103\_CKDN240021612-1A\_22HVGTLT4\_L2

Figure 4.1 Distribution of sequencing quality

The x-axis shows the base position within a sequencing read, and the y-axis shows the average phred score of all reads at each position.  
(Pair-end sequencing data are plotted together, with the first PE150 bp representing read 1 and the following PE150 bp for read 2.)

Result：02.QualityControl/QualityDistribution

---

  

###### 4.2.2 Sequencing Error Rate

Sequencing error rate and base quality can be affected by various factors such as sequencing platform, chemical reagent and sample quality. Due to the consumption of chemical reagents, error rate is increasing with read extension, which is a common feature of Illumina high throughput sequencing platforms. The sequencing error rate distribution can be featured as below:

(1) Error rate increases with the sequencing reads for consumption of sequencing reagent. It is common in the Illumina high-throughput sequencing platform.  
(2) The sequencing error rate is higher for the first several bases than at other positions, which is likely the result of reading errors during the first few cycles after calibration of the optical instruments.

  

DG4222\_CKDN240021609-1A\_22HTYHLT4\_L3
WRM101\_CKDN240021610-1A\_22HTVFLT4\_L1
WRM102\_CKDN240021611-1A\_22HVGTLT4\_L1
WRM103\_CKDN240021612-1A\_22HVGTLT4\_L2

Figure 4.2 Distribution of sequencing error rate

The x-axis shows the base position along each sequencing read and the y-axis shows the base error rate.  
(Pair-end sequencing data are plotted together, with the first PE150 bp representing read 1 and the following PE150 bp for read 2.)

Result：02.QualityControl/ErrorRate

---

  

###### 4.2.3 Sequencing Data Filtration

The sequencing reads/raw reads often contain low-quality reads or reads with adaptors, which will affect the quality of downstream analysis. To avoid this, it is necessary to filter the raw reads and obtain clean reads. Raw reads filtering is as follows:

      (1) Remove the paired reads when either read contains adapter contamination;  
      (2) Remove the paired reads when uncertain nucleotides (N) constitute more than 10 percent of either read;  
      (3) Remove the paired reads when low-quality nucleotides (base quality less than 5, Q ≤ 5) constitute more than 50 percent of either read.

DG4222\_CKDN240021609-1A\_22HTYHLT4\_L3
WRM101\_CKDN240021610-1A\_22HTVFLT4\_L1
WRM102\_CKDN240021611-1A\_22HVGTLT4\_L1
WRM103\_CKDN240021612-1A\_22HVGTLT4\_L2

Figure 4.3 Classification of the sequenced reads

Show Annotation

(1) Adapter related: (reads containing adapter) / (total raw reads).
(2) Containing N: (reads with more than 10% N) / (total raw reads).
(3) Low quality: (reads of low quality) / (total raw reads).
(4) Clean reads: (clean reads) / (total raw reads).

Result：02.QualityControl/ReadsClassification

---

  

###### 4.2.4 Statistics of Sequencing Data

According to the sequencing feature of Illumina platforms, for paired-end sequencing data we require that Q30 (the percent of bases with phred-scaled quality scores greater than 30) should be above 80%. Statistics of sequencing data are listed in **Table 4.3**.

Table 4.3 Statistics of Sequencing Data

| Sample name | Lane | Raw reads | Raw data (G) | Clean data (G) | Effective (%) | Error (%) | Q20 (%) | Q30 (%) | GC (%) |
| --- | --- | --- | --- | --- | --- | --- | --- | --- | --- |
| DG4222 | CKDN240021609-1A\_22HTYHLT4\_L3 | 29556418 | 4.4 | 4.4 | 98.43 | 0.01 | 98.87 | 96.94 | 35.67 |
| WRM101 | CKDN240021610-1A\_22HTVFLT4\_L1 | 29247208 | 4.4 | 4.3 | 98.49 | 0.01 | 98.90 | 97.19 | 35.91 |
| WRM102 | CKDN240021611-1A\_22HVGTLT4\_L1 | 40890294 | 6.1 | 6.0 | 98.08 | 0.01 | 98.96 | 97.09 | 36.77 |
| WRM103 | CKDN240021612-1A\_22HVGTLT4\_L2 | 46303002 | 6.9 | 6.8 | 98.61 | 0.01 | 98.90 | 96.84 | 39.75 |

Show Annotation

(1) Sample name: Sample name.  
(2) Lane: The flowcell ID and lane number during the sequencing (FlowcellID\_LaneNumber).  
(3) Raw reads: The number of sequencing reads pairs; four lines will be considered as one unit according to FASTQ format.   
(4) Raw data (G): The original sequence data volume.  
(5) Clean data (G): The sequence data volume calculated by clean data.  
(6) Effective (%): The ratio of clean data to raw data.  
(7) Error (%): Overall error rate of base.  
(8) Q20 (%): The percentage of bases with higher Phred score than 20.  
(9) Q30 (%): The percentage of bases with higher Phred score than 30.  
(10) GC: The percentage of G and C in the total bases.

Result：02.QualityControl/CleanData\_QCsummary

Show Q&A

Q.: As the sequencing error increases with the read length, what is the acceptable range of sequencing error rate?

A.: Currently for Illumina sequencers, the per base error rate is generally lower than 1%, and the highest acceptable threshold is 6%.

Q.: What are the data filtering criteria at Novogene? Are they strictly adhered?

A.: Novogene utilizes stringent data quality control procedures and strictly adheres to the high standard to guarantee the accuracy and reliability of the sequencing data. The detailed data quality control criteria are as follows:  
       **(1)** Discard the adapter-containing reads;  
       **(2)** Discard the paired reads when uncertain nucleotides (N) constitute more than 10 percent of either read;  
       **(3)** Discard the paired reads when low quality nucleotides (base quality less than 5, Q ≤ 5) constitute more than 50 percent of either read.

List of Related Terms:

**adapter**: the oligo nucleotides ligated to sample DNA at library preparation, for proper adhesion to the flow-cell via base-pairing in DNA sequencing.

**index**: the unique sequence tag for distinguishing each individual sample from multiplexing.

**Q20,Q30**: the proportion of bases with Phred score higher than 20 or 30. The Phred score, which is negatively correlated to the probability of incorrect base-calling, is calculated by the equation (Q_phred_=-10log_10_(e)) of sequencing error rate (e), indicating the sequencing quality.

**raw data/raw reads**: the original sequence data output by the instrument for a specific sample.

**clean data/clean reads**: the output of data from filtering raw data with quality control, which will be used for further analysis.

---

---

  

###### 4.3.2 Mapping Statistics with Reference Genome

The mapping rates of samples reflect the similarity between each sample and the reference genome. The depth and coverage are indicators of the evenness and homology with the reference genome.

Table 4.5 Statistics of mapping rate, depth and coverage

| Sample | Mapped reads | Total reads | Mapping rate (%) | Average depth(X) | Coverage at least 1X (%) | Coverage at least 4X (%) | Duplicate (%) |
| --- | --- | --- | --- | --- | --- | --- | --- |
| DG4222 | 29,042,108 | 29,092,026 | 99.83 | 37.71 | 99.97 | 99.94 | 13.08 |
| WRM101 | 28,399,240 | 28,805,826 | 98.59 | 32.16 | 99.97 | 99.93 | 24.15 |
| WRM102 | 37,289,939 | 40,106,664 | 92.98 | 45.03 | 99.97 | 99.95 | 19.20 |
| WRM103 | 33,326,262 | 45,658,508 | 72.99 | 38.56 | 99.97 | 99.94 | 22.53 |

Show Annotation

(1) Sample: Sample names.  
(2) Mapped reads: The number of clean reads mapped to the reference assembly, including both single-end reads and reads in pairs.  
(3) Total reads: Total number of effective reads in clean data.  
(4) Mapping rate: The ratio of the reference genome assembly mapped reads to the total sequenced clean reads.  
(5) Average depth: The average depth of mapped reads at each site, calculated by the total number of bases in the mapped reads dividing by size of the assembled genome.  
(6) Coverage at least 1X: The percentage of the assembled genome with more than one read at each site.  
(7) Coverage at least 4X: The percentage of the assembled genome with ≥4X coverage at each site.  
(8) Duplicate (%):Proportion of duplicated reads (data from Mapping).

DG4222
WRM101
WRM102
WRM103

Figure 4.4 The mean depth of each chromosome.

###### 4.3.3 Mapping Summary

For the current 100,286,401 bp reference genome, the mapping rate of each sample ranges from 72.99% to 99.83%. Refer to the reference genome (without Ns), the average depths are between 32.16X and 45.03X, and 1X coverages range from 99.97% to 99.97%. This result is in the qualified normal range and may serve in the subsequent variation detection and related analyses.

###### 4.3.4 Visualization of mapping results

We provide BAM format files of mapping results, and corresponding reference genome and annotation files for some species. IGV (integrated genomics viewer) browser is recommended to visualize BAM files.

IGV browser has the following features:

(1) It can display the position of single or multiple reading segments on the genome at different scales, including the distribution of reading segments on each chromosome and the distribution of exon, intron, splicing junction area and intergenic area of annotation, etc;

(2) It can display the read segment abundance of different regions in different scales to reflect the number of reads support of different regions;

(3) It can display other annotation information;

(4) You can download all kinds of annotation information from the remote server, and load the annotation information locally.

This guide describes the Integrative Genomics Viewer (IGV). To start IGV, go to the IGV downloads page: http://www.broadinstitute.org/igv/download

Look at a printer-friendly HTML version of the whole User Guide: http://software.broadinstitute.org/software/igv/book/export/html/6

Figure 4.5 IGV browser

Result：03.Mapping

Show Q&A

Q.: Which files are required for reference genome mapping?

A.: As for whole genome sequencing (WGS), only the corresponding reference genome file in FASTA format is required, while the whole exome sequencing (WES) also needs the target region file in BED format.

Q.: What are the potential causes of low mapping rate?

A.: The three major causes are: (1) with poorly assembled reference genome or a relatively far genetic relationship between the reference and the sample; (2) treatments to DNA that may change the sequence (e.g. bisulfite treatment) and herein affecting the mapping rate and (3) contamination of other species.

Q.: Could the full-length reads be mapped to the reference? or should we trim from the sequence ends?

A.: According to the standard Illumina pair-end (PE) sequencing protocol, DNA fragments were ligated with adapters at both ends. The adapter habors universal sequences for flow-cell binding, DNA sequencing, as well as unique index located upstream of the sequencing primer for multiplexing. As for the PE150 sequencing of 350 bp library, the sequenced reads of PE150 bp at either end are adapter-free, which could be directly subjected to quality control for low quality reads filtration. The retaining sequences in PE150 bp length (namely clean data) are qualified for mapping with the reference genome.

---

  

##### 4.4 SNP Detection & Annotation

Single nucleotide polymorphism (SNP) refers to a variation in a single nucleotide which may occur at some specific position in the genome, including transition and transversion of a single nucleotide. We detected the individual SNP variations using SAMtools (Li et al., 2009b) with the following parameter: 'mpileup -m 2 -F 0.002 -d 1000 -u -C 50'.

To reduce the error rate in SNP detection, we filtered the results with the criteria as follows:

       (1) The number of support reads for each SNP should be more than 4 and less than 1000;  
       (2) The mapping quality (MQ) of each SNP should be higher than 20.

###### 4.4.1 Statistics of SNP Detection & Annotation

ANNOVAR is a widely used software in variation annotation with multiple capabilities, including gene-based annotation, region-based annotation, filter-based annotation as well as other functionalities(Wang et al., 2010). Novogene used ANNOVAR to annotate the detected SNPs.

Table 4.6 Statistics of SNP detection and annotation

| Sample | Upstream | Exonic | | | | Intronic | Splicing | Downstream | Upstream/Downstream | Intergenic | 3'UTR | 5'UTR | 5'UTR/3'UTR | Others | ts | tv | ts/tv | Het rate(‰) | Total |
| --- | --- | --- | --- | --- | --- | --- | --- | --- | --- | --- | --- | --- | --- | --- | --- | --- | --- | --- | --- |
|  |  | Stop gain | Stop loss | Synonymous | Non-synonymous |  |  |  |  |  |  |  |  |  |  |  |  |  |  |
| DG4222 | 281 | 3 | 0 | 158 | 228 | 936 | 5 | 419 | 201 | 1795 | 38 | 9 | 0 | 99 | 2126 | 2079 | 1.023 | 0.033 | 4205 |
| WRM101 | 274 | 3 | 1 | 159 | 213 | 919 | 4 | 379 | 201 | 1663 | 40 | 14 | 1 | 88 | 1997 | 1989 | 1.004 | 0.031 | 3986 |
| WRM102 | 286 | 3 | 2 | 160 | 214 | 959 | 4 | 388 | 198 | 1829 | 49 | 8 | 0 | 97 | 2158 | 2074 | 1.041 | 0.033 | 4232 |
| WRM103 | 289 | 4 | 2 | 198 | 251 | 1059 | 5 | 433 | 225 | 1816 | 52 | 15 | 1 | 94 | 2227 | 2253 | 0.988 | 0.037 | 4480 |

Show Annotation

(1) Sample: Sample name.  
(2) Upstream: SNPs located within 1 kb upstream (away from transcription start site) of the gene.  
(3) Exonic: SNPs located in exonic region; Non-synonymous: single nucleotide mutation with changing amino acid sequence; Stop gain/loss: a nonsynonymous SNP that leads to the introduction/removal of stop codon at the variant site; Synonymous: single nucleotide mutation without changing amino acid sequence.  
(4) Intronic: SNPs located in intronic region.  
(5) Splicing: SNPs located in the splicing site (2 bp range of the intron/exon boundary).  
(6) Downstream: SNPs located within 1 kb downstream (away from transcription termination site) of the gene region.  
(7) Upstream/Downstream: SNPs located within the < 2 kb intergenic region, which is in 1 kb downstream or upstream of the genes.  
(8) Intergenic: SNPs located within the > 2 kb intergenic region.  
(9) 3'UTR: Variants within the 3'UTR.  
(10) 5'UTR: Variants within the 5'UTR.  
(11) 5'UTR/3'UTR: Variants within 3'UTR and 5'UTR jointly.  
(12) Others: SNPs located in other region.  
(13) ts: Transitions, a point mutation that changes a purine nucleotide to another purine (A ↔ G) or a pyrimidine nucleotide to another pyrimidine (C ↔ T). Approximately two out of three SNPs are transitions.  
(14) tv: Transversions, the substitution of a (two ring) purine for a (one ring) pyrimidine or *vice versa*.  
(15) ts/tv: The ratio of transitions to transversions.  
(16) Het rate: Genome-wide heterozygous rate, calculated by the ratio of heterozygous SNPs to the total number of genome bases.  
(17) Total: The total number of SNPs.  
\*Please note: The statistical data of Upstream、Intronic、Splicing、Downstream、Upstream/Downstream、Intergenic and Others is derived from genome-wide annotation file (\*snp.avinput.variant\_function.gz);The statistical data of Exonic including Stop gain, Stop loss, Synonymous, Non-synonymous comes from the exon annotation file (\*snp.avinput.exonic\_variant\_function.gz); The statistical data of ts, tv, and Total indicating variant sites number comes from the variants detection file (\*filter.snp.vcf.gz). Multiallelic sites of identical SNP site are possibly annotated to different annotation types. Additionally, the sum of the number of annotation types possibly unequal to the total number of variants.

DG4222
WRM101
WRM102
WRM103

Figure 4.6 The number of SNPs in different regions of the genome

The number of SNPs in different regions of the genome (left) and the number of SNPs of different types in the coding region (right) are distributed.

---

  

###### 4.4.2 SNP Quality Distribution

To assess the credibility of detected SNPs, we checked the distribution of support reads number, SNP quality, as well as the distance between adjacent SNPs. The results are shown in **Figure 4.7**.

SNP\_readsNum\_cumulative\_distribution
SNP\_distance\_cumulative\_distribution
SNP\_quality\_cumulative\_distribution

Figure 4.7 Cumulative distribution of SNP quality

Note: These figures show the quality distribution of SNPs by, from top to bottom, the distribution of SNP support reads number, the distribution of distances between adjacent SNPs, and the cumulative distribution of SNP quality.

---

  

###### 4.4.3 SNP Mutation Frequency

Take the T:A>C:G mutations as an example, this category includes mutations from T to C and A to G. When T>C mutation appears on either of the double-strand, the A>G mutation will be found in the same position of the other chain. Therefore the T>C and A>G mutations are classified into one category. Accordingly, the whole genome SNP mutations could be classified into six categories. The frequency of each type is shown in **Figure 4.8**.

Figure 4.8 Frequency of SNP mutations

The x-axis represents the number of the SNPs, and y-axis indicates the mutation types.

Result：04.SNP\_VarDetect

Show Q&A

Q.: What is the MQ quality for a SNP?

A.: The SNP quality is represented by the mapping quality of covering reads, calculated by the root-mean-square of the support reads' mapping quality.

Q.: What is QUAL value for a SNP?

A.: The QUAL value is the Phred quality score (QUAL), represents the probability (*p*) of SNP truly existing at certain position. The higher the QUAL value, the more likely the SNP exists. The relationship between QUAL and *p* is QUAL=-10\*log_10_ (1-*p*). Therefore, the QUAL value of 20 means the probability of the existence of this SNP is 99%.

Q.: What are transitions and transversions?

A.: Transitions refers to the changes between A and G, which are both purines, or between C and T, which are both pyrimidines; while transversions reprents changes between a purine and a pyrimidine, such as between A and T.

Q.: What is the heterozygous SNP?

A.: Heterozygous SNPs are those called with both REF (the same to reference) and ALT (different from reference) genotypes in a diploid species.

Q.: How to verify the SNP genotypes?

A.: The "golden standard" of SNP verification is PCR amplification followed by Sanger sequencing.

Q.: If the PCR-sequencing method failed to verify the detected SNP, did this mean that the NGS SNP calling is not reliable?

A.: SNP calling in NGS is based on the support reads and depended on the sufficient coverage depth, which ensures the accuracy of most but not all the detected SNPs. We recommend to double-check the PCR results first, then use a genome browser such as *Savant* and *IGV* to manually check the mapped reads of the NGS result.

---

  

##### 4.5 InDel Detection & Annotation

InDel refers to the small insertion or deletion in the DNA. We detected the individual SNP variations using SAMtools (Li et al., 2009b) with the following parameter: 'mpileup -m 2 -F 0.002 -d 1000 -u -C 50'. The filter conditions are same with SNPs.

###### 4.5.1 Statistics of InDel Detection & Annotation

Table 4.7 Statistics of InDel detection and annotation

| Sample | Upstream | Exonic | | | | | | Intronic | Splicing | Downstream | Upstream/Downstream | Intergenic | 3'UTR | 5'UTR | 5'UTR/3'UTR | Others | Insertion | Deletion | Het rate(‰) | Total |
| --- | --- | --- | --- | --- | --- | --- | --- | --- | --- | --- | --- | --- | --- | --- | --- | --- | --- | --- | --- | --- |
|  |  | Stop gain | Stop loss | Frameshift deletion | Frameshift insertion | Non-frameshift deletion | Non-frameshift insertion |  |  |  |  |  |  |  |  |  |  |  |  |  |
| DG4222 | 220 | 0 | 0 | 11 | 11 | 6 | 4 | 814 | 7 | 176 | 190 | 402 | 67 | 11 | 2 | 122 | 1094 | 948 | 0.010 | 2043 |
| WRM101 | 221 | 0 | 0 | 11 | 9 | 9 | 4 | 798 | 6 | 189 | 178 | 400 | 57 | 11 | 1 | 131 | 1070 | 955 | 0.010 | 2025 |
| WRM102 | 197 | 0 | 0 | 9 | 9 | 14 | 3 | 798 | 7 | 184 | 171 | 413 | 62 | 9 | 2 | 124 | 1071 | 925 | 0.010 | 2001 |
| WRM103 | 226 | 0 | 0 | 8 | 11 | 9 | 5 | 850 | 5 | 177 | 193 | 414 | 65 | 15 | 1 | 141 | 1133 | 985 | 0.011 | 2120 |

Show Annotation

(1) Sample: Sample names.  
(2) Upstream: InDels located within 1 kb upstream (away from transcription start site) of the gene.  
(3) Exonic: InDels located in exonic region; Stop gain/loss: InDel that leads to the introduction/removal of stop codon at the variant site; Frameshift deletion/insertion: InDel mutation changing the open reading frame with deletion or insertion; Non-Frameshift deletion/insertion:
InDel mutation without changing the open reading frame with deletion or insertion sequences of 3 or multiple of 3 bases;
  
(4) Intronic: InDel located in intronic region;  
(5) Splicing: InDel located in the splicing site (2 bp range of the intron/exon boundary).  
(6) Downstream: InDel located within 1 kb downstream (away from transcription termination site) of the gene region.  
(7) Upstream/Downstream: InDel SNPs located within the < 2 kb intergenic region, which is in 1 kb downstream or upstream of the genes.  
(8) Intergenic: InDel located within the > 2 kb intergenic region.  
(9) 3'UTR: Variants within the 3'UTR.  
(10) 5'UTR: Variants within the 5'UTR.  
(11) 5'UTR/3'UTR: Variants within 5'UTR and 3'UTR jointly.  
(12) Others: InDel located in other region.  
(13) Het rate: InDel heterozygous rate, calculated by the ratio of InDels to the total number of genome bases.  
(14) Total: The total number of InDels.  
\*Please note: The statistical data of Upstream、Intronic、Splicing、Downstream、Upstream/Downstream、Intergenic and Others is derived from genome-wide annotation file (\*indel.avinput.variant\_function.gz);The statistical data of Exonic including Stop gain, Stop loss, Frameshift deletion/insertion and Non-Frameshift deletion/insertion comes from the exon annotation file (\*indel.avinput.exonic\_variant\_function.gz);The statistical data of Total indicating variant sites number comes from variant detection file (\*filter.indel.vcf.gz);Multiallelic sites of identical InDel site are possibly annotated to different structural intervals.Additionally, the sum of the number of annotation types possibly unequal to the total number of variants.

DG4222
WRM101
WRM102
WRM103

Figure 4.9 The number of InDels in different regions of the genome

The number of InDels in different regions of the genome (left) and the number of InDels of different types in the coding region (right) are distributed.

---

  

###### 4.5.2 Length Distribution of InDels

The figure 4.10 show the length distribution of InDels for the random four samples in the sample list. Please refer to the statistical result file (./results/05.InDel\_VarDetect/Indel.GENOMEpercentage.xls and Indel.CDSpercentage.xls) for the details of all the samples.

Figure 4.10 Length distribution of InDels

The x-axis represents the proportion of the InDels with a certain length, and y-axis indicates the length of the InDels.

Result：05.InDel\_VarDetect

Show Q&A

Q.: What is the heterozygous rate for an InDel?

A.: Ratio of heterozygous InDels to total number of InDels, the heterozygous Indel is an Indel only located in one of the homologous chromosomes of a diploid sample.

Q.: What's the difference between an InDel and a SNP?

A.: SNP mutation refers to the change from one nucleotide to another nucleotide (eg. A↔T), while InDel is a type of mutation with insertion or deletion of one or more nucleotides (eg, A↔AT, with a T insertion).

---

  

##### 4.6 SV Detection & Annotation

Structural variants (SVs) are genomic variations with mutations of relatively larger size (>50 bp), including deletions, duplications, insertions, inversions and translocations. BreakDancer(Chen et al., 2009) software was used to detect insertion (INS), deletion (DEL), inversion (INV), intra-chromosomal translocation (ITX) and inter-chromosomal translocation (CTX) mutations, based on the reference genome mapping results and the detected insert size. The detected SVs were filtered by removing those with less than 2 supporting PE reads, the INS, DEL and INV were further annotated by ANNOVAR(Wang et al., 2010).

###### 4.6.1 Statistics of SV Detection & Annotation

Table 4.8 Statistics of SV detection and annotation

| Sample | Upstream | Exonic | Downstream | Intronic | Upstream/Downstream | Intergenic | Splicing | Others | INS | DEL | INV | ITX | CTX | Total |
| --- | --- | --- | --- | --- | --- | --- | --- | --- | --- | --- | --- | --- | --- | --- |
| DG4222 | 11 | 73 | 21 | 35 | 10 | 88 | 0 | 20 | 3 | 206 | 49 | 767 | 96 | 1121 |
| WRM101 | 14 | 100 | 30 | 31 | 13 | 87 | 0 | 21 | 2 | 252 | 42 | 703 | 92 | 1091 |
| WRM102 | 12 | 68 | 32 | 40 | 9 | 112 | 0 | 14 | 3 | 225 | 59 | 884 | 189 | 1360 |
| WRM103 | 8 | 70 | 25 | 23 | 11 | 84 | 0 | 20 | 2 | 195 | 44 | 797 | 117 | 1155 |

Show Annotation

(1) Sample: Sample names.  
(2) Upstream: SVs located within 1 kb upstream (away from transcription start site) of the gene.  
(3) Exonic: SVs located in exonic region.  
(4) Intronic: SVs located in intronic region.  
(5) Downstream: SVs located within 1 kb downstream (away from transcription termination site) of the gene region.  
(6) Upstream/Downstream: SVs located within the < 2 kb intergenic region, which is in 1 kb downstream or upstream of the genes.  
(7) Intergenic: SVs located within the > 2 kb intergenic region.  
(8) Splicing: SVs located in the splicing site (2 bp range of the intron/exon boundary).  
(9) Others: SVs located in other region.  
(10) INS: Insersion.  
(11) DEL: Deletion.  
(12) INV: Inversion.  
(13) ITX: Intra-chromosomal translocations.  
(14) CTX: Inter-chromosomal translocations.  
(15) Total: The total number of SVs.  
\*Please note: The statistical data of Upstream、Exonic、Downstream、Intronic、Upstream/Downstream、Intergenic and Others is derived from genome-wide annotation file (\*sv.avinput.variant\_function.gz);The statistical data of INS、DEL、INV、ITX、CTX、Total indicating variant sites number comes from variant detection file (\*.filted.ctx.gz).Additionally, the sum of the number of annotation types possible unequal to the total number of variants as the SV annotation only includes INS, DEL and INV.

DG4222
WRM101
WRM102
WRM103

Figure 4.11 The number of SVs in different regions of the genome

The number of SVs in different regions of the genome.

Figure 4.12 Variation type statistics distribution of SVs

The x-axis represents samples, and the y-axis indicates the proportion of each type of SVs.

---

  

###### 4.6.2 Length Distribution of SVs

The figure 4.13 shows the length distribution of SVs for the random four samples in the sample list. Please refer to the statistical result file (./results/06.SV\_VarDetect/SV.len.rate.xls) for the details of all the samples.

Figure 4.13 Length distribution of SVs

The x-axis represents samples, and the y-axis indicates the proportion of each type of SVs. Note, the length of DNA insert in library construction impacts the SVs detection greatly.

Result：06.SV\_VarDetect

Show Q&A

Q.: How did the SVs detected?

A.: There are four strategies for SV detection: (1) the read-pair technology, with detection of insertional or deletional mutations via aberrant insert size, and inversion via incorrect reading direction; (2) split-read approaches, which detects SVs with uncontinuously mapped reads on different positions of the reference genome; (3) the read-depth method, detecting CNVs caused by insertions and deletions; (4) *de novo* sequence assembly, with detection of SVs by comparison between the assembly and the reference genome. (For review, see *Alkan C, Coe BP, Eichler EE. Genome structural variation discovery and genotyping. Nature reviews Genetics. 2011;12(5):363-376.* from doi:10.1038/nrg2958.) Currently, the SV detection softwares are generally based on one of the principles, the Breakdancer software works with the read-pair method to detect the SVs.

Q.: Can Novogene provide the information of insertional or deletional fragments? Or, how could the SVs be verified?

A.: No, as limited by its read length and analysis, it's yet challenging for even the *de novo* assembling softwares with NGS data. The recommend strategy is to design specific primers and amplify the interested region and check the length variation on agarose gel or subject to Sanger sequencing.

Q.: Can Novogene provide us the flanking sequence adjacent to the mutation for designing primers?

A.: Novogene will provide the variant-related sequences on request.

---

  

##### 4.7 CNV Detection & Annotation

Copy-number variation (CNV) is a type of structural variation showing deletions or duplications in the genome. Based on the reads depth of the reference genome, CNVnator(Abyzov et al., 2011) was used to detect CNVs of potential deletions and duplications with the following parameter '-call 100'. The detected CNVs were further annotated by ANNOVAR(Wang et al., 2010).

Table 4.9 Statistics of CNV detection & annotation

| Sample | Upstream | Exonic | Intronic | Downstream | Upstream/Downstream | Intergenic | Others | Duplication | Deletion | Duplication length (bp) | Deletion length (bp) | Total |
| --- | --- | --- | --- | --- | --- | --- | --- | --- | --- | --- | --- | --- |
| DG4222 | 4 | 49 | 14 | 12 | 2 | 38 | 10 | 57 | 72 | 332900 | 340900 | 129 |
| WRM101 | 6 | 69 | 11 | 13 | 2 | 38 | 11 | 68 | 82 | 537100 | 518400 | 150 |
| WRM102 | 7 | 47 | 21 | 17 | 4 | 42 | 16 | 57 | 97 | 334500 | 296500 | 154 |
| WRM103 | 9 | 54 | 15 | 10 | 3 | 37 | 10 | 68 | 70 | 418200 | 353400 | 138 |

Show Annotation

(1) Sample: Sample names.  
(2) Upstream: CNVs located within 1 kb upstream (away from transcription start site) of the gene.  
(3) Exonic: CNVs located in exonic region.  
(4) Intronic: CNVs located in intronic region.  
(5) Downstream: CNVs located within 1 kb downstream (away from transcription termination site) of the gene region.  
(6) Upstream/Downstream: CNVs located within the < 2 kb intergenic region, which is in 1 kb downstream or upstream of the genes.  
(7) Intergenic: CNVs located within the > 2 kb intergenic region.  
(8) Others: CNVs located in other region.  
(9) Duplication: CNVs with increased copy number.  
(10) Deletion: CNVs with decreased copy number.  
(11) Duplication length (bp): The total length of CNV duplication.  
(12) Deletion length (bp): The total length of CNV deletion.  
(13) Total: The total number of CNVs.  
\*Please note:The statistical data of Upstream、Exonic、Downstream、Intronic、Upstream/Downstream、Intergenic and Others is derived from genome-wide annotation file (\*cnv.avinput.variant\_function.gz) ;The statistical data of Duplication、Deletion、Duplication length (bp)、Deletion length (bp) and Total indicating variant sites number comes from variant detection file (\*.cnv.vcf.gz).

DG4222
WRM101
WRM102
WRM103

Figure 4.14 The number of CNVs in different regions of the genome

The number of CNVs in different regions of the genome.

Figure 4.15 Variation type statistics distribution of CNVs

The x-axis represents samples, and the y-axis indicates the proportion of each type of CNVs.

Result：07.CNV\_VarDetect

Show Q&A

Q.: What is the strategy for CNV detection?

A.: In brief, CNVs are called using read depths of bins, with involving GC calibration and sequencing uniformity calibration.

Q.: What is the original copy number?

A.: For haploid, the original copy number is 1; while for diploid, the original copy number is 2.

---

  

##### 4.8 Visualization of Variation

For proper visualization of the structural variations on the whole genome, we present them according to mutation types with Circos(Krzywinski et al., 2009):

       (1) for SNP/InDel type, the density distribution is drawn.

       (2) for SV/CNV type, the location and size are drawn.

DG4222
WRM101
WRM102
WRM103

Figure 4.16 Whole genome variations distribution

From outer to inner, chromosome, SNP, InDel, CNV duplication, CNV deletion, SV insertion, SV deletion, SV inversion, SV ITX, SV CTX.

DG4222
WRM101
WRM102
WRM103

Figure 4.17 The SNP density

The SNP density on each window was counted with 100kb as the window.

DG4222
WRM101
WRM102
WRM103

Figure 4.18 The InDel density

The InDel density on each window was counted with 100kb as the window.

Result：08.VarDetect\_Visualization

---

  

#### 5 Appendix

##### 5.1 Software

Novogene provides the software list in bioinformatics analysis pipeline for your reference.

Table 5.1 List of software in WGS analyses

| Analysis | Software | Version | Parameter | Remarks |
| --- | --- | --- | --- | --- |
| Quality control | Fastp | 0.20.0 | -g -q 5 -u 50 -n 15 -l 150 | Quality control |
| Mapping | BWA | 0.7.17-r1188 | mem -t 4 -k 32 -M | Mapping clean reads to the reference genome and generation of bam result files. |
|  | SAMtools | 1.13 | sort -@ 6 -m 2G | Sorting the bam files. |
|  | Picard | 1.111 | VALIDATION\_STRINGENCY=SILENT | Merging the bam files from the same sample. |
| SNP/InDel Detection | SAMtools | 1.3.1 | mpileup -m 2 -F 0.002 -d 1000 -u -C 50 | Detection and filtration of SNPs and InDels. |
| SV Detection | Breakdancer | 1.4.4 | -q 20 | SV detection. |
| CNV Detection | CNVnator | V0.3 | -call 100 | CNV detection. |
| Variation Annotation | ANNOVAR | 2015Dec14 | - | Annotation of the detected variations. |

##### 5.2 Methods

Click to open the method(pdf), and the corresponding file can be viewed directly.

File catalog: methods

##### 5.3 Result File Decompression Method and Format Description

  

| Compressed.format | Customer.type | Uncompressed.method |
| --- | --- | --- |
| compressed files in the fomat of \*.tar: | Unix/Linux/Mac user | use tar -xvf \*.tar command |
|  | Windows user | use uncompressed software such as WinRAR, 7-Zip et al |
| compressed files in the format of \*.gz: | Unix/Linux/Mac user | use gzip –d \*.gz command |
|  | Windows user | use uncompressed software such as WinRAR, 7-Zip et al |
| compressed files in the format of \*.zip: | Unix/Linux/Mac user | use unzip \*.zip command |
|  | Windows user | use uncompressed software such as WinRAR, 7-Zip et al |

  
  

| File.type | Document.description | Open.mode |
| --- | --- | --- |
| file.readme.pdf | Results' construction files that help customers to undersdand all the results files better. | Windows/Mac user use Adobe Reader/Foxit reader/web browser to open. |
|  |  | Unix/Linux user use evince command to open. |
| file.fa/fasta/fna | Sequence file, in the fomat of fasta. | Unix/Linux/Mac user use less or more command can view sequences in the format of \*.fasta. |
|  |  | Windows user use editor Editplus/Notepad++ et al. |
| file.fq/fastq | Reads sequence file, in the fomat of fastq. | Unix/Linux/Mac user use less or more command. |
|  |  | Windows user use editor Editplus/Notepad++ et al. |
| file.vcf.gz | This file contains variants in bgzip-compressed VCF (Variant Call Format) format. VCF is a flexible and extendable line-oriented text format developed by the 1000 Genomes Project for releases of SNVs, INDELs, copy number variants and structural variants. | Unix/Linux/Mac user use less or more command. |
| file.pdf/svg | Figure result files; vectorgraph, can be zoom in or out. It is very convenient for customer to view or edit. Using Adobe Illustrator to edit,the picture can be used in paper for publishing. | Windows/Mac user use Adobe Reader/Foxit reader/web browser to open. |
|  |  | Unix/Linux user use evince command to open. |
| file.txt/xls | Table result file; files are separated by(Tab). | Unix/Linux/Mac user use less or more command. |
|  |  | Windows user use editor Editplus/Notepad++ et al, also can use Microsoft Excel to open. |
| file.bam | BAM is compressed format for SAM file which contains sequence alignment information. | Unix/Linux/Mac user use samtools software. |
| file.bai | Index file for bam. | No need to open, put it in the same directory with bam file when manipulate bam. |
| file.gff/gtf | GFF/GTF is general feature format. | Unix/Linux/Mac user use less or more command. |
|  |  | Windows user use editor Editplus/Notepad++ et al. |
| file.png | Image file, bitmap, loseless compression. | Unix/Linux/Mac user use display command. |
|  |  | Windows user use Image browser，such as photoshop et al. |

  

---

  
