## Supplemental Data Set 1 for "The *pos-1* 3ʹ untranslated region governs germline specification and proliferation to ensure reproductive robustness": Reference.README.txt

The detailed information of reference are as follows:

(1) Seq number: the total number of the assembled genomic sequences.
(2) Total length: the total length of the assembled genomic sequence.
(3) GC content: the GC content of the reference genome.
(4) Gap rate: the proportion of unknown sequence (N) in the reference genome assembly.
(5) N50 length: the length of scaffold N50, of which 50% of the sequence is higher than this level.
(6) N90 length: the length of scaffold N90, of which 90% of sequence is higher than this level.
(7) Total length NoN : total Length of Reference Genome Without N Bases.
