## Supplemental Data Set 1 for "The *pos-1* 3ʹ untranslated region governs germline specification and proliferation to ensure reproductive robustness": SAMv1.pdf

### Sequence Alignment/Map Format Specification

The SAM/BAM Format Specification Working Group

28 Feb 2014

The master version of this document can be found at `https://github.com/samtools/hts-specs`. This printing is version 7fd84b0 from that repository, last modified on the date shown above.

#### 1 The SAM Format Specification

SAM stands for Sequence Alignment/Map format. It is a TAB-delimited text format consisting of a header section, which is optional, and an alignment section. If present, the header must be prior to the alignments. Header lines start with '`@`', while alignment lines do not. Each alignment line has 11 mandatory fields for essential alignment information such as mapping position, and variable number of optional fields for flexible or aligner specific information.

##### 1.1 An example

Suppose we have the following alignment with bases in lower cases clipped from the alignment. Read `r001/1` and `r001/2` constitute a read pair; `r003` is a chimeric read; `r004` represents a split alignment.

```
Coor     12345678901234  5678901234567890123456789012345
ref      AGCATGTTAGATAA**GATAGCTGTGCTAGTAGGCAGTCAGCGCCAT

+r001/1        TTAGATAAAGGATA*CTG
+r002         aaaAGATAA*GGATA
+r003       gcctaAGCTAA
+r004                     ATAGCT..............TCAGC
-r003                            ttagctTAGGC
-r001/2                                        CAGCGGCAT
```

The corresponding SAM format is:

```
@HD VN:1.5 SO:coordinate
@SQ SN:ref LN:45
r001  163 ref  7 30 8M2I4M1D3M = 37  39 TTAGATAAAGGATACTG *
r002    0 ref  9 30 3S6M1P1I4M *  0   0 AAAAGATAAGGATA    *
r003    0 ref  9 30 5S6M       *  0   0 GCCTAAGCTAA       * SA:Z:ref,29,-,6H5M,17,0;
r004    0 ref 16 30 6M14N5M    *  0   0 ATAGCTTCAGC       *
r003 2064 ref 29 17 6H5M       *  0   0 TAGGC             * SA:Z:ref,9,+,5S6M,30,1;
r001   83 ref 37 30 9M         =  7 -39 CAGCGGCAT         * NM:i:1
```

#### 1.2 Terminologies and Concepts

**Template** A DNA/RNA sequence part of which is sequenced on a sequencing machine or assembled from raw sequences.

**Segment** A contiguous sequence or subsequence.

**Read** A raw sequence that comes off a sequencing machine. A read may consist of multiple segments. For sequencing data, reads are indexed by the order in which they are sequenced.

**Linear alignment** An alignment of a read to a single reference sequence that may include insertions, deletions, skips and clipping, but may not include direction changes (i.e. one portion of the alignment on forward strand and another portion of alignment on reverse strand). A linear alignment can be represented in a single SAM record.

**Chimeric alignment** An alignment of a read that cannot be represented as a linear alignment. A chimeric alignment is represented as a set of linear alignments that do not have large overlaps. Typically, one of the linear alignments in a chimeric alignment is considered the "representative" alignment, and the others are called "supplementary" and are distinguished by the supplementary alignment flag. All the SAM records in a chimeric alignment have the same `QNAME` and the same values for 0x40 and 0x80 flags (see Section 1.4). The decision regarding which linear alignment is representative is arbitrary.

**Read alignment** A linear alignment or a chimeric alignment that is the complete representation of the alignment of the read.

**Multiple mapping** The correct placement of a read may be ambiguous, e.g. due to repeats. In this case, there may be multiple read alignments for the same read. One of these alignments is considered primary. All the other alignments have the secondary alignment flag set in the SAM records that represent them. All the SAM records have the same `QNAME` and the same values for 0x40 and 0x80 flags. Typically the alignment designated primary is the best alignment, but the decision may be arbitrary.[1]

**1-based coordinate system** A coordinate system where the first base of a sequence is one. In this coordinate system, a region is specified by a closed interval. For example, the region between the 3rd and the 7th bases inclusive is $[3, 7]$. The SAM, VCF, GFF and Wiggle formats are using the 1-based coordinate system.

**0-based coordinate system** A coordinate system where the first base of a sequence is zero. In this coordinate system, a region is specified by a half-closed-half-open interval. For example, the region between the 3rd and the 7th bases inclusive is $[2, 7)$. The BAM, BCFv2, BED, and PSL formats are using the 0-based coordinate system.

**Phred scale** Given a probability $0 < p \leq 1$, the phred scale of $p$ equals $-10 \log_{10} p$, rounded to the closest integer.

#### 1.3 The header section

Each header line begins with character '`@`' followed by a two-letter record type code. In the header, each line is TAB-delimited and except the `@CO` lines, each data field follows a format '`TAG:VALUE`' where `TAG` is a two-letter string that defines the content and the format of `VALUE`. Each header line

[1] A chimeric alignment is primarily caused by structural variations, gene fusions, misassemblies, RNA-seq or experimental protocols. It is more frequent given longer reads. For a chimeric alignment, the linear alignments consisting of the aligment are largely non-overlapping; each linear alignment may have high mapping quality and is informative in SNP/INDEL calling. In contrast, multiple mappings are caused primarily by repeats. They are less frequent given longer reads. If a read has multiple mappings, all these mappings are almost entirely overlapping with each other; except the single-best optimal mapping, all the other mappings get mapping quality <Q3 and are ignored by most SNP/INDEL callers.

should match: `/^@[A-Za-z][A-Za-z](\t[A-Za-z][A-Za-z0-9]:[ -~]+)+$/` or `/^@CO\t.*/`. Tags containing lowercase letters are reserved for end users.

The following table give the defined record types and tags. Tags with '*' are required when the record type is present.

| Tag | Description |
|---|---|
| @HD | The header line. The first line if present. |
| VN* | Format version. *Accepted format*: `/^[0-9]+\.[0-9]+$/`. |
| SO | Sorting order of alignments. *Valid values*: unknown (default), unsorted, queryname and coordinate. For coordinate sort, the major sort key is the RNAME field, with order defined by the order of @SQ lines in the header. The minor sort key is the POS field. For alignments with equal RNAME and POS, order is arbitrary. All alignments with '*' in RNAME field follow alignments with some other value but otherwise are in arbitrary order. |
| @SQ | Reference sequence dictionary. The order of @SQ lines defines the alignment sorting order. |
| SN* | Reference sequence name. Each @SQ line must have a unique SN tag. The value of this field is used in the alignment records in RNAME and PNEXT fields. Regular expression: `[!-)+-<>-~][!-~]*` |
| LN* | Reference sequence length. *Range*: $[1,2^{31}-1]$ |
| AS | Genome assembly identifier. |
| M5 | MD5 checksum of the sequence in the uppercase, excluding spaces but including pads (as '*'s). |
| SP | Species. |
| UR | URI of the sequence. This value may start with one of the standard protocols, e.g http: or ftp:. If it does not start with one of these protocols, it is assumed to be a file-system path. |
| @RG | Read group. Unordered multiple @RG lines are allowed. |
| ID* | Read group identifier. Each @RG line must have a unique ID. The value of ID is used in the RG tags of alignment records. Must be unique among all read groups in header section. Read group IDs may be modified when merging SAM files in order to handle collisions. |
| CN | Name of sequencing center producing the read. |
| DS | Description. |
| DT | Date the run was produced (ISO8601 date or date/time). |
| FO | Flow order. The array of nucleotide bases that correspond to the nucleotides used for each flow of each read. Multi-base flows are encoded in IUPAC format, and non-nucleotide flows by various other characters. *Format*: `/\*\|[ACMGRSVTWYHKDBN]+/` |
| KS | The array of nucleotide bases that correspond to the key sequence of each read. |
| LB | Library. |
| PG | Programs used for processing the read group. |
| PI | Predicted median insert size. |
| PL | Platform/technology used to produce the reads. *Valid values*: CAPILLARY, LS454, ILLUMINA, SOLID, HELICOS, IONTORRENT and PACBIO. |
| PU | Platform unit (e.g. flowcell-barcode.lane for Illumina or slide for SOLiD). Unique identifier. |
| SM | Sample. Use pool name where a pool is being sequenced. |
| @PG | Program. |
| ID* | Program record identifier. Each @PG line must have a unique ID. The value of ID is used in the alignment PG tag and PP tags of other @PG lines. PG IDs may be modified when merging SAM files in order to handle collisions. |
| PN | Program name |
| CL | Command line |
| PP | Previous @PG-ID. Must match another @PG header's ID tag. @PG records may be chained using PP tag, with the last record in the chain having no PP tag. This chain defines the order of programs that have been applied to the alignment. PP values may be modified when merging SAM files in order to handle collisions of PG IDs. The first PG record in a chain (i.e. the one referred to by the PG tag in a SAM record) describes the most recent program that operated on the SAM record. The next PG record in the chain describes the next most recent program that operated on the SAM record. The PG ID on a SAM record is not required to refer to the newest PG record in a chain. It may refer to any PG record in a chain, implying that the SAM record has been operated on by the program in that PG record, and the program(s) referred to via the PP tag. |
| DS | Description. |
| VN | Program version |
| @CO | One-line text comment. Unordered multiple @CO lines are allowed. |

#### 1.4 The alignment section: mandatory fields

In the SAM format, each alignment line typically represents the linear alignment of a segment. Each line has 11 mandatory fields. These fields always appear in the same order and must be present, but their values can be '0' or '*' (depending on the field) if the corresponding information is unavailable. The following table gives an overview of the mandatory fields in the SAM format:

| Col | Field | Type | Regexp/Range | Brief description |
|---|---|---|---|---|
| 1 | QNAME | String | [!-?A-~]{1,255} | Query template NAME |
| 2 | FLAG | Int | [0,$2^{16}$-1] | bitwise FLAG |
| 3 | RNAME | String | *\|[!-()+-<>-~][!-~]* | Reference sequence NAME |
| 4 | POS | Int | [0,$2^{31}$-1] | 1-based leftmost mapping POSition |
| 5 | MAPQ | Int | [0,$2^{8}$-1] | MAPping Quality |
| 6 | CIGAR | String | *\|([0-9]+[MIDNSHPX=])+ | CIGAR string |
| 7 | RNEXT | String | *\|=\|[!-()+-<>-~][!-~]* | Ref. name of the mate/next read |
| 8 | PNEXT | Int | [0,$2^{31}$-1] | Position of the mate/next read |
| 9 | TLEN | Int | [-$2^{31}$+1,$2^{31}$-1] | observed Template LENgth |
| 10 | SEQ | String | *\|[A-Za-z=.]+ | segment SEQuence |
| 11 | QUAL | String | [!-~]+ | ASCII of Phred-scaled base QUALity+33 |

1. QNAME: Query template NAME. Reads/segments having identical QNAME are regarded to come from the same template. A QNAME '*' indicates the information is unavailable. In a SAM file, a read may occupy multiple alignment lines, when its alignment is chimeric or when multiple mappings are given.

2. FLAG: bitwise FLAG. Each bit is explained in the following table:

| Bit | Description |
|---|---|
| 0x1 | template having multiple segments in sequencing |
| 0x2 | each segment properly aligned according to the aligner |
| 0x4 | segment unmapped |
| 0x8 | next segment in the template unmapped |
| 0x10 | SEQ being reverse complemented |
| 0x20 | SEQ of the next segment in the template being reversed |
| 0x40 | the first segment in the template |
| 0x80 | the last segment in the template |
| 0x100 | secondary alignment |
| 0x200 | not passing quality controls |
| 0x400 | PCR or optical duplicate |
| 0x800 | supplementary alignment |

- For each read/contig in a SAM file, it is required that one and only one line associated with the read satisfies 'FLAG & 0x900 == 0'. This line is called the *primary line* of the read.
- Bit 0x100 marks the alignment not to be used in certain analyses when the tools in use are aware of this bit. It is typically used to flag alternative mappings when multiple mappings are presented in a SAM.
- Bit 0x800 indicates that the corresponding alignment line is part of a chimeric alignment. A line flagged with 0x800 is called as a *supplementary line.*
- Bit 0x4 is the only reliable place to tell whether the read is unmapped. If 0x4 is set, no assumptions can be made about RNAME, POS, CIGAR, MAPQ, bits 0x2, 0x10, 0x100 and 0x800, and the bit 0x20 of the previous read in the template.
- If 0x40 and 0x80 are both set, the read is part of a linear template, but it is neither the first nor the last read. If both 0x40 and 0x80 are unset, the index of the read in the template is unknown. This may happen for a non-linear template or the index is lost in data processing.
- If 0x1 is unset, no assumptions can be made about 0x2, 0x8, 0x20, 0x40 and 0x80.

3. RNAME: Reference sequence NAME of the alignment. If @SQ header lines are present, RNAME (if not '*') must be present in one of the SQ-SN tag. An unmapped segment without coordinate has a '*' at this field. However, an unmapped segment may also have an ordinary coordinate such that it can be placed at a desired position after sorting. If RNAME is '*', no assumptions can be made about POS and CIGAR.

4. POS: 1-based leftmost mapping POSition of the first matching base. The first base in a reference sequence has coordinate 1. POS is set as 0 for an unmapped read without coordinate. If POS is 0, no assumptions can be made about RNAME and CIGAR.

5. MAPQ: MAPping Quality. It equals $-10 \log_{10} \Pr\{\text{mapping position is wrong}\}$, rounded to the nearest integer. A value 255 indicates that the mapping quality is not available.

6. CIGAR: CIGAR string. The CIGAR operations are given in the following table (set '*' if unavailable):

| Op | BAM | Description |
|---|---|---|
| M | 0 | alignment match (can be a sequence match or mismatch) |
| I | 1 | insertion to the reference |
| D | 2 | deletion from the reference |
| N | 3 | skipped region from the reference |
| S | 4 | soft clipping (clipped sequences present in SEQ) |
| H | 5 | hard clipping (clipped sequences NOT present in SEQ) |
| P | 6 | padding (silent deletion from padded reference) |
| = | 7 | sequence match |
| X | 8 | sequence mismatch |

   - H can only be present as the first and/or last operation.
   - S may only have H operations between them and the ends of the CIGAR string.
   - For mRNA-to-genome alignment, an N operation represents an intron. For other types of alignments, the interpretation of N is not defined.
   - Sum of lengths of the M/I/S/=/X operations shall equal the length of SEQ.

7. RNEXT: Reference sequence name of the primary alignment of the NEXT read in the template. For the last read, the next read is the first read in the template. If @SQ header lines are present, RNEXT (if not '*' or '=') must be present in one of the SQ-SN tag. This field is set as '*' when the information is unavailable, and set as '=' if RNEXT is identical RNAME. If not '=' and the next read in the template has one primary mapping (see also bit 0x100 in FLAG), this field is identical to RNAME at the primary line of the next read. If RNEXT is '*', no assumptions can be made on PNEXT and bit 0x20.

8. PNEXT: Position of the primary alignment of the NEXT read in the template. Set as 0 when the information is unavailable. This field equals POS at the primary line of the next read. If PNEXT is 0, no assumptions can be made on RNEXT and bit 0x20.

9. TLEN: signed observed Template LENgth. If all segments are mapped to the same reference, the unsigned observed template length equals the number of bases from the leftmost mapped base to the rightmost mapped base. The leftmost segment has a plus sign and the rightmost has a minus sign. The sign of segments in the middle is undefined. It is set as 0 for single-segment template or when the information is unavailable.

10. SEQ: segment SEQuence. This field can be a '*' when the sequence is not stored. If not a '*', the length of the sequence must equal the sum of lengths of M/I/S/=/X operations in CIGAR. An '=' denotes the base is identical to the reference base. No assumptions can be made on the letter cases.

11. QUAL: ASCII of base QUALity plus 33 (same as the quality string in the Sanger FASTQ format). A base quality is the phred-scaled base error probability which equals $-10 \log_{10} \Pr\{\text{base is wrong}\}$. This field can be a '*' when quality is not stored. If not a '*', SEQ must not be a '*' and the length of the quality string ought to equal the length of SEQ.

#### 1.5 The alignment section: optional fields

All optional fields follow the `TAG:TYPE:VALUE` format where `TAG` is a two-character string that matches `/[A-Za-z][A-Za-z0-9]/`. Each `TAG` can only appear once in one alignment line. A `TAG` containing lowercase letters are reserved for end users. In an optional field, `TYPE` is a single case-sensitive letter which defines the format of `VALUE`:

| Type | Regexp matching `VALUE` | Description |
|---|---|---|
| A | `[!-~]` | Printable character |
| i | `[-+]?[0-9]+` | Singed 32-bit integer |
| f | `[-+]?[0-9]*\.?[0-9]+([eE][-+]?[0-9]+)?` | Single-precision floating number |
| Z | `[ !-~]+` | Printable string, including space |
| H | `[0-9A-F]+` | Byte array in the Hex format[2] |
| B | `[cCsSiIf](,[-+]?[0-9]*\.?[0-9]+([eE][-+]?[0-9]+)?)+` | Integer or numeric array |

For an integer or numeric array (type '`B`'), the first letter indicates the type of numbers in the following comma separated array. The letter can be one of '`cCsSiIf`', corresponding to **`int8_t`** (signed 8-bit integer), **`uint8_t`** (unsigned 8-bit integer), **`int16_t`**, **`uint16_t`**, **`int32_t`**, **`uint32_t`** and **`float`**, respectively[3]. During import/export, the element type may be changed if the new type is also compatible with the array.

Predefined tags are shown in the following table. You can freely add new tags, and if a new tag may be of general interest, you can email **``** to add the new tag to the specification. Note that tags starting with '`X`', '`Y`' and '`Z`' or tags containing lowercase letters in either position are reserved for local use and will not be formally defined in any future version of this specification.

| Tag[4] | Type | Description |
|---|---|---|
| `X?` | ? | Reserved fields for end users (together with `Y?` and `Z?`) |
| `AM` | i | The smallest template-independent mapping quality of segments in the rest |
| `AS` | i | Alignment score generated by aligner |
| `BC` | Z | Barcode sequence, with any quality scores stored in the `QT` tag. |
| `BQ` | Z | Offset to base alignment quality (BAQ), of the same length as the read sequence. At the $i$-th read base, $\mathrm{BAQ}_i = Q_i - (\mathrm{BQ}_i - 64)$ where $Q_i$ is the $i$-th base quality. |
| `CC` | Z | Reference name of the next hit; '`=`' for the same chromosome |
| `CM` | i | Edit distance between the color sequence and the color reference (see also `NM`) |
| `CO` | Z | Free-text comments |
| `CP` | i | Leftmost coordinate of the next hit |
| `CQ` | Z | Color read quality on the original strand of the read. Same encoding as QUAL; same length as `CS`. |
| `CS` | Z | Color read sequence on the original strand of the read. The primer base must be included. |
| `CT` | Z | Complete read annotation tag, used for consensus annotation dummy features[5]. |
| `E2` | Z | The 2nd most likely base calls. Same encoding and same length as QUAL. |
| `FI` | i | The index of segment in the template. |
| `FS` | Z | Segment suffix. |
| `FZ` | B,S | Flow signal intensities on the original strand of the read, stored as `(uint16_t) round(value * 100.0)`. |
| `LB` | Z | Library. Value to be consistent with the header `RG-LB` tag if `@RG` is present. |
| `H0` | i | Number of perfect hits |
| `H1` | i | Number of 1-difference hits (see also `NM`) |
| `H2` | i | Number of 2-difference hits |
| `HI` | i | Query hit index, indicating the alignment record is the i-th one stored in SAM |
| `IH` | i | Number of stored alignments in SAM that contains the query in the current record |
| `MC` | Z | CIGAR string for mate/next segment |
| `MD` | Z | String for mismatching positions. *Regex*: `[0-9]+(([A-Z]|\^[A-Z]+)[0-9]+)*`[6] |
| `MQ` | i | Mapping quality of the mate/next segment |
| `NH` | i | Number of reported alignments that contains the query in the current record |
| `NM` | i | Edit distance to the reference, including ambiguous bases but excluding clipping |

[2]For example, a byte array {`0x1a,0xe3,0x1`} corresponds to a Hex string '`1AE301`'.

[3]Explicit typing eases format parsing and helps to reduce the file size when SAM is converted to BAM.

| | | |
|---|---|---|
| OQ | Z | Original base quality (usually before recalibration). Same encoding as QUAL. |
| OP | i | Original mapping position (usually before realignment) |
| OC | Z | Original CIGAR (usually before realignment) |
| PG | Z | Program. Value matches the header PG-ID tag if @PG is present. |
| PQ | i | Phred likelihood of the template, conditional on both the mapping being correct |
| PT | Z | Read annotations for parts of the padded read sequence[7] |
| PU | Z | Platform unit. Value to be consistent with the header RG-PU tag if @RG is present. |
| QT | Z | Phred quality of the barcode sequence in the BC (or RT) tag. Same encoding as QUAL. |
| Q2 | Z | Phred quality of the mate/next segment sequence in the R2 tag. Same encoding as QUAL. |
| R2 | Z | Sequence of the mate/next segment in the template. |
| RG | Z | Read group. Value matches the header RG-ID tag if @RG is present in the header. |
| RT | Z | Deprecated alternative to BC tag originally used at Sanger. |
| SA | Z | Other canonical alignments in a chimeric alignment, in the format of: (*rname*,*pos*,*strand*,*CIGAR*,*mapQ*,*NM*;)+. Each element in the semi-colon delimited list represents a part of the chimeric alignment. Conventionally, at a supplementary line, the first element points to the primary line. |
| SM | i | Template-independent mapping quality |
| TC | i | The number of segments in the template. |
| U2 | Z | Phred probility of the 2nd call being wrong conditional on the best being wrong. The same encoding as QUAL. |
| UQ | i | Phred likelihood of the segment, conditional on the mapping being correct |

[4]The GS, GC, GQ, MF, S2 and SQ are reserved for backward compatibility.

[5]The MD field aims to achieve SNP/indel calling without looking at the reference. For example, a string '10A5^AC6' means from the leftmost reference base in the alignment, there are 10 matches followed by an A on the reference which is different from the aligned read base; the next 5 reference bases are matches followed by a 2bp deletion from the reference; the deleted sequence is AC; the last 6 bases are matches. The MD field ought to match the CIGAR string.

[6]The CT tag is intended primarily for annotation dummy reads, and consists of a *strand*, *type* and zero or more *key*=*value* pairs, each separated with semicolons. The *strand* field has four values as in GFF3, and supplements FLAG bit 0x10 to allow unstranded ('.'), and stranded but unknown strand ('?') annotation. For these and annotation on the forward strand (*strand* set to '+'), do not set FLAG bit 0x10. For annotation on the reverse strand, set the *strand* to '-' and set FLAG bit 0x10. The *type* and any *keys* and their optional *values* are all percent encoded according to RFC3986 to escape meta-characters '=', '%', ';', '|' or non-printable characters not matched by the isprint() macro (with the C locale). For example a percent sign becomes '%2C'. The CT record matches: "*strand*;*type*(;*key*(=*value*))*".

[7]The PT tag value has the format of a series of tags separated by |, each annotating a sub-region of the read. Each tag consists of *start*, *end*, *strand*, *type* and zero or more *key*=*value* pairs, each separated with semicolons. *Start* and *end* are 1-based positions between one and the sum of the M/I/D/P/S/=/X CIGAR operators, i.e. SEQ length plus any pads. Note any editing of the CIGAR string may require updating the 'PT' tag coordinates, or even invalidate them. As in GFF3, *strand* is one of '+' for forward strand tags, '-' for reverse strand, '.' for unstranded or '?' for stranded but unknown strand. The *type* and any *keys* and their optional *values* are all percent encoded as in the CT tag. Formally the entire PT record matches: "*start*;*end*;*strand*;*type*(;*key*(=*value*))*(\|*start*;*end*;*strand*;*type*(;*key*(=*value*))*)*".

#### 2 Recommended Practice for the SAM Format

This section describes the best practice for representing data in the SAM format. They are not required in general, but may be required by a specific software package for it to function properly.

1. The header section
   1 The `@HD` line should be present with the `SO` tag specified.
   2 The `@SQ` lines should be present if reads have been mapped.
   3 When a `RG` tag appears anywhere in the alignment section, there should be a single corresponding `@RG` line with matching `ID` tag in the header.
   4 When a `PG` tag appears anywhere in the alignment section, there should be a single corresponding `@PG` line with matching `ID` tag in the header.
2. Adjacent CIGAR operations should be different.
3. No alignments should be assigned mapping quality 255.
4. Unmapped reads
   1 For a unmapped paired-end or mate-pair read whose mate is mapped, the unmapped read should have RNAME and POS identical to its mate.
   2 If all segments in a template are unmapped, their RNAME should be set as '*' and POS as 0.
   3 If POS plus the sum of lengths of `M/=/X/D/N` operations in CIGAR exceeds the length specified in the `LN` field of the `@SQ` header line (if exists) with an SN equal to RNAME, the alignment should be unmapped.
5. Multiple mapping
   1 When one segment is present in multiple lines to represent a multiple mapping of the segment, only one of these records should have the secondary alignment flag bit (0x100) unset. RNEXT and PNEXT point to the primary line of the next read in the template.
   2 SEQ and QUAL of secondary alignments should be set to '*' to reduce the file size.
6. Optional tags:
   1 If the template has more than 2 segments, the `TC` tag should be present.
   2 The `NM` tag should be present.
7. Annotation dummy reads: These have SEQ set to `*`, FLAG bits 0x100 and 0x200 set (secondary and filtered), and a `CT` tag.
   1 If you wish to store free text in a `CT` tag, use the *key* value `Note` (uppercase N) to match GFF3.
   2 Multi-segment annotation (e.g. a gene with introns) should be described with multiple lines in SAM (like a multi-segment read). Where there is a clear biological direction (e.g. a gene), the first segment (FLAG bit 0x40) is used for the first section (e.g. the 5′ end of the gene). Thus a GenBank entry location like `complement(join(85052..85354, 85441..85621, 6097..86284))` would have three lines in SAM with a common QNAME:
      i. The 5′ fragment FLAG 883, POS 86097, CIGAR `188M`, and tags `FI:i:1` and `TC:i:3`
      ii. Middle fragment FLAG 819, POS 85441, CIGAR `181M`, and tags `FI:i:2` and `TC:i:3`
      iii. The 3′ fragment FLAG 947, POS 85052, CIGAR `303M`, and tags `FI:i:3` and `TC:i:3`

3 If converting GFF3 to SAM, store any *key*, *values* from column 9 in the `CT` tag, except for the unique ID which is used for the QNAME. GFF3 columns 1 (seqid), 4 (start) and 5 (end) are encoded using SAM columns RNAME, POS and CIGAR to hold the length. GFF3 columns 3 (type) and 7 (strand) are stored explicitly in the `CT` tag. Remaining GFF3 columns 2 (source), 6 (score), and 8 (phase) are stored in the `CT` tag using *key* values `FSource`, `FScore` and `FPhase` (uppercase keys are restricted in GFF3, so these names avoid clashes). Split location features are described with multiple lines in GFF3, and similarly become multi-segment dummy reads in SAM, with the RNEXT and PNEXT columns filled in appropriately. In the absence of a convention in SAM/BAM for reads wrapping the origin of a circular genome, any GFF3 feature line wrapping the origin must be split into two segments in SAM.

### 3 Guide for Describing Assembly Sequences in SAM

#### 3.1 Unpadded versus padded representation

To describe alignments, we can regard the reference sequence with no respect to other alignments against it. Such a reference sequence is called an *unpadded reference*. A position on an unpadded reference, referred to as an *unpadded position*, is not affected by any alignments. When we use unpadded references and positions to describe alignments, we say we are using the *unpadded representation*.

Alternatively, to describe the same alignments, we can modify the reference sequence to contain pads that make room for sequences inserted relative to the reference. A pad is effectively a gap and conventionally represented by an asterisk '*'. A reference sequence containing pads is called a *padded reference*. A position which counts the *'s is referred to as a *padded position*. A padded reference sequence may be affected by the query alignments and because of gap insertions is typically longer than the unpadded reference. The padded position of one query alignment may be affected by other query alignments.

Unpadded and padded are different representations of the same alignments. They are convertible to each other with no loss of any information. The unpadded representation is more common due to the convenience of a fixed coordinate system, while the padded representation has the advantage that alignments can be simply described by the start and end coordinates without using complex CIGAR strings. SAM traditionally uses the padded representation for *de novo* assembly. The ACE assembly format uses the padded representation exclusively.

#### 3.2 Padded SAM

The SAM format is typically used to describe alignments against an unpadded reference sequence, but it is also able to describe alignments against a padded reference. In the latter case, we say we are using a *padded SAM*. A padded SAM is a valid SAM, but with the difference that the reference and positions in use are padded. There may be more than one way to describe the padded representation. We recommend the following.

In a padded SAM, alignments and coordinates are described with respect to the padded reference sequence. Unlike traditional padded representations like the ACE file format where pads/gaps are recorded in reads using *'s, we do not write *'s in the SEQ field of the SAM format[1]. Instead, we describe pads in the query sequences as deletions from the padded reference using the CIGAR 'D' operation. In a padded SAM, the insertion and padding CIGAR operations ('I' and 'P') are not used because the padded reference already considers all the insertions.

The following shows the padded SAM for the example alignment in Section 1.1. Notably, the length of `ref` is 47 instead of 45. POS of the last three alignments are all shifted by 2. CIGAR of alignments bridging the 2bp insertion are also changed.

```
@HD VN:1.3 SO:coordinate
@SQ SN:ref LN:47
ref  516 ref  1  0 14M2D31M   *  0   0 AGCATGTTAGATAAGATAGCTGTGCTAGTAGGCAGTCAGCGCCAT *
r001 163 ref  7 30 14M1D3M    = 39  41 TTAGATAAAGGATACTG *
*    768 ref  8 30 1M         *  0   0 *                 *  CT:Z:.;Warning;Note=Ref wrong?
r002   0 ref  9 30 3S6M1D5M   *  0   0 AAAAGATAAGGATA    *  PT:Z:1;4;+;homopolymer
r003   0 ref  9 30 5H6M       *  0   0 AGCTAA            *  NM:i:1
r004   0 ref 18 30 6M14N5M    *  0   0 ATAGCTTCAGC       *
r003  16 ref 31 30 6H5M       *  0   0 TAGGC             *  NM:i:0
r001  83 ref 39 30 9M         =  7 -41 CAGCGCCAT         *
```

Here we also exemplify the recommended practice for storing the reference sequence and the reference annotations in SAM when necessary. For a reference sequence in SAM, QNAME should be

[1]Writing pads/gaps as *'s in the SEQ field might have been more convenient, but this caused concerns for backward compatibility.

identical to RNAME, POS set to 1 and FLAG to 516 (filtered and unmapped); for an annotation, FLAG should be set to 768 (filtered and secondary) with no restriction to QNAME. Dummy reads for annotation would typically have an 'CT' tag to hold the annotation information, see Section 2.

### 4 The BAM Format Specification

#### 4.1 The BGZF compression format

BGZF is block compression implemented on top of the standard gzip file format. The goal of BGZF is to provide good compression while allowing efficient random access to the BAM file for indexed queries. The BGZF format is 'gunzip compatible', in the sense that a compliant gunzip utility can decompress a BGZF compressed file[2].

A BGZF file is a series of concatenated BGZF blocks. Each BGZF block is itself a spec-compliant gzip archive which contains an "extra field" in the format described in RFC1952. The gzip file format allows the inclusion of application-specific extra fields and these are ignored by compliant decompression implementation. The gzip specification also allows gzip files to be concatenated. The result of decompressing concatenated gzip files is the concatenation of the uncompressed data.

Each BGZF block contains a standard gzip file header with the following standard-compliant extensions:

1. The F.EXTRA bit in the header is set to indicate that extra fields are present.
2. The extra field used by BGZF uses the two subfield ID values 66 and 67 (ascii 'BC').
3. The length of the BGZF extra field payload (field LEN in the gzip specification) is 2 (two bytes of payload).
4. The payload of the BGZF extra field is a 16-bit unsigned integer in little endian format. This integer gives the size of the containing BGZF block minus one.

On disk, a complete BGZF file is a series of blocks as shown in the following table. (All integers are little endian as is required by RFC1952.)

| Field | Description | Type | Value |
|---|---|---|---|
| *List of compression blocks (until the end of the file)* | | | |
| ID1 | gzip IDentifier1 | `uint8_t` | 31 |
| ID2 | gzip IDentifier2 | `uint8_t` | 139 |
| CM | gzip Compression Method | `uint8_t` | 8 |
| FLG | gzip FLaGs | `uint8_t` | 4 |
| MTIME | gzip Modification TIME | `uint32_t` | |
| XFL | gzip eXtra FLags | `uint8_t` | |
| OS | gzip Operating System | `uint8_t` | |
| XLEN | gzip eXtra LENgth | `uint16_t` | |
| *Extra subfield(s) (total size=XLEN)* | | | |
| *Additional RFC1952 extra subfields if present* | | | |
| SI1 | Subfield Identifier1 | `uint8_t` | 66 |
| SI2 | Subfield Identifier2 | `uint8_t` | 67 |
| SLEN | Subfield LENgth | `uint16_t` | 2 |
| BSIZE | total Block SIZE minus 1 | `uint16_t` | |
| *Additional RFC1952 extra subfields if present* | | | |
| CDATA | Compressed DATA by zlib::deflate() | `uint8_t`[BSIZE-XLEN-19] | |
| CRC32 | CRC-32 | `uint32_t` | |
| ISIZE | Input SIZE (length of uncompressed data) | `uint32_t` | |

##### 4.1.1 Random access

BGZF files support random access through the BAM file index. To achieve this, the BAM file index uses *virtual file offsets* into the BGZF file. Each virtual file offset is an unsigned 64-bit integer, defined as: `coffset<<16|uoffset`, where `coffset` is an unsigned byte offset into the BGZF file to

[2]It is worth noting that there is a known bug in the Java GZIPInputStream class that concatenated gzip archives cannot be successfully decompressed by this class. BGZF files can be created and manipulated using the built-in Java util.zip package, but naive use of GZIPInputStream on a BGZF file will not work due to this bug.

the beginning of a BGZF block, and `uoffset` is an unsigned byte offset into the uncompressed data stream represented by that BGZF block. Virtual file offsets can be compared, but subtraction between virtual file offsets and addition between a virtual offset and an integer are both disallowed.

###### 4.1.2 End-of-file marker

An end-of-file (EOF) trailer or marker block should be written at the end of BGZF files, so that unintended file truncation can be easily detected. The EOF marker block is a particular empty[3] BGZF block encoded with the default `zlib` compression level setttings, and consists of the following 28 hexadecimal bytes:

```
1f 8b 08 04 00 00 00 00 00 ff 06 00 42 43 02 00 1b 00 03 00 00 00 00 00 00 00 00 00
```

The presence of this EOF marker at the end of a BGZF file indicates that the immediately following physical EOF is the end of the file as intended by the program that wrote it. Empty BGZF blocks are not otherwise special; in particular, the presence of an EOF marker block does not by itself signal end of file.[4]

The absence of this final EOF marker should trigger a warning or error soon after opening a BGZF file where random access is available.[5] When reading a BGZF file in sequential streaming fashion, ideally this EOF check should be performed when the end of the stream is reached. Checking that the final BGZF block in the file decompresses to empty or checking that the last 28 bytes of the file are exactly the bytes above are both sufficent tests; each is likely more convenient in different circumstances.

[3] Empty in the sense of having been formed by compressing a data block of length zero.

[4] An implementation that supports reopening a BAM file in append mode could produce a file by writing headers and alignment records to it, closing it (adding an EOF marker); then reopening it for append, writing more alignment records, and closing it (adding an EOF marker). The resulting BAM file would contain an embedded insignificant EOF marker block that should be effectively ignored when it is read.

[5] It is useful to produce a diagnostic at the beginning of reading a file, so that interactive users can abort lengthy analysis of potentially-corrupted files. Of course, this is only possible if the stream in question supports random access.

#### 4.2 The BAM format

BAM is compressed in the BGZF format. All multi-byte numbers in BAM are little-endian, regardless of the machine endianness. The format is formally described in the following table where values in brackets are the default when the corresponding information is not available; an underlined word in uppercase denotes a field in the SAM format.

<table>
<tr><th>Field</th><th>Description</th><th>Type</th><th>Value</th></tr>
<tr><td>magic</td><td>BAM magic string</td><td>char[4]</td><td>BAM\1</td></tr>
<tr><td>l_text</td><td>Length of the header text, including any NULL padding</td><td>int32_t</td><td></td></tr>
<tr><td>text</td><td>Plain header text in SAM; not necessarily NULL terminated</td><td>char[l_text]</td><td></td></tr>
<tr><td>n_ref</td><td># reference sequences</td><td>int32_t</td><td></td></tr>
<tr><td colspan="4">List of reference information (n=n_ref)</td></tr>
<tr><td>l_name</td><td>Length of the reference name plus 1 (including NULL)</td><td>int32_t</td><td></td></tr>
<tr><td>name</td><td>Reference sequence name; NULL terminated</td><td>char[l_name]</td><td></td></tr>
<tr><td>l_ref</td><td>Length of the reference sequence</td><td>int32_t</td><td></td></tr>
<tr><td colspan="4">List of alignments (until the end of the file)</td></tr>
<tr><td>block_size</td><td>Length of the remainder of the alignment record</td><td>int32_t</td><td></td></tr>
<tr><td>refID</td><td>Reference sequence ID, $-1 \le$ refID $<$ n_ref; -1 for a read without a mapping position.</td><td>int32_t</td><td>[-1]</td></tr>
<tr><td>pos</td><td>0-based leftmost coordinate (= <u>POS</u> $-$ 1)</td><td>int32_t</td><td>[-1]</td></tr>
<tr><td>bin_mq_nl</td><td>bin<<16|<u>MAPQ</u><<8|l_read_name; bin is computed by the reg2bin() function in Section 5.3; l_read_name is the length of read_name below (= length(<u>QNAME</u>) + 1).</td><td>uint32_t</td><td></td></tr>
<tr><td>flag_nc</td><td><u>FLAG</u><<16|n_cigar_op; n_cigar_op is the number of operations in <u>CIGAR</u>.</td><td>uint32_t</td><td></td></tr>
<tr><td>l_seq</td><td>Length of <u>SEQ</u></td><td>int32_t</td><td></td></tr>
<tr><td>next_refID</td><td>Ref-ID of the next segment ($-1 \le$ mate_refID $<$ n_ref)</td><td>int32_t</td><td>[-1]</td></tr>
<tr><td>next_pos</td><td>0-based leftmost pos of the next segment (= <u>PNEXT</u> $-$ 1)</td><td>int32_t</td><td>[-1]</td></tr>
<tr><td>tlen</td><td>Template length (= <u>TLEN</u>)</td><td>int32_t</td><td>[0]</td></tr>
<tr><td>read_name</td><td>Read name[1], NULL terminated (<u>QNAME</u> plus a tailing '\0')</td><td>char[l_read_name]</td><td></td></tr>
<tr><td>cigar</td><td>CIGAR: op_len<<4|op. 'MIDNSHP=X'→'012345678'</td><td>uint32_t[n_cigar_op]</td><td></td></tr>
<tr><td>seq</td><td>4-bit encoded read: '=ACMGRSVTWYHKDBN'→ $[0, 15]$; other characters mapped to 'N'; high nybble first (1st base in the highest 4-bit of the 1st byte)</td><td>uint8_t[(l_seq+1)/2]</td><td></td></tr>
<tr><td>qual</td><td>Phred base quality (a sequence of 0xFF if absent)</td><td>char[l_seq]</td><td></td></tr>
<tr><td colspan="4">List of auxiliary data (until the end of the alignment block)</td></tr>
<tr><td>tag</td><td>Two-character tag</td><td>char[2]</td><td></td></tr>
<tr><td>val_type</td><td>Value type: AcCsSiIfZHB[2,3]</td><td>char</td><td></td></tr>
<tr><td>value</td><td>Tag value</td><td>(by val_type)</td><td></td></tr>
</table>

[1]For backward compatibility, a QNAME '*' is stored as a C string "*\0".

[2]An integer may be stored as one of 'cCsSiI' in BAM, representing int8_t, uint8_t, int16_t, uint16_t, int32_t and uint32_t, respectively. In SAM, all single integer types are mapped to int32_t.

[3]A 'B'-typed (array) tag–value pair is stored as follows. The first two bytes keep the two-character tag. The 3rd byte is always 'B'. The 4th byte, matching /^[cCsSiIf]$/, indicates the type of an element in the array. Bytes from 5 to 8 encode a little-endian 32-bit integer which gives the number of elements in the array. Bytes starting from the 9th store the array in the little-endian byte order; the number of these bytes is determined by the type and the length of the array.

### 5 Indexing BAM

Indexing aims to achieve fast retrieval of alignments overlapping a specified region without going through the whole alignments. BAM must be sorted by the reference ID and then the leftmost coordinate before indexing.

#### 5.1 Algorithm

##### 5.1.1 Basic binning index

The UCSC binning scheme was suggested by Richard Durbin and Lincoln Stein and is explained by Kent et al. (2002). In this scheme, each bin represents a contiguous genomic region which is either fully contained in or non-overlapping with another bin; each alignment is associated with a bin which represents the smallest region containing the entire alignment. The binning scheme is essentially a representation of R-tree. A distinct bin uniquely corresponds to a distinct internal node in a R-tree. Bin A is a child of Bin B if the region represented by A is contained in B.

To find the alignments that overlap a specified region, we need to get the bins that overlap the region, and then test each alignment in the bins to check overlap. To quickly find alignments associated with a specified bin, we can keep in the index the start file offsets of chunks of alignments which all have the bin. As alignments are sorted by the leftmost coordinates, alignments having the same bin tend to be clustered together on the disk and therefore usually a bin is only associated with a few chunks. Traversing all the alignments having the same bin usually needs a few seek calls. Given the set of bins that overlap the specified region, we can visit alignments in the order of their leftmost coordinates and stop seeking the rest when an alignment falls outside the required region. This strategy saves half of the seek calls in average.

In BAM, each bin may span $2^{29}$, $2^{26}$, $2^{23}$, $2^{20}$, $2^{17}$ or $2^{14}$ bp[6]. Bin 0 spans a 512Mbp region, bins 1–8 span 64Mbp, 9–72 8Mbp, 73–584 1Mbp, 585–4680 128Kbp and bins 4681–37449 span 16Kbp regions.

##### 5.1.2 Reducing small chunks

Around the boundary of two adjacent bins, we may see many small chunks with some having a shorter bin while the rest having a larger bin. To reduce the number of seek calls, we may join two chunks having the same bin if they are close to each other. After this process, a joined chunk will contain alignments with different bins. We need to keep in the index the file offset of the end of each chunk to identify its boundaries.

##### 5.1.3 Combining with linear index

For an alignment starting beyond 64Mbp, we always need to seek to some chunks in bin 0, which can be avoided by using a linear index. In the linear index, for each tiling 16384bp window on the reference, we record the smallest file offset of the alignments that start in the window. Given a region [rbeg,rend), we only need to visit a chunk whose end file offset is larger than the file offset of the 16kbp window containing rbeg.

With both binning and linear indices, we can retrieve alignments in most of regions with just one seek call.

##### 5.1.4 A conceptual example

Suppose we have a genome shorter than 144kbp. we can design a binning scheme which consists of three types of bins: bin 0 spans 0-144kbp, bin 1, 2 and 3 span 48kbp and bins from 4 to 12 span 16kbp each:

An alignment starting at 65kbp and ending at 67kbp would have a bin number 8, which is the smallest bin containing the alignment. Similarly, an alignment starting at 51kbp and ending at 70kbp

[6] Due to a limitation in the current indexing scheme, a chromosome sequence longer than $2^{29} - 1$ is not supported during indexing.

| | | | | | | | | |
|---|---|---|---|---|---|---|---|---|
| 0 (0–144kbp) | | | | | | | | |
| 1 (0–48kbp) | | | 2 (48–96kbp) | | | 1 (96–144kbp) | | |
| 4 (0–16k) | 5 (16–32k) | 6 (32–48k) | 7 (48–64k) | 8 (64–80k) | 9 (80–96k) | 10 | 11 | 12 |

would go to bin 2, while an alignment between [40k,49k] to bin 0. Suppose we want to find all the alignments overlapping region [65k,71k). We first calculate that bin 0, 2 and 8 overlap with this region and then traverse the alignments in these bins to find the required alignments. With a binning index alone, we need to visit the alignment at [40k,49k] as it belongs to bin 0. But with a linear index, we know that such an alignment stops before 64kbp and cannot overlap the specified region. A seek call can thus be saved.

#### 5.2 The BAM indexing format

| Field | Description | Type | Value |
|---|---|---|---|
| magic | Magic string | `char[4]` | `BAI\1` |
| n_ref | # reference sequences | `int32_t` | |
| *List of indices (n=n_ref)* | | | |
| n_bin | # distinct bins (for the binning index) | `int32_t` | |
| *List of distinct bins (n=n_bin)* | | | |
| bin | Distinct bin | `uint32_t` | |
| n_chunk | # chunks | `int32_t` | |
| *List of chunks (n=n_chunk)* | | | |
| chunk_beg | (Virtual) file offset of the start of the chunk | `uint64_t` | |
| chunk_end | (Virtual) file offset of the end of the chunk | `uint64_t` | |
| n_intv | # 16kbp intervals (for the linear index) | `int32_t` | |
| *List of intervals (n=n_intv)* | | | |
| ioffset | (Virtual) file offset of the first alignment in the interval | `uint64_t` | |

#### 5.3 C source code for computing bin number and overlapping bins

```
/* calculate bin given an alignment covering [beg,end) (zero-based, half-close-half-open) */
int reg2bin(int beg, int end)
{
    --end;
    if (beg>>14 == end>>14) return ((1<<15)-1)/7 + (beg>>14);
    if (beg>>17 == end>>17) return ((1<<12)-1)/7 + (beg>>17);
    if (beg>>20 == end>>20) return ((1<<9)-1)/7  + (beg>>20);
    if (beg>>23 == end>>23) return ((1<<6)-1)/7  + (beg>>23);
    if (beg>>26 == end>>26) return ((1<<3)-1)/7  + (beg>>26);
    return 0;
}
/* calculate the list of bins that may overlap with region [beg,end) (zero-based) */
#define MAX_BIN (((1<<18)-1)/7)
int reg2bins(int beg, int end, uint16_t list[MAX_BIN])
{
    int i = 0, k;
    --end;
    list[i++] = 0;
    for (k =    1 + (beg>>26); k <=    1 + (end>>26); ++k) list[i++] = k;
    for (k =    9 + (beg>>23); k <=    9 + (end>>23); ++k) list[i++] = k;
    for (k =   73 + (beg>>20); k <=   73 + (end>>20); ++k) list[i++] = k;
    for (k =  585 + (beg>>17); k <=  585 + (end>>17); ++k) list[i++] = k;
    for (k = 4681 + (beg>>14); k <= 4681 + (end>>14); ++k) list[i++] = k;
    return i;
}
```
