## Supplemental Data Set 1 for "The *pos-1* 3ʹ untranslated region governs germline specification and proliferation to ensure reproductive robustness": Readme.txt

Supplemental Data Set 1
Varderesian et al., 2025

File:
AF1_Exonic_Indels_SNPs.xlsx (Microsoft Office)

This file contains exonic SNPs and Indels mapped in strain DG4222, WRM101, and WRM102 against the reference genome ce11 (WBcel35). It also contains a comparison of the three strains and was used to make Supplemental Figure 1.

Folder:
03.Result_X202SC24112711-Z01-F001_C_elegans

This folder contains the complete report and results from Novogene's whole genome sequencing service, including quality control, mapping statistics, SNP detection, Indel detection, SV detection, and CNV detection for DG4222, WRM101, and WRM102. It also contains visualizations and a result tree. Please note that strain WRM103 was not used in this work (it's relevant to a different project). As such, all data for WRM103 was removed from this folder, but may be referenced in the pre-compiled reports.
