## Supplementary figures and images for "The *pos-1* 3ʹ untranslated region governs germline specification and proliferation to ensure reproductive robustness"

### DG4222.Circos.png

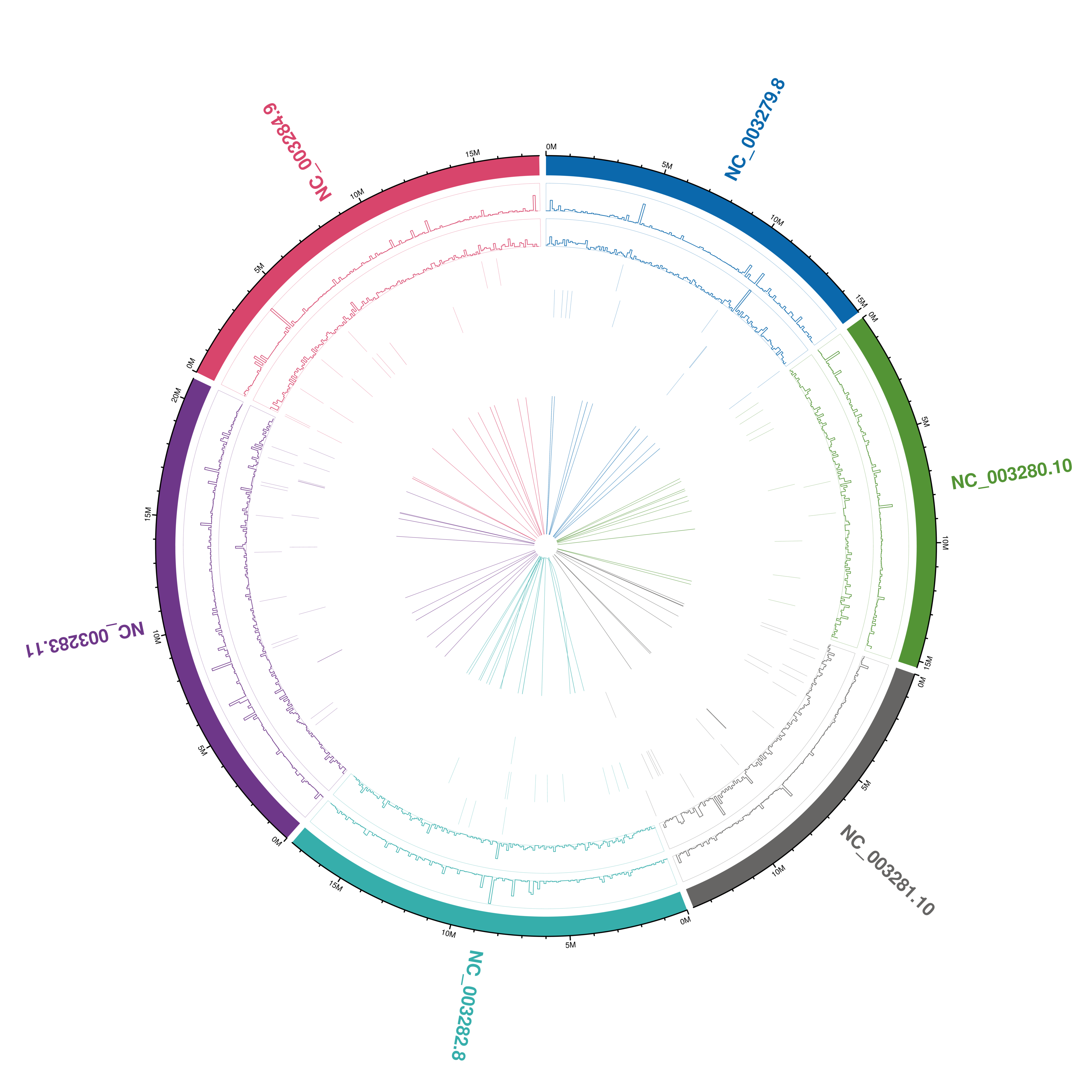

### DG4222.indDensity.pdf

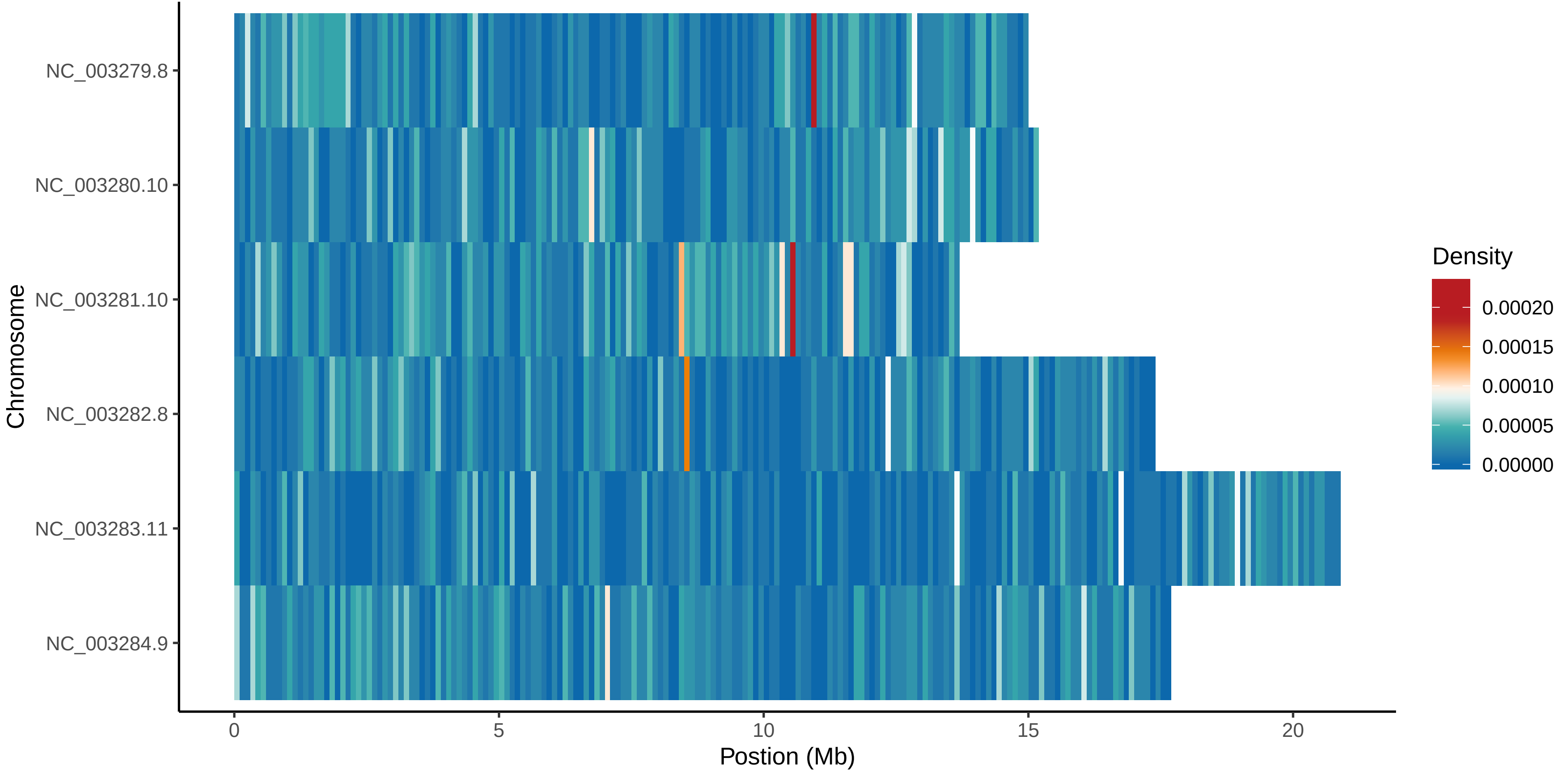

### DG4222.mapbychrdepth.pdf

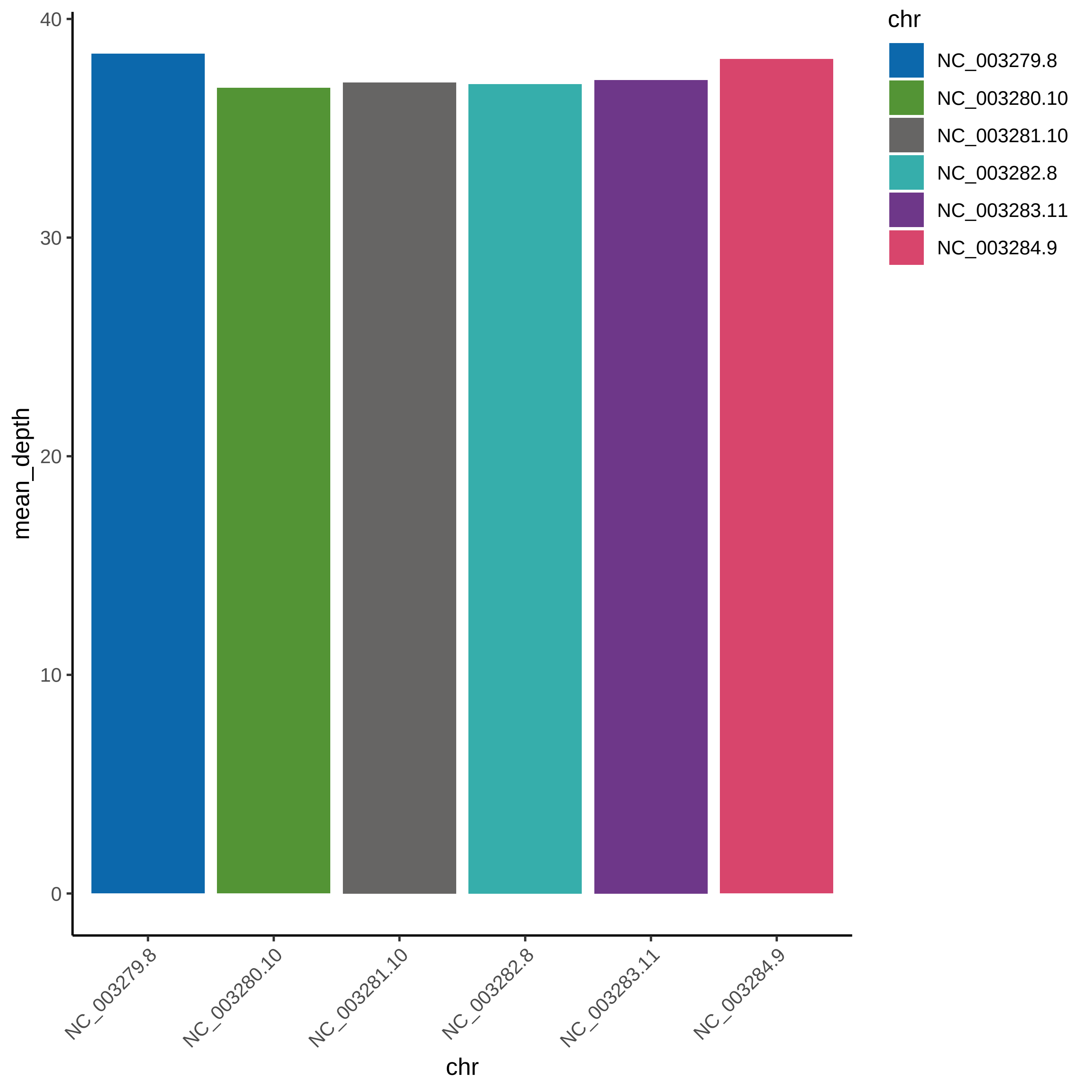

### DG4222.mapbychrdepth.png

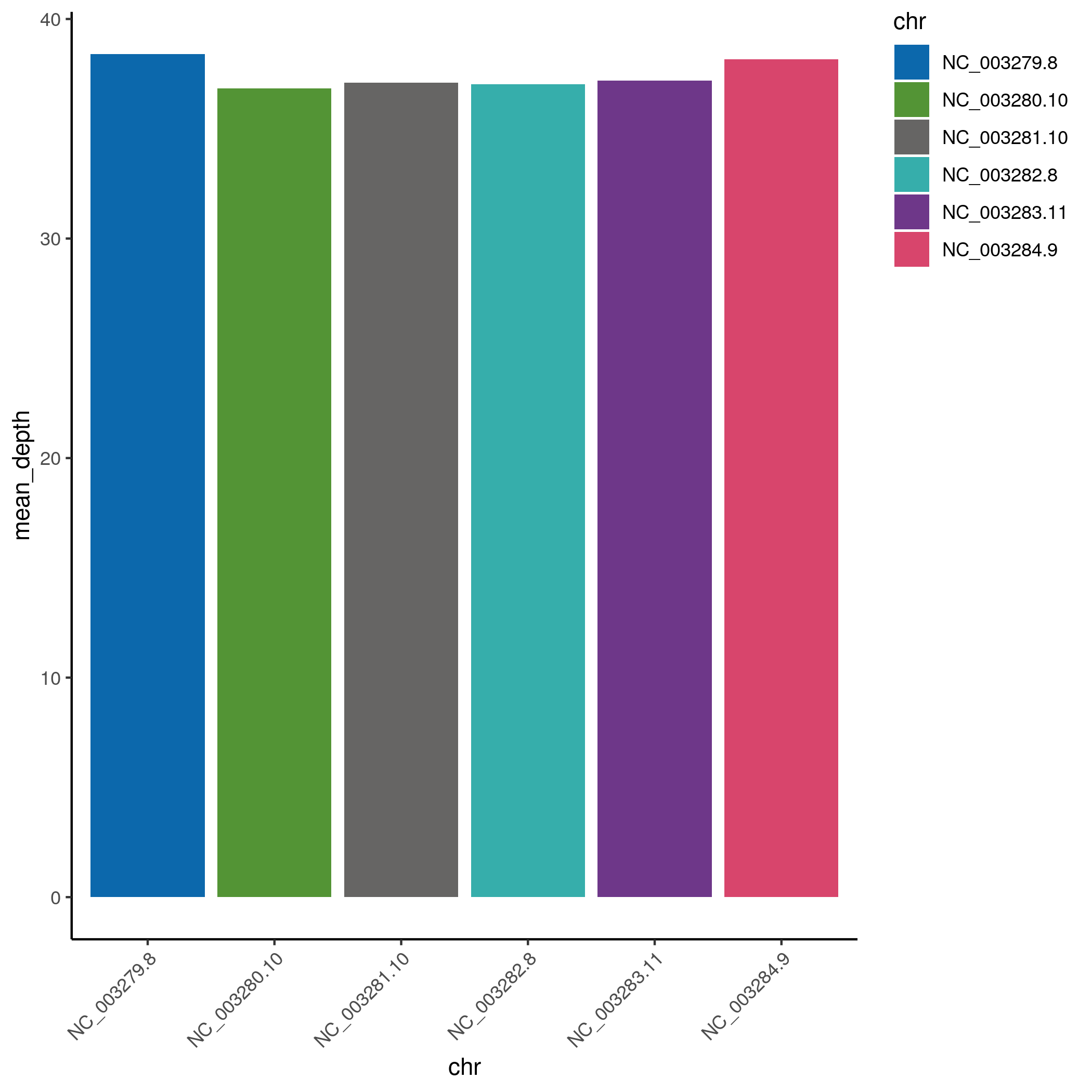

### sort_asc.png

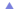

### WRM101.mapbychrdepth.pdf

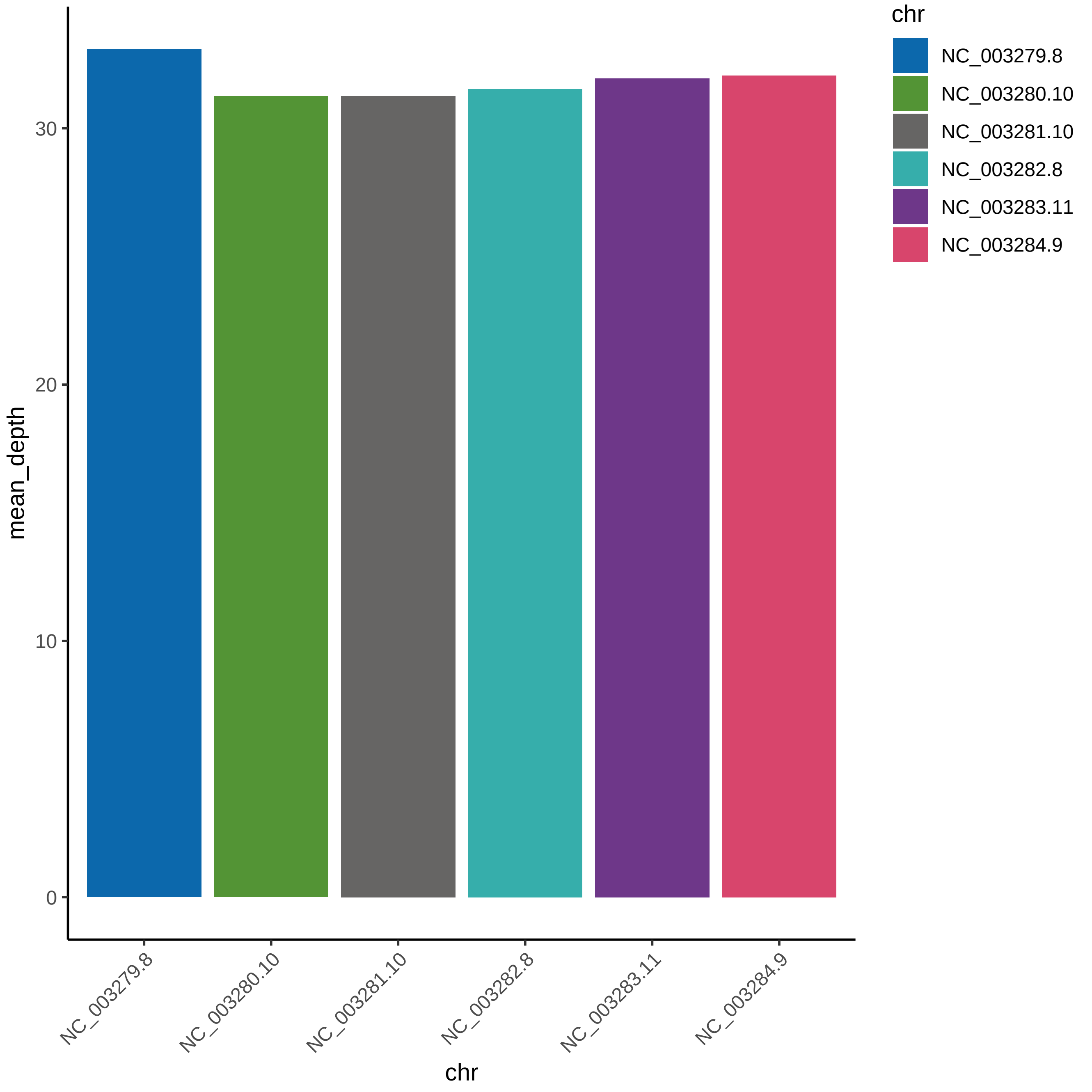

### WRM101.mapbychrdepth.png

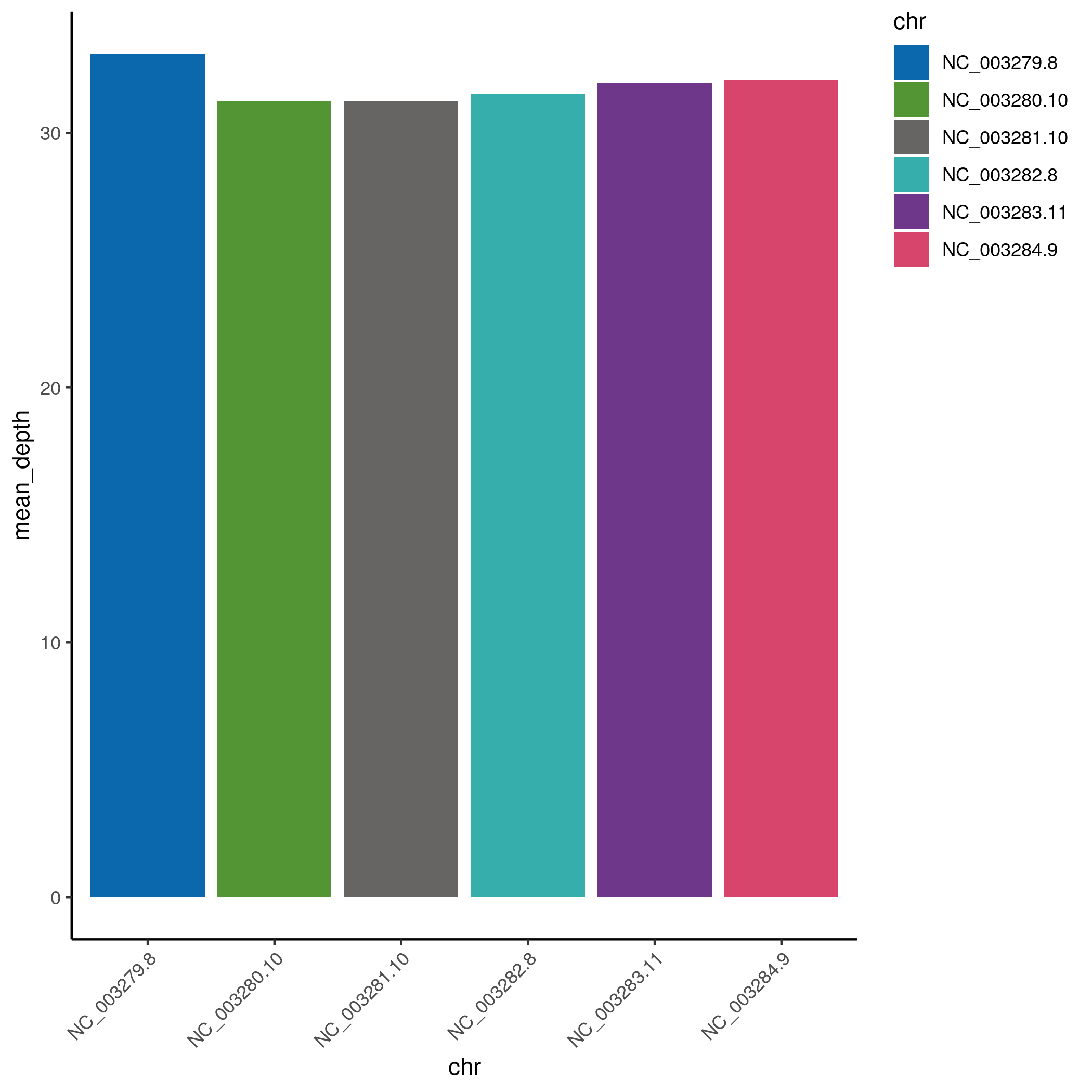

### WRM101.snpDensity.pdf

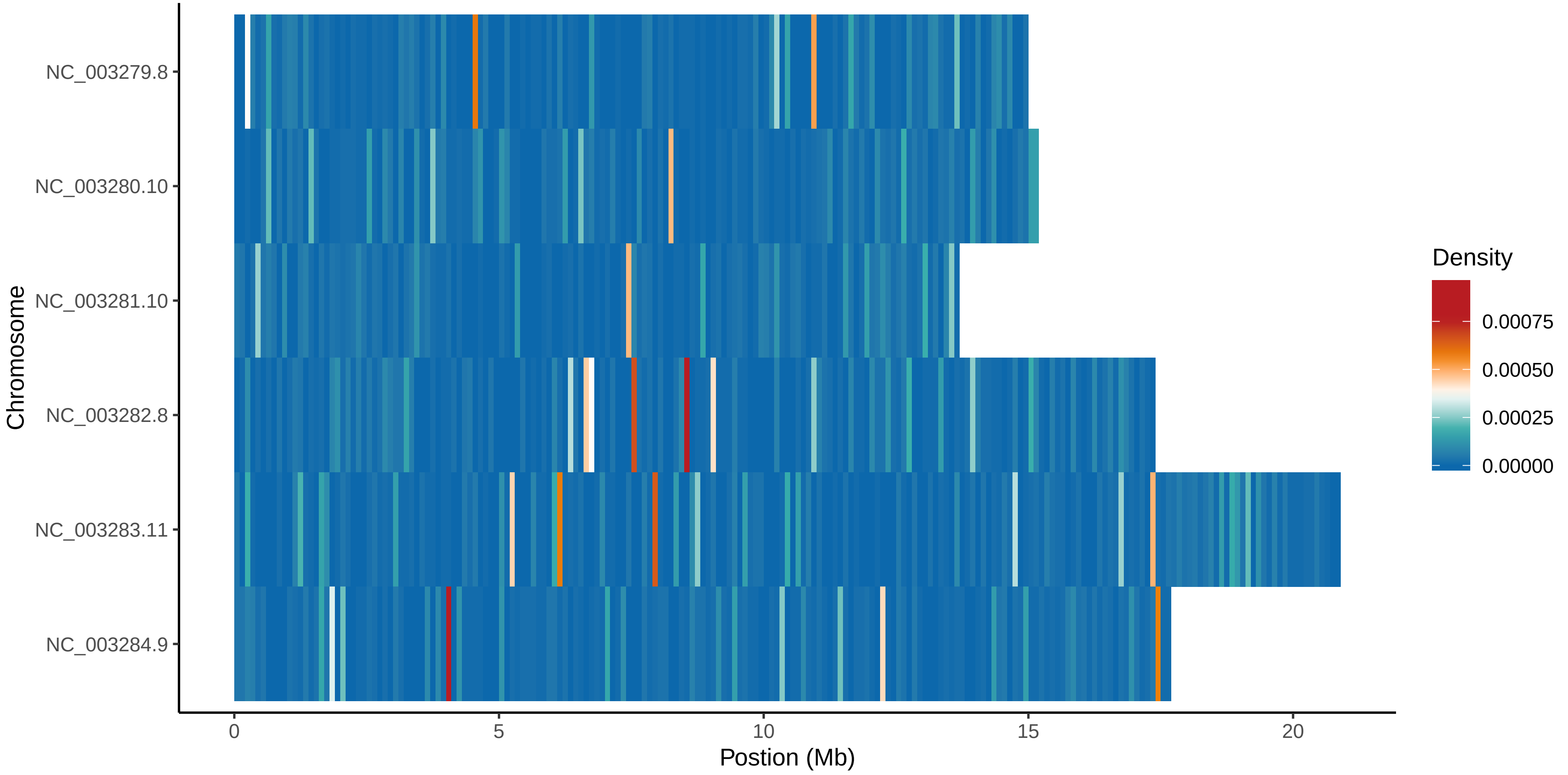

### WRM102.Circos.pdf

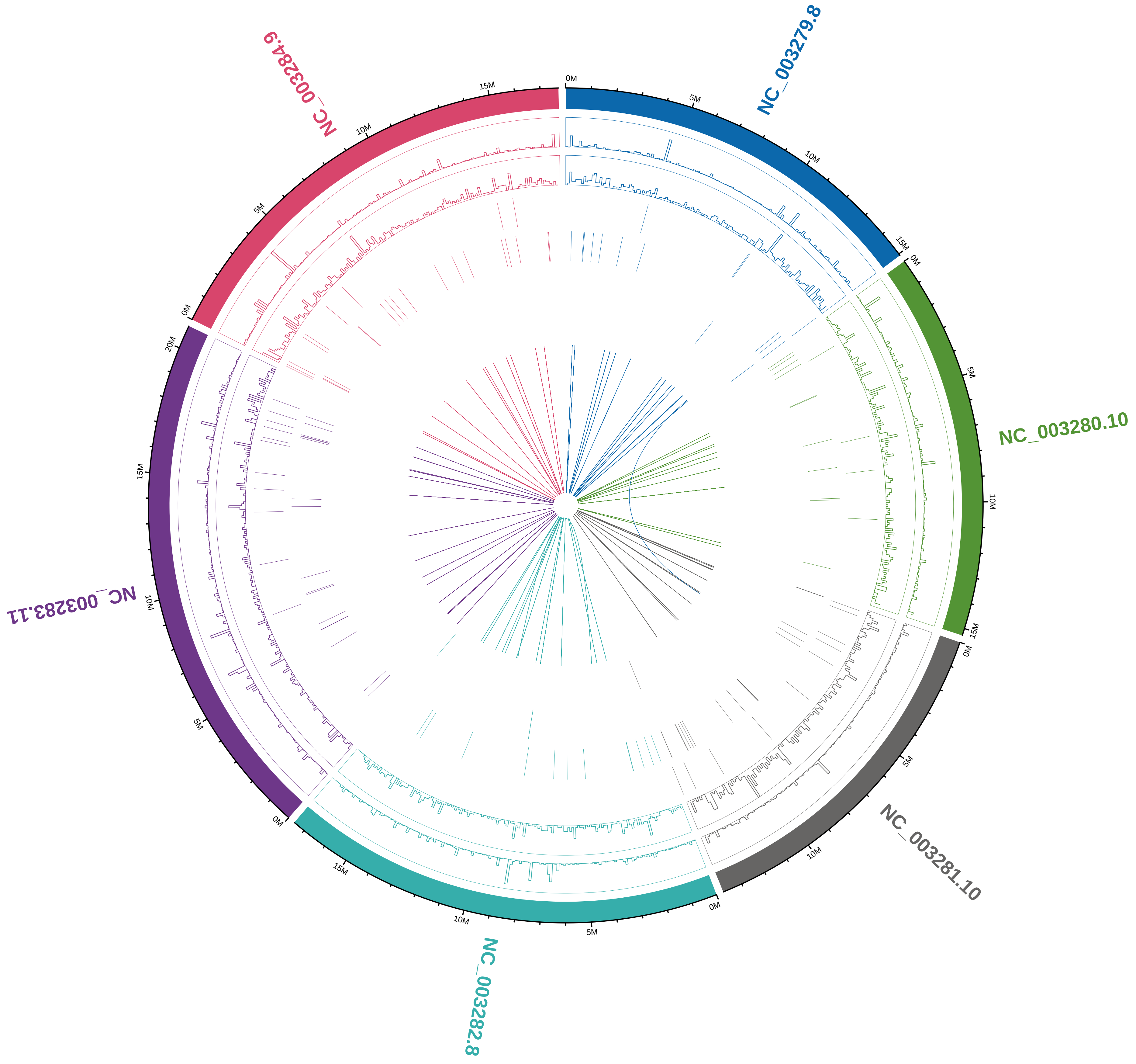

### WRM102.Circos.png

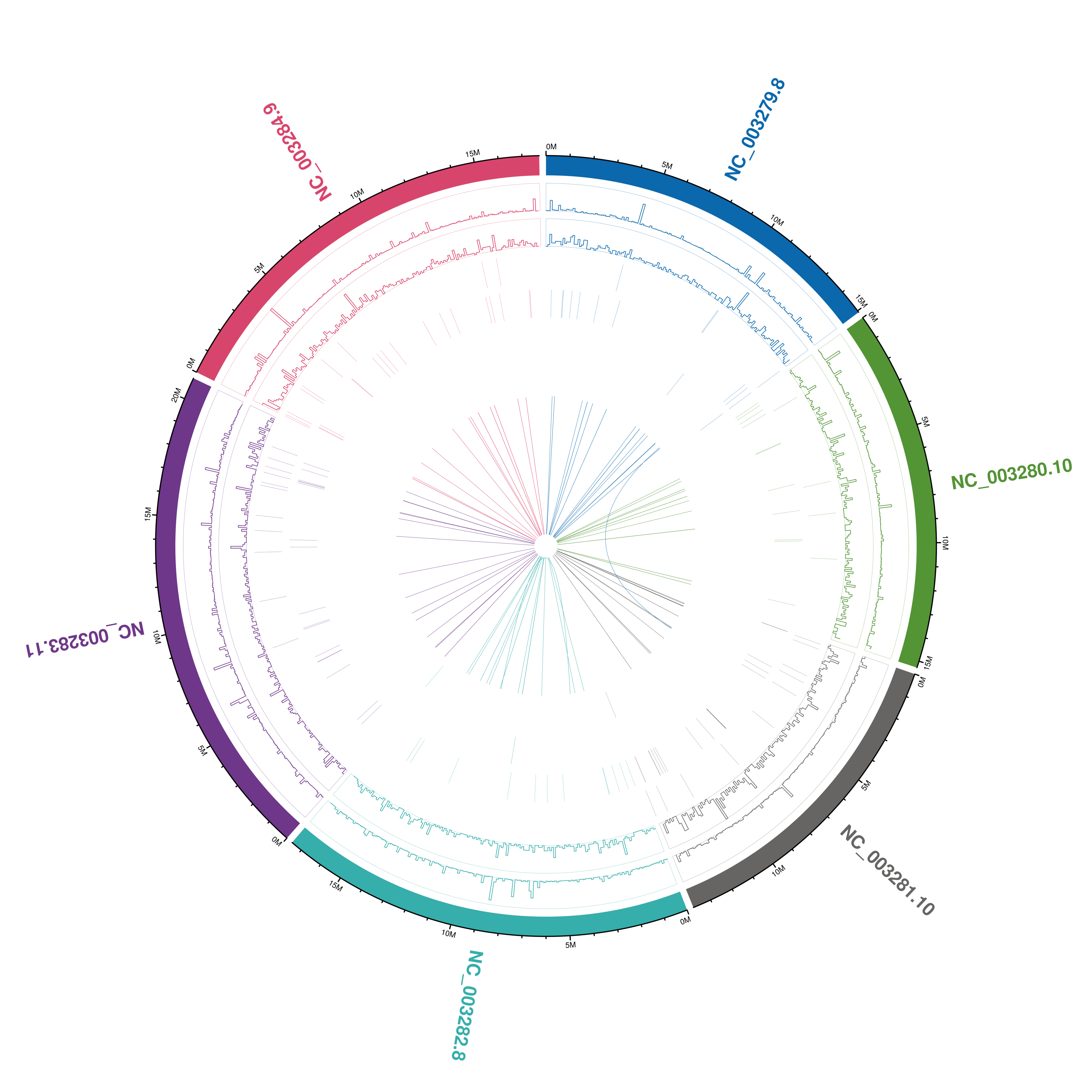

### WRM102.mapbychrdepth.pdf

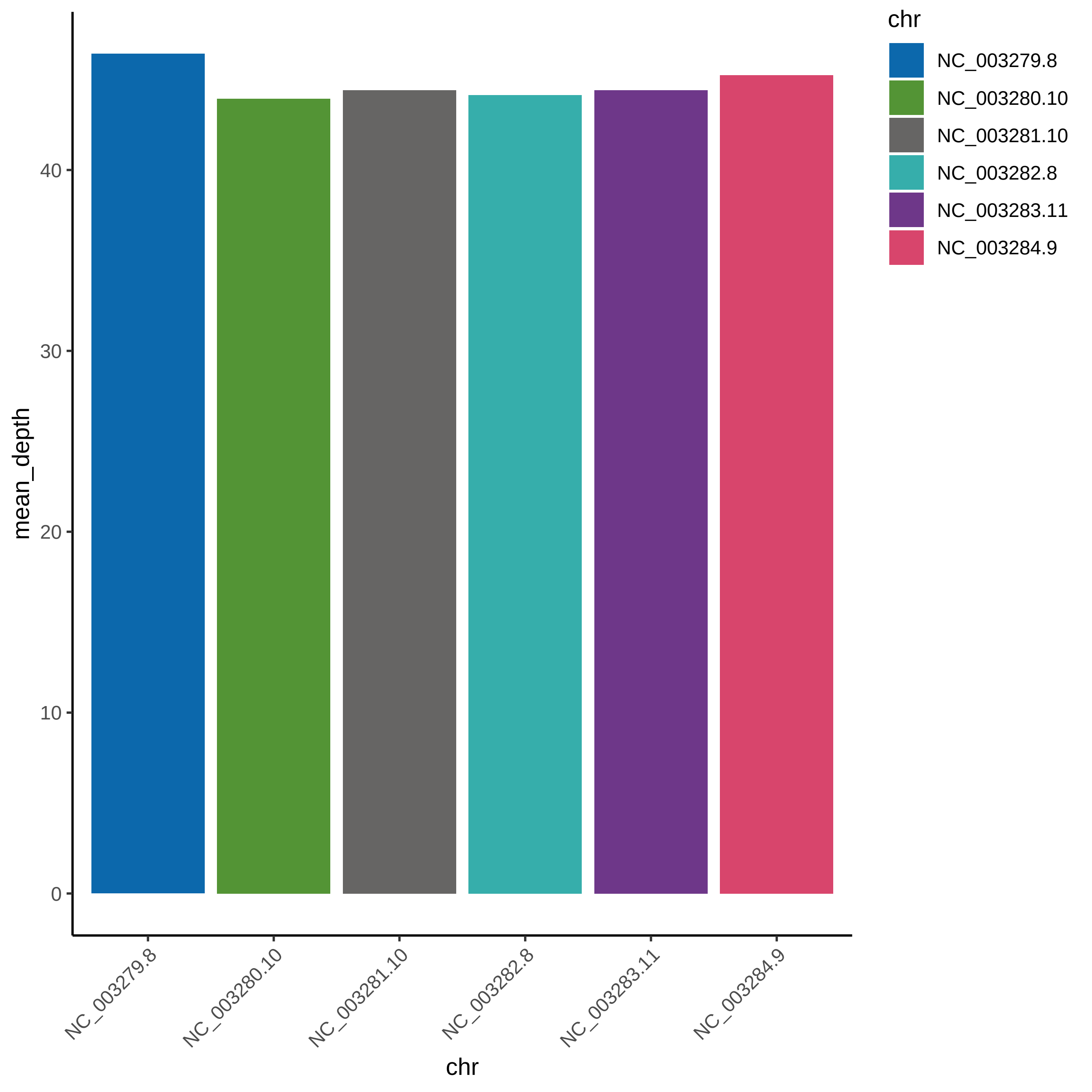

### WRM102.mapbychrdepth.png

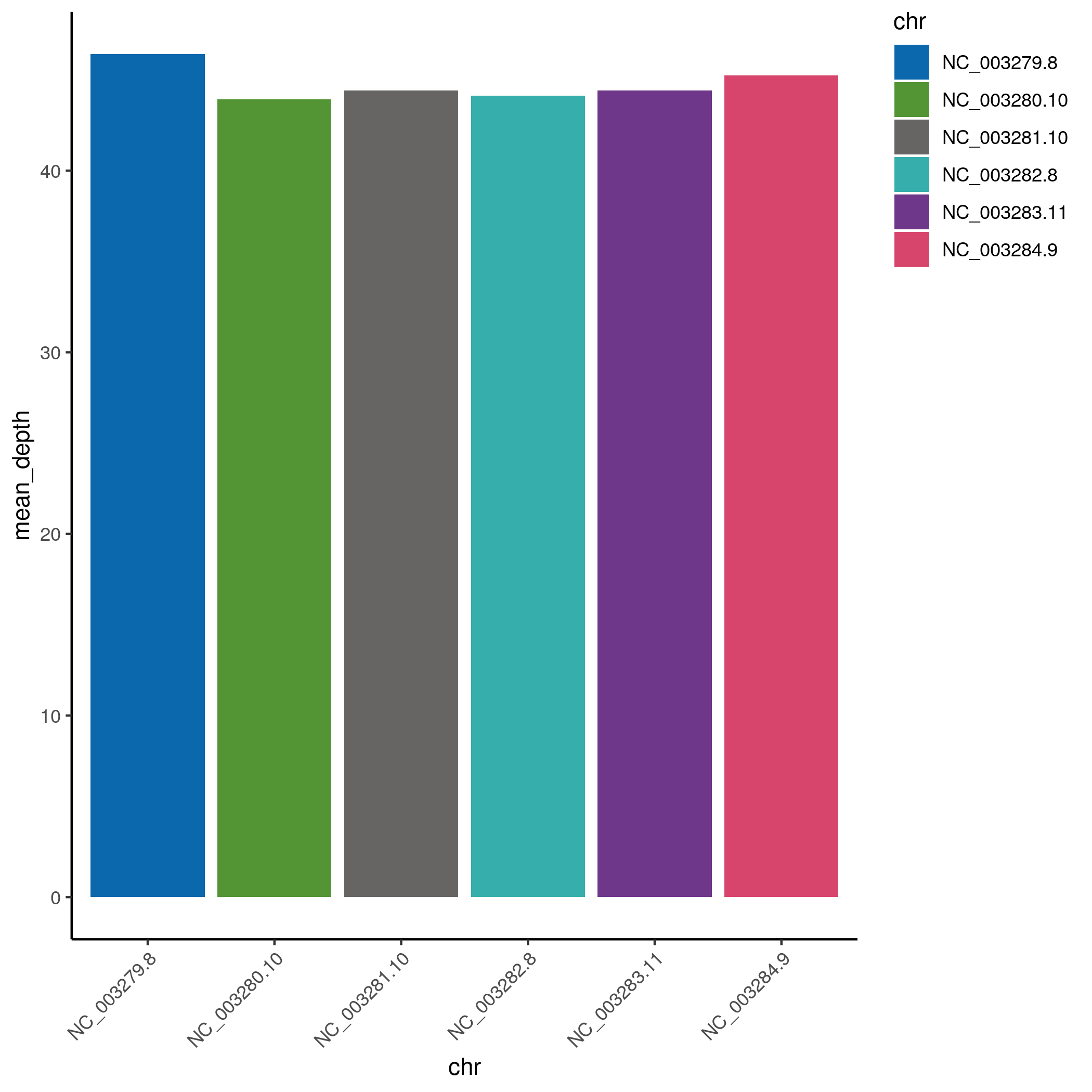

### WRM102.snpDensity.pdf

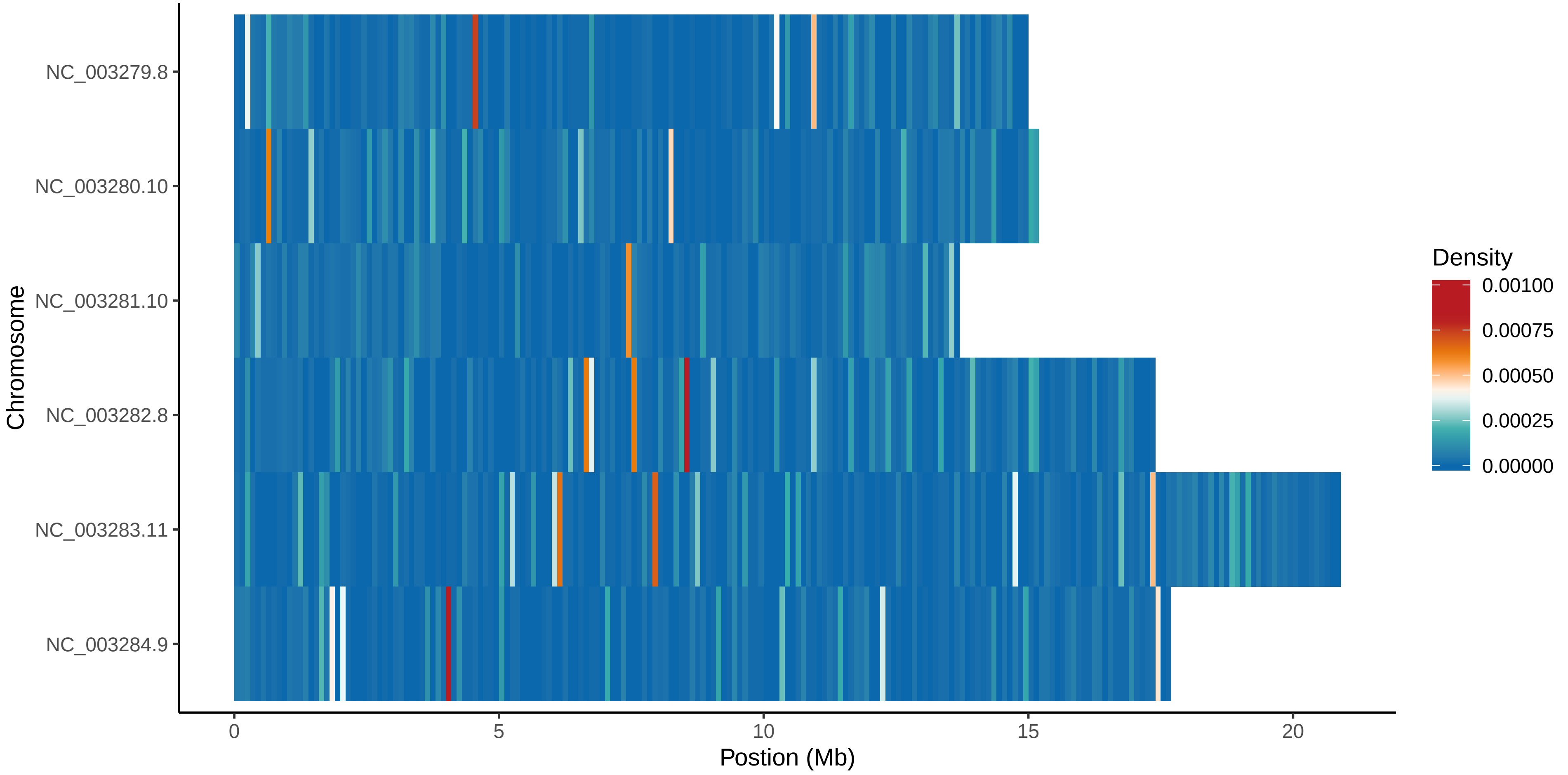

### WRM103.mapbychrdepth.pdf

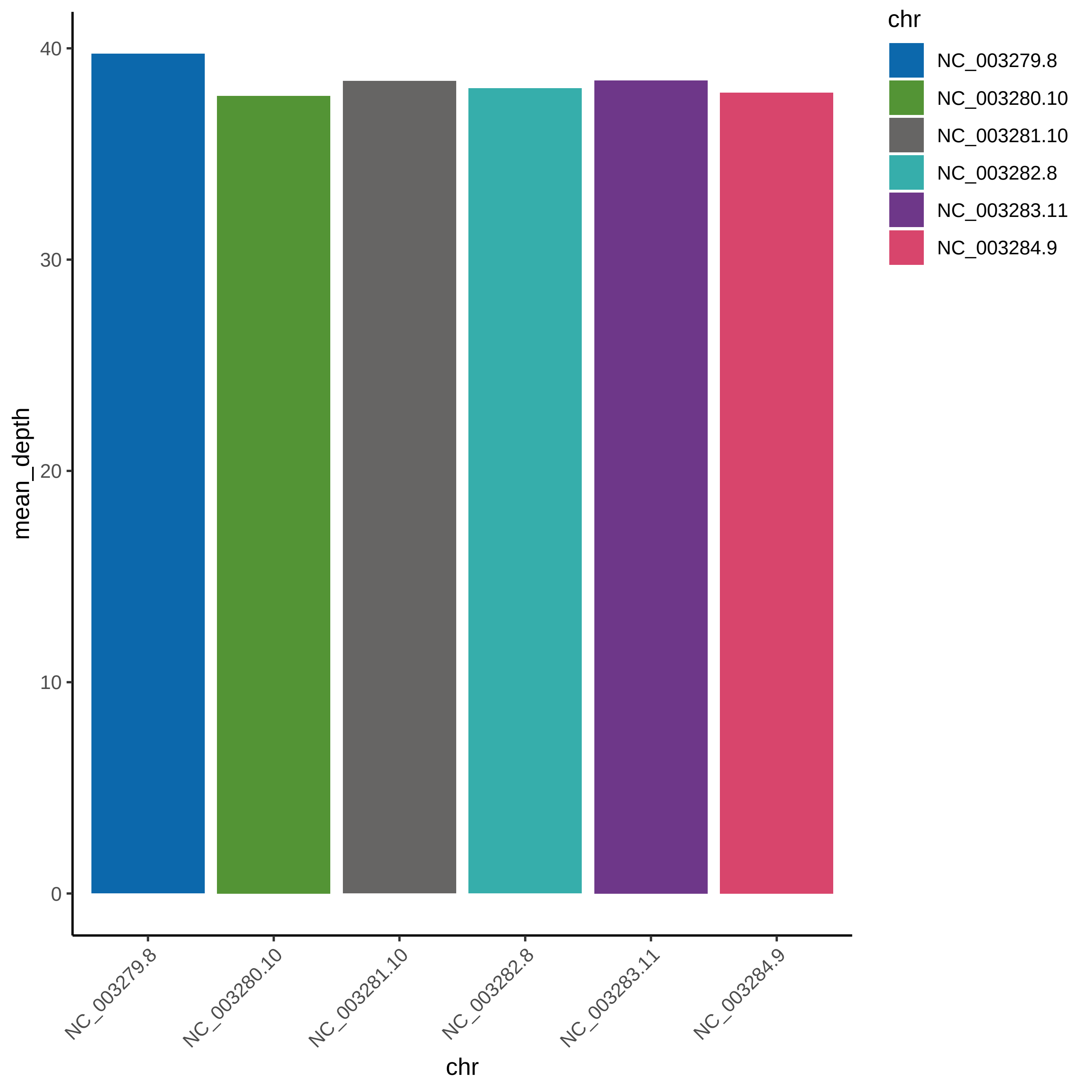

### WRM103.mapbychrdepth.png

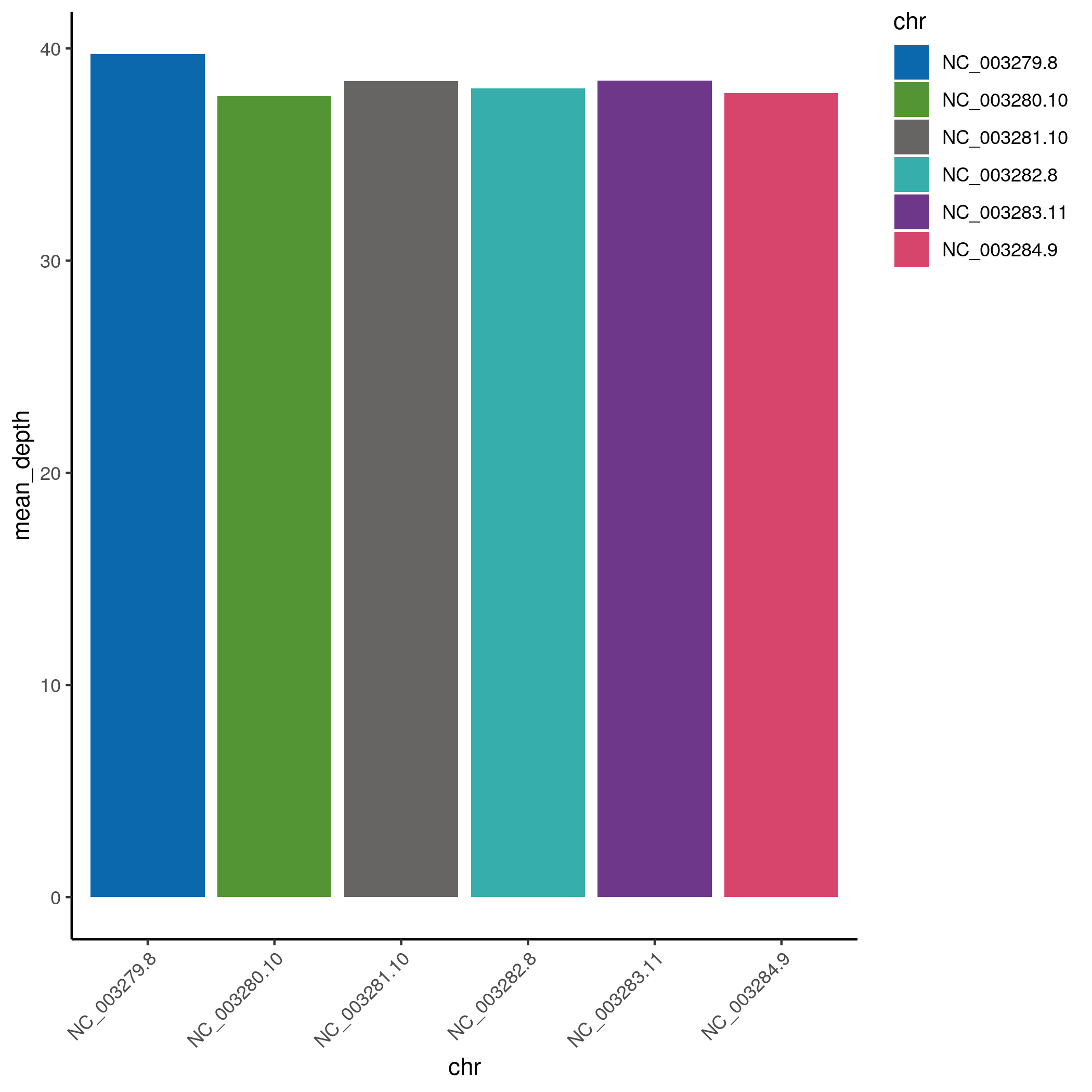

### WRM103.snpDensity.pdf

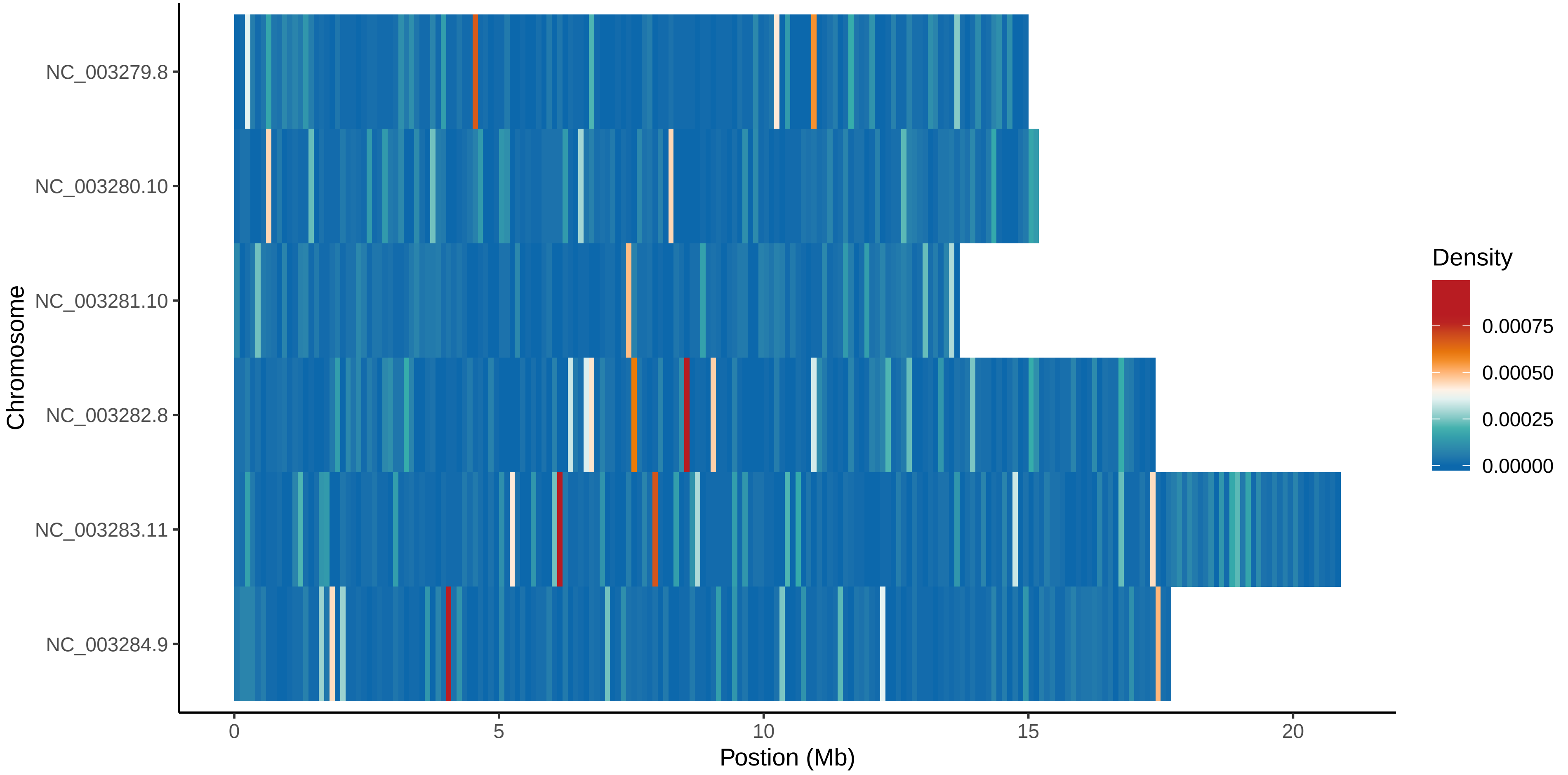
