## Supplementary figures and images for "The *pos-1* 3ʹ untranslated region governs germline specification and proliferation to ensure reproductive robustness"

### album-slider-arrow_box.png

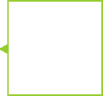

### album-slider-button.png

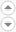

### bioinfoWorkflow.adv.png

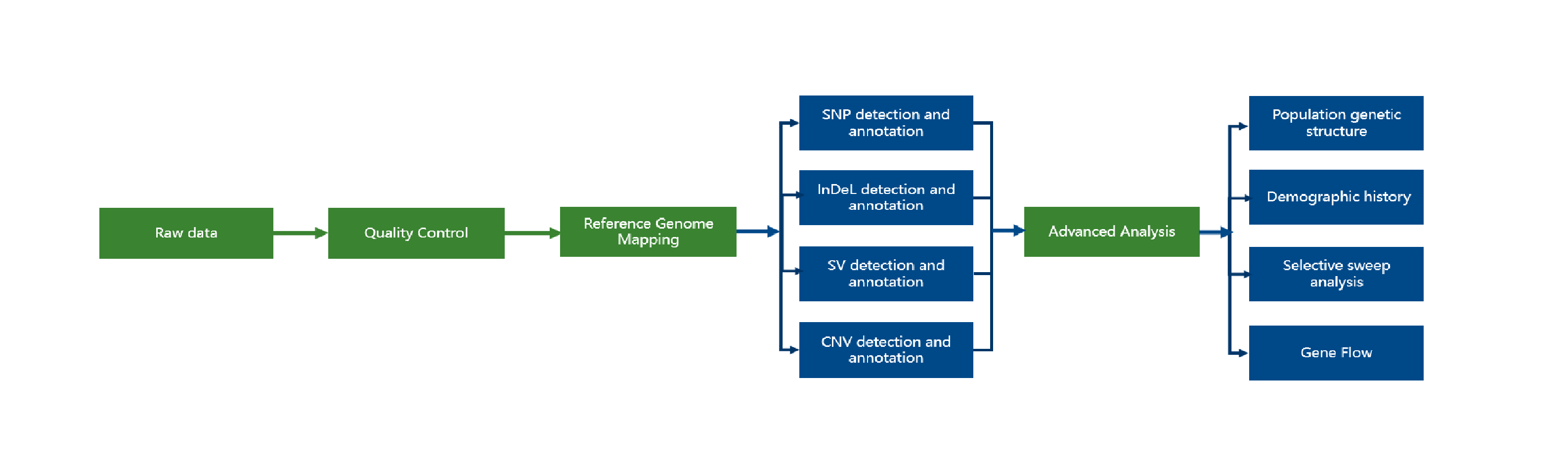

### bioinfoWorkflow.png

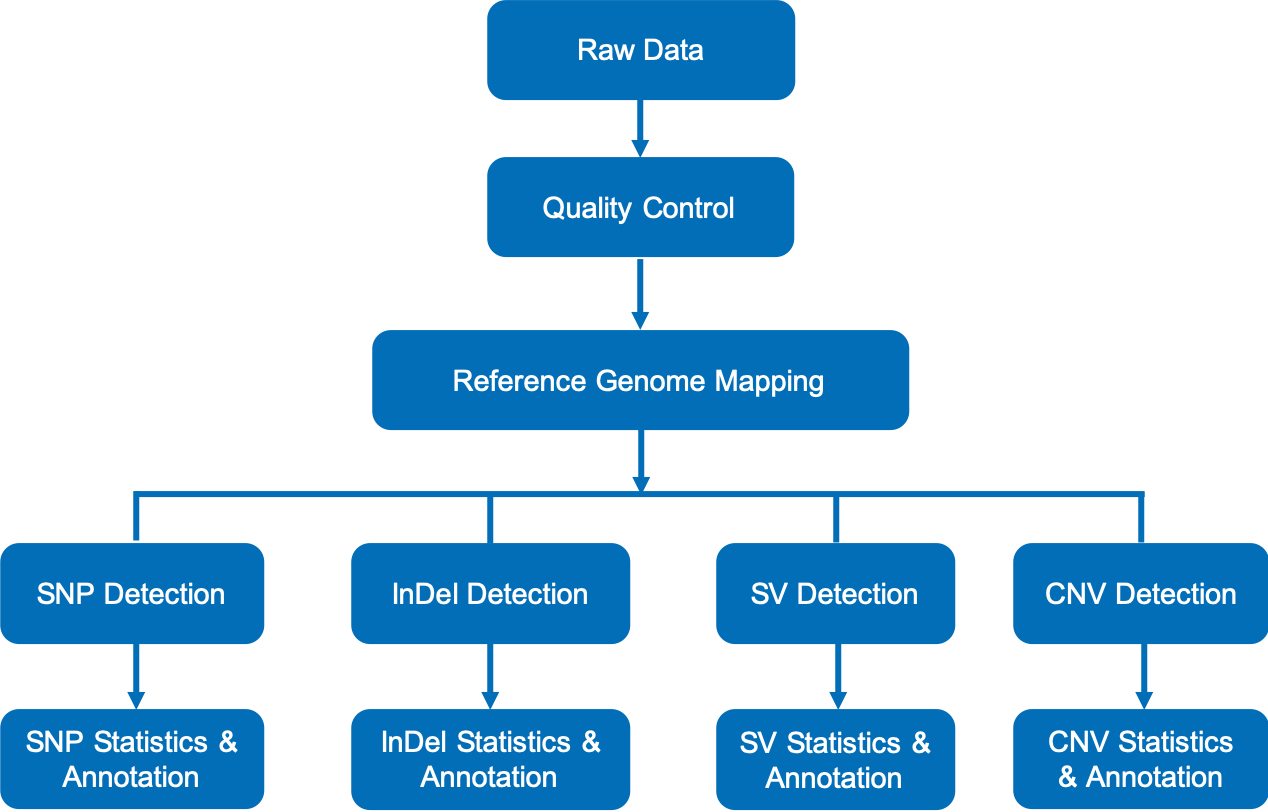

### close.gif

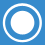

### CNV_ann_Variation_type_statistics_distribution.png

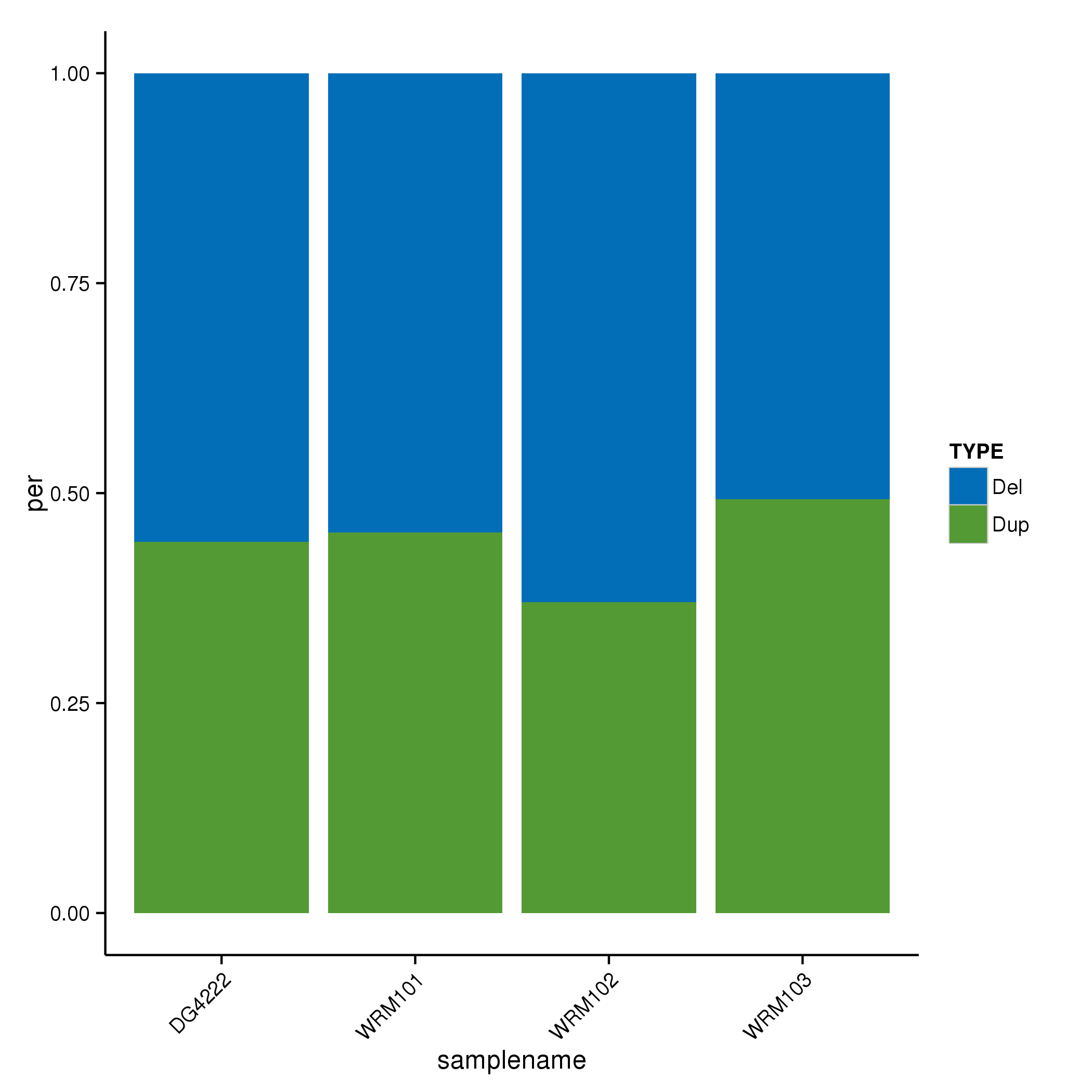

### DG4222.CNV.table.png

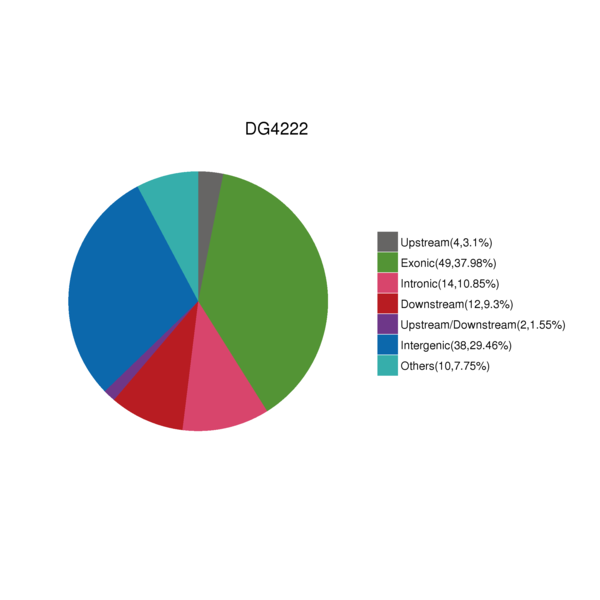

### DG4222.indDensity.png

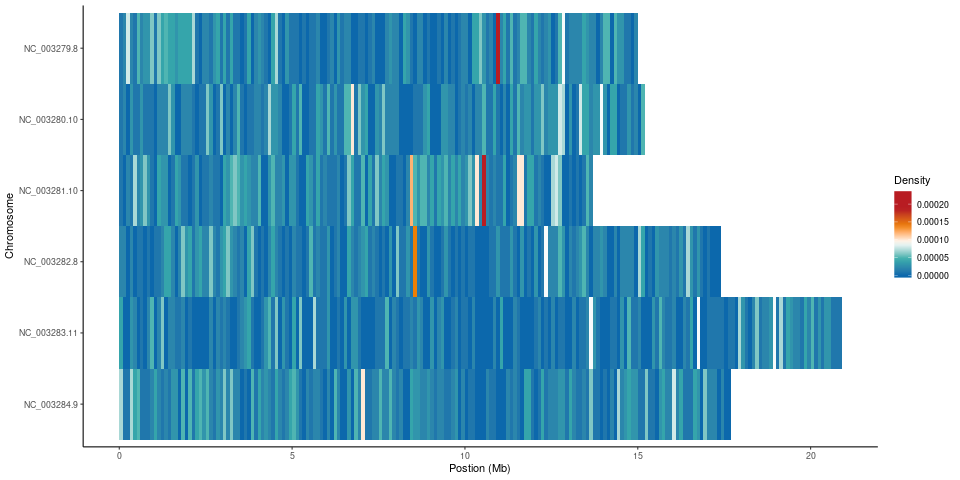

### DG4222.InDel.table.png

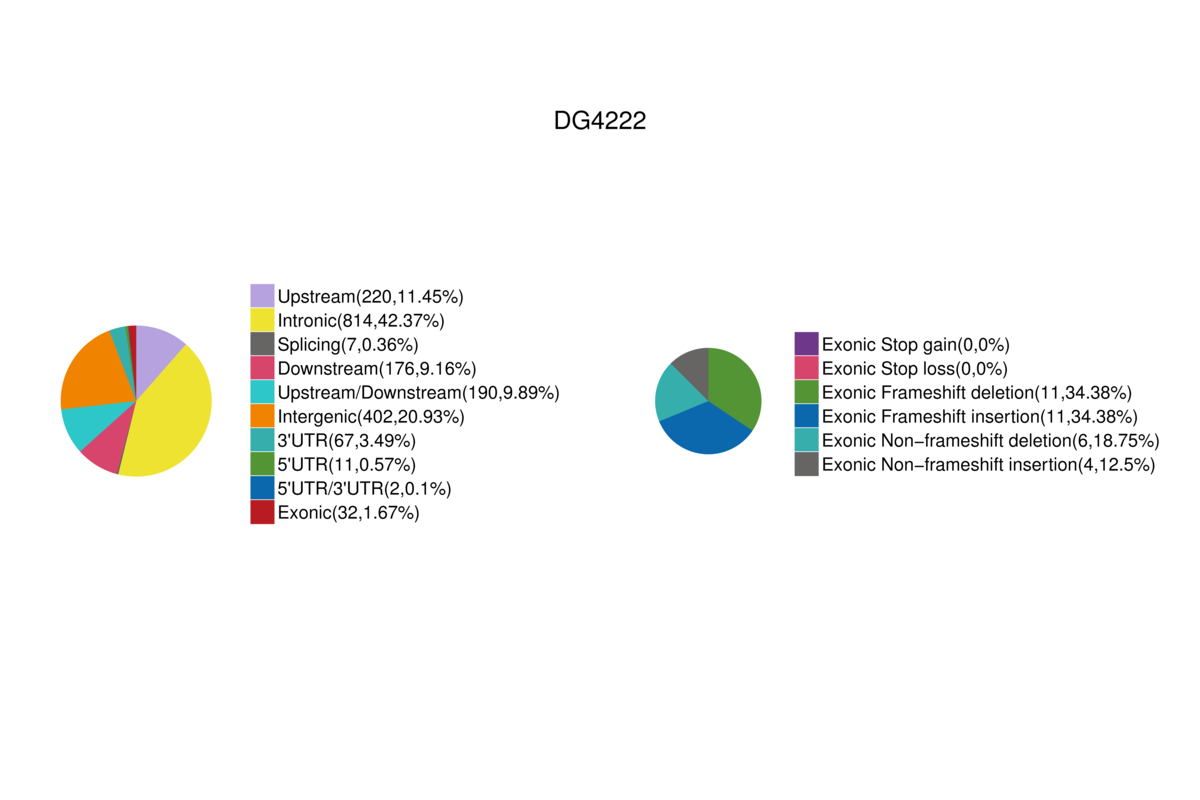

### DG4222.JPEG

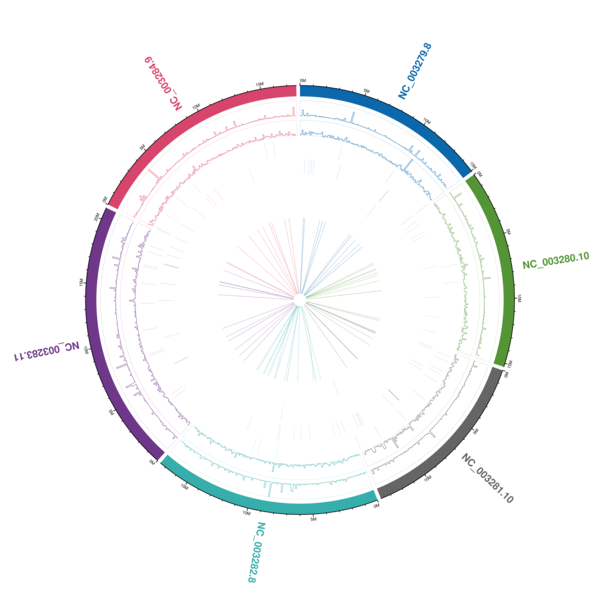

### DG4222.mapbychrdepth.png

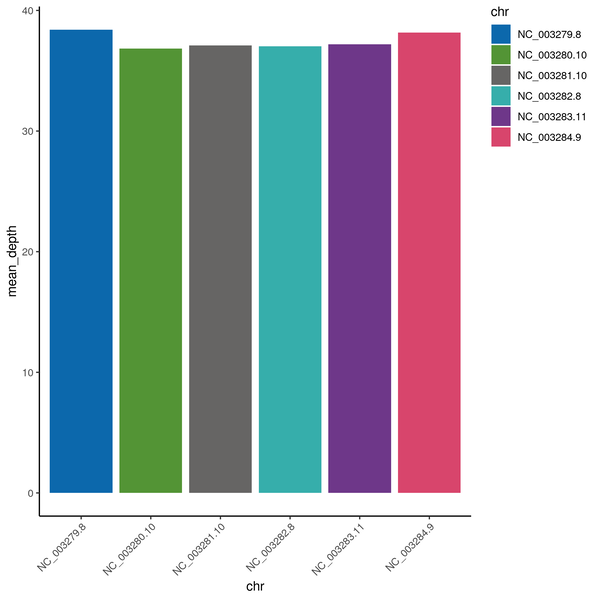

### DG4222.SNP.table.png

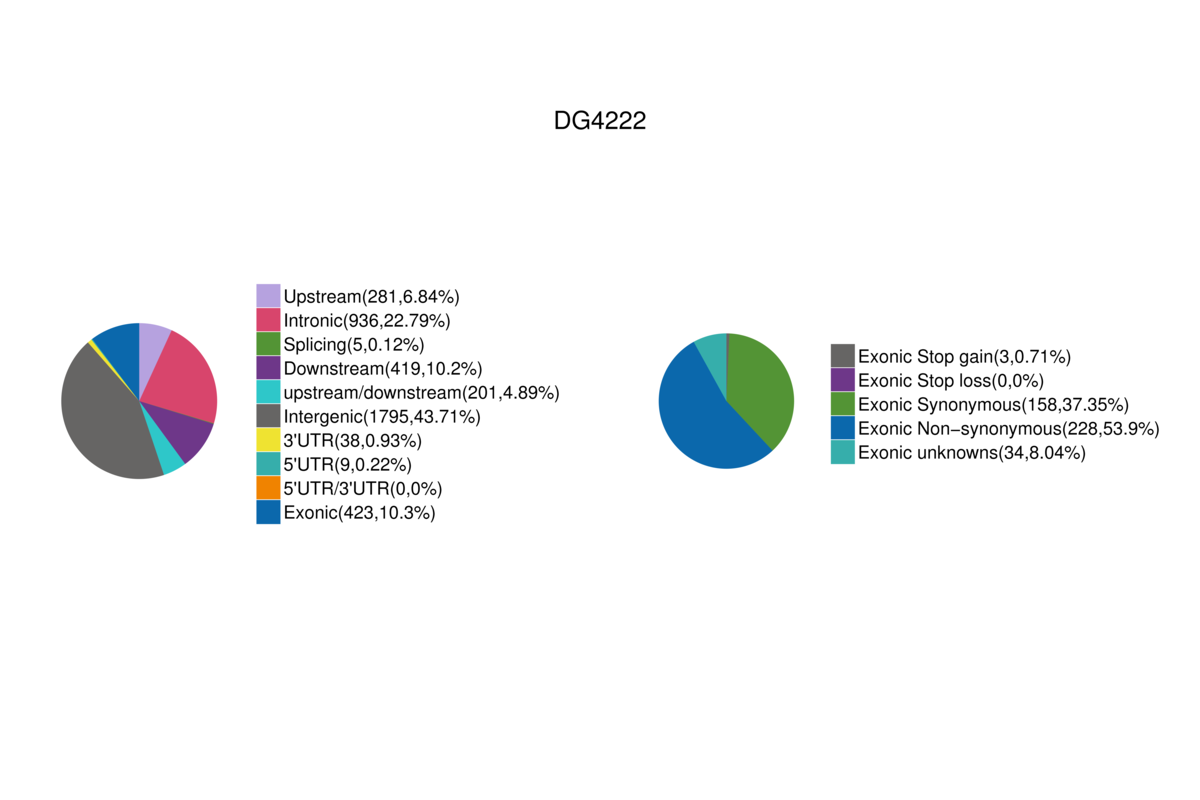

### DG4222.snpDensity.png

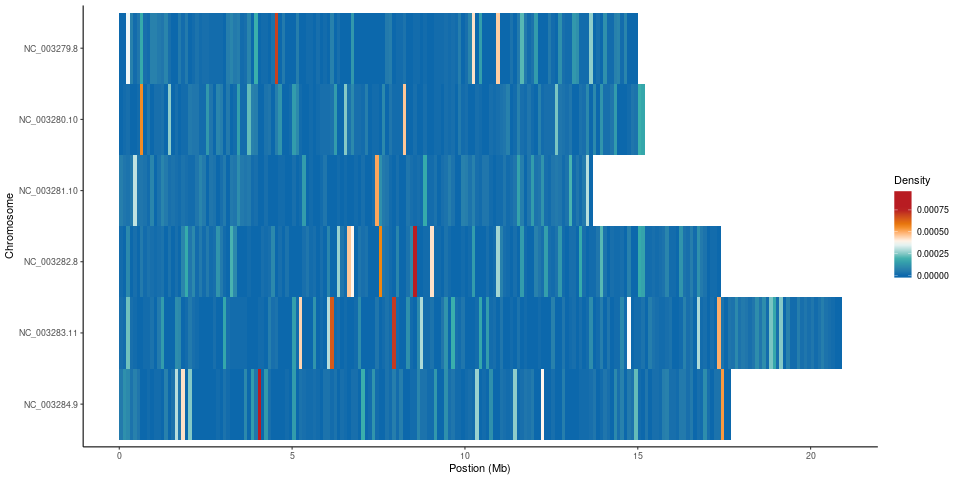

### DG4222.SV.table.png

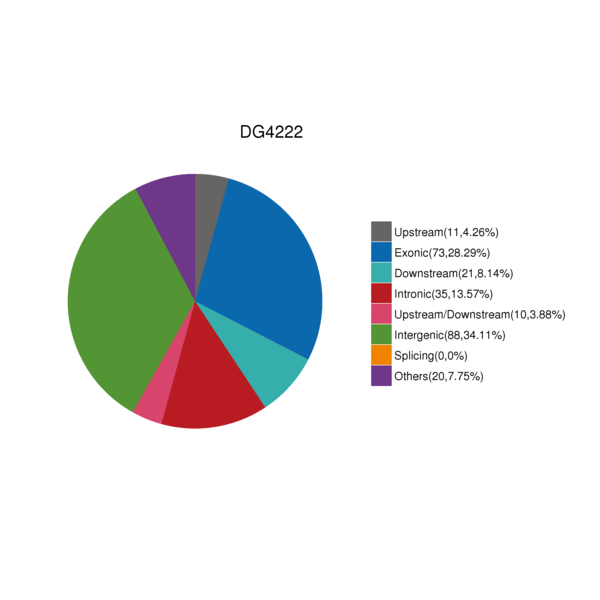

### DG4222_CKDN240021609-1A_22HTYHLT4_L3.Error.JPEG

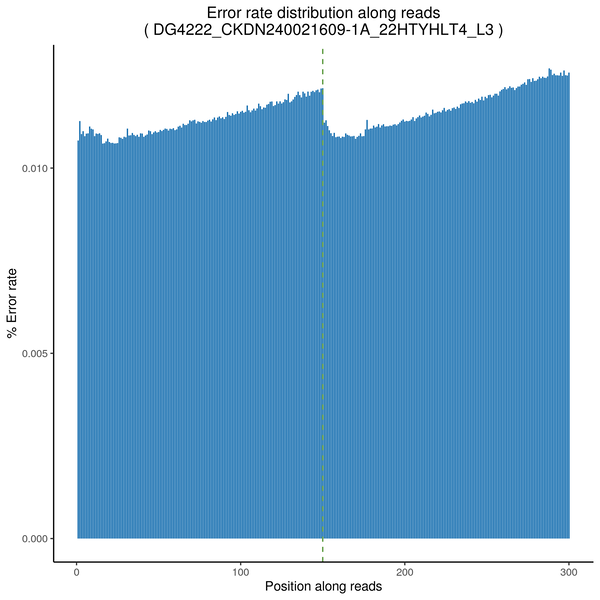

### DG4222_CKDN240021609-1A_22HTYHLT4_L3.GC.png

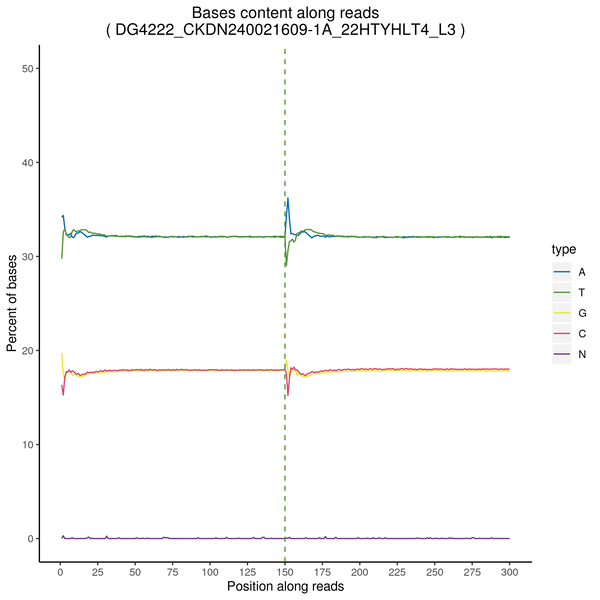

### DG4222_CKDN240021609-1A_22HTYHLT4_L3.pie3d.png

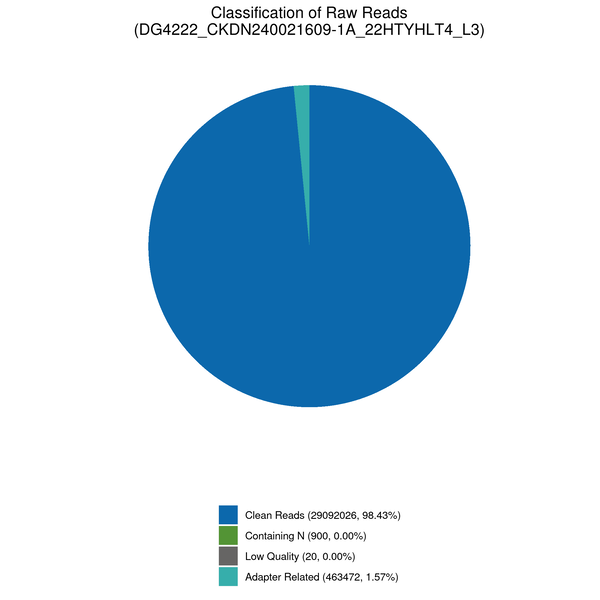

### DG4222_CKDN240021609-1A_22HTYHLT4_L3.QM.png

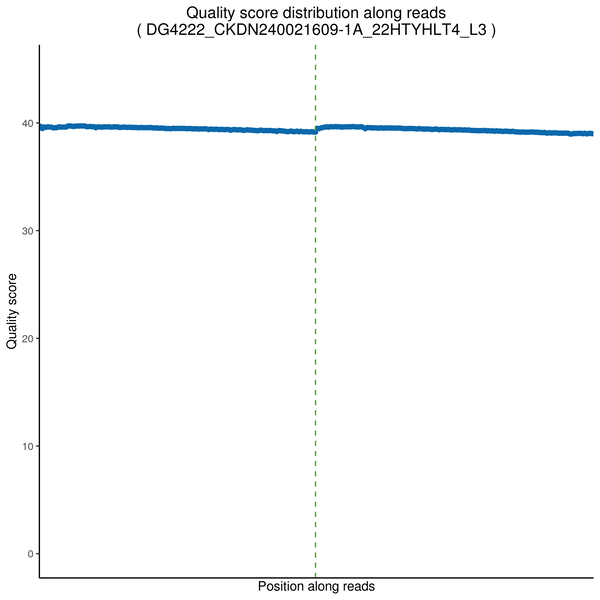

### experimentWorkflow.png

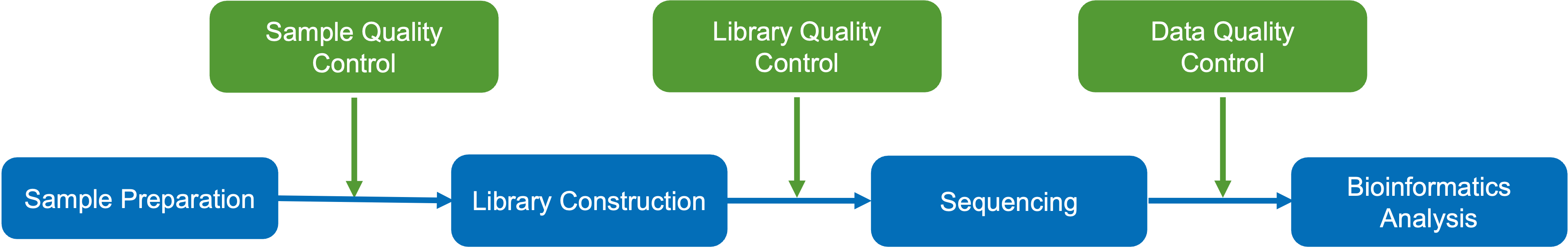

### fancy_close.png

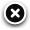

### fancy_loading.png

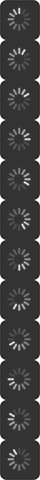

### fancy_nav_left.png

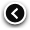

### fancy_nav_right.png

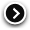

### fancy_shadow_ne.png

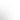

### fancy_shadow_nw.png

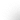

### fancy_shadow_se.png

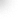

### fancy_shadow_sw.png

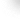

### fancy_title_left.png

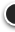

### fancy_title_right.png

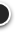

### fancybox.png

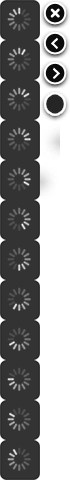

### flow_snpindel.adv.png

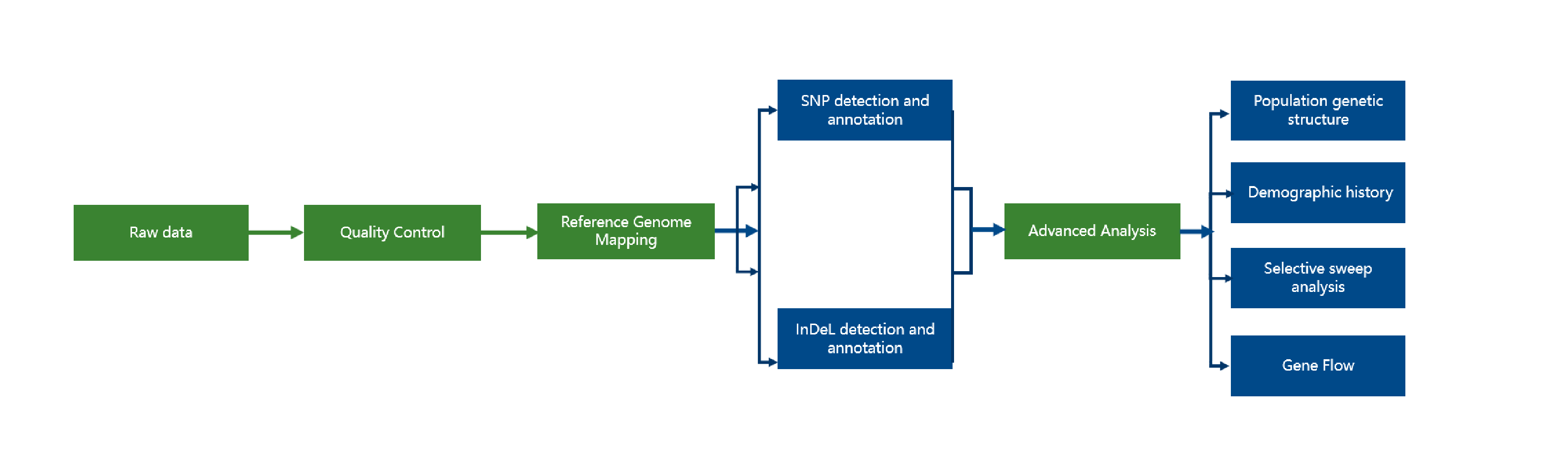

### flow_snpindel.png

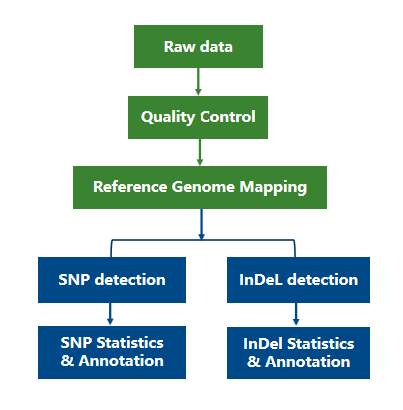

### goTop.jpg

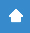

### igv.png

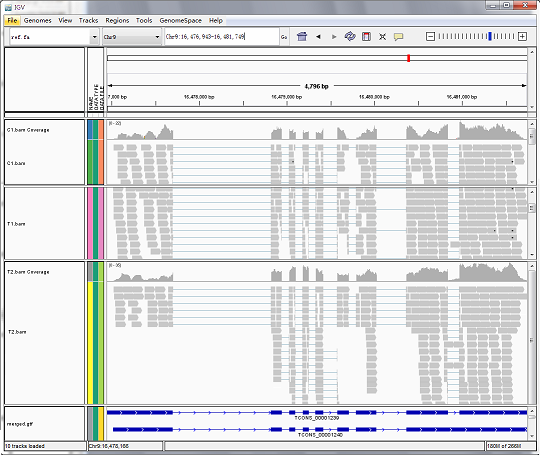

### illumina_experiment_pipeline.png

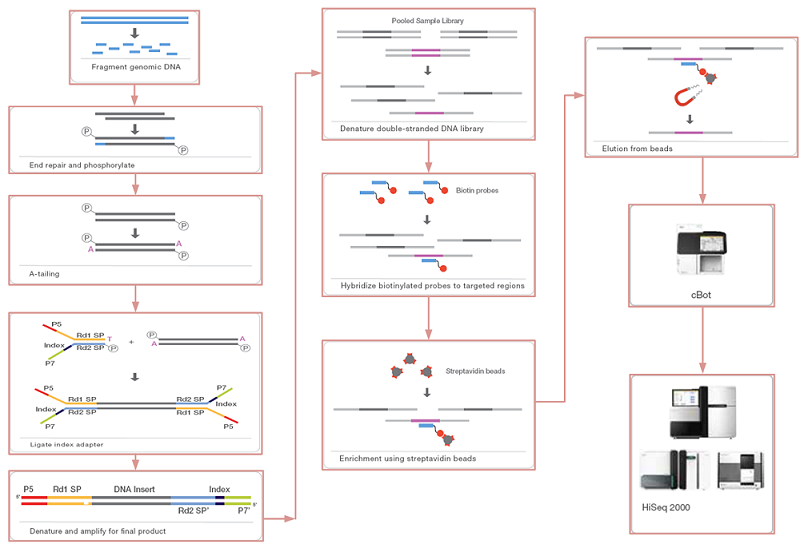

### InDel_CDSpercentage.png

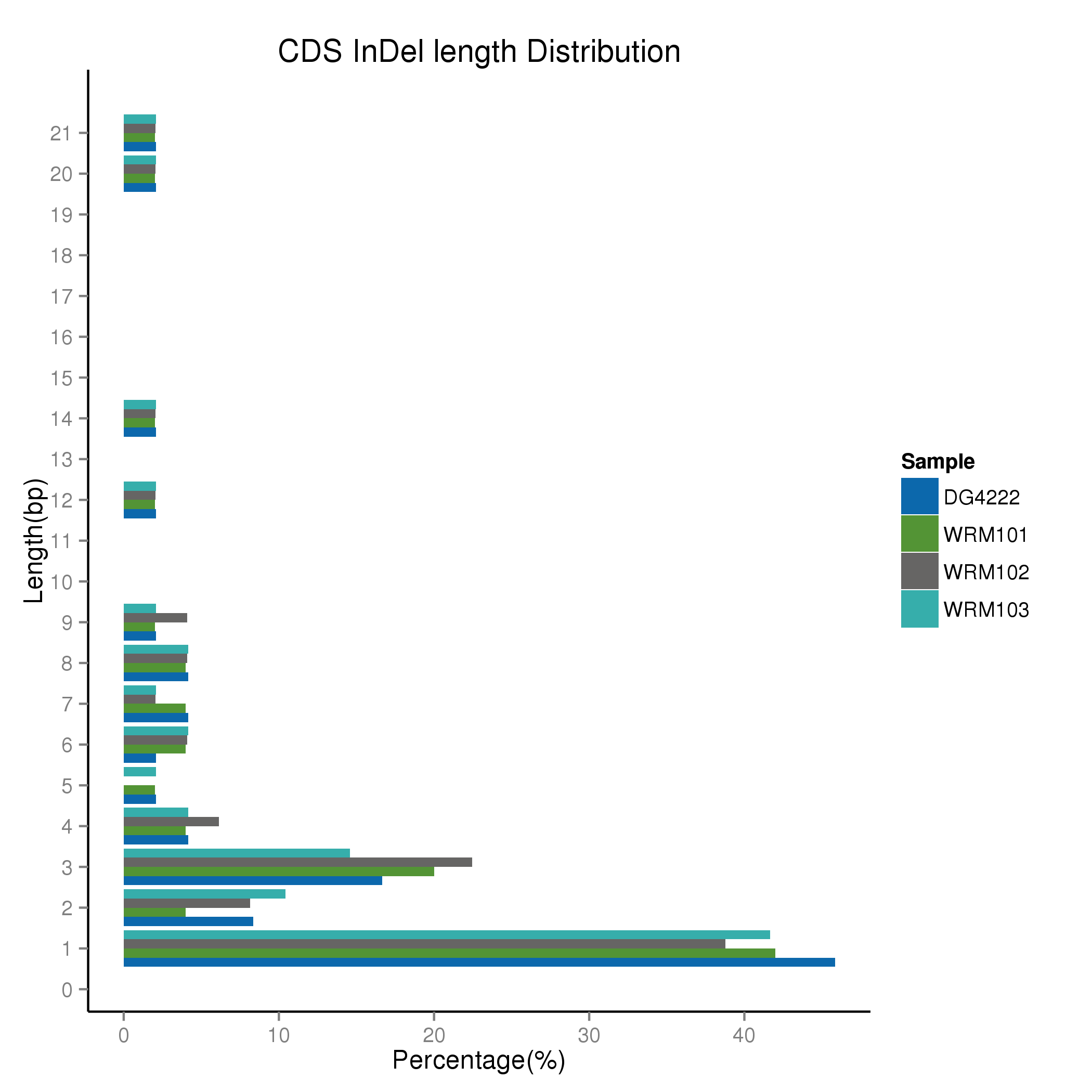

### InDel_GENOMEpercentage.png

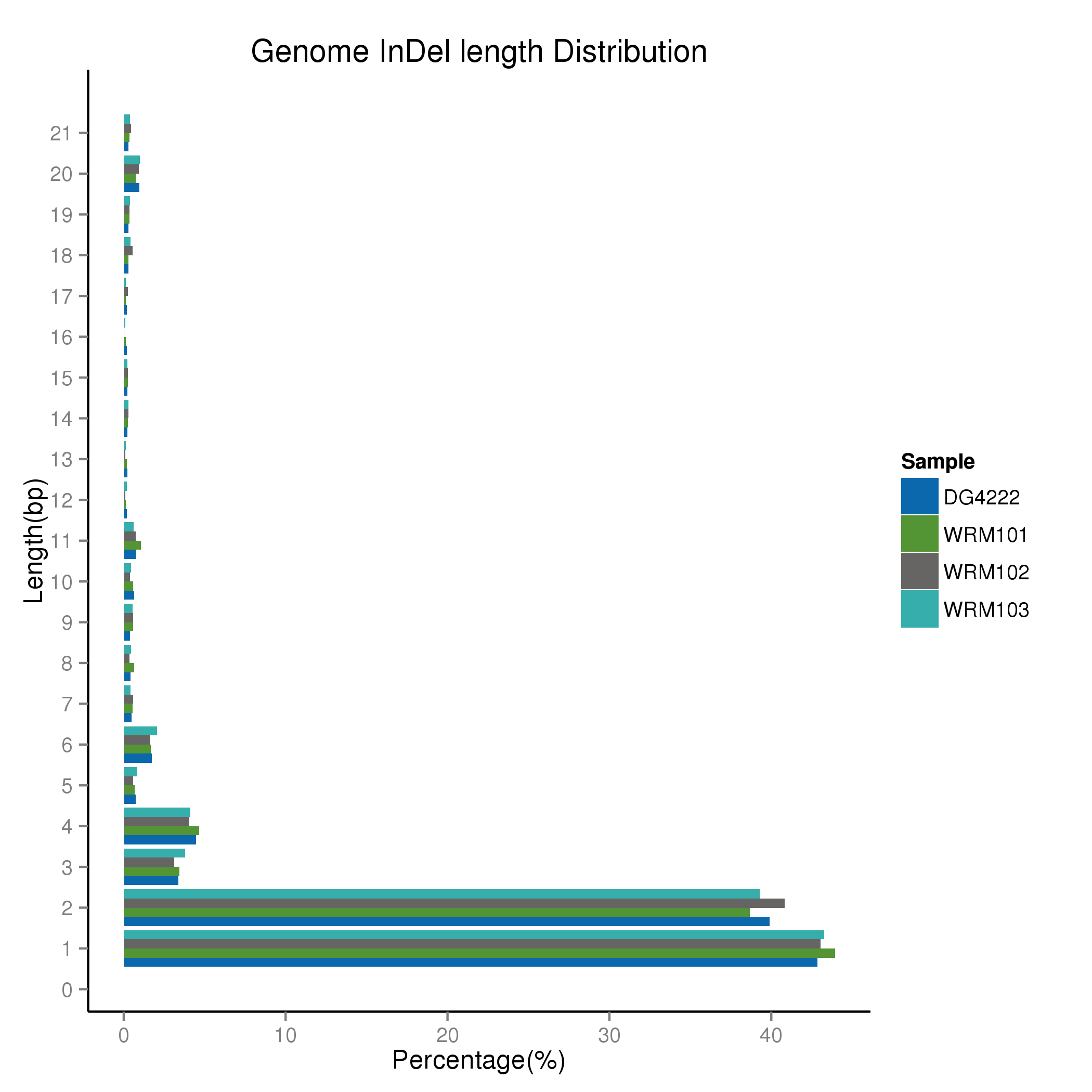

### LibPrep_PCR-free.png

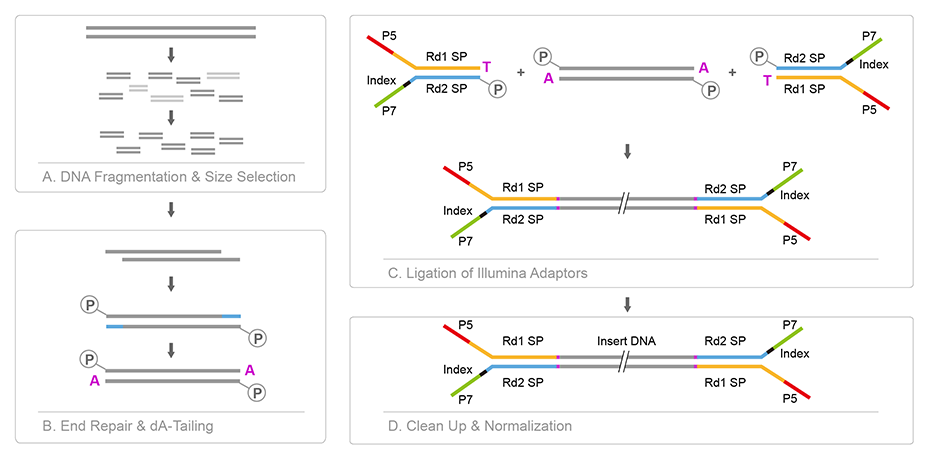

### logo1.png

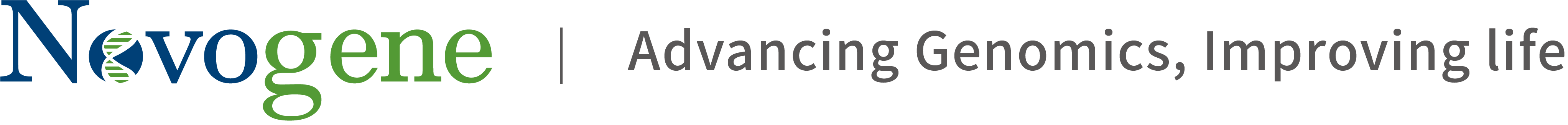

### logo.png

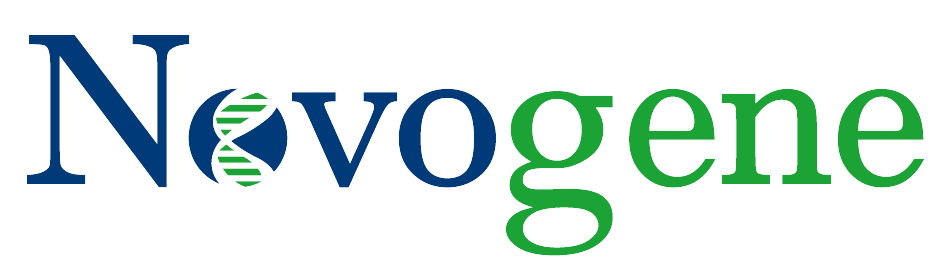

### mapping.png

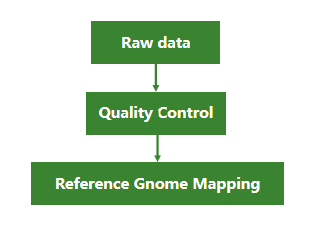

### novogeneLogo.png

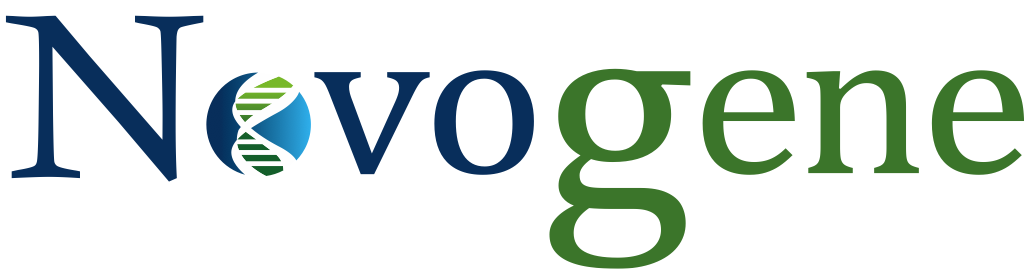

### open.gif

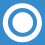

### QC.png

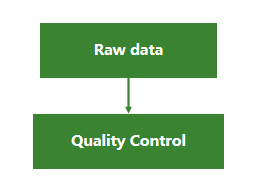

### savant.png

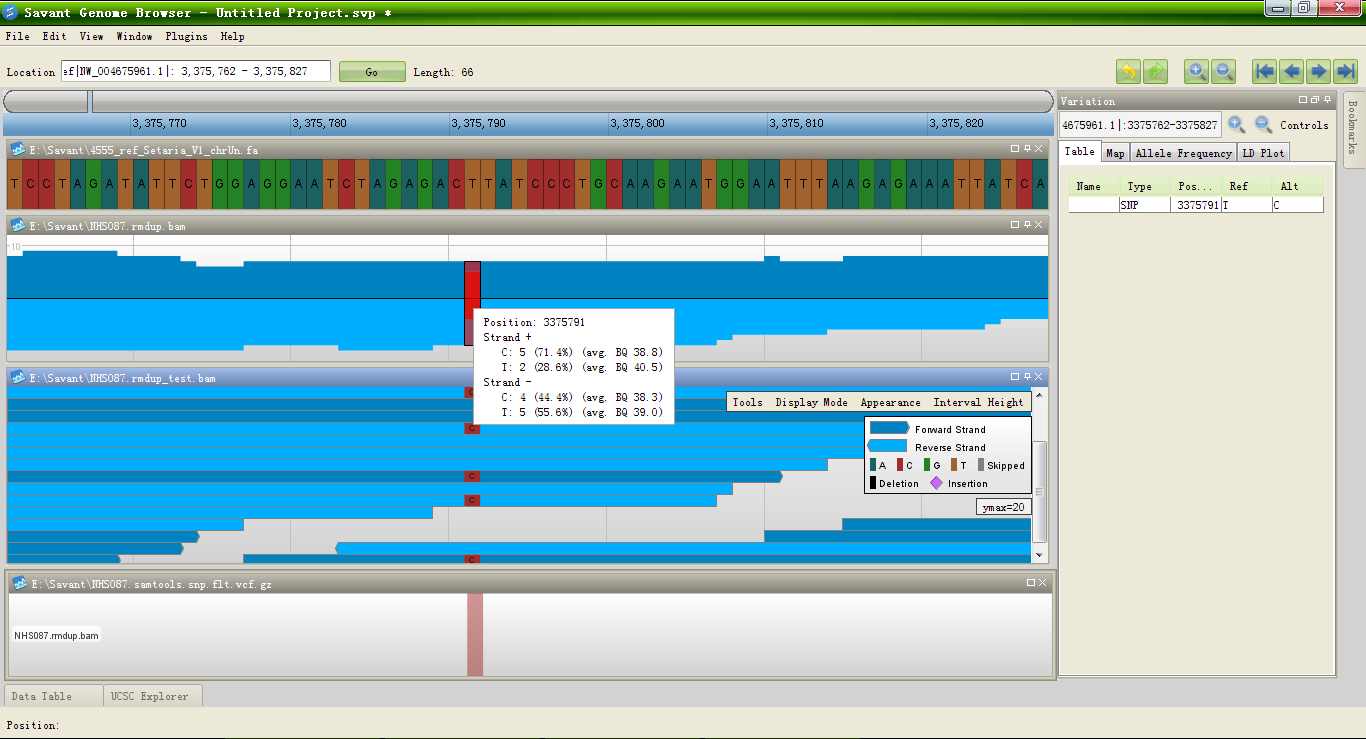

### SNP.frequency.xls.barplot.png

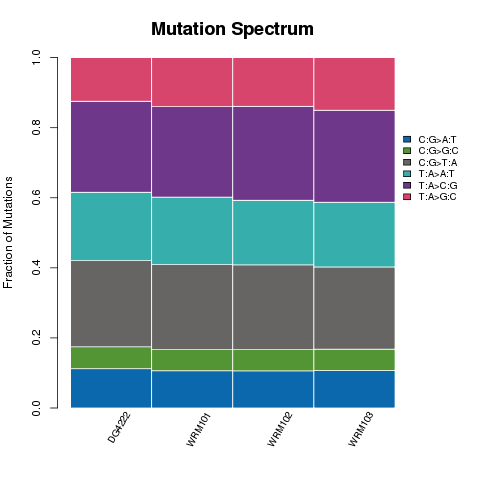

### SNP_distance_cumulative_distribution.JPEG

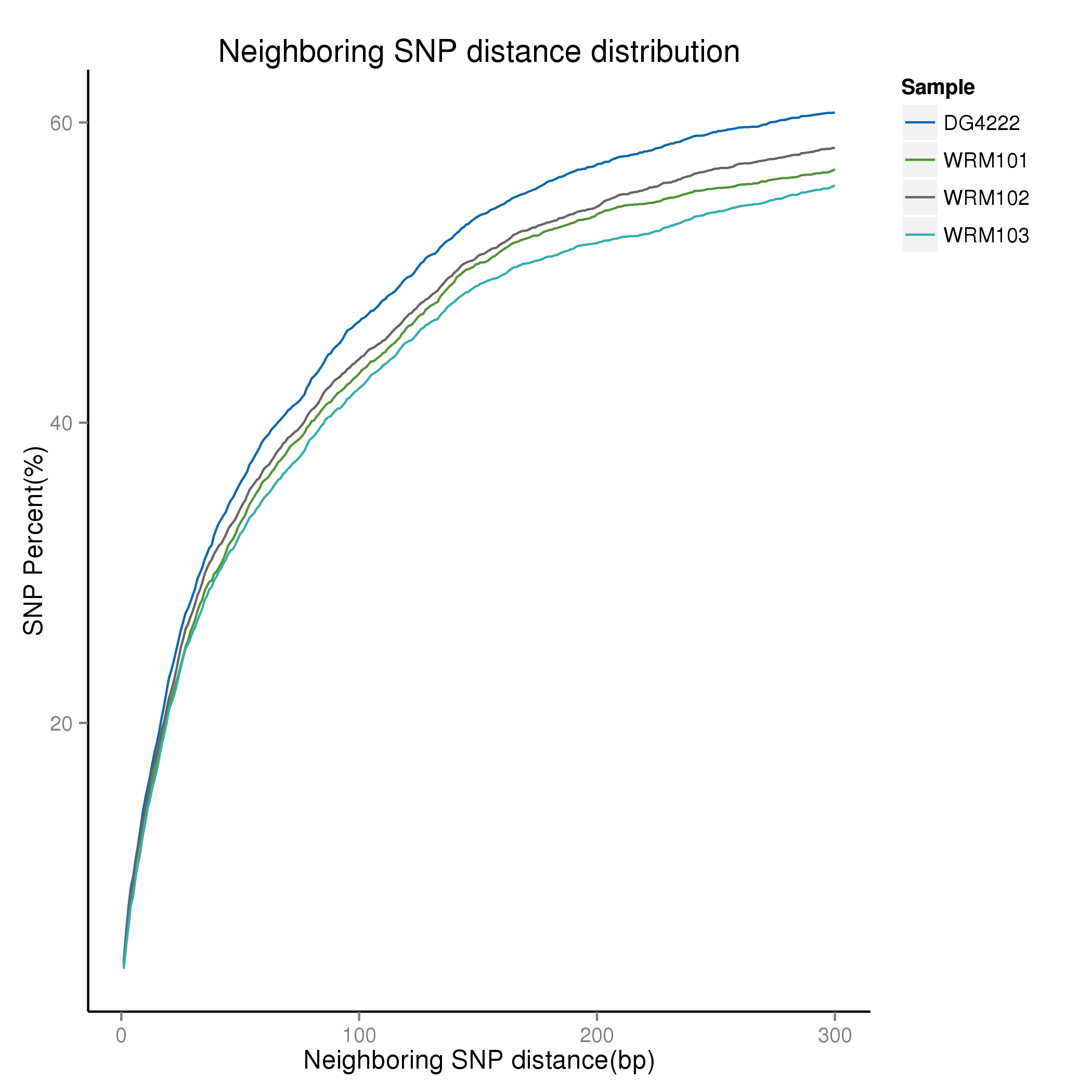

### SNP_frequency.png

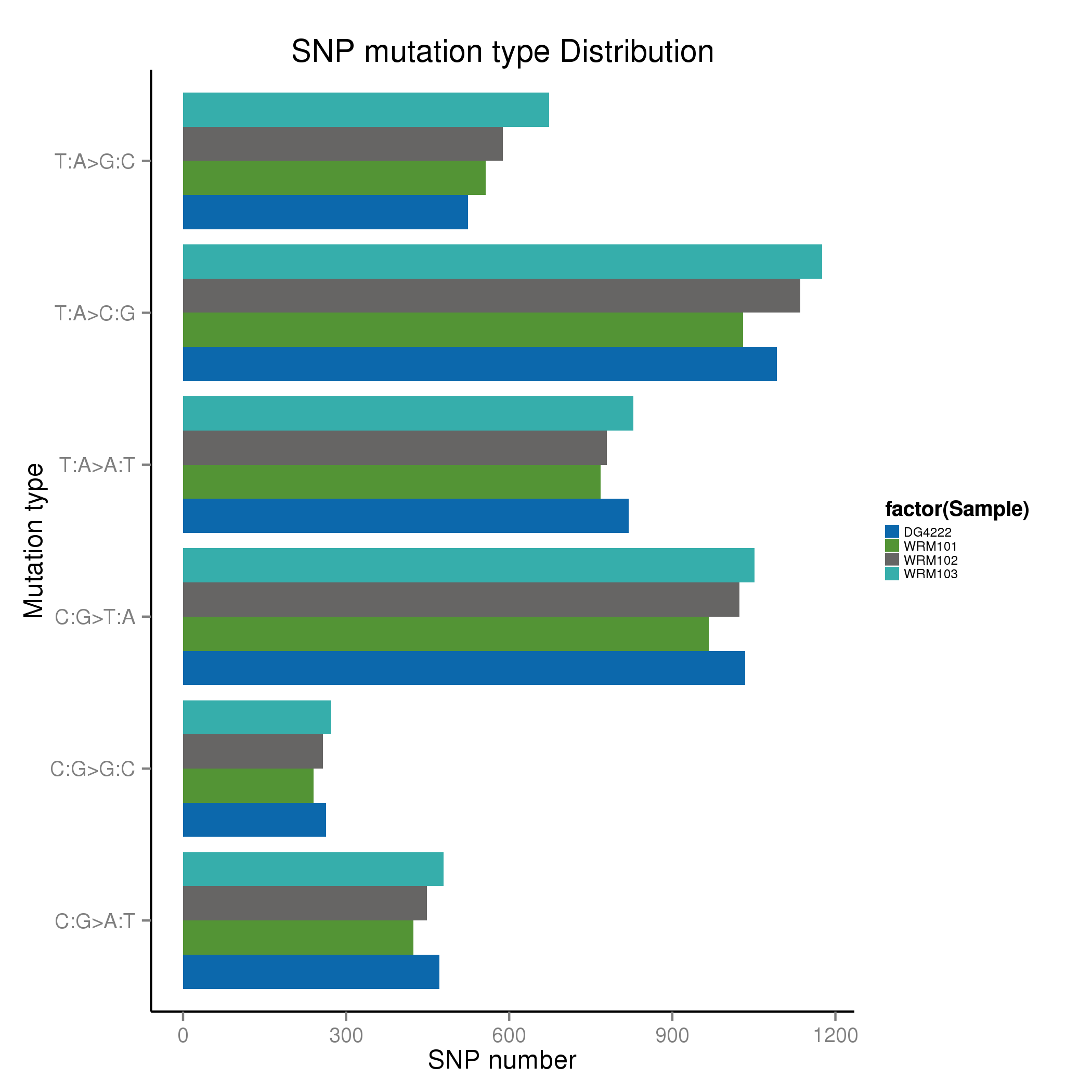

### SNP_quality_cumulative_distribution.JPEG

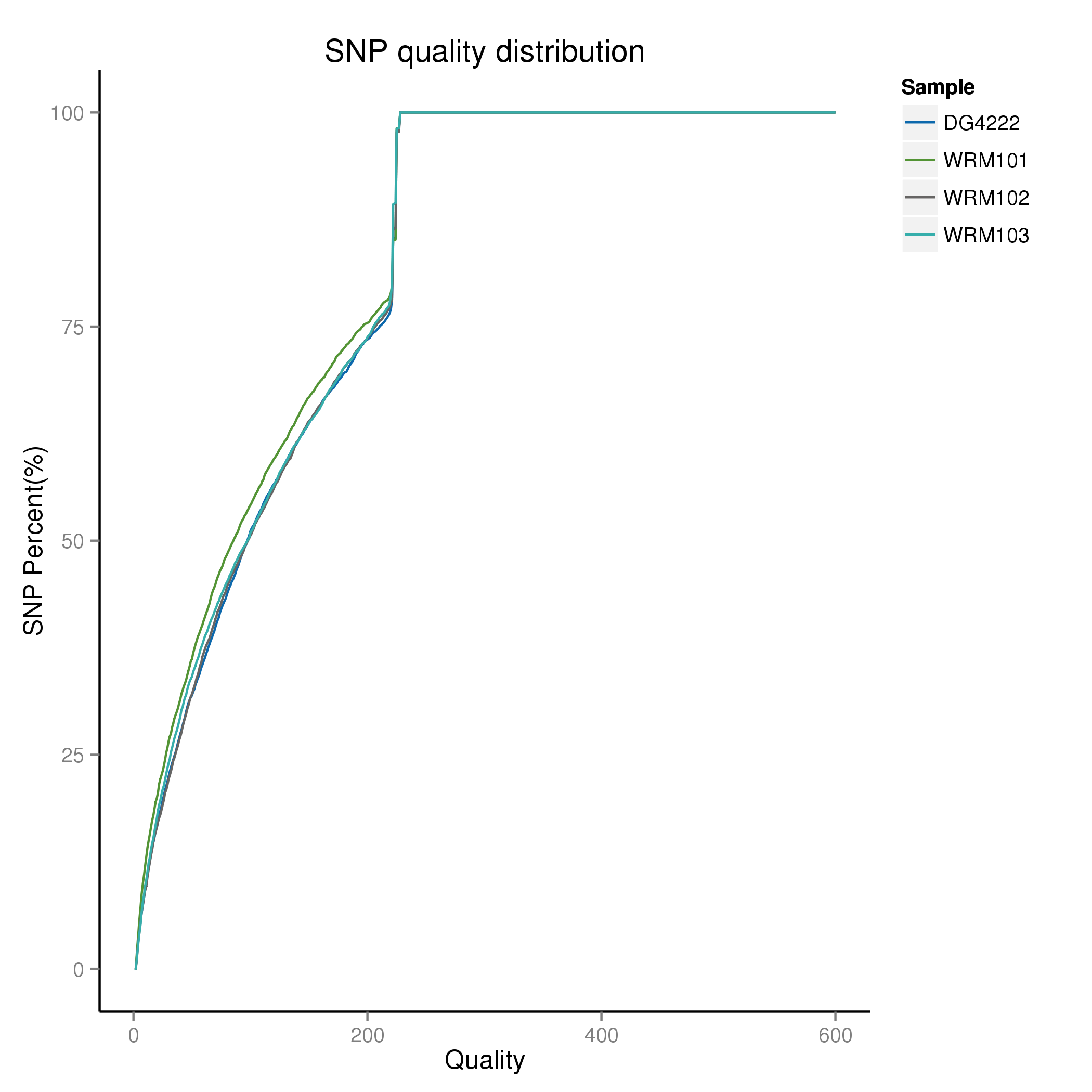

### SNP_readsNum_cumulative_distribution.JPEG

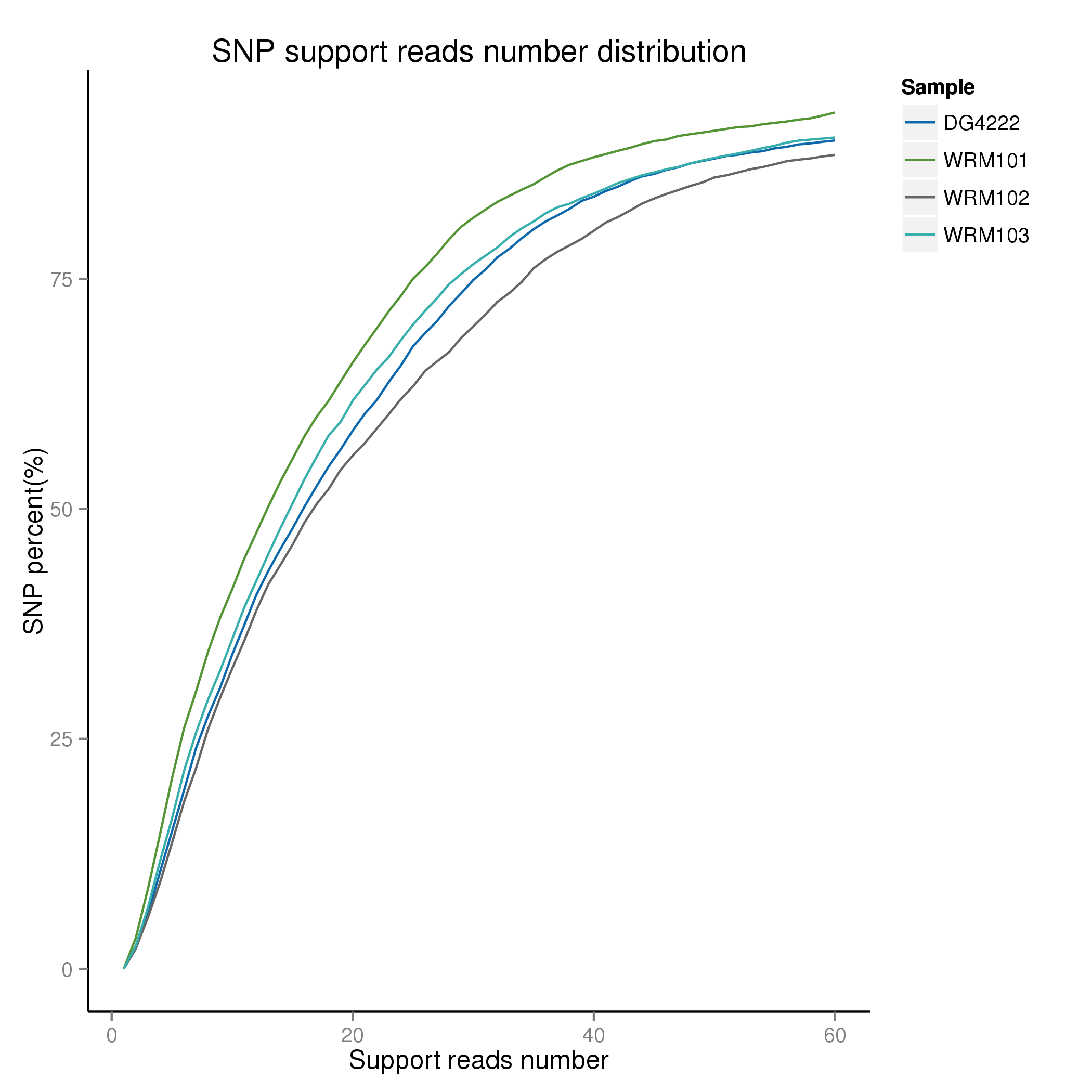

### sort_both.png

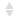

### sort_desc.png

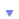

### SV_ann_Variation_type_statistics_distribution.png

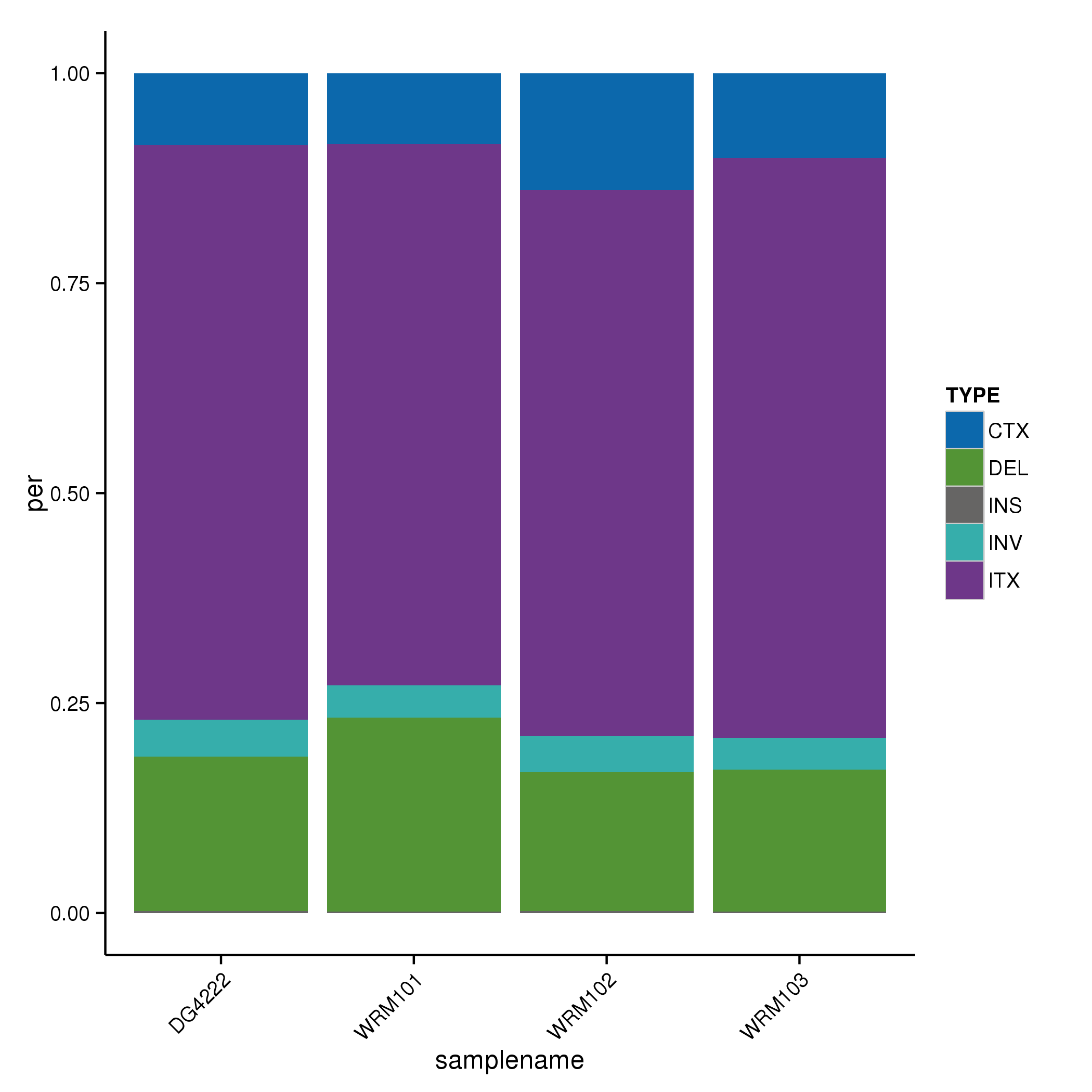

### SV_length_distribution.png

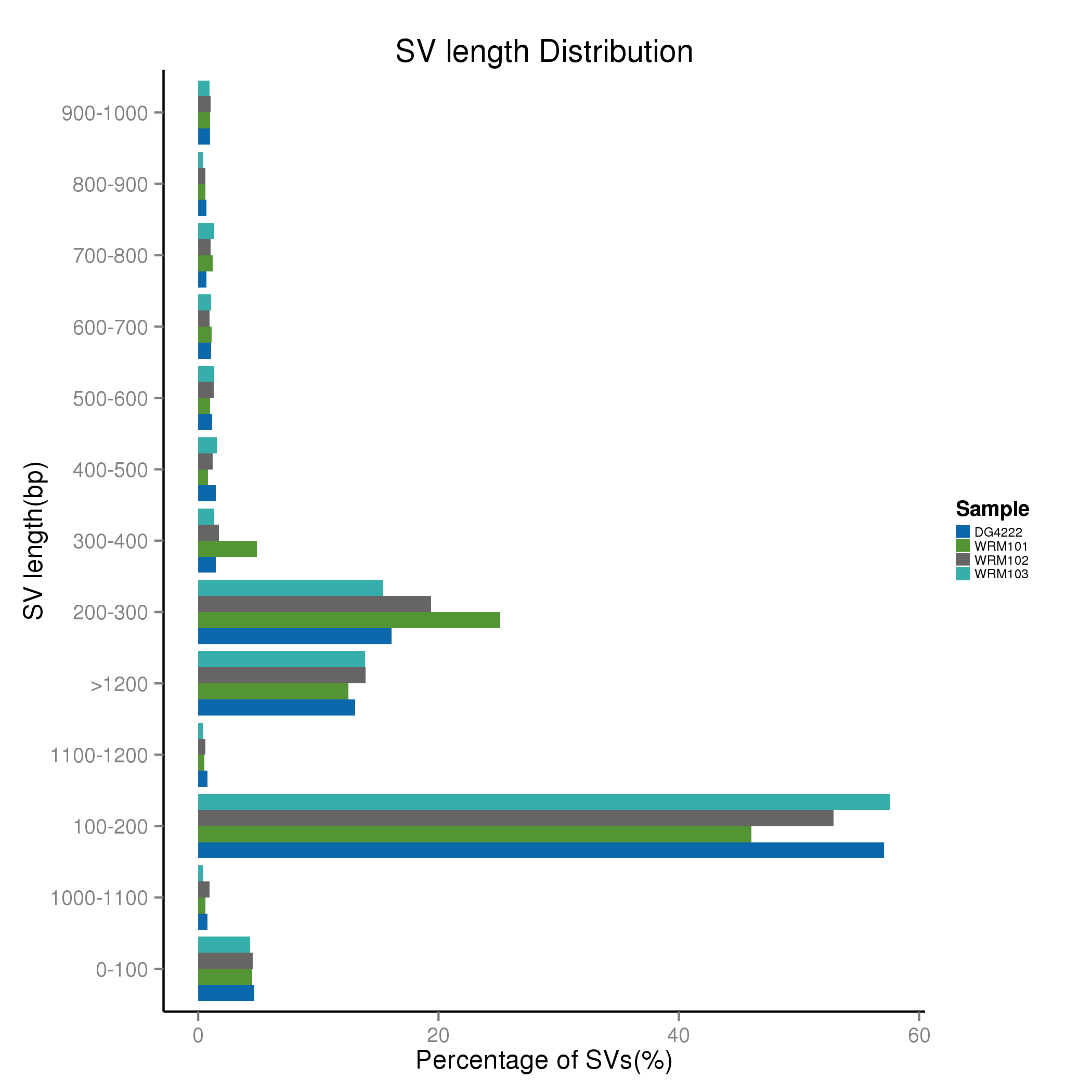

### wgs_experiment_pipeline.png

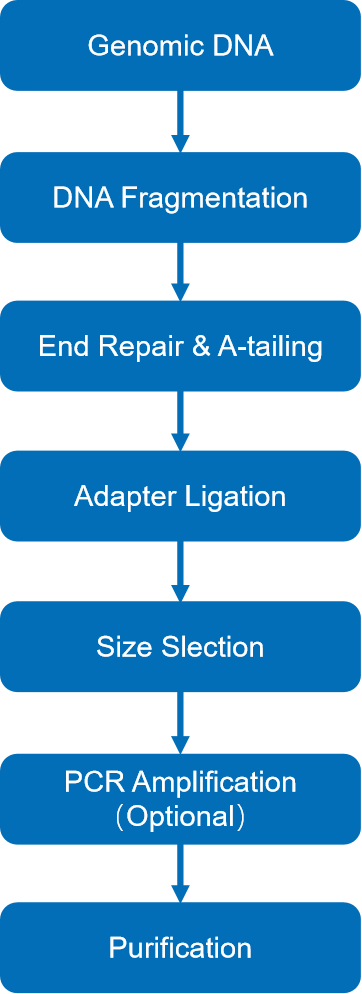

### WRM101.CNV.table.png

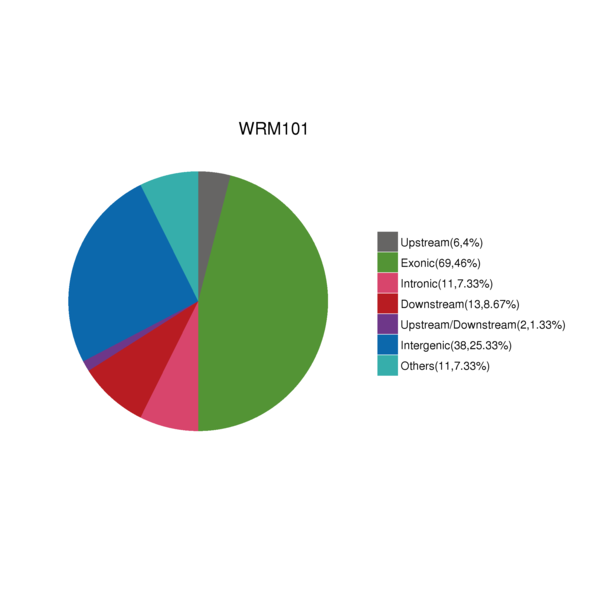

### WRM101.indDensity.png

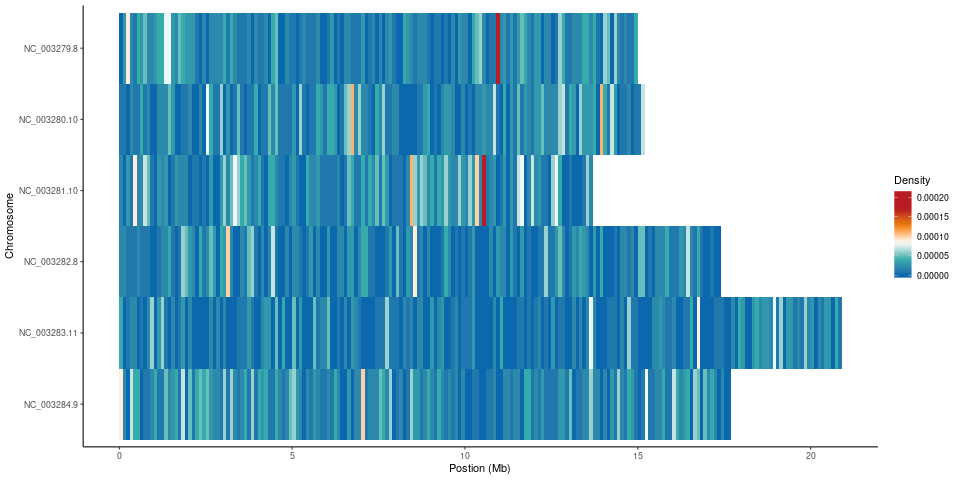

### WRM101.InDel.table.png

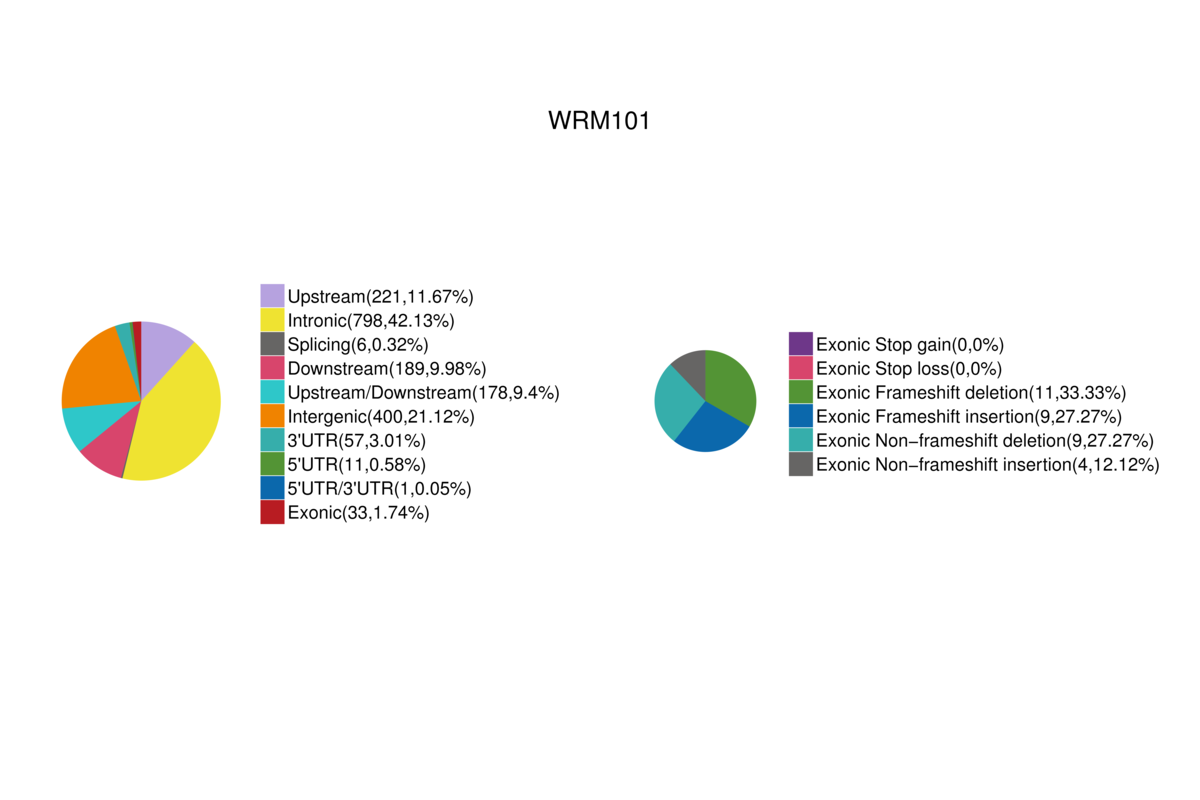

### WRM101.JPEG

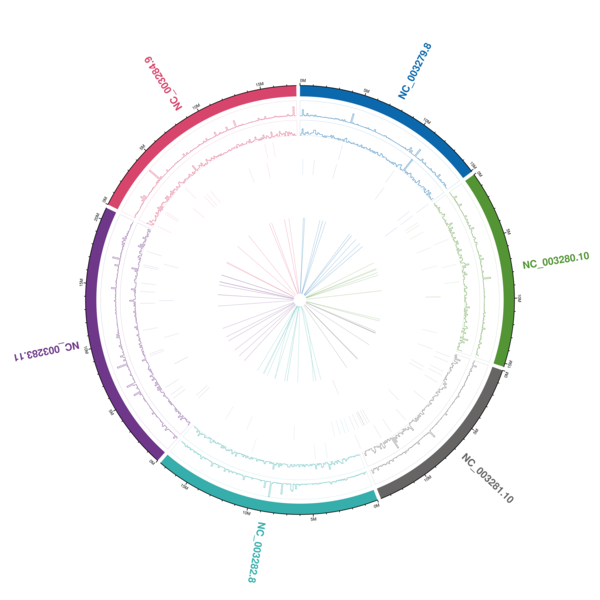

### WRM101.mapbychrdepth.png

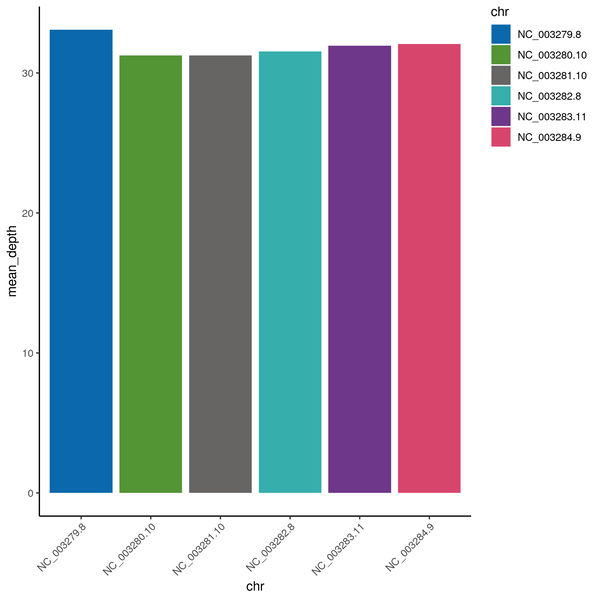

### WRM101.SNP.table.png

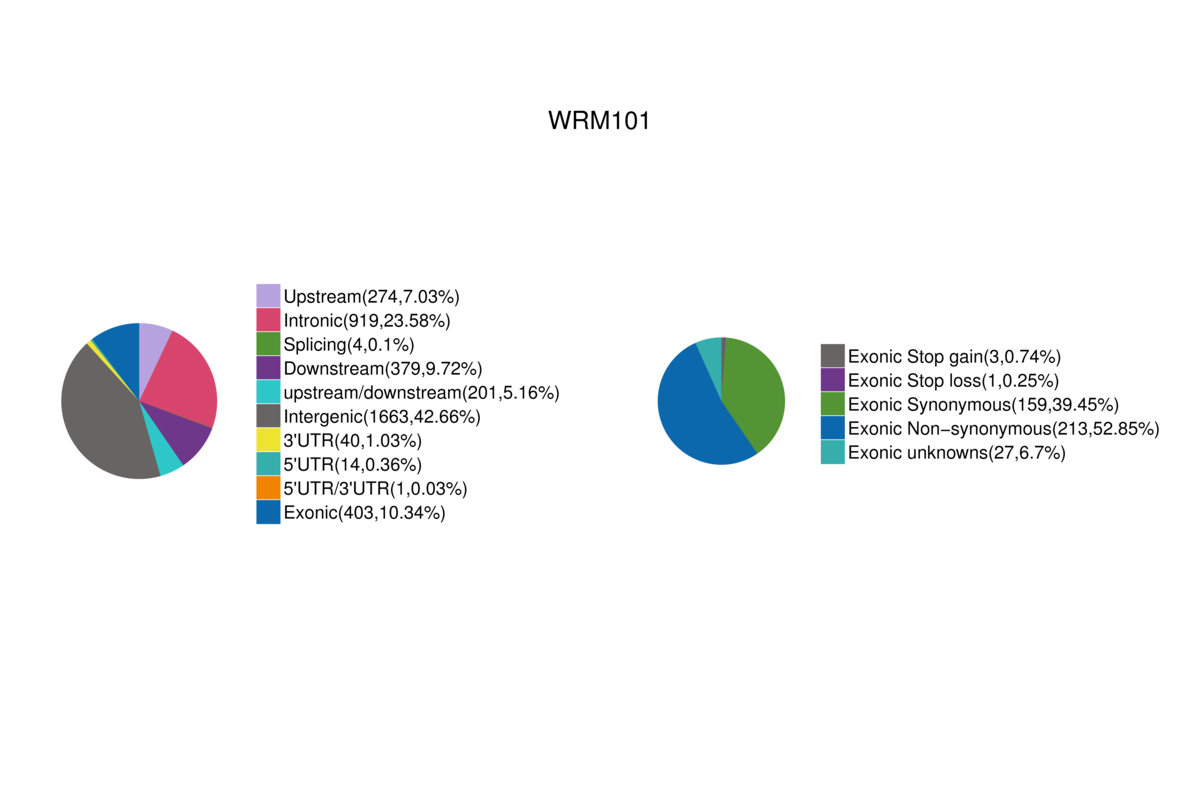

### WRM101.snpDensity.png

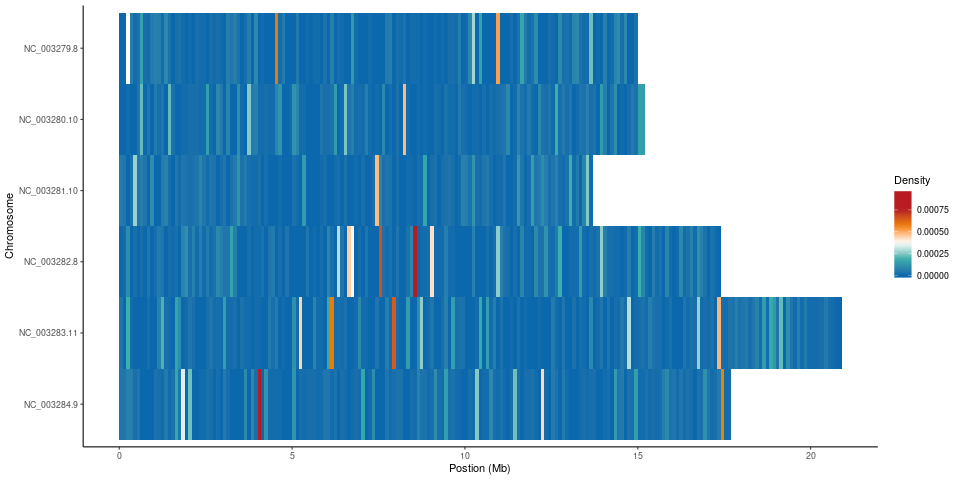

### WRM101.SV.table.png

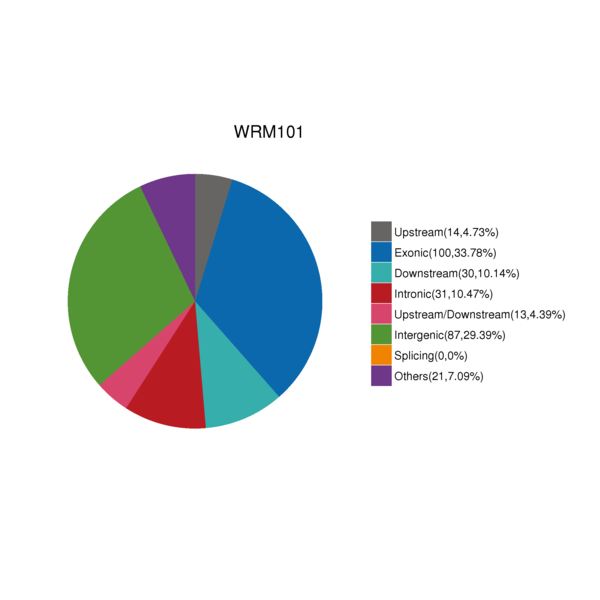

### WRM101_CKDN240021610-1A_22HTVFLT4_L1.Error.JPEG

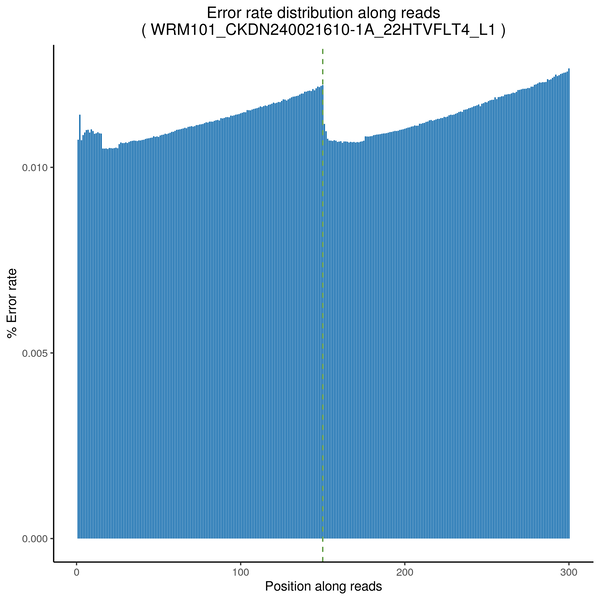

### WRM101_CKDN240021610-1A_22HTVFLT4_L1.GC.JPEG

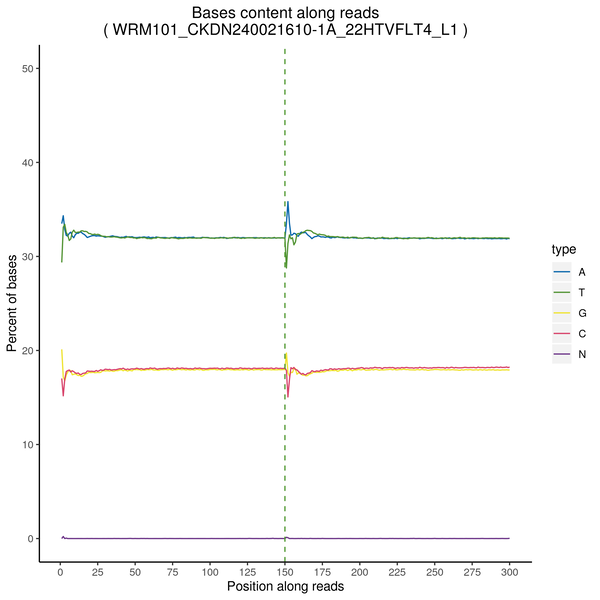

### WRM101_CKDN240021610-1A_22HTVFLT4_L1.pie3d.png

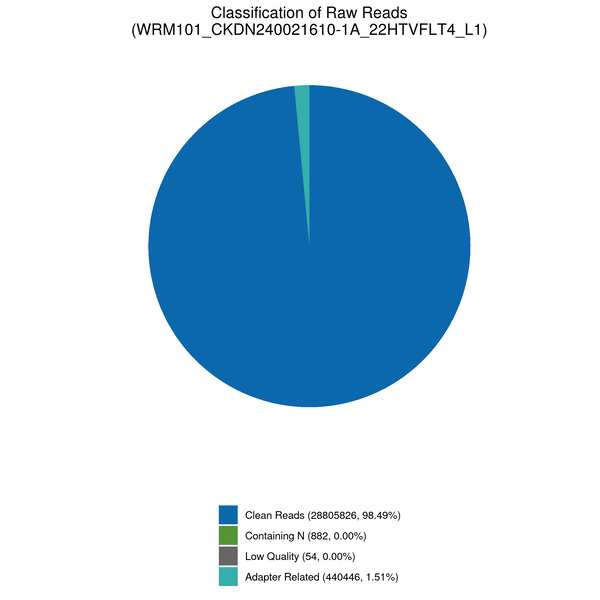

### WRM101_CKDN240021610-1A_22HTVFLT4_L1.QM.JPEG

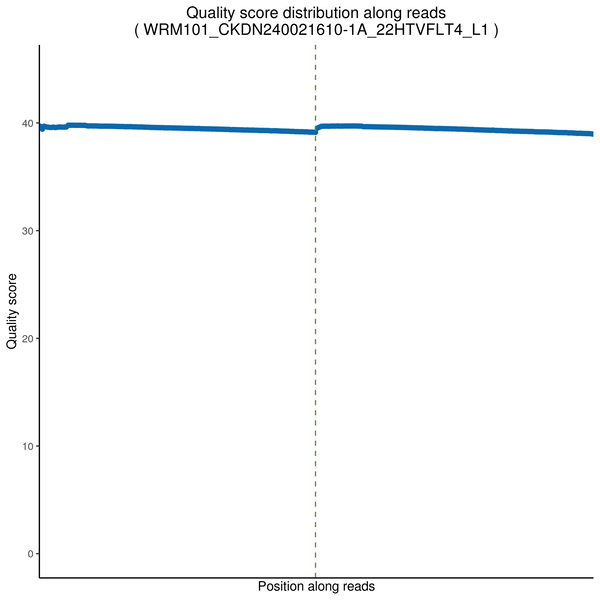

### WRM102.CNV.table.png

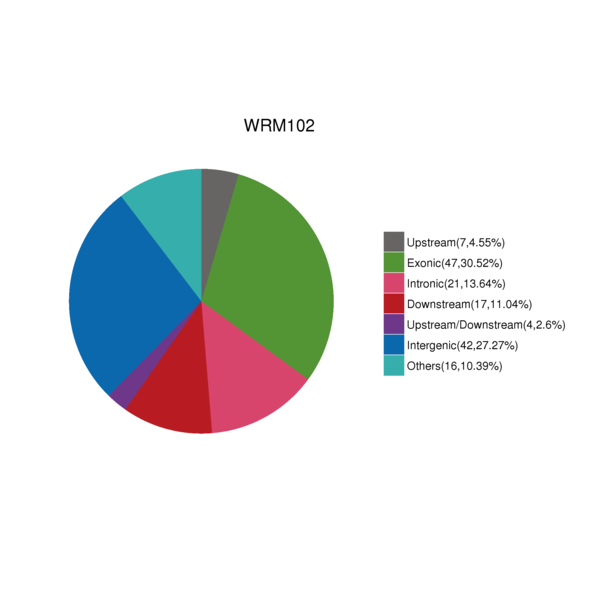

### WRM102.indDensity.png

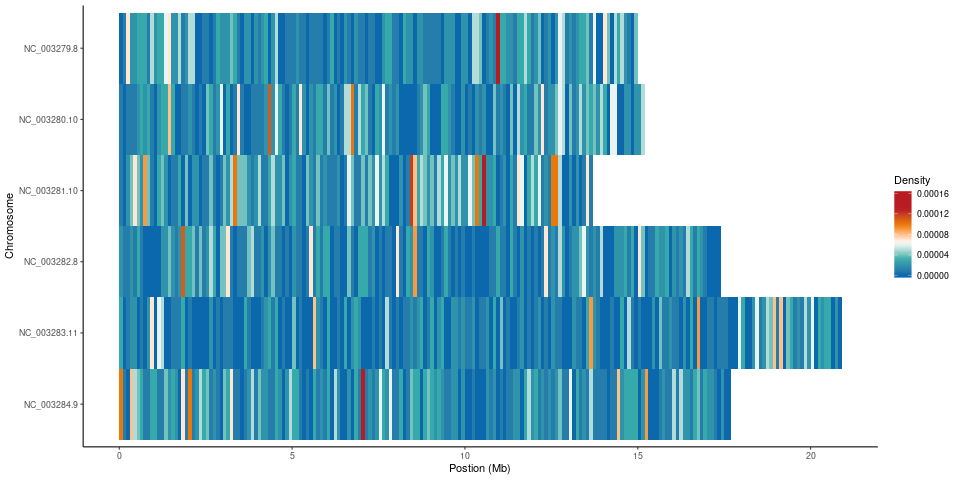

### WRM102.InDel.table.png

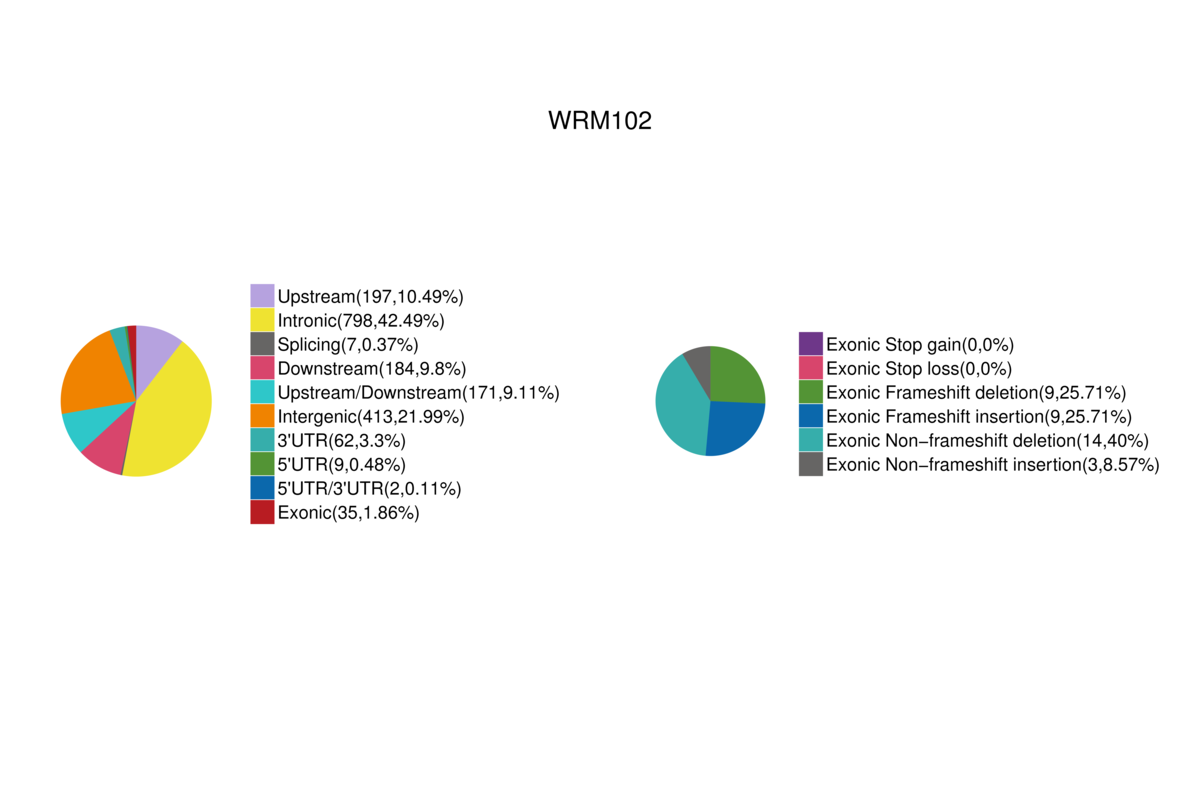

### WRM102.JPEG

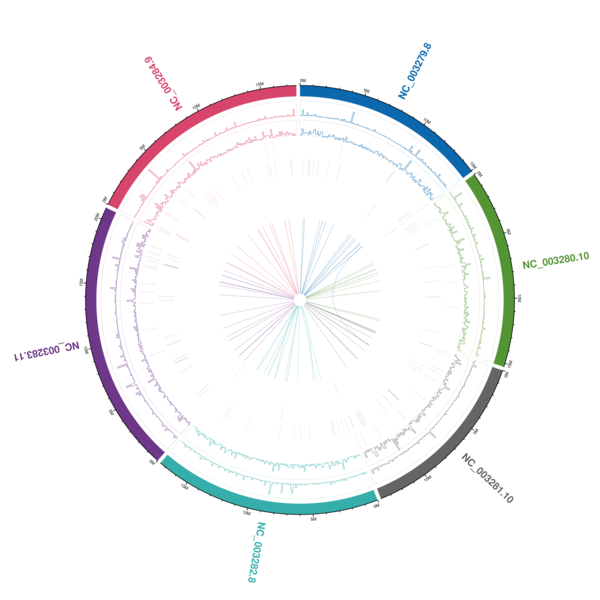

### WRM102.mapbychrdepth.png

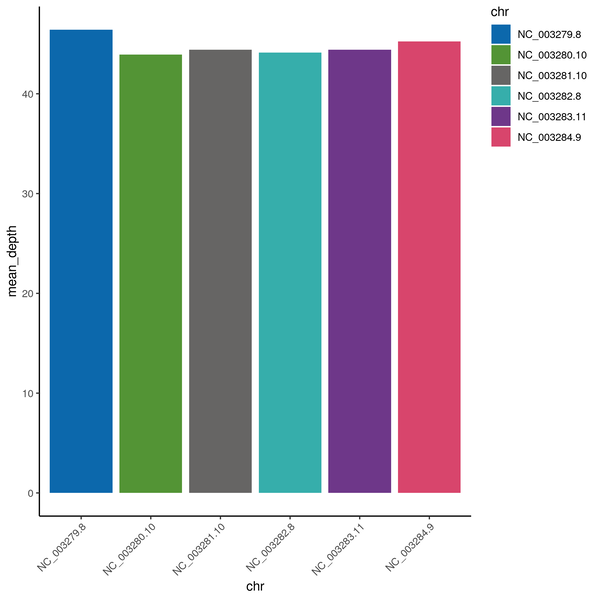

### WRM102.SNP.table.png

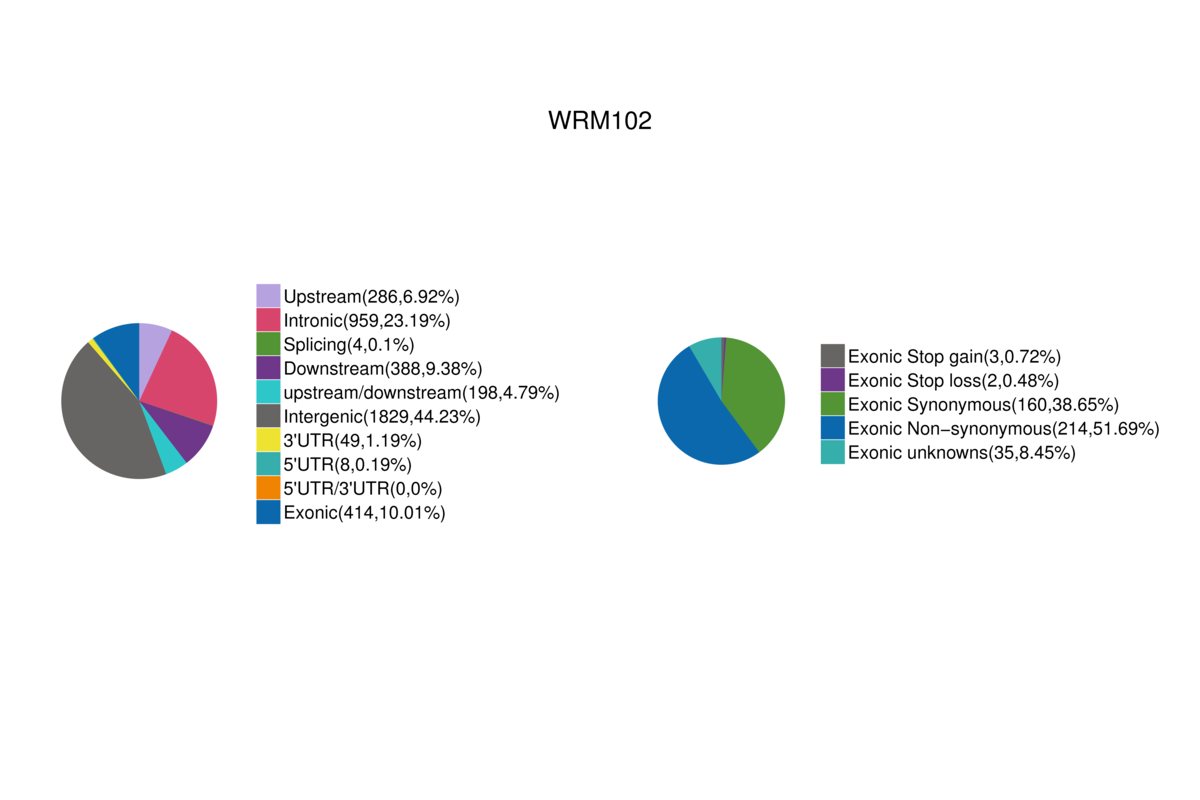

### WRM102.snpDensity.png

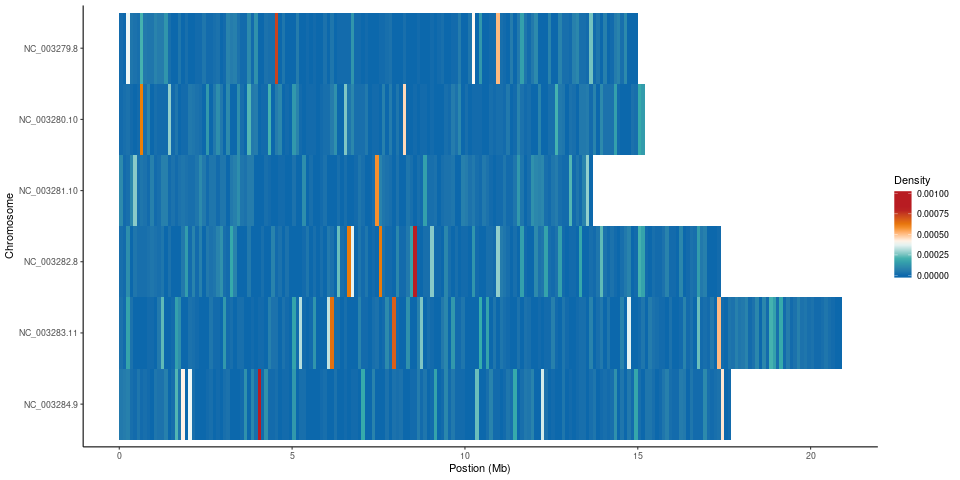

### WRM102.SV.table.png

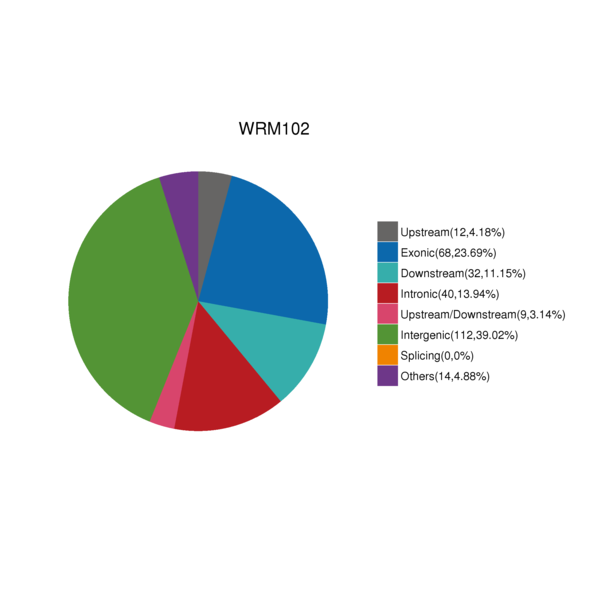

### WRM102_CKDN240021611-1A_22HVGTLT4_L1.Error.png

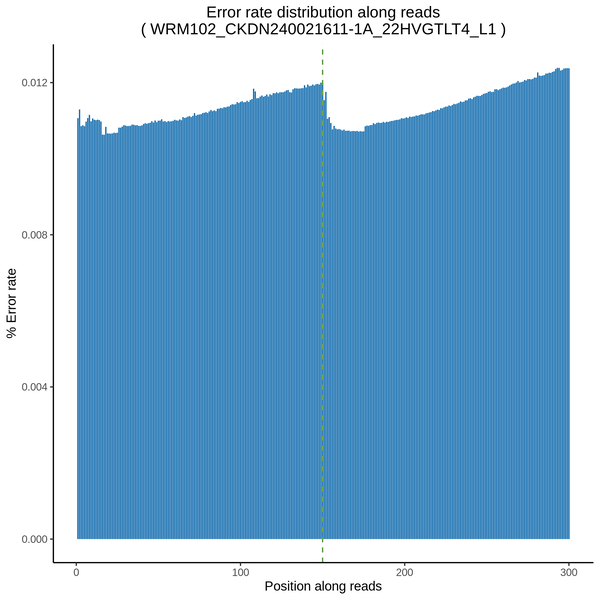

### WRM102_CKDN240021611-1A_22HVGTLT4_L1.GC.png

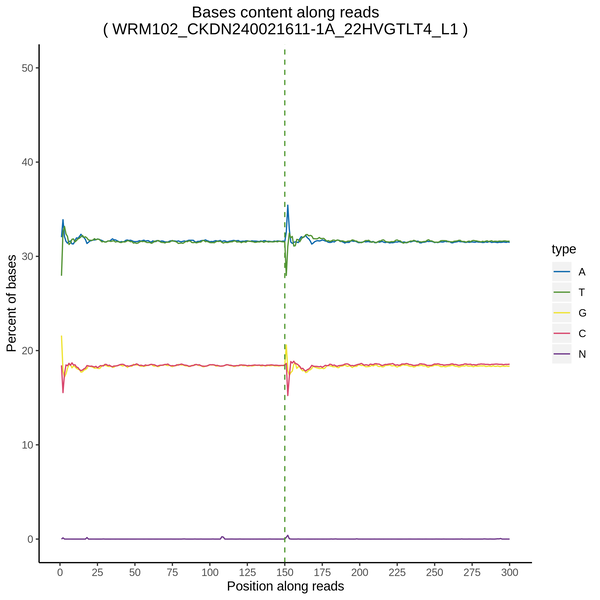

### WRM102_CKDN240021611-1A_22HVGTLT4_L1.pie3d.png

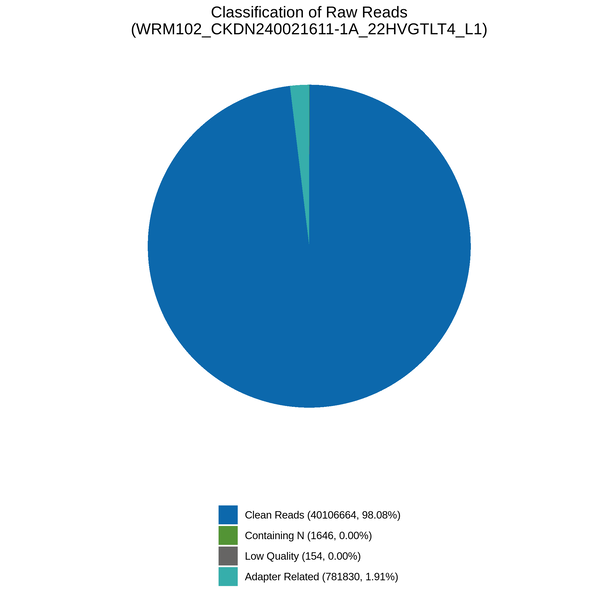

### WRM102_CKDN240021611-1A_22HVGTLT4_L1.QM.png

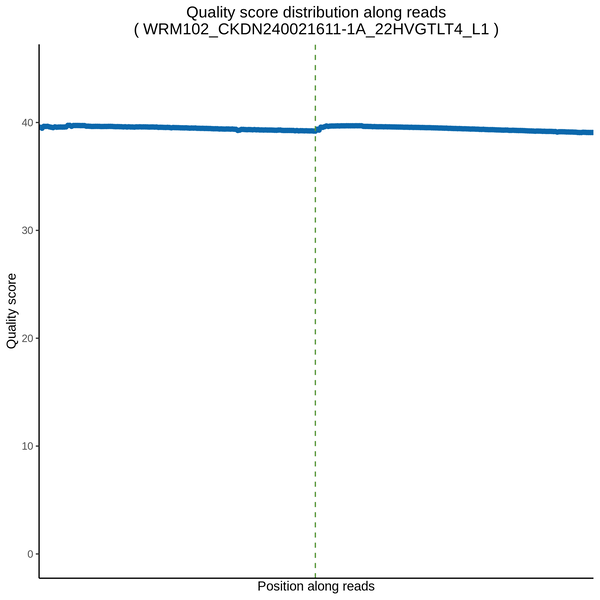

### WRM103.CNV.table.png

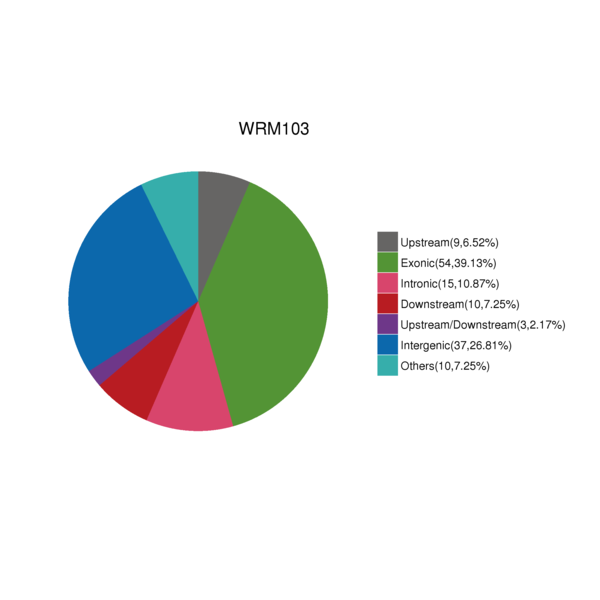

### WRM103.indDensity.png

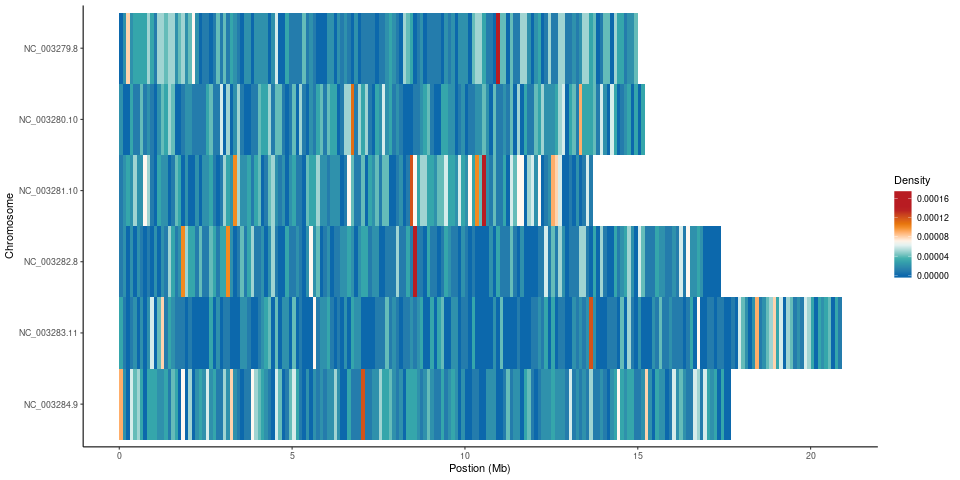

### WRM103.InDel.table.png

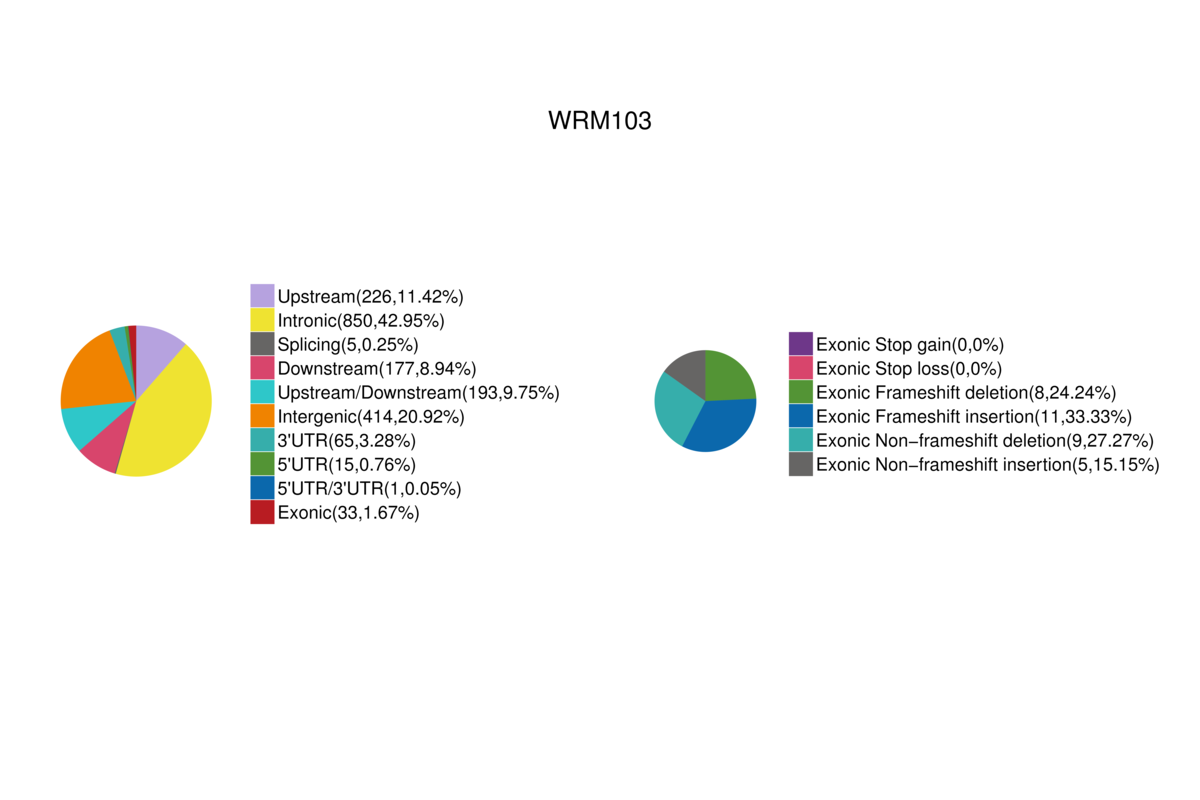

### WRM103.JPEG

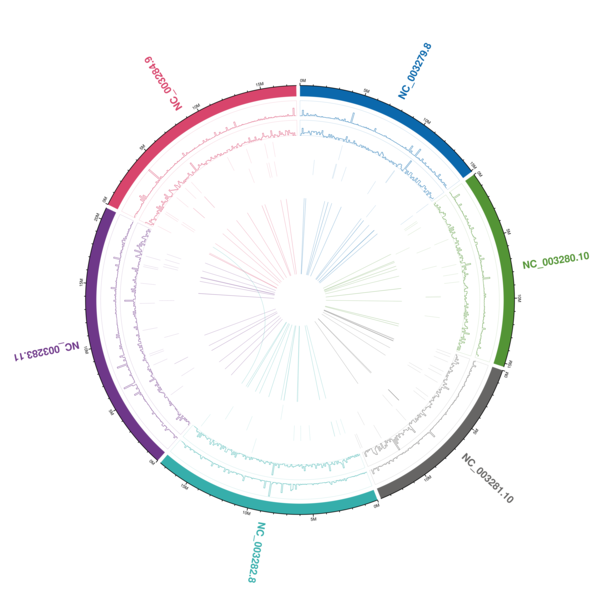

### WRM103.mapbychrdepth.png

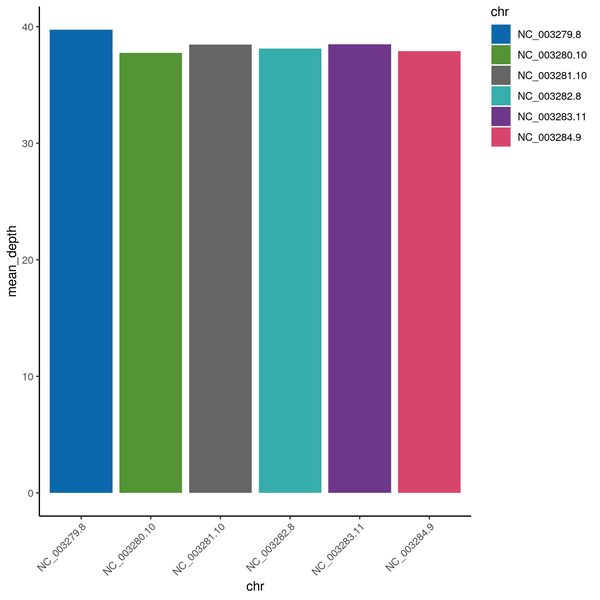

### WRM103.SNP.table.png

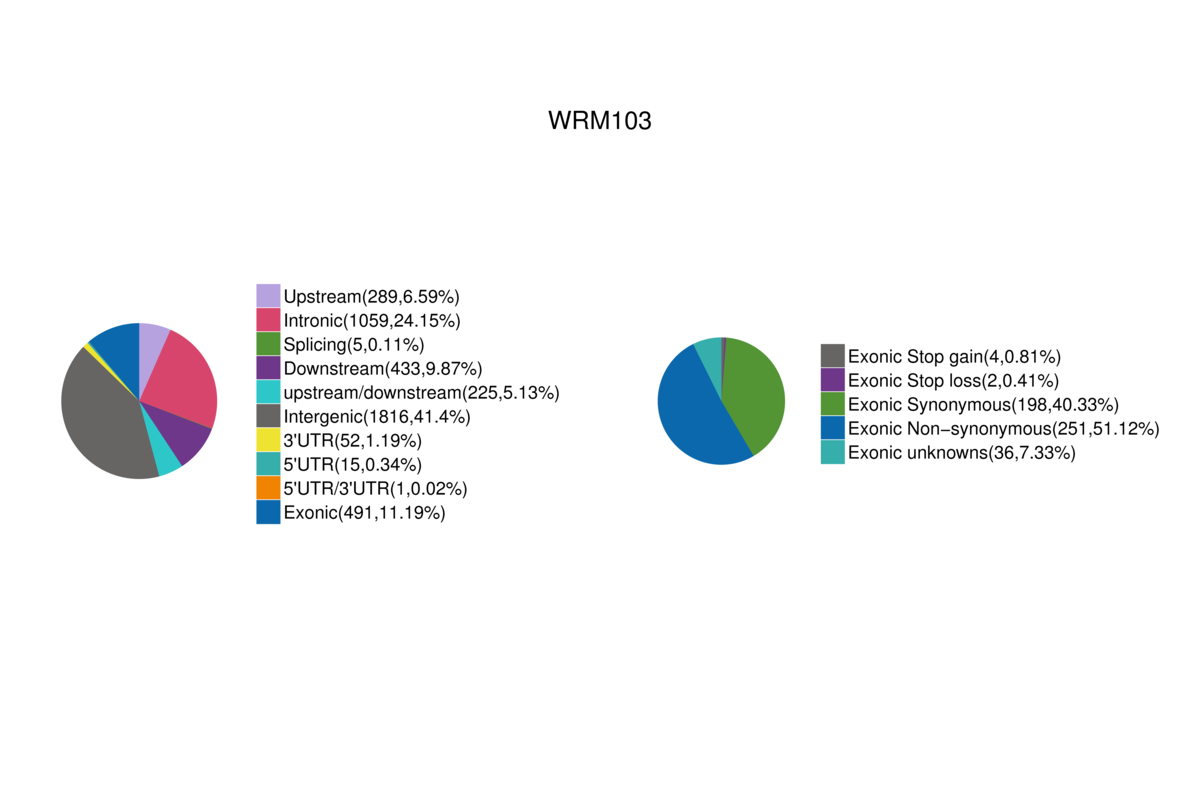

### WRM103.snpDensity.png

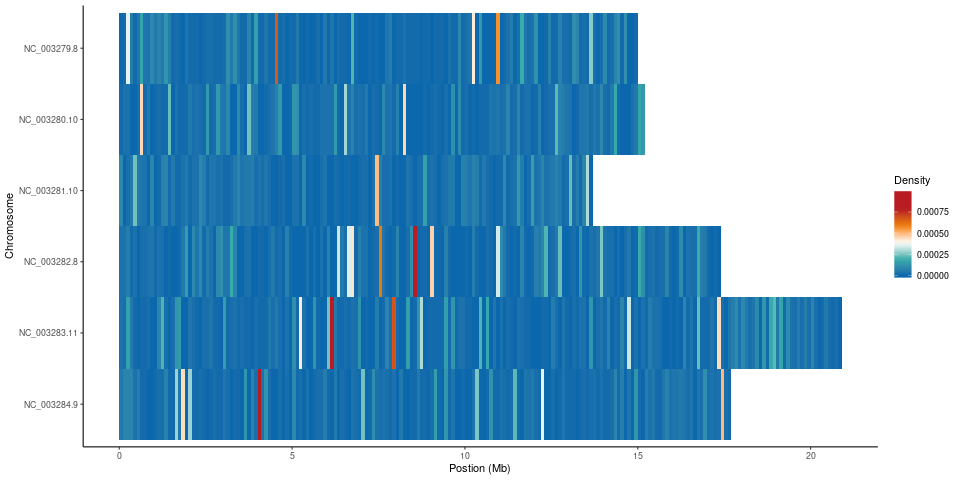

### WRM103.SV.table.png

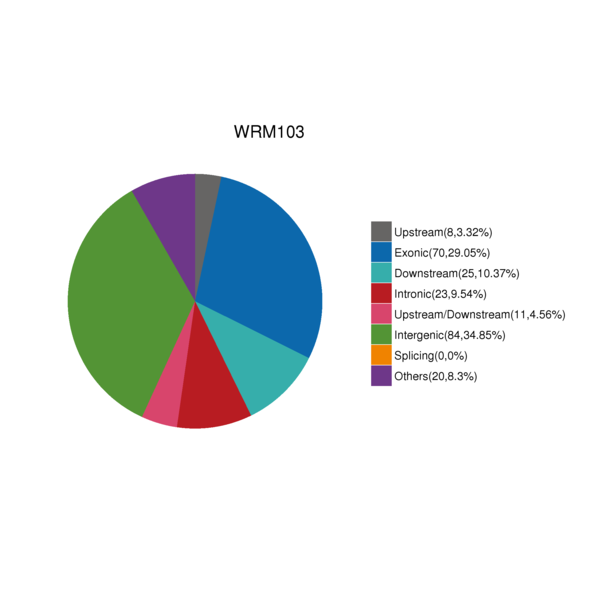

### WRM103_CKDN240021612-1A_22HVGTLT4_L2.Error.png

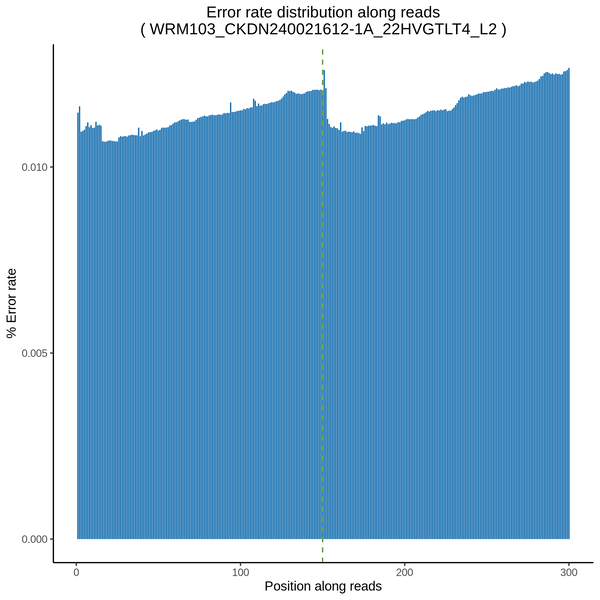

### WRM103_CKDN240021612-1A_22HVGTLT4_L2.GC.JPEG

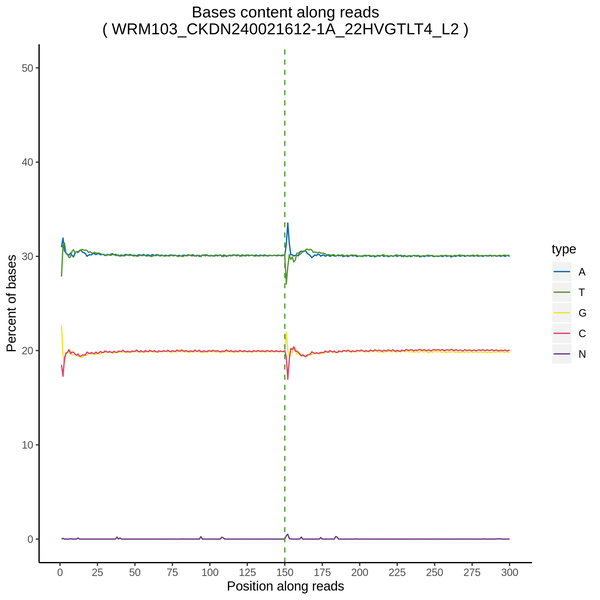

### WRM103_CKDN240021612-1A_22HVGTLT4_L2.pie3d.png

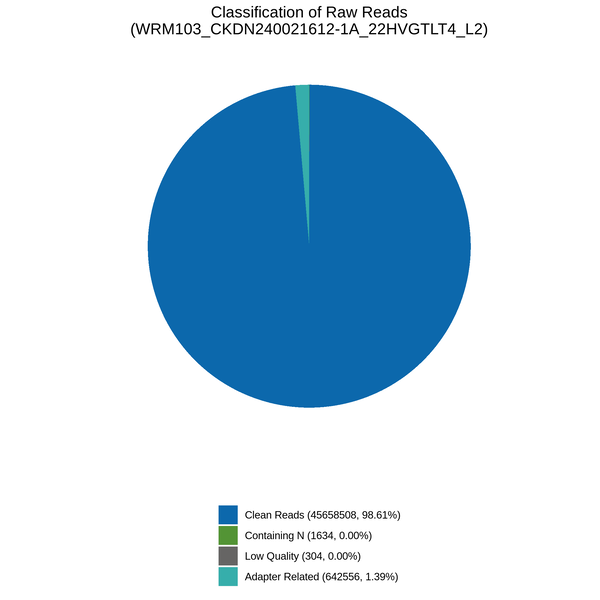

### WRM103_CKDN240021612-1A_22HVGTLT4_L2.QM.png

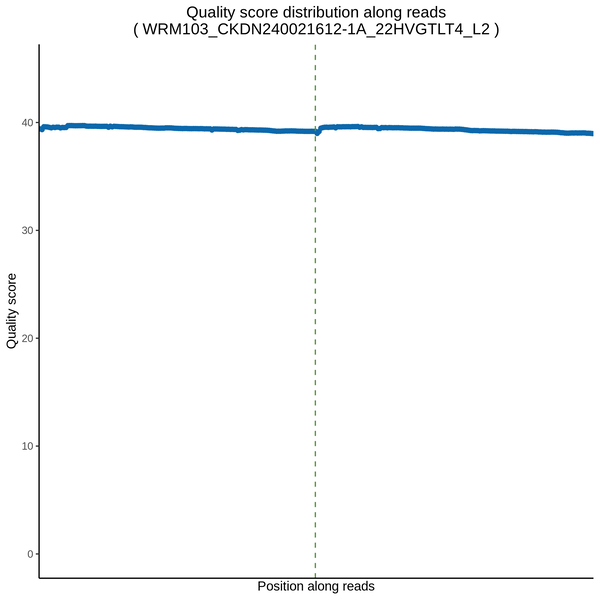
